# Active morphodynamics of intracellular organelles in the trafficking pathway

**DOI:** 10.1101/2024.09.27.615486

**Authors:** S. Alex Rautu, Richard G. Morris, Madan Rao

## Abstract

From the Golgi apparatus to endosomes, organelles in the endomembrane system exhibit complex and varied morphologies that are often related to their function. Such membrane-bound organelles operate far from equilibrium due to directed fluxes of smaller trafficking vesicles; the physical principles governing the emergence and maintenance of these structures have thus remained elusive. By understanding individual fission and fusion events in terms of active mechano-chemical cycles, we show how such trafficking manifests at the hydrodynamic scale, resulting not only in fluxes of material— such as membrane area and encapsulated volume— but also in active stresses that drive momentum transfer between an organelle and its cytosolic environment. Due to the fluid and deformable nature of the bounding membrane, this gives rise to novel physics, coupling nonequilibrium forces to organelle composition, morphology and hydrodynamic flows. We demonstrate how both stable compartment drift and ramified sac-like morphologies, each reminiscent of Golgi-cisternae, emerge naturally from the same underlying nonequilibrium dynamics of fission and fusion.

## I. INTRODUCTION

Non-equilibrium self-assembly is thought to play a crucial role in the emergence and spatial-temporal patterning of intracellular structures [1–5], such as organelles, mitotic spindle [4, 6], centrioles [7], filopodia, flagella and cilia [8, 9]. A key feature of such non-equilibrium structure formation and maintenance is the ATP/GTP driven addition-removal of its subunits; for instance, actin or microtubule based structures form and adapt dynamically via the polymerization-depolymerization of monomers maintained out of equilibrium [10, 11].

In this work, we explore such ideas in the context of the organelles in the endomembrane system [12], such as the Golgi apparatus, lysosomes, endosomes, and vacuoles. Such organelles are often identified by their lipid and protein composition, governed by local material fluxes and synthesis [13–17]. They are also identified by their morphology, which remarkably is maintained in the face of continual non-equilibrium driving from the flux of transport vesicles [16–21]. Indeed, perturbations to vesicular flux have been observed to dramatically alter the shape of Golgi cisternae [22–27], highlighting the role of non-equilibrium driving in determining the steady-state shape.

Due to the complexity and intricate nature of the relevant soft-matter physics, a principled physical description of the non-equilibrium dynamics of such micron-size membrane compartments embedded in the cytosol and subject to vesicular trafficking has been a challenge. Although a few studies have treated the out-of-equilibrium shape changes induced by the incorporation of additional membrane material [28–36], these omit several aspects of fundamental physics underpinning the emergence, maintenance and adaptation of organelle morphologies.

To this end, we incorporate three fundamental features that are common to the dynamics of membrane-bound organelles in the trafficking pathway. First, fission and fusion of transport vesicles are active processes [28, 37], that require specific energy-consuming macromolecules, associated with *non-equilibrium mechanochemical cycles* [38]. Second, each fission and fusion cycle is not only associated with a flux of material that transfers membrane area and encapsulated volume, but also with momentum transfer to/from the ambient fluid [39, 40]. The latter induces active membrane stresses, *i*.*e*. each fission and fusion event is associated with an *active internal force* [28, 37]. Third, membrane shape deformations couple hydrodynamically to the surrounding medium, inducing both membrane and fluid flows, which dynamically change the local membrane composition. By incorporating these features, we find that organelle morphodynamics present a rich context for the physics of non-equilibrium soft matter, involving a complex interplay between fluid dynamics, deformable interfaces, composition, and active stresses arising from the microscopic transduction of chemical energy [41–43].

## II. RESULTS

We derive active hydrodynamic equations for a closed membrane compartment embedded in a viscous solvent subject to active fission and fusion of transport vesicles, in terms of shape, composition and fluid velocity (Supplementary Section I). We find that each fission–fusion event is associated with active force moments whose signs depend on whether it is fission or fusion (Fig. 1). This contributes to a dynamical renormalization of membrane tension, which can potentially lead to shape instabilities, and spontaneous curvature that can lead to segregation of fission and fusion components— our first main result. Linear stability analysis of the derived hydrodynamic equations (Supplementary Section II) about a uniform spherical membrane compartment in the presence of isotropic vesicular flux leads to our second result— a spontaneous drift instability of the compartment beyond a threshold activity (Fig. 2). Such motion can be either anterograde or retrograde relative to the incident vesicular flux, depending on the magnitude of the active force moment and the relative fission and fusion rates (Fig. 3). Indeed the active stresses control a combined drift-shape instability, leading to moving spheres, prolates, and oblates. Importantly, the drift instability does not require a fore-aft symmetry breaking in shape and can be driven by a vectorial mass flux. This has important implications for cisternal progression [16, 17, 44, 45]. In addition, the compartment exhibits higher-order shape instabilities in the form of flattenedsacs or tubular ramified structures (Fig. 2 & 3). Such cisternal morphologies, commonly exhibited in a variety of cell types [17, 46, 47], constitutes our third result. By extending the linear stability analysis to include leading order nonlinearities (Supplementary Section III), we derive amplitude equations for the shape and composition to second-order in the fields. A weakly nonlinear analysis [48] shows a stable drift velocity of the organelle in a regime of the parameter space (Fig. 4). We emphasise that these results are contingent on a hydrodynamic description of non-equilibrium fission and fusion processes and will not arise from a purely kinetic treatment.

**FIG. 1.**
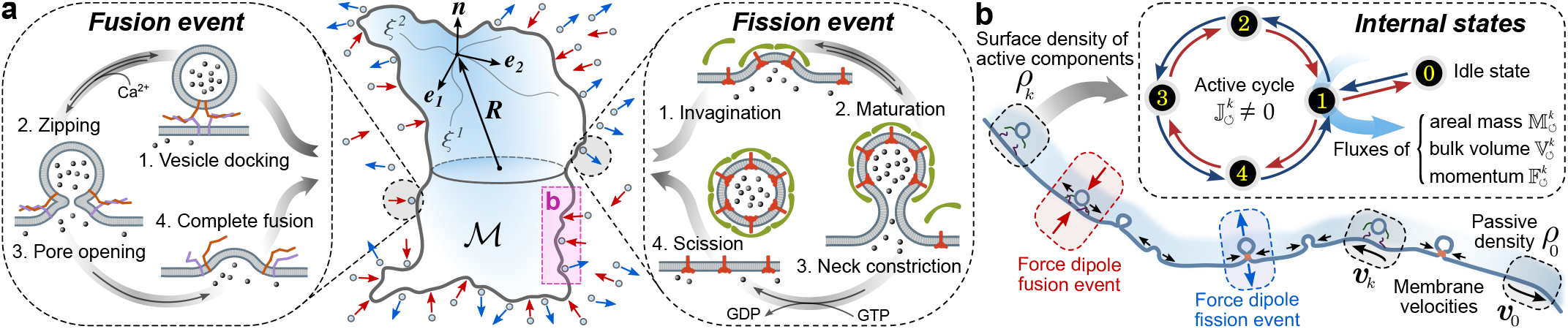
Organelle subject to active fission and fusion of transport vesicles. (a) A subcellular membrane organelle that is continually subject to the fission and fusion of small transport vesicles (small circles). The blue arrows indicate outgoing vesicles (fission), while the red arrows label incoming vesicles (fusion). Fission and fusion are independent processes arising from detailed balance violating biochemical reaction cycles (insets depict 4 states). (b) Cross-section of a small patch of the deformable membrane ℳ, showing the boundary layer at the membrane in which numerous fission and fusion events may occur. On large length and time scales, these events can be described by local densities *ρ*_*k*_, *ρ*_0_ and velocities ***v***_*k*_, ***v***_0_ of active (fissogens and fusogens) and passive components. Each cycle, when averaged over their internal states, carries a nonzero loop current that in turn drives local fluxes of area, volume, and momentum onto the membrane organelle; see inset. Integrated over a cycle, the fission and fusion events can be associated with extensile and contractile force dipoles normal to the membrane.

**FIG. 2.**
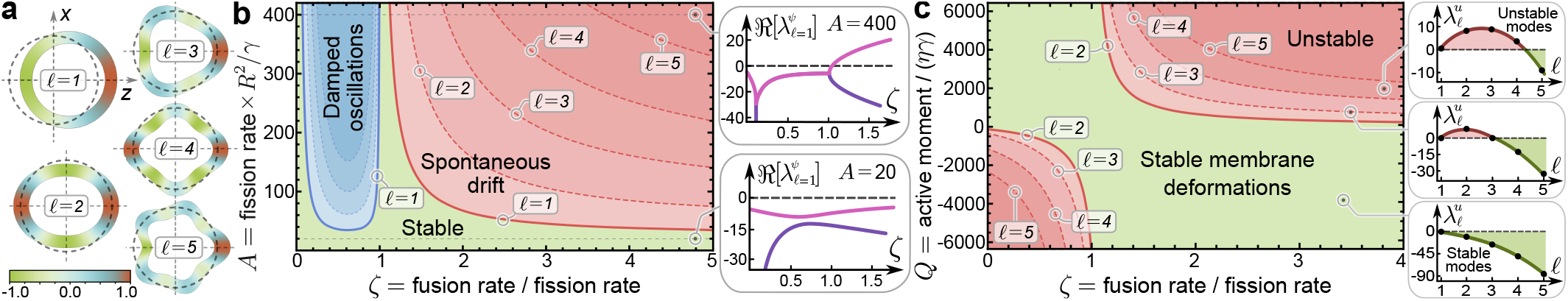
Generic instabilities in the composition and shape of membrane organelle. (a) Normalized spherical harmonics *Y*_*𝓁*,*m*_(*θ, Φ*) for *m* = 0 and *Φ* = 0, illustrating the spherical decomposition of surface densities (color map) and shape distortions from a homogeneous sphere (dashed circle). (b) Stability diagram corresponding to composition perturbations for each spherical harmonic, in terms of the fission and fusion rates, made dimensionless by the surface diffusion time *R*^2^*/γ*. This stability diagram is independent of the active moment *Q*. The onset of spontaneous drift instability (*𝓁* = 1) is marked by the solid red line, while the boundaries of the higher mode (*𝓁 >* 1) shape instabilities are shown as red dashed lines. Both green and blue regions correspond to the linearly stable regime, with the latter showing stable damped oscillations (each mode is shown by the dotted lines in the blue region). Insets show the growth rates 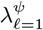 of composition at two fixed values of *A* as a function of *ζ*. (c) Stability diagram corresponding to shape perturbations, with growth rates 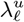 as a function of *𝓁* at fixed values of *ζ*, dimensionless active moment *Q*, and dimensionless bending rigidity 𝒦= *κ/*(*Rγη*) = 10 (see insets). Red lines show the onset of shape instabilities for the different *𝓁* modes. This stability diagram is independent of the dimensionless fission rate *A*.

**FIG. 3.**
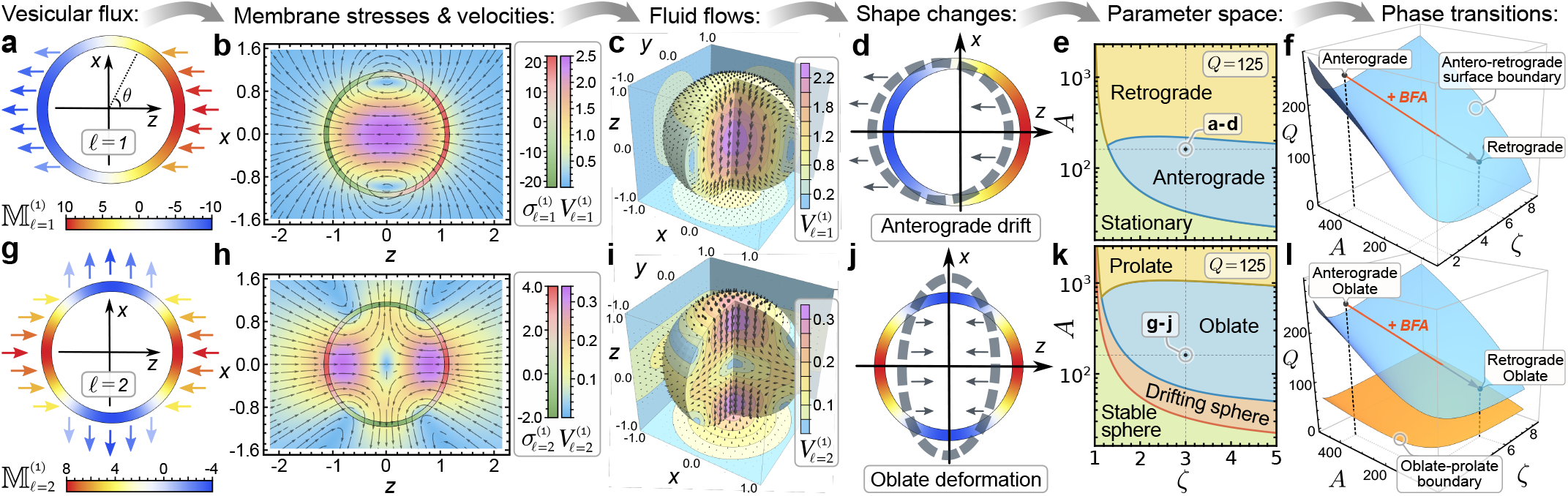
Vesicular flux drives shape changes by inducing membrane stresses and fluid flows. An initial perturbation about the homogeneous sphere, described by the net mass flux 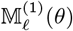, drives instabilities in composition at different *𝓁*, which induce membrane stresses and fluid flows, further driving shape changes. Here, we choose *ζ* = 3 and *A* = 180, such that only *𝓁* = 1 and *𝓁* = 2 modes are unstable, see Fig. 2(b), at *Q* = 125 and 𝒦 = 10. (a) Angular profile of 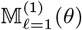 with arrows denoting its directionality. (b) Vesicular flux induces a normal surface stress 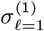 that results in membrane and bulk fluid flows (streamlines) with speed 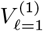. Both (a) and (b) represent the cross-section *y* = 0. (c) Three-dimensional bulk flows for *l* = 1 mode. Far-field flows decay as 1*/r*^2^ from the centre of the organelle, see Supplementary Section II.I. (d) Membrane flows incur shape distortions, and lead, for this choice of parameters, to an anterograde drift, moving away from the source of vesicular flux. (e) Phase diagram in parameters *A* and *ζ*, showing the transitions between stationary, anterograde and retrograde drift at fixed *Q*. The point in the phase diagram corresponding to (a)–(d) is shown. Reducing *Q* moves the anterograde–retrograde transition line to lower values of *A*, see Supplementary Figures S.14 and S.15. (f) Phase diagram in *A*–*ζ*–*Q* showing the boundary surface between anterograde and retrograde regimes. Starting from parameter values corresponding to an anterograde phase, a three-fold decrease in the fission flux 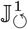 corresponds to proportionate changes in *ζ, A*, and *Q* (see Supplementary Section I.F), driving a transition to a retrograde drift (red arrow). This is consistent with *in vivo* experiments using Brefeldin A (BFA) treatment. (g) Angular profile of 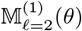. (h) Normal stress 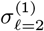 induced by 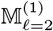 and their associated fluid flows 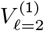. Both (g) and represent the cross-section *y* = 0. (h) Three-dimensional bulk flows for *𝓁* = 2, which decays as 1*/r*^2^ in the far-field. (j) For this choice parameters, the *𝓁* = 2 flows lead to an oblate distortion. (k) Phase diagram in *A* and *ζ* at fixed *Q* shows transitions between sphere, prolate and oblate. The red line demarcates the transition between a stationary and a drifting sphere, see Supplementary Figure S.14 and S.15. The point corresponding to (g)–(h) is indicated. (l) Phase diagram in *A*–*ζ*–*Q* showing the boundary surfaces between anterograde and retrograde drifts (blue) and oblate and prolate shapes (orange). As in (f), a change in the fission rate can drive a transition from anterograde oblate to retrograde oblate (or even to a retrograde prolate for sufficiently large changes). All the plots (a)–(d) and (g)–(j) are computed at time *t* = 0.1 *R*^2^*/γ*, with initial 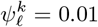.

**FIG. 4.**
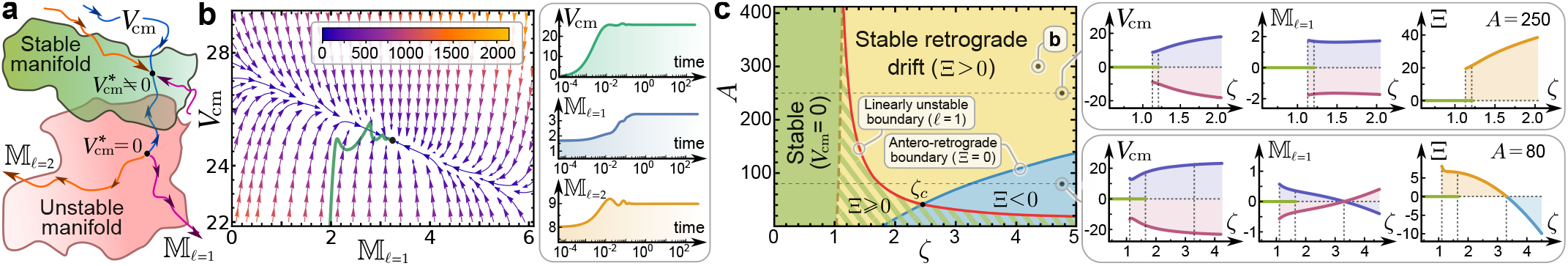
Nonlinear analysis shows stable anterograde and retrograde motion of organelle. (a) Schematic of stable and unstable manifolds in the space of mass fluxes 𝕄_*𝓁*_ and shape amplitudes for modes *𝓁* = 0, 1, 2, …, subject to the constraint that vesicular flux does not increase the net membrane mass of organelle. This illustrates that the stationary state 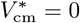 lies within an unstable manifold. Certain trajectories can bring the system onto a stable manifold, with *V*_cm_ ≠ 0. (b) Restricting to the lowest nonlinearities in the governing equations for composition, together with a truncation up to *𝓁* = 2 mode, yields stable fixed points with *V*_cm_ ≠ 0. We project all amplitudes at the fixed point except for 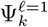, which allows us to construct a plot of *V*_cm_ versus 𝕄_*𝓁*=1_, converging to a stable fixed point with *V*_cm_ ≠ 0. The green curve gives the projection onto the *V*_cm_– 𝕄_*𝓁*=1_ plane, of the trajectory found by solving the dynamical equations, starting from initial conditions. Inset shows the time evolution of the amplitudes, starting from *V*_cm_ = 0 and with the other mass fluxes chosen to be in the basin of attraction of the stable fixed point. Here, *ζ* = 4, *A* = 300, and *Q* = − 8000, and time is in units of *R*^2^*/γ*. (c) Phase diagram in *A* and *ζ*, at fixed *Q* = − 8000. Red curve is the linearly unstable boundary, above which the organelle spontaneously drifts. We find regions of stable retrograde (yellow) and anterograde (blue) drift, which extend to the hatched region where anterograde and retrograde solutions coexist with the homogeneous stationary solution (*V*_cm_ = 0). This phase coexistence shows up in the bifurcation plots of *V*_cm_, 𝕄_*𝓁*=1_, and their product Ξ, as a function of *ζ* at fixed *A* and *Q*. The sign of Ξ decides whether the motion is anterograde (Ξ *<* 0) or retrograde (Ξ *>* 0). The bifurcation plots show that the linearly unstable line (red) in the phase diagram corresponds to a subcritical bifurcation (first-order phase boundary), whereas the antero-retrograde line (blue), with Ξ = 0, is a second-order phase boundary. The point *ζ*_*c*_, where these two lines intersect, is a tricritical point.

### A. Fission and fusion processes are active mechanochemical pumps

Our coarse-grained hydrodynamic equations for membrane composition and shape (Supplementary Section I) rely on a natural separation of spatial–temporal scales between active mechanochemistry and membrane mechanics. The timescales of vesicle fission and fusion events (0.1 – 1 s) [31, 49–51] are smaller than the passive relaxation times of the membrane, such as the lateral diffusion time (*∼*100 s) and the viscous shape relaxation (*∼* 10 s), see Supplementary Section I.A. The lateral size of organelle (*∼*5 *µm*) is much larger than the size of transport vesicles (*∼*50 nm). This allows us to treat the active events as being local in time and space. In cases where the waiting time between fission and fusion events is much longer than the membrane relaxation times, the membrane adiabatically equilibrates at a different area and volume between each fission and fusion event.

The composition fields on the membrane consist of passive membrane components (lipids and proteins) and the protein machinery associated with fission and fusion, referred throughout as *fissogens* and *fusogens*. Locally, these compositional fields evolve via recruitment, active mechanochemistry, and dynamics along the membrane. We describe the internal mechanochemical kinetics as a *fast* biochemical Markov cycle (Supplementary Section I.A and I.E) with many intermediate states, see Fig. 1(a). The transitions between these states do not respect detailed balance, as many of these reactions are catalysed by energy-consuming enzymes. For simplicity, we model the biochemical cycles as first-order kinetics with constant effective rates. This ignores potential feedback control of the reaction rates through membrane elasticity, such as surface tension, or limiting levels of fissogens and fusogens [35, 52, 53]. Violation of detailed balance leads to a finite non-equilibrium loop current associated with each cycle, denoted by 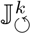 where *k* = 1, 2 labels fission and fusion, respectively. The completion of a fission (fusion) cycle leads to the removal (addition) of membrane area, and volume comprising both solvent and solutes as cargo. Thus, fissogens and fusogens are *active pumps* [37, 54, 55], for membrane area and lumenal volume; the strength of these fluxes depend on the non-equilibrium currents 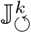 and local area fractions, see Fig. 1(b).

In addition, each cycle drives a flux of momentum exchanged with the surrounding medium, which arises due to internal active forces within a small boundary layer normal to the membrane [39, 40, 56], whose scale is set by the size of transport vesicles. Integrating over the cycle time allows us to coarse-grain the dynamical equations over the boundary layer, projecting the hydrodynamic fields onto the two-dimensional membrane. Under such coarse-graining, the momentum flux due to fission and fusion events in the boundary layer leads to a local active membrane stress (see Supplementary Section I.H).

### B. Active hydrodynamics of membranes subject to fission and fusion

The density of active membrane components is given by *ρ*_*k*_, where *k* = 1, 2 denotes fissogens and fusogens, respectively, whilst the density of passive membrane components is *ρ*_0_ (Fig. 1). In our coarse-grained description, the membrane is incompressible, with a constant total density *ρ* = *ρ*_0_+Σ_*k*_*ρ*_*k*_. By defining the local fractions Φ_0_ = *ρ*_0_*/ρ* and Φ_*k*_ = *ρ*_*k*_*/ρ* and assuming a dilute concentration (Φ_*k*_ ≪ Φ_0_), the free-energy of a close membrane ℳ is ℱ = ∫_ℳ_ *F* d*S* (Supplementary Section I.D), with

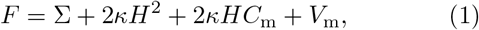

where Σ is the surface tension, *κ* is the bending rigidity, *H* is the mean curvature, and the spontaneous curvature *C*_m_ = 𝒞_0_Φ_0_ + Σ_*k*_ 𝒞_*k*_Φ_*k*_, with 𝒞_0_ and 𝒞_*k*_ as curvature coupling strengths. The composition free-energy density in the dilute limit, 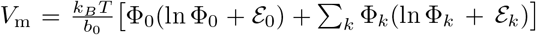, contains entropy of mixing and chemical potentials ε_0_ and ε_*k*_ in units of thermal energy *k*_*B*_*T*, and *b*_0_ is a coarse-grained area.

The evolution of membrane components is determined by passive forces, obtained from *ℱ*, and active fluxes associated with mechanochemical cycles (Fig. 1):

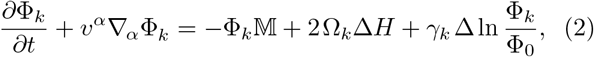

where *γ*_*k*_ is surface diffusivity, Ω_*k*_ is curvature coupling strength, Δ is Laplace–Beltrami operator, and ∇_*α*_ is the covariant derivative, with summation convention implied. The active vesicular flux 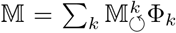, where 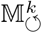 is the membrane area transferred during the fission and fusion cycles (see Supplementary Section I.F). Due to membrane incompressibility, 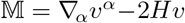, with the right-hand-side being the surface divergence of membrane ve-locity 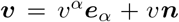, which is decomposed along the tangent basis ***e***_*α*_ and the outwards unit normal ***n***.

The evolution of membrane shape is determined by a combination of passive elastic forces, active stresses associated with fission and fusion cycles, and external forces arising from the ambient fluids. By conservation of linear and angular momentum, the force balance at the membrane is given by (Supplementary Section I.H):

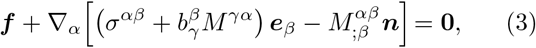

where the external force ***f*** arises from stresses imposed by the surrounding fluids (Supplementary Section I.H), while 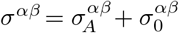 is the total symmetrized in-plane stress tensor, and 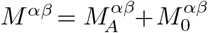 is the total bending moment tensor. The passive elastic stresses are given by

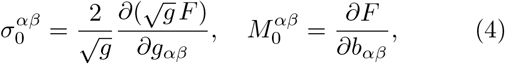

where *g*_*αβ*_ and *b*_*αβ*_ are the metric and curvature tensors, respectively, and *g* = det[*g*_*αβ*_]. The active stresses due to fission and fusion at the membrane interface with the outer fluid are derived by associating with each cycle a force density 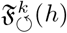, where *h* is the normal height away from the membrane. By momentum conservation, the monopole contribution must vanish, 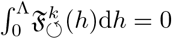, where Λ is the boundary layer thickness. A moment expansion of the force density 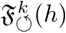, to second order in *h*, gives the active symmetric in-plane stress and the active bending moment stress (Supplementary Section I.G):

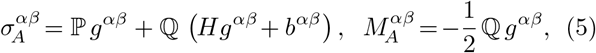

where 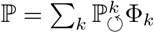 and 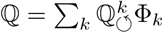 are the first and second moments of the active force distributions, with variables 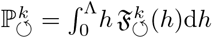 and 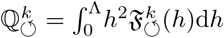.

The inner (−) and outer (+) ambient fluids satisfy the incompressible Stokes equations [57]: **∇** *·* **Ω**_***±***_ = **0** and **∇** *·* ***V***_***±***_ = 0, where fluid stress tensors **Ω**_***±***_ = −*p*_***±***_ ***I*** + *η* [(**∇*V***_***±***_) +(**∇*V***_***±***_)^T^], *p*_***±***_ and ***V***_***±***_ are the corresponding fluid pressure and velocity. For simplicity we choose the inner and outer fluids to have the same shear viscosity *η*. The force ***f*** in Eq. (3) is given by the stress jump across the membrane interface: ***f*** = ( **Ω**_+_ − **Ω**_**−**_) *·* ***n*** | _ℳ_ (Supplementary Section I.I). Additionally, the Stokes equations must satisfy velocity matching conditions:

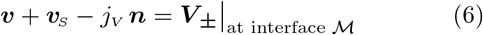

where the slip velocity 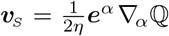 is a consequence of flows within the boundary layer induced by active surface stresses (Supplementary Section I.G and I.I). The boundary condition Eq. (6) ensures that we maintain the same far field flows as in a semi-microscopic description, in which the active stresses at membrane enter the bulk fluid equations as sources [56, 58–60]. The volume flux *j*_*V*_ accounts for transport across the interface through passive osmosis and active volume exchange from fission and fusion (Supplementary Section I.K).

The composition dynamics (2), force–balance (3), incompressible Stokes equations, and boundary condition (6) form a closed set of equations that describe the organelle morphodynamics, see flowchart in Supplementary Figure S.1. Even before solving the governing equations, we can establish a few key results. By collecting the coefficients linear in mean curvature from Eq. (3) along the normal direction, we find an activity renormalized surface tension that depends explicitly on the first moment 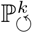 of the active force distribution (see Supplementary Equations S.123). With the simplest active force realisation as a force-dipole, the sign of the active moments 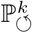 and 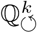 are fixed by 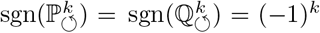 (see Supplementary Equation S.109). By this sign convention, we find an increase (decrease) of surface tension during fission (fusion). Thus excess fusion can drive the surface tension to negative values, leading to shape instabilities consistent with earlier theoretical work [28, 29] and *in vitro* membrane experiments [61, 62]. Likewise, we find an activity renormalized local spontaneous curvature (see Supplementary Equations S.124) that depends explicitly on the second moment 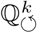. The same sign convention for 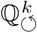 implies that local spontaneous curvature increases (decreases) during fission (fusion), which can drive segregation of fissogens to highly curved regions, as observed at the Golgi cisternal rims [63]. As described in Supplementary Section I.G, the signs of 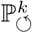 and 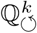 for more elaborate active force realisations can be independent in general. We emphasise that both the activity renormalized surface tension and spontaneous curvature have contributions arising from the active force moments, in addition to the expected contribution from the mass flux of fissogens and fusogens.

### C. Spontaneous drift and shape instabilities

The closed set of nonlinear equations can be solved using a suitable numerical scheme (Supplementary Figure S.1), such as boundary integral methods [64]. Instead, we find it instructive to study the dynamics perturbatively about the homogeneous spherical non-equilibrium steady-state (Supplementary Section II). The shape deformation 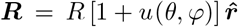, where 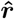 is the radial unit vector, and *u*(*θ, Φ*) is a small radial distortion about the undeformed sphere of radius *R*, with *θ* and *ϕ* as the inclination and azimuthal angles. We perturb the composition fields 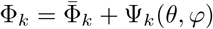 about the homogeneous value 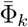, *maintaining the total mass fixed*. The velocity fields of the fluid and membrane are expanded to linear order, with Eq. (6) appearing as a boundary condition on the undeformed sphere. Lamb’s solution to the Stokes equations [39] allows us to solve for the leading order fluid flows, 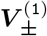, in terms of radial velocity, 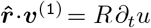, and the surface divergence, 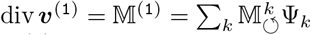. By expanding Eqs. (2) and (3) in spherical harmonics, we derive the equations for the shape *u*_*𝓁*_, composition 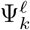, and surface tension Σ_*𝓁*_, corresponding to the *𝓁*-th spherical harmonic mode, as shown in Fig. 2(a). For further details, see Supplementary Section II.A–F.

This approach allows us to study the instabilities of membrane composition and shape (Supplementary Section II.G), arising from spatial modulations in composition and coupling to hydrodynamic flows induced by membrane stresses (Fig. 2 and 3). The far-field fluid flows decay as 1*/r*^2^ (Supplementary Section II.I) from the centre of the organelle, see Fig. 3(c) and (i). Our key result is the spontaneous drift instability of the organelle (*𝓁* = 1), along with higher–order shape instabilities (*𝓁* ≥ 2) in the form of flattened-sacs and tubular ramified structures. We construct stability diagrams by tuning the imbalance 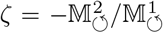 between fusion and fission rates, the dimensionless fission rate 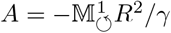, and the dimensionless active moment 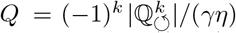, at fixed dimensionless bending rigidity 𝒦= κ*/*(*Rγη*), as shown in Fig. 2(b) and (c). For simplicity, we set the magnitude of 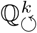 to be independent of *k*, the diffusivity *γ*_*k*_ = *γ*, and neglect spontaneous curvature terms and volume fluxes.

The spontaneous drift instability appears generically when *ζ >* 1 and 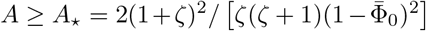, where 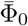 is the areal fraction of passive components. The critical value *A*_*\**_ represents the organelle size *R* at which surface diffusion is not fast enough to homogenize the composition relative to the vesicular flux. To understand the spontaneous organelle drift, we solve the linear stability equations for initial data in the amplitudes 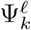 and *u*_*𝓁*_, to obtain membrane stresses and fluid flows to linear order. Fig. 3 highlights this interplay between vesicular flux and shape deformations via the induced fluid flows and membrane stresses.

To linear order, the magnitude of the drift velocity is given by 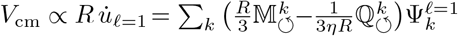. The direction in which the organelle moves with respect to the incident flux 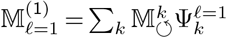 is set by an interplay between mass transfer and active stresses; the organelle motion is *retrograde* (*anterograde*) if it is towards (away from) the source of vesicular flux. Fig. 3(e) shows transitions from stationary →anterograde → retrograde as we increase *A*, keeping *ζ* and *Q* fixed. Similarly, the higher-order (*𝓁* ≥ 2) shape instabilities are driven by the imbalance *ζ* and active moment *Q*, as shown in Fig. 2(c), which shows the critical lines above which instabilities occur for each *𝓁*-th mode. To find the shape corresponding to the ellipsoidal mode *𝓁* = 2, we solve the linear equations for *u*_*𝓁*=2_ and 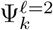; Fig. 3(k) shows transitions from stationary sphere → moving sphere →moving oblate → moving prolate as we increase *A* by holding *ζ* and *Q* fixed.

We highlight two crucial aspects that emerge from our analysis. First we note that the oblate organelle moves in an anterograde fashion, reminiscent of the Golgi cisternal progression [16, 17]. Second, we emphasise that varying the fission loop current 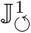 leads to proportionate changes in *ζ, A*, and *Q*. Therefore, by perturbing the organelle through a reduction in the fission current, the initial anterograde drift (or even stationary) could transition to a retrograde drift, see Fig. 3(f) and (l). At the same time, the activity–renormalized surface tension may be driven to negative values, leading to a potential break-up of the cisterna [28, 61, 62]. This is consistent with Brefeldin A experiments [19, 65, 66] that inhibit vesicle coat formation, hence fission, and is observed to cause retrograde motion of the Golgi cisterna, its fragmentation and ultimately its fusion with the endoplasmic reticulum. This is also consistent with the observed fragmentation of the Golgi cisternae in fibroblast cells upon loss of adhesion from a substrate [27]. To the best of our knowledge, this is the first theoretical justification from first principles that may explain these experimental observations.

### D. Weakly nonlinear analysis reveals a stable fixed point with finite organelle velocity

The generic drift and shape instabilities predicted by our linear stability analysis shows that the stationary state (a homogeneous sphere with *V*_cm_ = 0) lies within an unstable manifold, see Fig. 4(a). To find the shape parameters and centre-of-mass velocity at the non-equilibrium steady-state, we perform a stable manifold analysis [48] on the system of nonlinear amplitude equations in shape *u* and composition Ψ_*k*_ (see Supplementary Section III). To carry out this analysis, we restrict to a lower dimensional phase space and retain only quadratic terms in the amplitudes of *u* and Ψ_*k*_. We truncate the resulting mode–coupling equations to *𝓁* = 2 and take only the *m* = 0 component of the spherical harmonic modes. For simplicity, we neglect the nonlinearities associated with the shape amplitude equations (Supplementary Section III.A and B). This allows us to write down a dynamical equation for the drift *V*_cm_ and study its dynamical flows (Supplementary Section III.C) together with the nonlinear equations for composition 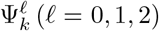. We seek steady-state solutions that conserve the total membrane area of the organelle, enforced here as a constraint. Fig. 4(b) shows a section of the phase portrait in the plane of *V*_cm_ and membrane flux 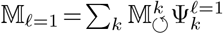 stable fixed point with *V*_cm_ ≠ 0 is approached by moving along a centre manifold [48], see Supplementary Section III.D. This has implications for the time evolution of the amplitudes starting from an initial stationary configuration, see inset in Fig. 4(b).

We find a *non-equilibrium phase diagram* of stable retrograde and anterograde solutions in terms of *A* and *ζ*, at fixed *Q* [67]; see Fig. 4(c) and Supplementary Section III.D. The product of *V*_cm_ and 𝕄_*𝓁*=1_ provides us with an order parameter Ξ, whose sign distinguishes between the two types of motion, as shown by the bifurcation plots of Fig. 4(c). The linearly unstable line is a subcritical bifurcation (first-order transition) from a stationary to a moving organelle. The hatched region shows coexistence between the stationary and the moving phases, as is reflected in the bifurcation plots in the insets of Fig. 4(c). The transition from anterograde to retrograde motion is smooth, corresponding to a second-order phase boundary at Ξ = 0. The first and second order lines meet at a tricritical point (*ζ*_*c*_, *A*_*c*_).

Thus, under vesicular trafficking, an organelle acts a force–free swimmer, that moves either retrograde or anterograde with respect to the incident flux.

## III. DISCUSSION, CAVEATS AND OUTLOOK

In this paper, we have provided a systematic analysis of organelle shapes subjected to non-equilibrium forces and hydrodynamic flows driven by energy-consuming active fission and fusion. In doing so, our work takes a first step towards a quantitative study of the non-equilibrium mechanics of the shapes of membrane-bound organelles, such as the Golgi apparatus, in the trafficking pathways. Here, we focus on the issue of maintenance and instabilities of the cisternal shape under continuous trafficking, leaving the crucial question of *de novo* non-equilibrium assembly of Golgi cisternae for later work. Thus, the perturbations that drive shape instabilities are such that there is no change in net membrane area from the initial steady-state configuration. We have shown that the fluid, deformable nature of the membrane provides a coupling between composition and shape, which drives spontaneous drift and shape instabilities; nonlinearities in the fields result in stable organelle motion that depends on the levels of activity and mass fluxes.

Over the years, there have been several proposed models of Golgi trafficking, such as vesicle transport, cisternal progression and their variants, often taken to be “diametrically” opposed [16, 68]. Recent studies have provided strong evidence for cisternal progression in the budding yeast [44, 45]. An outstanding question in the field has been: *“What drives cisternal progression?”* [16]; our work may provide an answer to this, and identifies, for the firsttime, non-equilbrium fission-fusion as a driving force for cisternal movement. An important result of our study, as shown in Fig. 4(c), is that cisternal progression and vesicle transport can be realised as the moving and stationary phases, respectively, arising from the same underlying non-equilibrium dynamics of fission and fusion.

Moreover, the ramified sac-like morphology of stacked Golgi cisternae emerges as a natural outcome of the higher-order shape instabilities driven by nonequilibrium stresses induced by fission and fusion. Indeed, an implication of our study is that the Golgi morphology and dynamics is intimately associated with its primary function, namely glycosylation, which is dependent on the nonequilibrium trafficking of cargo vesicles [13, 69]. We emphasise that the key results highlighted here could not have been derived from a purely kinetic approach and require a careful treatment of active mechanics and hydrodynamics, as attempted here. For instance, in the absence of active force moments ℙ and ℚ, both the anterograde–retrograde drift and oblate–prolate shape transitions do not occur.

We now list the caveats regarding the present work and suggest ways of addressing them. One major short-coming is that the transition rates in the fission–fusion cycles have been taken to be independent of state of the organelle, such as surface tension and local composition or the availability of fissogens and fusogens. This mechano-chemical feedback is required for the maintenance of homeostatic control of the organelle size.

Another important cellular aspect that we have ignored is the influence of the cytoskeleton. The presence of the cytoskeletal meshwork will result in a screening of hydrodynamic flows in the exterior fluid generated by active membrane stresses on the organelle. Direct interactions between the cytoskeleton and the membrane could suppress higher-order shape instabilities. In addition, in cells with stacked Golgi cisternae, the cytoskeletal scaffolding could lead to a suppression of the trafficking-induced cisternal drift instability.

We hope the ideas presented here will stimulate further theoretical studies and careful biophysical experiments on a system of organelles, such as Golgi and endosomes, in the trafficking pathway. An exciting challenge is to construct synthetic realisations of nonequilibrium soft fluidics capable of controlled fission and fusion.

## ACKNOWLEDGEMENTS

We acknowledge contributions from K. Gowrishankar to an earlier study. AR acknowledges support from the Simons Foundation, and stimulating discussions with M. Shelley and B. Chakrabarti. MR acknowledges support and funding from the Department of Atomic Energy (India), under project no. RTI4006, the Simons Foundation (Grant No. 287975), and the JC Bose Fellowship from DST-SERB (India). RGM acknowledges support from the EMBL Australia program and the Australian Research Council Centre of Excellence for Mathematical Analysis of Cellular Systems (CE230100001).

