## Supplementary Information for "Active morphodynamics of intracellular organelles in the trafficking pathway"

#### CONTENTS

I. Non-equilibrium Model and Active Hydrodynamic Equations 1
A. Mass balance and kinematics 3
B. Force balance and kinetics 4
C. Surface variation and membrane stresses 6
D. Free-energy and constitutive relations 7
E. Adiabatic approximation 9
F. Dynamics of the areal densities 10
G. Active membrane stresses 11
H. Membrane equations of motion 14
I. Coupling to the ambient fluid 15
J. Boundary integral formulation 16
K. Osmosis and volume dynamics 18

II. Quasi-spherical Perturbation Analysis 20
A. Surface parameterization 20
B. Lamb's general solution of Stokes equations 20
C. Flows around a slightly deformed sphere 22
1. Zeroth order perturbation 23
2. First order perturbation 24
D. Dynamics of the excess areal fractions 26
E. Area and volume dynamics 27
F. Membrane shape equations 29
1. At zeroth order 29
2. At first order 31
G. Linear stability analysis 33
H. Solutions to membrane shape and composition 36
I. Solutions to the surrounding fluid flows 39

III. Nonlinear Perturbative Analysis 41
A. Weakly nonlinear compositional dynamics 41
B. Galerkin projection and truncation method 41
C. Dynamics of the centre-of-mass velocity 43
D. Bifurcation diagrams and phase portraits 43

IV. Mathematical Prerequisites 47
A. Differential Geometry of Surfaces 47
B. Calculus of Moving Surfaces 49
C. Integral and Transport Theorems 51

References 51

#### I. NON-EQUILIBRIUM MODEL AND ACTIVE HYDRODYNAMIC EQUATIONS

We study the role of membrane trafficking on the morphology of subcellular compartments, such as those encountered within the eukaryotic cell. These continuously interchange material amongst themselves via fission and fusion of small transport vesicles. The fission and fusion are orchestrated by a highly complex biochemical network of proteins driven by energy-consuming processes.

Table I contains a list of the experimentally measured quantities and their physiological range, which helps us to determine the appropriate length and time scales in our system. This includes the time-scale associated with the lateral diffusion of membrane components ($T_d$); the time-scale corresponding to the relaxation of membrane shape deformations ($T_s$); and lastly the time-scale associated with the inter-fission and fusion events of transport vesicles ($T_a$). We find that the lateral diffusion of material $T_d$ is much slower than the active processes of fission and fusion, $T_a$. Further, we find that $T_a$ is comparable to the shape relaxation time $T_s$ for membrane organelles of size $R \lesssim 1$ $\mu$m, while $T_a$ is considerably less than $T_s$ for larger values of $R$. This implies that fission and fusion could provide a continuous flux that maintains the cisternal membrane out of equilibrium.

We develop a general theory which allows us to study the dynamical interplay of such active processes of fission and fusion with the membrane morphology as well as the hydrodynamics of the ambient fluid. This is facilitated by an appropriate coarse-graining over the microscopic biochemical reactions, which allows us to derive hydrodynamic equations of such membrane compartments ap-

| | |
|---|---|
| Transport vesicle radius | $r = 30-40$ nm [1–3] |
| Golgi cisternal length scale | $R = 1-2$ $\mu$m [4–6] |
| Bending rigidity of organelle | $\kappa = 20\ k_BT \approx 10^{-19}$ J [7] |
| Cytoplasmic viscosity | $\eta = 10^{-2}$ Pa·s [8–10] |
| Membrane surface diffusion | $\gamma = 0.1-1$ $\mu$m$^2$/s [11, 12] |
| Total area of Golgi | $\mathcal{A}_0 = 10-250$ $\mu$m$^2$ [4, 6, 13] |
| Turnover time of Golgi | $\tau = 10$–$25$ mins [14, 15] |
| Trafficking event time | $T_a = 4\pi r^2\tau/\mathcal{A}_0 \approx 0.1-2$ s |
| Lateral diffusion time | $T_d = 4\pi R^2/\gamma \approx 10$–$100$ s |
| Shape relaxation (lowest mode) | $T_s = 2\eta R^3/\kappa \approx 0.25$–$2$ s |

TABLE I. Length scales, membrane mechanical and transport properties, and estimated time-scales relevant to the trafficking dynamics of a Golgi cisterna.

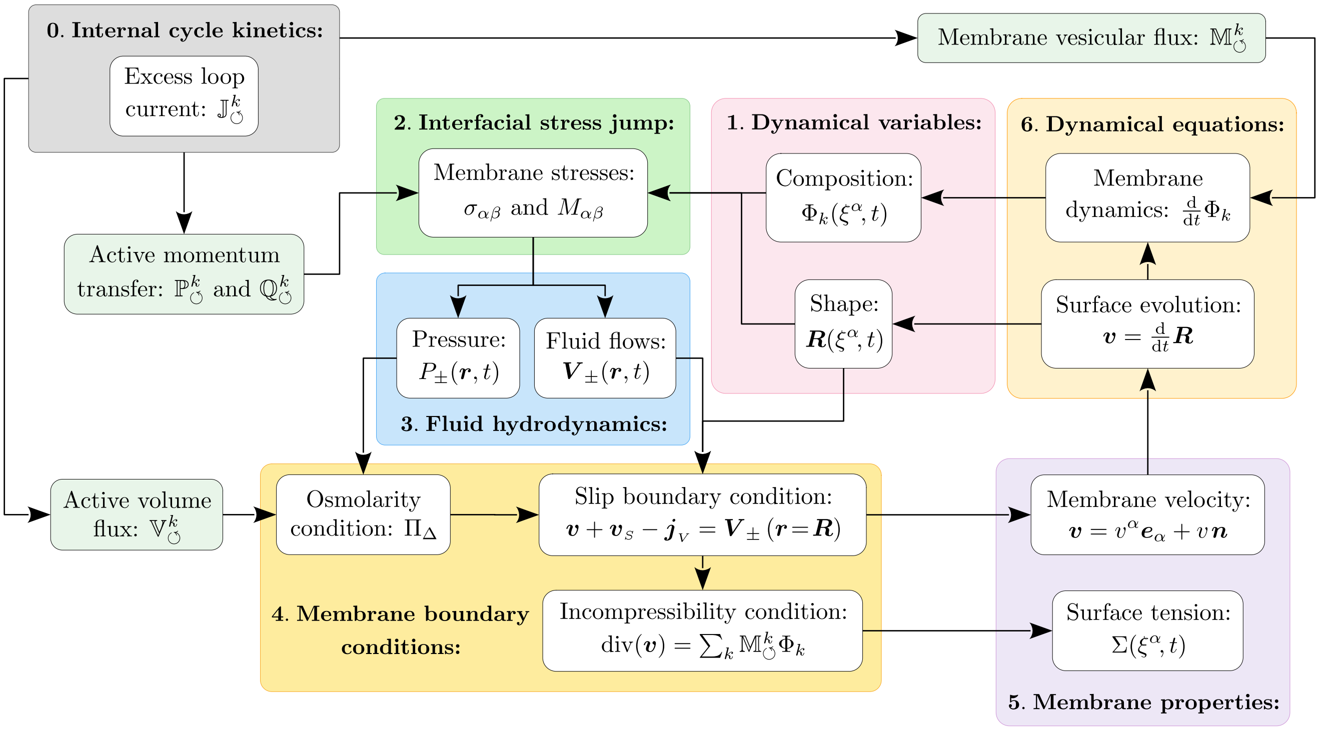


FIG. S.1. Flow chart summarizing the model. Starting at box 0, the excess loop current $\mathbb{J}^k_{\circlearrowleft}$ due to the biochemical reaction cycles associated with the fissogens and fusogens (labeled by $k$) drives proportional local fluxes of membrane mass $\mathbb{M}^k_{\circlearrowleft}$, volume flux $\mathbb{V}^k_{\circlearrowleft}$, as well as the active transfer of momentum. The latter is described in terms of the lowest nontrivial moments of the transverse force distribution to the membrane; namely, its first and second moments $\mathbb{P}^k_{\circlearrowleft}$ and $\mathbb{Q}^k_{\circlearrowleft}$, respectively. The dynamical variables are the area fraction $\Phi_k(\xi^\alpha, t)$ of active components (fissogens and fusogens) that reside on the membrane surface, which itself evolves, being described by the shape $\boldsymbol{R}(\xi^\alpha, t)$ embedded in a fluid with shear viscosity $\eta$. Here, $\xi^\alpha$ are curvilinear coordinates that parameterize the two-dimensional surface of the compartment. Initial data of the composition and shape allows us to compute the in-plane stress tensor $\sigma^{\alpha\beta}$ and the bending moment tensor $M^{\alpha\beta}$, which include passive contributions (due to the membrane elasticity and entropy-of-mixing terms), as well as active stresses that depend on $\mathbb{P}^k_{\circlearrowleft}$ and $\mathbb{Q}^k_{\circlearrowleft}$. The interfacial stress jump at the membrane is balanced by the viscous forces associated with the inner and outer ambient fluids, which satisfy the incompressible Stokes equations. Knowledge of the stress jump allows us to obtain the fluid pressures $P_\pm(\boldsymbol{r}, t)$ and the fluid flows $\boldsymbol{V}_\pm(\boldsymbol{r}, t)$ at each point $\boldsymbol{r}$, using a boundary integral representation. All of the terms in $\sigma^{\alpha\beta}$ and $M^{\alpha\beta}$ are known, apart from the local surface tension $\Sigma(\xi^\alpha, t)$, which acts as a Lagrange multiplier for the membrane incompressibility condition; the surface divergence of membrane flows $\boldsymbol{v}$ must equal the total vesicular flux of membrane area, $\mathbb{M} = \sum_k \mathbb{M}^k_{\circlearrowleft}\Phi_k$. To determine the surface tension, we use the slip boundary condition at the membrane ($\boldsymbol{r} = \boldsymbol{R}$) together with the incompressibility condition. The slip velocity $\boldsymbol{v}_S = \frac{1}{2\eta}\boldsymbol{e}^\alpha \nabla_\alpha \mathbb{Q}$, where the active moment $\mathbb{Q} = \sum_k \mathbb{Q}^k_{\circlearrowleft}\Phi_k$, whereas the volume flux $\boldsymbol{j}_V$ depends on the passive leakage of solvent through the membrane and the active volume flux $\mathbb{V} = \sum_k \mathbb{V}^k_{\circlearrowleft}\Phi_k$. The passive volume flux is driven by the difference between the osmotic pressure $\Pi_\Delta$ and the hydrostatic pressure across the interface. Once the surface tension $\Sigma$ is computed, the membrane velocity $\boldsymbol{v}$ can be readily obtained from the slip condition. This velocity can be decomposed along its surface components, namely $v^\alpha \boldsymbol{e}_\alpha$, which tells us how the membrane constituents are advected on the deformed surface. The material derivative of the compositional fields is thus given by $\frac{\mathrm{d}}{\mathrm{d}t} = \frac{\partial}{\partial t} + v^\alpha \nabla_\alpha$, where $\nabla_\alpha$ denotes the covariant derivative. Here, the evolution of compositional fields $\frac{\mathrm{d}}{\mathrm{d}t}\Phi_k$ is governed by the surface diffusion of membrane components and the active exchange of membrane material via the vesicular flux with the rates given by $\mathbb{M}^k_{\circlearrowleft}$. From an algorithmic viewpoint, this allows us to compute the mass area fractions $\Phi_k$ at the next time-step. Similarly, by using the normal velocity $v\boldsymbol{n}$, we can evolve the surface to determine its shape at the subsequent time point. This method of time-marching describes an explicit scheme to numerically solve the system of equations, starting from some initial data for the compositional fields and the shape. Note that we have assumed that the rates entering the internal cycle kinetics are *a priori* given and not regulated by feedback from the membrane mechanics, allowing us to operate in a regime where the transitions can be modelled as first-order kinetics with constant rates.

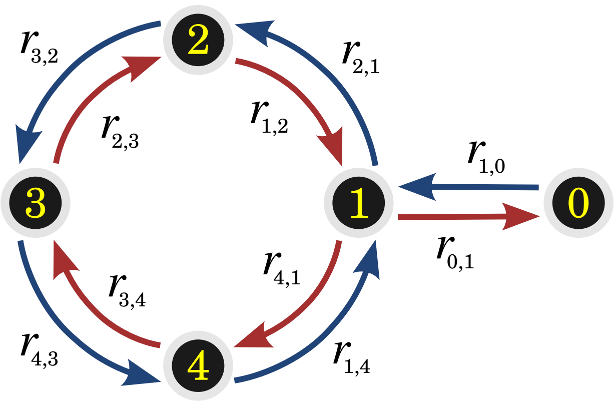


FIG. S.2. Kinetic diagram corresponding to the biochemical cycles of fission or fusion, where the number of internal active states is $\aleph_k = 4$; thus, there are 10 possible transitions, including the idle, zeroth state. Here, the forward (backward) transitions are indicated by the blue (red) arrow, and each has a first-order rate constant associated with it. We denote the latter by $r_{nm}$ for $m \rightarrow n$ (superscript $k$ is omitted here).

propriate to large spatial and temporal scales.

For convenience, we display a flow chart diagram in Fig. S.1 that outlines the model with its key elements and the connections between them. This can be viewed as an algorithmic, explicit scheme to compute and time-march the physical and dynamical properties of the model, such as membrane surface velocities, membrane stresses and surface tension, bulk fluid flows and pressures, as well as membrane composition and shape.

Note that the important equations have been boxed for ease of reading.

#### A. Mass balance and kinematics

We consider a closed membrane that is described by $\boldsymbol{R}(\xi^\alpha, t)$ at time $t$, where $\xi^\alpha$ are the local surface coordinates parametrizing the membrane. The membrane comprises three distinct components: active components that take part solely in the process of membrane fission and fusion, denoted by the superscript $k = 1$ (fissogens) and $k = 2$ (fusogens), respectively; and the passive membrane background component, indicated by the subscript '0'. Fission and fusion involve highly specific proteins and lipids, which undergo a number of biochemical reactions that primarily require the hydrolysis of either adenosine triphosphate (ATP) or guanosine triphosphate (GTP).

At a coarse level, we can describe the internal mechochemical processes of fission and fusion within a non-equilibrium Markov chain model, which assigns a state space to the various intermediary stages of the process, as well as transitions among those states that form at least one closed loop, or cycle, in the state space, as shown in Fig. S.2. Here, the realization of a single fusion or fission event is effectively characterized by the completion of an entire cycle on their corresponding kinetic diagrams. Upon the activation from an idle state (that we denote here by the subscript $n = 0$), the membrane constituents for each $k$ can transition into a sequence of intermediate states that are indicated by the index $n \in \{1, 2, 3, \ldots, \aleph_k\}$, where $\aleph_k$ is number of internal states associated with that cycle. Notably, this final state winds back to the first state, terminating the cycle. This could immediately continue into a new cycle, or it might be first deactivated into the idle state ($n = 0$), and after a dwell time a new cycle is initiated again, as in Fig. S.2. Note that this dwell time represents the waiting time between two distinct fission (or fusion) events.

The mass density for each $k$ at time $t$, within the $n$-th internal state, is denoted by $\rho_n^k(\xi^\alpha, t)$, whereas the mass density of $k = 0$ is given by $\rho_0(\xi^\alpha, t)$. Hence, the total mass density on the membrane can be written as

$$\rho(\xi^\alpha, t) = \rho_0(\xi^\alpha, t) + \sum_{k,n} \rho_n^k(\xi^\alpha, t), \tag{S.1}$$

where we use the notation that $\sum_{k,n}(\cdot) = \sum_k \sum_{n=0}^{\aleph_k}(\cdot)$. Moreover, the barycentric velocity of the membrane in the local surface basis is given by $\boldsymbol{v} = v^\alpha \boldsymbol{e}_\alpha + v\boldsymbol{n}$, with $\boldsymbol{n}$ and $\boldsymbol{e}_\alpha$ as the normal and in-plane tangent vectors to the membrane, respectively (see Supplementary Section IV.A for definitions). This centre-of-mass velocity is defined by

$$\rho\boldsymbol{v} = \rho_0\boldsymbol{v}_0 + \sum_{k,n} \rho_n^k \boldsymbol{v}_n^k, \tag{S.2}$$

where the background membrane velocity can be decomposed as $\boldsymbol{v}_0 = v_0^\alpha \boldsymbol{e}_\alpha + v\boldsymbol{n}$, and the velocity of $k$-th species in state $n$ is given by $\boldsymbol{v}_n^k = v_n^{k\alpha} \boldsymbol{e}_\alpha + v\boldsymbol{n}$. Notice that the normal velocities of $\boldsymbol{v}$, $\boldsymbol{v}_0$, and $\boldsymbol{v}_n^k$ must all be the same for all species and their associated internal states.

Furthermore, we denote the in-plane diffusive flux of $k$-th species, in state $n$, by $\boldsymbol{j}_n^k$, defined as follows:

$$\boldsymbol{j}_n^k = \rho_n^k \left(\boldsymbol{v}_n^k - \boldsymbol{v}\right) = \rho_n^k \left(v_n^{k\alpha} - v^\alpha\right) \boldsymbol{e}_\alpha = j_n^{k\alpha} \boldsymbol{e}_\alpha, \tag{S.3}$$

which shows that this flux lies solely within the tangent plane of the membrane, with components $j_n^{k\alpha}$. Similarly, the diffusive flux corresponding to $\rho_0$ is defined by

$$\boldsymbol{j}_0 = \rho_0(\boldsymbol{v}_0 - \boldsymbol{v}). \tag{S.4}$$

As a corollary, it can be shown that the summation over all diffusive fluxes in Eqs. (S.3) and (S.4) gives

$$\boldsymbol{j}_0 + \sum_{k,n} \boldsymbol{j}_n^k = \boldsymbol{0}. \tag{S.5}$$

The net mass of a $k$-th species within the state $n$ over a membrane patch $\mathcal{S}$ changes due to the diffusive mass flux at the boundary $\partial\mathcal{S}$ of that patch, as well as due to the chemical reactions that lead to forward and backward transitions between the states $n$ and $m$ for every $k$. Here, we denote $r^k_{nm}$ to be the transition rate from the state $m$ to state $n$ for the $k$-th species (see Fig. S.2). Thus,

$$\frac{\mathrm{d}}{\mathrm{d}t}\int\limits_{\mathcal{S}} \rho^k_n \,\mathrm{d}S = -\int\limits_{\partial\mathcal{S}} \boldsymbol{j}^k_n \cdot \boldsymbol{\nu}\,\mathrm{d}l + \int\limits_{\mathcal{S}} \sum_{m\in\mathfrak{I}^k_n} \rho\,\mathbb{J}^k_{nm}\,\mathrm{d}S, \quad \text{(S.6)}$$

where $\boldsymbol{\nu}$ is the tangent unit vector normal to the boundary $\partial\mathcal{S}$, with $\boldsymbol{\nu}\cdot\boldsymbol{n}=0$. The index set $\mathfrak{I}^k_n$ gives all the internal states of the $k$-cycle which are directly connected to state $n$; for example, the states one and zero in Fig. S.2 have $\mathfrak{I}^k_1=\{0,2,4\}$ and $\mathfrak{I}^k_0=\{1\}$, respectively. Here, we define the material derivative to be $\frac{\mathrm{d}}{\mathrm{d}t}(\cdot)=\frac{\partial}{\partial t}+v^\alpha(\cdot)_{,\alpha}$. In the last term of Eq. (S.6), $\mathbb{J}^k_{nm}$ denotes the probability current of the internal cycles, namely

$$\mathbb{J}^k_{nm} = r^k_{nm}\phi^k_m - r^k_{mn}\phi^k_n, \quad \text{(S.7)}$$

where the mass fraction $\phi^k_n=\rho^k_n/\rho$. Note that $\mathbb{J}^k_{nm}$ is an antisymmetric object with respect to the lower indices, that is, we have $\mathbb{J}^k_{nm}=-\mathbb{J}^k_{mn}$. By using the Reynolds transport theorem on the left-hand-side of (S.6), namely

$$\frac{\mathrm{d}}{\mathrm{d}t}\int_{\mathcal{S}}\rho\,\mathrm{d}S = \int_{\mathcal{S}}\left(\frac{\mathrm{d}\rho}{\mathrm{d}t}+\rho\,\mathrm{div}\,\boldsymbol{v}\right)\mathrm{d}S, \quad \text{(S.8)}$$

and then by applying the divergence theorem to the first term on the right-hand-side of Eq. (S.6), the local form of the mass conservation is found to be

$$\boxed{\dot\rho^k_n + \rho^k_n\,\mathrm{div}\,\boldsymbol{v} = -\mathrm{div}\big(\boldsymbol{j}^k_n\big) + \rho\sum_{m\in\mathfrak{I}^k_n}\mathbb{J}^k_{nm},} \quad \text{(S.9)}$$

where the dot notation denotes the material derivative. Here, the divergence of the diffusive current can be also written as $\mathrm{div}\big(\boldsymbol{j}^k_n\big)=j^{k\alpha}_{n;\alpha}$, where the semi-colon is used to indicate a covariant derivative, cf. Eq. (S.416). At equilibrium, $\mathbb{J}^k_{mn}$ must be identically zero due to detailed balance, but in a non-equilibrium steady-state the currents do not need to vanish as detailed balance is violated.

Similarly, the rate of change in the membrane mass of the background component is given by

$$\frac{\mathrm{d}}{\mathrm{d}t}\int\limits_{\mathcal{S}}\rho_0\,\mathrm{d}S = -\int\limits_{\partial\mathcal{S}}\boldsymbol{j}_0\cdot\boldsymbol{\nu}\,\mathrm{d}l + \int\limits_{\mathcal{S}}\rho\,\mathbb{M}\,\mathrm{d}S, \quad \text{(S.10)}$$

where the last term is the total membrane mass removed from $(k=1)$ and brought onto $(k=2)$ the surface $\mathcal{S}$, as a result of the internal chemical reactions, where the net rate of mass flux over both $k$-cycles is given by

$$\mathbb{M} = \sum_{k,n,m}\mathbb{M}^k_{nm}r^k_{nm}\phi^k_m, \quad \text{(S.11)}$$

with $\sum_{k,n,m}(\cdot)=\sum_k\sum_{n=0}^{\aleph_k}\sum_{m\in\mathfrak{I}^k_n}(\cdot)$, and $\mathbb{M}^k_{nm}$ is the amount of mass exchanged in a transition from the state $m$ to $n$ for the $k$-cycle. Again, by means of the Reynolds transport theorem and the divergence theorem, this leads to the following local form of the mass balance:

$$\boxed{\dot\rho_0 + \rho_0\,\mathrm{div}\,\boldsymbol{v} = -\mathrm{div}\big(\boldsymbol{j}_0\big) + \rho\,\mathbb{M},} \quad \text{(S.12)}$$

with $\mathrm{div}\big(\boldsymbol{j}_0\big)=j^\alpha_{0;\alpha}$. Thus, by summing over all densities from Eqs. (S.9) and (S.12), the local mass balance of the total density can be written as follows:

$$\boxed{\dot\rho + \rho\,\mathrm{div}\,\boldsymbol{v} = \rho\,\mathbb{M}.} \quad \text{(S.13)}$$

Notice that the diffusive currents vanish due to Eq. (S.5), and also $\sum_n\sum_{m\in\mathfrak{I}^k_n}\mathbb{J}^k_{nm}=0$ as $\mathbb{J}^k_{nm}=-\mathbb{J}^k_{mn}$.

By means of the local mass balance in Eq. (S.13), the dynamics of the mass fraction $\Phi_0=\rho_0/\rho$ is governed by

$$\dot\Phi_0 = (1-\Phi_0)\,\mathbb{M} + \frac{1}{\rho}\sum_{k,n}\mathrm{div}\big(\boldsymbol{j}^k_n\big), \quad \text{(S.14)}$$

where Eq. (S.5) and the chain rule, $\dot\rho_0=\rho\,\dot\Phi_0+\dot\rho\,\Phi_0$, are used to derive the above expression. Note that $\Phi_0$ is linked to the other mass fractions through Eq. (S.1),

$$1-\Phi_0 = \sum_{k,n}\phi^k_n, \quad \text{(S.15)}$$

where $k=1,2$. Thus there are only $\sum_k(1+\aleph_k)$ linearly independent mass fractions. As analogous to Eq. (S.14), a dynamical equation for the mass fractions $\phi^k_n$ can be derived, that is,

$$\dot\phi^k_n = -\phi^k_n\,\mathbb{M} + \sum_{m\in\mathfrak{I}^k_n}\mathbb{J}^k_{nm} - \frac{1}{\rho}\mathrm{div}\big(\boldsymbol{j}^k_n\big). \quad \text{(S.16)}$$

#### B. Force balance and kinetics

The membrane patch $\mathcal{S}$ that is subjected to a traction $\boldsymbol{T}$ at the boundary $\partial\mathcal{S}$ of the domain. As a consequence, the total linear momentum over all membrane constituents, *i.e.* $\int_{\mathcal{S}}\big(\rho_0\boldsymbol{v}_0+\sum_{k,n}\rho^k_n\boldsymbol{v}^k_n\big)\,\mathrm{d}S$, changes as follows:

$$\frac{\mathrm{d}}{\mathrm{d}t}\int\limits_{\mathcal{S}}\rho\boldsymbol{v}\,\mathrm{d}S = \int\limits_{\mathcal{S}}\boldsymbol{f}\,\mathrm{d}S + \int\limits_{\partial\mathcal{S}}\boldsymbol{T}^\alpha\nu_\alpha\,\mathrm{d}l + \int\limits_{\mathcal{S}}\rho\boldsymbol{v}\,\mathbb{M}\,\mathrm{d}S, \quad \text{(S.17)}$$

where $\boldsymbol{\nu}=\nu_\alpha\boldsymbol{e}^\alpha$ is the unit vector normal to the boundary $\partial\mathcal{S}$ and also tangent to the surface patch $\mathcal{S}$, and $\boldsymbol{f}$ is the overall (mass-weighted) body force acting on all membrane species. The last term in Eq. (S.17) gives the momentum transfer due to the mass flux of membrane material carried in (and out) of the membrane from (and to) its surroundings, which results from the chemical reaction cycles. This assumes that the added, or removed, material moves with a velocity given by the barycentric velocity $\boldsymbol{v}(t)$. By means of Reynolds transport theorem, $\frac{\mathrm{d}}{\mathrm{d}t}\int_{\mathcal{S}}(\rho\boldsymbol{v})\,\mathrm{d}S=\int_{\mathcal{S}}\big[\frac{\mathrm{d}}{\mathrm{d}t}(\rho\boldsymbol{v})+\rho\boldsymbol{v}\,\mathrm{div}\,\boldsymbol{v}\big]\,\mathrm{d}S$, together with the mass balance relation in Eq. (S.13), we obtain that

$$\frac{\mathrm{d}}{\mathrm{d}t}\int\limits_{\mathcal{S}}\rho\boldsymbol{v}\,\mathrm{d}S = \int\limits_{\mathcal{S}}(\rho\dot{\boldsymbol{v}}+\rho\boldsymbol{v}\,\mathbb{M})\,\mathrm{d}S. \quad \text{(S.18)}$$

Moreover, by applying the divergence theorem on the boundary term of Eq. (S.17), this results in the following local form of linear momentum balance:

$$\rho\dot{\boldsymbol{v}} = \boldsymbol{f} + \boldsymbol{T}^{\alpha}_{;\alpha}. \tag{S.19}$$

The stress vectors $\boldsymbol{T}^{\alpha}$ are the projections of the tractions onto the constant surface lines, parametrized by $\xi^{\alpha}$. This can be further decomposed into the basis of the surface:

$$\boldsymbol{T}^{\alpha} = T^{\alpha\beta}\boldsymbol{e}_{\beta} + N^{\alpha}\boldsymbol{n}, \tag{S.20}$$

where $N^{\alpha}$ describes the out-of-plane shear stresses, and $T^{\alpha\beta}$ are the components of the in-plane stress tensor. By also decomposing the body force $\boldsymbol{f}$ in the basis $\{\boldsymbol{e}_{\alpha}, \boldsymbol{n}\}$, *i.e.* $\boldsymbol{f} = f^{\alpha}\boldsymbol{e}_{\alpha} + f\boldsymbol{n}$, then the normal component of the momentum balance relation in Eq. (S.19) is found to be

$$\rho\dot{\boldsymbol{v}}\cdot\boldsymbol{n} = f + T^{\alpha\beta}b_{\alpha\beta} + N^{\alpha}_{;\alpha}, \tag{S.21}$$

whereas the in-plane component is given by

$$\rho\dot{\boldsymbol{v}}\cdot\boldsymbol{e}^{\beta} = f^{\beta} + T^{\alpha\beta}_{;\alpha} - N^{\alpha}b^{\beta}_{\alpha}, \tag{S.22}$$

by using *Gauss-Weingarten* equations, cf. Eq. (S.431).

In addition, the rate of change of angular momentum is given by the sum of the torques that arise from the body forces and tractions, as well as a director traction $\boldsymbol{M}$ that acts on the boundary of membrane patch $\mathcal{S}$. In our case, the coarse-grained membrane is assumed to be an object of zero thickness, and its associated director field is chosen to be the same as the unit vector normal $\boldsymbol{n}$ to the surface, which is known as the Kirchhoff–Love assumption [16]. At every point along the boundary $\partial\mathcal{S}$, the director traction $\boldsymbol{M}$ exerts equal and opposite forces on the director field $\boldsymbol{n}$, which results in a moment per unit length $\boldsymbol{m} = \boldsymbol{n}\times\boldsymbol{M}$, giving rise to an overall internal torque $\int_{\partial\mathcal{S}}\boldsymbol{m}\,\mathrm{d}l$. This implies that $\boldsymbol{m}$ has only tangential components, even though the director traction may have in general both normal and in-plane components. Hence, the corresponding stress vectors $\boldsymbol{M}^{\alpha}$, which are obtained by projecting $\boldsymbol{M}$ along the curves of constant $\xi^{\alpha}$ (that is, $\boldsymbol{M} = \boldsymbol{M}^{\alpha}\nu_{\alpha}$), can be chosen without any loss of generality to be in the tangent plane of the membrane; namely, we define

$$\boldsymbol{M}^{\alpha} = -M^{\alpha\beta}\boldsymbol{e}_{\beta}, \tag{S.23}$$

where $M^{\alpha\beta}$ is known as the bending moment tensor. As a result, the conservation of angular momentum yields

$$\frac{\mathrm{d}}{\mathrm{d}t}\int_{\mathcal{S}}\boldsymbol{R}\times\rho\boldsymbol{v}\,\mathrm{d}S = \int_{\mathcal{S}}\boldsymbol{R}\times(\boldsymbol{f}+\rho\boldsymbol{v}\,\mathbb{M})\,\mathrm{d}S + \int_{\partial\mathcal{S}}(\boldsymbol{m}+\boldsymbol{R}\times\boldsymbol{T})\cdot\boldsymbol{\nu}\,\mathrm{d}l, \tag{S.24}$$

which, by applying Reynolds transport theorem, becomes

$$\int_{\mathcal{S}}\boldsymbol{R}\times\rho\dot{\boldsymbol{v}}\,\mathrm{d}S = \int_{\mathcal{S}}\boldsymbol{R}\times\boldsymbol{f}\,\mathrm{d}S + \int_{\partial\mathcal{S}}(\boldsymbol{m}+\boldsymbol{R}\times\boldsymbol{T})\cdot\boldsymbol{\nu}\,\mathrm{d}l. \tag{S.25}$$

By means of divergence theorem, a local form of the angular momentum balance can be written as

$$\begin{aligned}\boldsymbol{R}\times\rho\dot{\boldsymbol{v}} = \boldsymbol{R}\times\boldsymbol{f} + \boldsymbol{e}_{\alpha}\times\boldsymbol{T}^{\alpha} + \boldsymbol{R}\times\boldsymbol{T}^{\alpha}_{;\alpha}\\ + \boldsymbol{n}\times\boldsymbol{M}^{\alpha}_{;\alpha} - b^{\beta}_{\alpha}\boldsymbol{e}_{\beta}\times\boldsymbol{M}^{\alpha},\end{aligned} \tag{S.26}$$

where the covariant derivatives have been expanded out and the Gauss-Weingarten equations have been used to derive the above expression. By subtracting out the cross product of $\boldsymbol{R}$ with the momentum balance relation in Eq. (S.19), then $\boldsymbol{e}_{\alpha}\times\boldsymbol{T}^{\alpha} + \boldsymbol{n}\times\boldsymbol{M}^{\alpha}_{;\alpha} - b^{\beta}_{\alpha}\boldsymbol{e}_{\beta}\times\boldsymbol{M}^{\alpha} = \boldsymbol{0}$. Therefore, by using the definition of the bending moment tensor in Eq. (S.23), the latter equation leads to

$$\Big[\big(T^{\alpha\beta} - b^{\beta}_{\gamma}M^{\gamma\alpha}\big)\boldsymbol{e}_{\beta} + \Big(N^{\alpha} + M^{\beta\alpha}_{;\beta}\Big)\boldsymbol{n}\Big]\times\boldsymbol{e}_{\alpha} = \boldsymbol{0}, \tag{S.27}$$

which readily implies that

$$\sigma^{\alpha\beta} = T^{\alpha\beta} - b^{\beta}_{\gamma}M^{\gamma\alpha} \tag{S.28}$$

must be a symmetric tensor, and

$$N^{\alpha} = -M^{\beta\alpha}_{;\beta}, \tag{S.29}$$

so that the linear and angular momentum are conserved. Notice that in a system where the boundary moment $\boldsymbol{m}$ vanishes identically, the in-plane stress tensor $T^{\alpha\beta}$ must be symmetric, and the out-of-plane shear stress $N^{\alpha} = 0$.

As a result, the equation of motion of the membrane along the normal direction is found to be

$$\boxed{\rho\dot{\boldsymbol{v}}\cdot\boldsymbol{n} = f + \sigma^{\alpha\beta}b_{\alpha\beta} + M^{\alpha\beta}b_{\alpha\gamma}b^{\gamma}_{\beta} - M^{\alpha\beta}_{;\alpha\beta},} \tag{S.30}$$

which is known as *the shape equation.* It is noteworthy to mention that the identity $b_{\alpha\gamma}b^{\gamma}_{\beta} = 2H\,b_{\alpha\beta} - K\,g_{\alpha\beta}$ can be used to further unpack the above expression. On the other hand, the force-balance along the surface now reads

$$\boxed{\rho\dot{\boldsymbol{v}}\cdot\boldsymbol{e}^{\alpha} = f^{\alpha} + \sigma^{\alpha\beta}_{;\beta} + (b^{\alpha}_{\beta}M^{\beta\gamma})_{;\gamma} + b^{\alpha}_{\beta}M^{\gamma\beta}_{;\gamma}.} \tag{S.31}$$

To solve these equations of motion, the symmetrized form of the in-plane stress tensor $\sigma^{\alpha\beta}$ and the bending moment tensor $M^{\alpha\beta}$ must be first prescribed. In general, these stress tensors include both passive (*e.g.* elasticity and viscous dissipation) and active contributions.

Hereinafter, the passive contributions to the traction force and the director traction are denoted by $\boldsymbol{T}_0$ and $\boldsymbol{M}_0$, respectively. These terms are found from an elastic free-energy of membranes, and this will be discussed in the next section. On the other hand, the active terms are indicated herein by the subscript $A$; namely, the tractions $\boldsymbol{T}_A$ and $\boldsymbol{M}_A$, which are induced on the membrane surface as a result of the active forces that act in the surrounding fluid across a finite separation transverse to the membrane. Thus, the net traction force is $\boldsymbol{T} = \boldsymbol{T}_0 + \boldsymbol{T}_A$, while the total director traction is given by $\boldsymbol{M} = \boldsymbol{M}_0 + \boldsymbol{M}_A$.

#### C. Surface variation and membrane stresses

We consider that the elastic free-energy $\mathcal{F}$ of a membrane patch $\mathcal{S}$ depends solely on its geometry, *i.e.* $\mathcal{F}$ is only a function of the metric tensor $g_{\alpha\beta}$ and the curvature tensor $b_{\alpha\beta}$. Given this free-energy functional form,

$$\mathcal{F}(g_{\alpha\beta}, b_{\alpha\beta}) = \int_{\mathcal{S}} F(g_{\alpha\beta}, b_{\alpha\beta})\, \mathrm{d}S, \tag{S.32}$$

with $F$ as the surface free-energy density, we seek to obtain the mechanical response of the membrane, namely its free-energy change $\delta\mathcal{F}$, due to a shape deformation $\boldsymbol{R} \mapsto \boldsymbol{R} + \delta\boldsymbol{R}$. This procedure requires us to compute the subsequent changes in the basis vectors, $\delta\boldsymbol{e}_\alpha$, and its normal, $\delta\boldsymbol{n}$, as well as the corresponding variations in both the metric tensor, $\delta g_{\alpha\beta}$, and the curvature tensor, $\delta b_{\alpha\beta}$. However, computing these objects is typically cumbersome [17], and therefore here we adopt a Lagrange multiplier technique [18] which circumvents this difficulty by treating the variables $\boldsymbol{R}$, $\boldsymbol{n}$, $\boldsymbol{e}_\alpha$, $g_{\alpha\beta}$, and $b_{\alpha\beta}$ as independent, including the known geometrical relationships between these variables as constraints. Thus, a Lagrange function can be constructed by the following free-energy functional, $\mathcal{F}_C[\boldsymbol{R}, \boldsymbol{n}, \boldsymbol{e}_\alpha, g_{\alpha\beta}, b_{\alpha\beta}]$, which is defined by

$$\begin{aligned}\mathcal{F}_C = \mathcal{F} \,+\, &\int_{\mathcal{S}} \mathrm{d}S\, \left[\boldsymbol{\Upsilon}^\alpha \cdot (\boldsymbol{e}_\alpha - \boldsymbol{R}_{,\alpha}) + \lambda^\alpha\, (\boldsymbol{e}_\alpha \cdot \boldsymbol{n})\right] \\ &+ \int_{\mathcal{S}} \mathrm{d}S\, \left[\frac{\lambda}{2}\left(\boldsymbol{n}^2 - 1\right) + \frac{\lambda^{\alpha\beta}}{2}\left(g_{\alpha\beta} - \boldsymbol{e}_\alpha \cdot \boldsymbol{e}_\beta\right)\right] \\ &+ \int_{\mathcal{S}} \mathrm{d}S\, \left[\Lambda^{\alpha\beta}\left(b_{\alpha\beta} + \boldsymbol{e}_\alpha \cdot \boldsymbol{n}_{,\beta}\right)\right],\end{aligned} \tag{S.33}$$

where the Lagrange multiplier $\boldsymbol{\Upsilon}^\alpha$ dictates the relation between the basis $\boldsymbol{e}_\alpha$ and the embedding function $\boldsymbol{R}$, $\lambda^\alpha$ imposes the orthogonality condition between $\boldsymbol{e}_\alpha$ and $\boldsymbol{n}$, and $\lambda$ enforces the normalization of the unit vector $\boldsymbol{n}$. Similarly, the multipliers $\Lambda^{\alpha\beta}$ and $\lambda^{\alpha\beta}$ ensure the definitions of the metric tensor $g_{\alpha\beta}$ and the curvature tensor $b_{\alpha\beta}$, respectively, in terms of the basis vectors $\boldsymbol{e}_\alpha$ and the normal vector $\boldsymbol{n}$. Note that both these surface tensors are required to be symmetric, namely we have $\lambda^{\alpha\beta} = \lambda^{\beta\alpha}$ and $\Lambda^{\alpha\beta} = \Lambda^{\beta\alpha}$. The advantage of this approach is that we do not need to explicitly consider the changes in $g_{\alpha\beta}$ and $b_{\alpha\beta}$ due to a variation in the embedding $\boldsymbol{R}$, as these tensors are now treated as independent variables [18].

To determine $\boldsymbol{\Upsilon}^\alpha$, the functional derivative of $\mathcal{F}_C$ with respect to $\boldsymbol{e}_\alpha$ must vanish ($\delta\mathcal{F}_C/\delta\boldsymbol{e}_\alpha = 0$), which yields

$$\boldsymbol{\Upsilon}^\alpha = \left(\lambda^{\alpha\beta} + \Lambda^{\alpha\gamma} b^\beta_\gamma\right)\boldsymbol{e}_\beta - \lambda^\alpha \boldsymbol{n}, \tag{S.34}$$

where the *Gauss–Weingarten* equations have been used, *i.e.* $\boldsymbol{e}_{\alpha;\beta} = b_{\alpha\beta}\,\boldsymbol{n}$, and $\boldsymbol{n}_{,\alpha} = -b^\beta_\alpha\,\boldsymbol{e}_\beta$, which themselves can be obtained respectively from the constraints given by $\delta\mathcal{F}_C/\delta\Lambda^{\alpha\beta} = 0$ and $\delta\mathcal{F}_C/\delta\lambda = 0$. The boundary terms in Eq. (S.34) vanish due to the orthogonality of the basis vectors. Likewise, $\lambda^{\alpha\beta}$ and $\Lambda^{\alpha\beta}$ are given by the Euler-Lagrange equations associated to $g_{\alpha\beta}$ and $b_{\alpha\beta}$, namely,

$$\lambda^{\alpha\beta} = -\frac{2}{\sqrt{g}}\frac{\partial(\sqrt{g}F)}{\partial g_{\alpha\beta}}, \quad \text{and} \quad \Lambda^{\alpha\beta} = -\frac{\partial F}{\partial b_{\alpha\beta}}, \tag{S.35}$$

respectively. Since the variable $\boldsymbol{R}$ appears only in the second term of Eq. (S.33), the infinitesimal variation of the functional $\mathcal{F}_C$ due to a deformation $\delta\boldsymbol{R}$ is written as

$$\delta_{\boldsymbol{R}}(\mathcal{F}_C) = \int_{\mathcal{S}} \mathrm{d}S\, \left(\boldsymbol{\Upsilon}^\alpha_{\;;\alpha} \cdot \delta\boldsymbol{R}\right) - \oint_{\partial\mathcal{S}} \mathrm{d}l\; \nu_\alpha\left(\boldsymbol{\Upsilon}^\alpha \cdot \delta\boldsymbol{R}\right), \tag{S.36}$$

with $\boldsymbol{\nu} = \nu_\alpha \boldsymbol{e}^\alpha$ being the unit tangent vector normal to the boundary. If the shape variation $\delta\boldsymbol{R}$ vanishes on $\partial\mathcal{S}$, that is, the position vector $\boldsymbol{R}$ is fixed at the boundary, then the last term in Eq. (S.36) is identically zero, and thus we have that

$$\frac{\delta\mathcal{F}_C}{\delta\boldsymbol{R}} = \boldsymbol{\Upsilon}^\alpha_{\;;\alpha} = 0, \tag{S.37}$$

which states that at equilibrium the stress vector $\boldsymbol{\Upsilon}^\alpha$ is a covariantly conserved variable on the membrane surface.

Similarly, the local variation of $\mathcal{F}_C$ due to a infinitesimal change $\delta\boldsymbol{n}$ in the normal unit vector is given by

$$\begin{aligned}\delta_{\boldsymbol{n}}(\mathcal{F}_C) = &\int_{\mathcal{S}} \mathrm{d}S\, \left[\left(\lambda + \Lambda^{\alpha\beta} b_{\alpha\beta}\right)\boldsymbol{n} + \left(\lambda^\alpha - \Lambda^{\alpha\beta}_{\;;\beta}\right)\boldsymbol{e}_\alpha\right] \cdot \delta\boldsymbol{n} \\ &+ \oint_{\partial\mathcal{S}} \mathrm{d}l\; \nu_\beta\left(\Lambda^{\alpha\beta}\boldsymbol{e}_\alpha \cdot \delta\boldsymbol{n}\right).\end{aligned} \tag{S.38}$$

If the normal unit vector is fixed at the boundary, that is, $\delta\boldsymbol{n} = 0$ on $\partial\mathcal{S}$, then the associated Euler-Lagrange equation with respect to $\boldsymbol{n}$ is found to be

$$\frac{\delta\mathcal{F}_C}{\delta\boldsymbol{n}} = \left(\lambda + \Lambda^{\alpha\beta} b_{\alpha\beta}\right)\boldsymbol{n} + \left(\lambda^\alpha - \Lambda^{\alpha\beta}_{\;;\beta}\right)\boldsymbol{e}_\alpha = 0, \tag{S.39}$$

and therefore we have that

$$\lambda = -\Lambda^{\alpha\beta} b_{\alpha\beta} \quad \text{and} \quad \lambda^\alpha = \Lambda^{\alpha\beta}_{\;;\beta}. \tag{S.40}$$

However, in general, neither $\boldsymbol{n}$ and $\boldsymbol{R}$ are anchored on the boundary of the patch $\mathcal{S}$, but they are free variables. Consequently, the boundary terms give rise to traction forces and boundary moments. Using the Euler-Lagrange equations in (S.37) and (S.39), then the total infinitesimal variation of the free-energy $\mathcal{F}_C$ is given by

$$\delta\mathcal{F}_C = \oint_{\partial\mathcal{S}} \mathrm{d}l\; \nu_\beta\left(\Lambda^{\alpha\beta}\boldsymbol{e}_\alpha \cdot \delta\boldsymbol{n} - \boldsymbol{\Upsilon}^\beta \cdot \delta\boldsymbol{R}\right). \tag{S.41}$$

This must be balanced by the traction force $\boldsymbol{T}_0$ and the direction traction $\boldsymbol{M}_0$ at the boundary $\partial\mathcal{S}$, which are (by definition) the passive contributions to the tractions $\boldsymbol{T}$ and $\boldsymbol{M}$, respectively. Thus, $\delta\mathcal{F}_C$ equals the virtual work,

$$\delta\mathcal{W} = \oint_{\partial\mathcal{S}} \mathrm{d}l\; \nu_\alpha\left(\boldsymbol{T}^\alpha_0 \cdot \delta\boldsymbol{R} + \boldsymbol{M}^\alpha_0 \cdot \delta\boldsymbol{n}\right), \tag{S.42}$$

where $\boldsymbol{T}_0^\alpha$ and $\boldsymbol{M}_0^\alpha$ are the projections of the tractions onto the surface lines of constant $\xi^\alpha$. From Eq. (S.41) and (S.42), we find that $\boldsymbol{T}_0^\alpha = -\boldsymbol{\Upsilon}^\alpha$ and $\boldsymbol{M}_0^\alpha = \Lambda^{\alpha\beta}\boldsymbol{e}_\alpha$. By decomposing these stress vectors into the surface basis, as analogous to Eqs. (S.23) and (S.20), *i.e.* $\boldsymbol{M}_0^\alpha = -M_0^{\alpha\beta}\boldsymbol{e}_\beta$ and $\boldsymbol{T}_0^\alpha = T_0^{\alpha\beta}\boldsymbol{e}_\beta + N_0^\alpha\boldsymbol{n}$, then we have that the bending moment tensor $M_0^{\alpha\beta} = -\Lambda^{\alpha\beta}$, the in-plane stress tensor $\sigma_0^{\alpha\beta} = T_0^{\alpha\beta} - b_\gamma^\beta M_0^{\gamma\alpha} = -\lambda^{\alpha\beta}$, cf. Eq. (S.28), and the out-of-plane shear stress $N_0^\alpha = \lambda^\alpha$. The latter, together with the second equation in (S.40), gives $N_0^\alpha = -M_0^{\alpha\beta}{}_{;\beta}$, as found previously in Eq. (S.29) for the overall shear stress $N^\alpha$. Since $\sigma_0^{\alpha\beta}$ is symmetric, this shows that the passive contributions to the tractions, which arise from the free-energy $\mathcal{F}$, respect the conservation of the angular momentum. Thus, the passive stresses are given by

$$\boxed{\sigma_0^{\alpha\beta} = \frac{2}{\sqrt{g}}\frac{\partial(\sqrt{g}F)}{\partial g_{\alpha\beta}}, \quad \text{and} \quad M_0^{\alpha\beta} = \frac{\partial F}{\partial b_{\alpha\beta}},} \tag{S.43}$$

and the stress vector reduces to $\boldsymbol{\Upsilon}^\alpha = -T_0^{\alpha\beta}\boldsymbol{e}_\beta + M_0^{\alpha\beta}{}_{;\beta}\,\boldsymbol{n}$.

#### D. Free-energy and constitutive relations

At a coarse-grained level, the Helmholtz free-energy $\mathcal{F}$ of a multi-component biomembrane (described by three distinct membrane components, and their internal states) includes an elastic energetic cost due to surface bending and stretching (*i.e.* the Helfrich energy density) [19], an entropy of mixing of the various species, as well as the interaction terms given by a phenomenological coupling between the mean curvature $H$ of the surface and the local density of membrane components [20]. We consider that the membrane forms a closed surface, a membrane compartment, and we denote its surface by $\mathcal{M}$ (formally, we say that $\mathcal{M}$ is homeomorphic to the Euclidean 2-sphere). The overall free-energy of the compartment is therefore given by $\mathcal{F} = \int_\mathcal{M} F\,\mathrm{d}S$, where the free-energy density

$$\boxed{F = \Sigma + 2\kappa H^2 + 2\kappa H C_\mathrm{m} + V_\mathrm{m},} \tag{S.44}$$

with $\kappa$ and $\Sigma$ as the bending rigidity and surface tension of the membrane, respectively [17]. The latter can be in general a function of the local coordinates $\xi^\alpha$. Here, $V_\mathrm{m}$ is the mean-field free-energy associated with membrane components in the dilute limit (expanded to lowest order in the mass fractions of each membrane component):

$$V_\mathrm{m} = \frac{k_BT}{b_0}\Big[\Phi_0\big(\ln\Phi_0 + \mathcal{E}_0\big) + \sum_{k,n}\phi_n^k\big(\ln\phi_n^k + \mathcal{E}_n^k\big)\Big], \tag{S.45}$$

which contains the entropy of mixing terms, as well as the linear order energetic penalties for each membrane component. Herein, $\mathcal{E}_0$ and $\mathcal{E}_n^k$ are dimensionless constants, representing the standard chemical potential, measured in units of $k_BT$ (thermal energy), $b_0$ is the characteristic area of the membrane components, commensurate with the coarse-graining length-scale in our model ($\sim$50 nm). The local spontaneous curvature $C_\mathrm{m}$ gives the interaction term via the coupling to curvature, namely

$$C_\mathrm{m} = \Phi_0\mathcal{C}_0 + \sum_{k,n}\phi_n^k\mathcal{C}_n^k = \mathcal{C}_0 + \sum_{k,n}\phi_n^k\big(\mathcal{C}_n^k - \mathcal{C}_0\big), \tag{S.46}$$

where $\mathcal{C}_0$ and $\mathcal{C}_n^k$ are coefficients which specify the coupling strengths to curvature for each membrane component and their respective internal states. Moreover, the subsequent equality in (S.46) follows from Eq. (S.15), and $\mathcal{C}_0$ can be interpreted as a global spontaneous curvature of the membrane, often included in the Helfrich theory [19]. Strictly positive values of $\mathcal{C}_0$ and $\mathcal{C}_n^k$ indicates a tendency to enrich regions of negative curvature with the corresponding membrane components. Notice that the standard Helfrich free-energy functional [19] can also contain a Gaussian curvature term of the form $\int_\mathcal{M}\bar{\kappa}K\,\mathrm{d}S$ as given by a quadratic expansion in the geometric invariants of the surface [17]. Nevertheless, this term can be ignored here, as its associated energy contributes to the overall energy only through changes in the topology of the surface; a result that is known as the *Gauss–Bonnet theorem* [21]. For the sake of completeness, the Gaussian energy contribution can be easily computed for $\mathcal{M}$ (*i.e.* a closed surface of genus zero), which is found to be the constant energy $4\pi\bar{\kappa}$.

Using Eq. (S.43), the membrane surface stresses can be thus calculated from the free-energy density $F$, which is a function of the metric $g_{\alpha\beta}$ and the curvature tensor $b_{\alpha\beta}$. However, the construction of the free-energy functional in Eq. (S.44) has implicitly assumed that $\mathcal{F}$ is invariant under coordinate transformations. The free-energy density $F$ depends on the tensors $g_{\alpha\beta}$ and $b_{\alpha\beta}$ only via the mean curvature $H$. Hence, by chain rule, we have that

$$\frac{\partial F}{\partial g_{\alpha\beta}} = -\frac{1}{2}\,b^{\alpha\beta}\frac{\partial F}{\partial H} \quad \text{and} \quad \frac{\partial F}{\partial b_{\alpha\beta}} = \frac{1}{2}\,g^{\alpha\beta}\frac{\partial F}{\partial H}, \tag{S.47}$$

which follow from the derivative identities $\frac{\partial H}{\partial g_{\alpha\beta}} = -\frac{1}{2}b^{\alpha\beta}$ and $\frac{\partial H}{\partial b_{\alpha\beta}} = \frac{1}{2}g^{\alpha\beta}$. The derivative of $F$ with respect to the mean curvature $H$ is obtained by using Eq. (S.44),

$$\frac{\partial F}{\partial H} = 4\kappa H\left(1 + \frac{1}{2}\frac{\partial C_\mathrm{m}}{\partial H}\right) + 2\kappa C_\mathrm{m} + \frac{\partial V_\mathrm{m}}{\partial H}, \tag{S.48}$$

then the in-plane stress tensor $\sigma_0^{\alpha\beta}$ readily follows:

$$\sigma_0^{\alpha\beta} = Fg^{\alpha\beta} - \frac{\partial F}{\partial H}\,b^{\alpha\beta}, \tag{S.49}$$

by applying the identity $\partial g/\partial g_{\alpha\beta} = g\,g^{\alpha\beta}$. Similarly, the tensor $M_0^{\alpha\beta}$, as defined in Eq. (S.43), is given by

$$M_0^{\alpha\beta} = \frac{1}{2}\frac{\partial F}{\partial H}\,g^{\alpha\beta}. \tag{S.50}$$

As a result, $T_0^{\alpha\beta}$ and $N_0^\alpha$ are found to be

$$T_0^{\alpha\beta} = Fg^{\alpha\beta} - \frac{1}{2}\frac{\partial F}{\partial H}\,b^{\alpha\beta}, \quad \text{and} \quad N_0^\alpha = -\frac{1}{2}\left(\frac{\partial F}{\partial H}\right)^{;\alpha}. \tag{S.51}$$

In addition to the passive contribution to the mechanical stresses, the free-energy density $F$ also characterizes the chemical potential of all membrane species at a specific mass fraction. In general, this is defined as the functional derivative of $\mathcal{F}$ with respect to the mass fractions, but in this particular case it is simply given by the corresponding partial derivative of $F$. Since $\phi_n^k$ and $\Phi_0$ are not independent, being coupled through Eq. (S.15), only the change of the chemical potentials with respect to the background component,

$$\boxed{\mu_n^k = \frac{\partial F}{\partial \phi_n^k} - \frac{\partial F}{\partial \Phi_0},} \tag{S.52}$$

gives the local free-energy cost per-unit-area of adding (or removing) a particle of species $k$ within the $n$-th state.

A local chemical potential imbalance leads to the gradients in $\mu_n^k$, which result in the migration of membrane components in order to decrease the total free-energy of the system. Thus, the diffusive currents $j_n^{k\,\alpha}$ must be directly related to the gradients $\mu_n^{k\,;\alpha}$. For general irreversible processes in the linear regime (*i.e.* close to equilibrium), the Onsager's principle holds [22], which establishes a linear relationship between the generalized forces and their corresponding thermodynamic fluxes [16]. For isothermal systems, this principle is shown to reduce to the principle of least energy dissipation [23]; namely, the system evolves dynamically so that it minimizes the competition between dissipation and energy release.

Therefore, this can be cast as a variational principle problem, where we seek to minimize a functional that is known as the *Rayleighian*, and which is defined by

$$\mathfrak{R} = \mathfrak{D} + \dot{\mathcal{F}}, \tag{S.53}$$

with $\mathfrak{D}$ as the dissipation potential [24]. $\mathcal{F}$ is typically a function of the state variables, such as the metric tensor $g_{\alpha\beta}$, the curvature tensor $b_{\alpha\beta}$, and the mass fractions $\phi_n^k$, describing the state of the system [24]. On the other hand, $\mathfrak{D}$ is a function of the *process variables*, such as the diffusive fluxes $\boldsymbol{j}_n^k$, which characterize how the system changes its state and generates dissipation [24]. It is noteworthy to mention that the minimization procedure of the Rayleighian is carried out with respect to the process variables, rather than the state variables of the system. In general, the Rayleighian $\mathfrak{R}$ may include terms that correspond to external power sources, as well as any other constraints that the process variables must additionally satisfy [24]. Lastly, we note that, the dissipation potential $\mathfrak{D}$, in the Onsager's variational approach, is assumed to be a quadratic function of the process variables [25]. Nevertheless, this may be further generalized to any nonlinear function that satisfies the following thermodynamic condition [24]: $\frac{\mathrm{d}}{\mathrm{d}t}\mathcal{F} \leq 0$. This restricts $\mathfrak{D}$ to be any convex, positive function that becomes zero only when all of the process variables vanish identically [24].

Since the free-energy density $F \equiv F\big[g_{\alpha\beta}, b_{\alpha\beta}, \phi_n^k\big]$, we find the rate of change of the total free-energy to be

$$\dot{\mathcal{F}} = \int_{\mathcal{M}} \mathrm{d}S \left( \frac{1}{2}\sigma_0^{\alpha\beta}\, \dot{g}_{\alpha\beta} + M_0^{\alpha\beta}\, \dot{b}_{\alpha\beta} + \sum_{k,n} \mu_n^k\, \dot{\phi}_n^k \right), \tag{S.54}$$

which follows by Reynolds transport theorem (by noting that $\mathcal{M}$ is a closed surface), and together with the results in Eq. (S.43) and the definition of $\mu_n^k$ in Eq. (S.52).

In general, the in-plane membrane flow and the out-of-plane bending may also result in energy dissipation, *e.g.* the in-plane dissipation due to the two-dimensional shear and bulk viscosities of the membrane [26]. Here we neglect their contributions, since the viscosity of the ambient fluid, adjacent to the membrane, provides the dominant mechanism of dissipation in the system.

However, the associated dissipation involved in the diffusive motion of the membrane species within the background component cannot be ignored. Consequently, in order to construct the dissipation potential associated to this mechanism, we assume that the drag force $\boldsymbol{d}_n^k$ (a force per unit area) on a membrane species $k$ and internal state $n$, within an infinitesimal membrane patch (which is commensurate to the coarse-grained membrane area $b_0$) is given by $\boldsymbol{d}_n^k = -\zeta_n^k \phi_n^k \big(\boldsymbol{v}_n^k - \boldsymbol{v}\big)$, where $\zeta_n^k$ are the drag coefficients. Here, $\phi_n^k \big(\boldsymbol{v}_n^k - \boldsymbol{v}\big)$ represents the collective velocity of the respective membrane species, relative to the barycentric velocity, within a small membrane patch. Thus the rate of energy loss per unit membrane area can be written as follows: $\mathfrak{D}_n^k = \frac{1}{2}\zeta_n^k \phi_n^k \left|\boldsymbol{v}_n^k - \boldsymbol{v}\right|^2$. Since the concentration of membrane species is assumed to follow a dilute distribution of non-interacting particles, then the drag on an individual component is unaffected by the presence of the others; namely, the net local energy dissipation is proportional to the sum $\sum_{k,n} \mathfrak{D}_n^k$. As a result, we write the total dissipation potential as follows:

$$\mathfrak{D} = \int_{\mathcal{M}} \mathrm{d}S\, \frac{1}{2\rho} \sum_{k,n} \zeta_n^k \left|\boldsymbol{j}_n^k\right|^2, \tag{S.55}$$

using Eq. (S.3). The minimization of the Rayleighian $\mathfrak{R}$ with respect to each diffusive flux $\boldsymbol{j}_n^k$ gives us that

$$-\zeta_n^k\, \boldsymbol{j}_n^k = \rho\, \frac{\delta \dot{\mathcal{F}}}{\delta \boldsymbol{j}_n^k} = \rho\, \frac{\delta}{\delta \boldsymbol{j}_n^k} \left\{ \int_{\mathcal{M}} \mathrm{d}S \sum_{k,n} \mu_n^k\, \dot{\phi}_n^k \right\}. \tag{S.56}$$

As the dynamics of $\phi_n^k$ depends on their diffusive fluxes, via Eq. (S.16), we find, by integration by parts, that

$$\boxed{j_n^{k\,\alpha} = -\frac{\rho}{\zeta_n^k} \left( \frac{\mu_n^k}{\rho} \right)^{;\alpha},} \tag{S.57}$$

which linearly relates the diffusive flux with the gradient in the chemical potential if the total density $\rho\,(\xi^\alpha, t)$ is a covariantly conserved quantity, namely $\rho_{;\alpha} = 0$.

#### E. Adiabatic approximation

Henceforth, only cyclic transitions are considered for the biochemical cycles associated with fusion and fission events, as depicted in Fig. S.2. In other words, the index set $\mathfrak{I}^k_n$, whose elements are the states $m$ which directly connect to state $n$ for a given $k$-th component (fissogens or fusogens), is defined to be

$$\mathfrak{I}^k_n = \begin{cases} \{1\} & \text{if } n = 0, \\ \{0, 2, \aleph_k\} & \text{if } n = 1, \\ \{n-1, n+1\} & \text{otherwise}, \end{cases} \tag{S.58}$$

with the cyclic condition that state $1+\aleph_k$ is the same as, or maps back to, state one. Here, the state zero plays the role of an idle or inactivated state. In the case of a fusion cycle, the transition out of this inactive state physically corresponds to the arrival of a small transport vesicle, such as COPII, in the close vicinity ($< 10\,\text{nm}$) of a cellular compartment, *e.g.* a Golgi cisterna; while the transition into the idle state signifies the inactivation of the membrane components that were previously recruited at that site, *e.g.* the SNARE-complex disassembly. Similarly, in the case of a fission event, the transition out of the idle state gives the rate of association of proteins, at a point on the membrane compartment, which could initiate the fission process, such as coatomers; whereas a transition into the idle state represents the scission of a newly formed vesicle from the membrane of the mother compartment.

In the physiological regime, the characteristic times associated with the mechanochemical kinetics of the internal, cyclic transitions are much smaller (on the order of milliseconds) than the relaxation times of the membrane dynamics for wavelengths which are greater than the coarse-graining length scale ($\sim 50$ nm). Typically, we find the relaxation times of the latter to be in the range of 1–100 s (see Table I). This separation of scales allows us to consider the transitional dynamics of the internal states as an adiabatic process with respect to the shape deformations of the membrane, and the diffusion of its components within the membrane.

By using this adiabatic approximation, the densities $\rho^k_n$ can be treated as fast varying fields, adjusting rapidly to a (non-equilibrium) steady-state, which must satisfy the following flux relations:

$$\sum_{m \in \mathfrak{I}^k_n} (r^k_{nm}\rho^k_m - r^k_{mn}\rho^k_n) = 0, \ \text{ or } \sum_{m \in \mathfrak{I}^k_n} \mathbb{J}^k_{nm} = 0, \tag{S.59}$$

for all $n$ in a $k$-cycle. This follows from Eq. (S.9) in which both the inertial and diffusive terms are neglected with respect to the dynamics of the biochemical kinetics. Note that the internal currents $\mathbb{J}^k_{nm} = r^k_{nm}\phi^k_m - r^k_{mn}\phi^k_n$ must vanish identically at thermodynamic equilibrium due to the detailed balance condition. On the other hand, in a nonequilibrium steady-state the latter is no longer satisfied, with the flux currents $\mathbb{J}^k_{nm} \neq 0$ in general. Since the transitions are cyclic (see Fig. S.2), following the kinetic rules in Eq. (S.58), then Eq. (S.59) can be reduced to:

$$\begin{cases} \mathbb{J}^k_{0,1} = 0, & \text{if } n = 0, \\ \mathbb{J}^k_{1,0} + \mathbb{J}^k_{1,2} + \mathbb{J}^k_{1,\aleph_k} = 0, & \text{if } n = 1, \\ \mathbb{J}^k_{n,n-1} + \mathbb{J}^k_{n,n+1} = 0, & \text{otherwise}, \end{cases} \tag{S.60}$$

Due to the antisymmetric property, that is, $\mathbb{J}^k_{nm} = -\mathbb{J}^k_{mn}$, we can easily see that the only non-trivial solution to this recurrence equation is a constant current solution, *i.e.*

$$\boxed{\mathbb{J}^k_{n+1,n} = \mathbb{J}^k_{\circlearrowleft} \ \text{ and } \ \mathbb{J}^k_{n,n+1} = -\mathbb{J}^k_{\circlearrowleft},} \tag{S.61}$$

for all $k$, and $n \in \{1, 2, \ldots, \aleph_k\}$, with the property that the state $1 + \aleph_k$ is the same as state 1. Here, the loop flux current $\mathbb{J}^k_{\circlearrowleft} \geq 0$, such that the internal currents $\mathbb{J}^k_{nm}$ along a forward cyclic direction $(1 \to 2 \to 3 \to \cdots \to 1)$ are strictly positive, and negative otherwise. In addition to this, we must have the transitional flux currents

$$\mathbb{J}^k_{0,1} = -\mathbb{J}^k_{1,0} = 0, \tag{S.62}$$

for each of the $k$-th species. Note that if only two internal states exist, then $\mathbb{J}^k_{\circlearrowleft} = 0$, even in the absence of detailed balance. Therefore, we restrict henceforth to fusion and fission cycles with at least three internal states ($\aleph_k > 2$). The nonzero current $\mathbb{J}^k_{\circlearrowleft}$ is a non-equilibrium feature of the system which describes its steady-state distribution out of thermodynamic equilibrium (vanishing identically at equilibrium due to the detailed balance).

The mass fractions $\phi^k_n$ are determined by solving the simultaneous equations in (S.61) and (S.62), which can be rewritten in the matrix form

$$\underbrace{\begin{bmatrix} -r^k_{1,0} & r^k_{0,1} & 0 & 0 & \cdots & 0 \\ 0 & -r^k_{2,1} & r^k_{1,2} & 0 & \cdots & 0 \\ 0 & 0 & -r^k_{3,2} & r^k_{2,3} & \cdots & 0 \\ \vdots & \vdots & \vdots & \vdots & \ddots & \vdots \\ 0 & r^k_{\aleph_k,1} & 0 & 0 & \cdots & -r^k_{1,\aleph_k} \end{bmatrix}}_{=\,\mathsf{R}_k \text{ (a square matrix of order } 1+\aleph_k)} \overbrace{\begin{bmatrix} \phi^k_0 \\ \phi^k_1 \\ \phi^k_2 \\ \vdots \\ \phi^k_{\aleph_k} \end{bmatrix}}^{=\,\mathsf{f}_k} = \overbrace{\begin{bmatrix} 0 \\ \mathbb{J}^k_{\circlearrowleft} \\ \mathbb{J}^k_{\circlearrowleft} \\ \vdots \\ \mathbb{J}^k_{\circlearrowleft} \end{bmatrix}}^{=\,\mathsf{j}_k}$$

and thus succinctly expressed as follows: $\mathsf{R}_k \mathsf{f}_k = \mathsf{j}_k$. This can be easily solved by a matrix inversion, provided that $\det \mathsf{R}^k \neq 0$; namely, $\mathsf{f}_k = \mathsf{R}^{-1}_k \mathsf{j}_k$, where $\mathsf{R}^{-1}_k$ is the inverse matrix of $\mathsf{R}_k$. This yields the solution of the mass fractions $\phi^k_n$ in terms of the loop current $\mathbb{J}^k_{\circlearrowleft}$; namely,

$$\phi^k_n = \mathbb{J}^k_{\circlearrowleft} \,/\, \mathcal{R}^k_n, \tag{S.63}$$

where $\mathcal{R}^k_n$ is a coefficient that depends solely on the transition rates $r^k_{nm}$. The loop current $\mathbb{J}^k_{\circlearrowleft}$ is not an independent parameter, and its explicit form can be obtained in terms of the overall mass-fraction of the $k$-th membrane species, which we define as follows:

$$\boxed{\Phi_k = \sum_{n=0}^{\aleph_k} \phi_n^k.} \tag{S.64}$$

Therefore, the internal loop current can be rewritten as

$$\mathbb{J}_{\circlearrowleft}^k = \Phi_k \mathcal{R}_{\circlearrowleft}^k , \tag{S.65}$$

with the parameter $\mathcal{R}_{\circlearrowleft}^k$ satisfying $1/\mathcal{R}_{\circlearrowleft}^k = \sum_{n=0}^{\aleph_k} 1/\mathcal{R}_n^k$. At thermodynamic equilibrium, the parameter $\mathcal{R}_{\circlearrowleft}^k$ must vanish identically as detailed balance is being satisfied. Hence, the expression of the mass fractions $\phi_n^k$ now reads

$$\boxed{\phi_n^k = \Phi_k \mathcal{R}_{\circlearrowleft}^k / \mathcal{R}_n^k,} \tag{S.66}$$

which tells us that each of the areal fractions $\phi_n^k/\Phi_k$ is set entirely by the biochemical transition rates. Nonetheless, the net mass fractions $\Phi_k$ themselves are not determined by the kinetic rates, and they are slow dynamical variables whose dynamics will be discussed in the next section.

#### F. Dynamics of the areal densities

Within the adiabatic approximation, the total rate of mass flux over a reaction cycle, as given by Eq. (S.11), can be expressed in terms of the loop current as follows:

$$\mathbb{M} = \sum_k \mathbb{J}_{\circlearrowleft}^k \sum_{n=0}^{\aleph_k} \mathbb{M}_{n,n-1}^k. \tag{S.67}$$

This has been obtained by using the fact that the transition rates are cyclic, and invoking the microscopic reversibility, $\mathbb{M}_{nm}^k = -\mathbb{M}_{mn}^k$; namely, the rate of membrane mass added or removed, as a result of the biochemical transition $m \to n$, must have the same magnitude as the mass rate due to the $n \to m$ transition, but with opposite sign. Notice that the rate of mass flux $\mathbb{M}$ vanishes, as expected, if the system is at equilibrium. Thus, by using Eq. (S.65), $\mathbb{M}$ can be rewritten as

$$\boxed{\mathbb{M} = \sum_k \Phi_k \mathbb{M}_{\circlearrowleft}^k ,} \tag{S.68}$$

where $\mathbb{M}_{\circlearrowleft}^k$ is the flux rate of the total membrane mass over a transition cycle for the $k$-th component, namely

$$\mathbb{M}_{\circlearrowleft}^k = \mathcal{R}_{\circlearrowleft}^k \sum_{n=0}^{\aleph_k} \mathbb{M}_{n,n-1}^k, \tag{S.69}$$

whose sign is negative for $k = 1$ (fissogens) and positive for $k = 2$ (fusogens); specifically, we say that

$$\operatorname{sgn}\big(\mathbb{M}_{\circlearrowleft}^k\big) = (-1)^k, \tag{S.70}$$

with sgn$(\cdot)$ denoting the *signum* function. Hereinafter, we consider that the system is driven out of equilibrium, and consequently we have the active rates $\mathbb{M}_{\circlearrowleft}^k \neq 0$.

Moreover, we assume that the net membrane density $\rho$ is a constant (say $\bar{\rho}$) for wavelengths greater than the coarse-graining length scale in our model ($\sim 50$ nm). In other words, the membrane is assumed to be locally incompressible, and thus gradients in the total density $\rho$ are not supported at that scale. As a corollary, Eq. (S.13) leads to the following condition:

$$\boxed{\operatorname{div} \boldsymbol{v} = v^{\alpha}_{;\alpha} - 2Hv = \mathbb{M},} \tag{S.71}$$

where the identity (S.437) is used. This equation acts as an imposed local constraint for the membrane area, with the surface tension $\Sigma$ being the Lagrange multiplier.

As a result, the kinematic equation for the mass fraction $\Phi_0$ in Eq. (S.14), together with Eq. (S.57), yields

$$\dot{\Phi}_0 = (1 - \Phi_0)\,\mathbb{M} - \sum_{k,n} \frac{\Delta\mu_n^k}{\bar{\rho}\,\zeta_n^k}, \tag{S.72}$$

where $\Delta$ is the two-dimensional Laplace–Beltrami operator. On the other hand, the kinematics of the mass fractions $\Phi_k$ are obtained from Eq. (S.16), and given by

$$\dot{\Phi}_k = -\Phi_k\,\mathbb{M} + \sum_{n=0}^{\aleph_k} \frac{\Delta\mu_n^k}{\bar{\rho}\,\zeta_n^k}. \tag{S.73}$$

Notice that by summing this equation with respect to $k$, and using that $1 - \Phi_0 = \Phi_1 + \Phi_2$, we retrieve Eq. (S.72).

From the free-energy density $F$ in Eq. (S.44), the local chemical potential difference is found to be

$$\mu_n^k = 2\kappa H\big(\mathcal{C}_n^k - \mathcal{C}_0\big) + \frac{k_B T}{b_0}\Big[\big(\mathcal{E}_n^k - \mathcal{E}_0\big) + \ln\frac{\phi_n^k}{\Phi_0}\Big], \tag{S.74}$$

which must vanish identically at equilibrium. By using the form of the chemical potential in Eq. (S.74), this allows us to write the last term in Eq. (S.73) as follows:

$$\sum_{n=0}^{\aleph_k} \frac{\Delta\mu_n^k}{\bar{\rho}\,\zeta_n^k} = 2\Omega_k\,\Delta H + \gamma_k\,\Delta\Big(\ln\frac{\Phi_k}{\Phi_0}\Big), \tag{S.75}$$

where the mobility coefficients $\gamma_k$ are defined by

$$\gamma_k = \frac{k_B T}{\bar{\rho} b_0} \sum_{n=0}^{\aleph_k} \frac{1}{\zeta_n^k}, \tag{S.76}$$

and the effective constants $\Omega_k$ are given by

$$\Omega_k = \frac{\kappa}{\bar{\rho}} \sum_{n=0}^{\aleph_k} \frac{\mathcal{C}_n^k - \mathcal{C}_0}{\zeta_n^k}. \tag{S.77}$$

Accordingly, Eq. (S.73) can be rewritten as follows:

$$\boxed{\dot{\Phi}_k = -\Phi_k\,\mathbb{M} + 2\Omega_k\,\Delta H + \gamma_k\,\Delta\Big(\ln\frac{\Phi_k}{\Phi_0}\Big),} \tag{S.78}$$

where the Laplacian term, by chain rule, expands to

$$\Delta\Big(\ln\frac{\Phi_k}{\Phi_0}\Big)=\frac{\Delta\Phi_k}{\Phi_k}-\frac{\Phi_{k\,;\alpha}\Phi_k^{;\alpha}}{(\Phi_k)^2}+\frac{\Phi_{0\,;\alpha}\Phi_0^{;\alpha}}{(\Phi_0)^2}. \tag{S.79}$$

Thus, Eq. (S.78) represents the governing equation (at large scales) of the areal concentration of membrane components, *i.e.* fissogens ($k=1$) and fusogens ($k=2$).

### G. Active membrane stresses

The active surface contributions to the membrane traction force and the director traction are denoted by $\boldsymbol{T}_A$ and $\boldsymbol{M}_A$ [27]. Both of these are induced by the active forces that act in the neighboring cytosolic fluid, associated with each of the internal biochemical transitions of the fission and fusion events. The active forces are generated across a finite distance away from the mid-plane of the lipid bilayer, along a direction transverse to the membrane. The characteristic thickness of this boundary layer is on order of the coarse-graining length ($\sim$50 nm).

We write $\boldsymbol{F}_A$ to be the overall active force at position $\boldsymbol{r}(\xi^1,\xi^2,h)$, where $h$ parameterizes the normal coordinate away from the membrane, and $\xi^\alpha$ are surface coordinates of membrane compartment $\mathcal{M}$; namely, we write

$$\boldsymbol{F}_A(\boldsymbol{r})=\int_{\mathcal{M}}\mathrm{d}S\int_0^\infty \mathrm{d}h\;\boldsymbol{\mathfrak{F}}_A(\xi^\alpha,h)\,\delta\big(\boldsymbol{r}-\tilde{\boldsymbol{R}}\big), \tag{S.80}$$

where the position vector $\tilde{\boldsymbol{R}}$ is defined by

$$\tilde{\boldsymbol{R}}(\xi^1,\xi^2,h)=h\boldsymbol{n}(\xi^1,\xi^2)+\boldsymbol{R}(\xi^1,\xi^2), \tag{S.81}$$

and $\boldsymbol{\mathfrak{F}}_A(\xi^\alpha,h)$ is a force per unit volume that captures the force distribution away from the membrane. In other words, the integral in Eq. (S.80) can be seen as a mapping from the Cartesian position $\boldsymbol{r}$ to the coordinate system associated with the membrane and its height away from its surface, that is, $(\xi^1,\xi^2,h)$. The force density $\boldsymbol{\mathfrak{F}}_A$ is obtained by averaging over the volume of a coarse-graining pillbox located at position $\boldsymbol{R}$, with a height thickness $\Lambda$ that is set by the size of transport vesicles. Thus, $\mathfrak{F}^k_{\circlearrowright}$ is zero for $h>\Lambda$, being restricted to the interval $[0,\Lambda]$. In averaging over the pillbox, we assume that the distribution of forces along the tangential direction to the membrane is isotropic, with the active force density $\boldsymbol{\mathfrak{F}}_A$ primarily acting along the normal direction of the membrane. However, one may consider in general a more detailed description that models specifically the positional and orientational distribution of forces within the pillbox. Herein, we specify that the active force density

$$\boldsymbol{\mathfrak{F}}_A(\xi^\alpha,h)=\mathfrak{F}_A(\xi^\alpha,h)\boldsymbol{n}, \tag{S.82}$$

where the magnitude along the normal is given by

$$\mathfrak{F}_A(\xi^\alpha,h)=\sum_k\Phi_k\,\mathfrak{F}^k_{\circlearrowright}(\xi^\alpha,h), \tag{S.83}$$

which includes the corresponding active force distributions $\mathfrak{F}^k_{\circlearrowright}$ due to fissogens ($k=1$) and fusogens ($k=2$) over their entire cycle. Analogous to the derivation of Eq. (S.67), each force distribution can be written in terms of the loop currents:

$$\mathfrak{F}^k_{\circlearrowright}(\xi^\alpha,h)=\mathbb{J}^k_{\circlearrowright}\sum_{n=0}^{\aleph_k}\mathbb{F}^k_{n,n-1}, \tag{S.84}$$

where $\mathbb{F}^k_{nm}$ is the momentum flux corresponding to an active transition from the internal state $m$ to $n$ for the $k$-th cycle, which depends on the normal height away from the membrane. In deriving this expression, we assume that the transitions are cyclic and the microscopic reversibility $\mathbb{F}^k_{nm}=-\mathbb{F}^k_{mn}$ holds.

The active force $\boldsymbol{F}_A$ acts in the neighboring cytosolic fluid, whose momentum balance equation is given by

$$\boldsymbol{\nabla}\cdot\boldsymbol{\Omega}_+ +\boldsymbol{F}_A=\boldsymbol{0}, \tag{S.85}$$

where $\boldsymbol{\Omega}_+$ is the stress associated with the ambient fluid external to the membrane compartment (see forthcoming section I, where the bulk fluid stresses are discussed in more details). As written in Eq. (S.85), $\boldsymbol{F}_A$ appears as a local momentum source; however, globally we must demand that there is no production of momentum arising from the term $\boldsymbol{F}_A$ (for every force we must have an equal and opposite counter force). This condition is imposed by requiring that

$$\int_0^\Lambda \mathrm{d}h\;\mathfrak{F}^k_{\circlearrowright}(\xi^\alpha,h)=0, \tag{S.86}$$

at each position along the membrane. This is equivalent to demanding the sum of all forces within the pillbox is zero. Thus, the first and second moment with respect to the height $h$ are the first nontrivial contributions.

For illustrative purposes, we consider a simple point-force dipolar model:

$$\mathfrak{F}^k_{\circlearrowright}(h)=f^k_0\,[\delta(h-\Lambda)-\delta(h)], \tag{S.87}$$

where $f^k_0$ gives the momentum flux distribution over a full cycle for each $k$ component. Note that the monopole of this force distribution is identically zero, by construction. This is a simple model that captures only the magnitude of the momentum transfer and the characteristic transverse distance of the point-force dipole. In this simple model, the first and second moments of the force density are given by

$$\mathbb{P}^k_{\circlearrowright}=\int_0^\Lambda h\,\mathfrak{F}^k_{\circlearrowright}(h)\,\mathrm{d}h=f^k_0\Lambda \quad\text{and} \tag{S.88}$$

$$\mathbb{Q}^k_{\circlearrowright}=\int_0^\Lambda h^2\,\mathfrak{F}^k_{\circlearrowright}(h)\,\mathrm{d}h=f^k_0\Lambda^2, \tag{S.89}$$

respectively. Note that the sign of $\mathbb{P}^k_{\circlearrowright}$ and $\mathbb{Q}^k_{\circlearrowright}$ is determined in this case by the sign of $f^k_0$. This is specific for this model and it is not true in general.

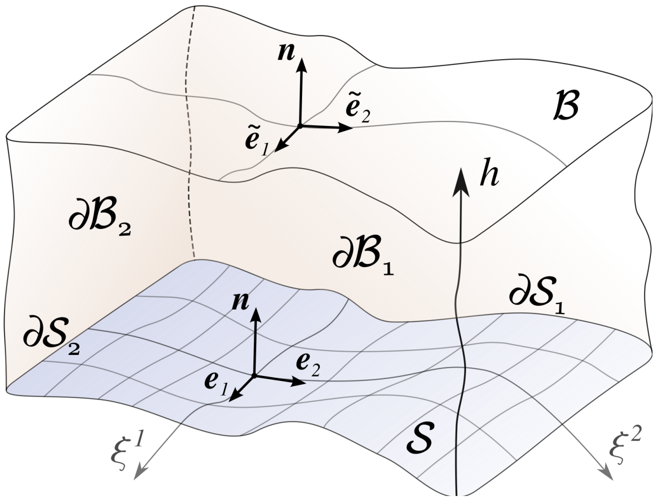


FIG. S.3. Diagram of a three-dimensional cell that includes the ambient fluid above a membrane surface patch $\mathcal{S}$, where the side boundaries are labeled by $\partial\mathcal{S}_\alpha$ with $\alpha \in \{1, 2\}$. Each point above the membrane is parameterized by the internal curvilinear coordinates $\xi^\alpha$ and the height $h$ along the surface normal $\boldsymbol{n}$. Also, the side faces of the volume cell $\mathcal{B}$, that are associated with the boundaries $\partial\mathcal{S}_\alpha$, are indicated by $\partial\mathcal{B}_\alpha$. Note that local tangent basis at height $h$ above the membrane surface are slightly different than those on $\mathcal{S}$, due to the local change in curvature $b_{\alpha\beta}$; namely, $\tilde{\boldsymbol{e}}_\alpha = \boldsymbol{e}_\beta\left(\delta^\beta_\alpha - h\, b^\beta_\alpha\right)$.

As another illustrative example, we consider the force density $\mathfrak{F}^k_\circlearrowright$ to be an expansion in $h/\Lambda$, that is,

$$\mathfrak{F}^k_\circlearrowright(h) = f^k_0 + f^k_1\left[\frac{h}{\Lambda}\right] + f^k_2\left[\frac{h}{\Lambda}\right]^2, \tag{S.90}$$

where $f^k_0$, $f^k_1$ and $f^k_2$ are some constants. By imposing that force monopoles vanish, from Eq. (S.86), we find

$$f^k_0 = -\frac{1}{6}\left(3f^k_1 + 2f^k_2\right), \tag{S.91}$$

which yields the first moment

$$\mathbb{P}^k_\circlearrowright = \frac{1}{12}(f^k_1 + f^k_2)\Lambda^2, \tag{S.92}$$

while the second moment is

$$\mathbb{Q}^k_\circlearrowright = \frac{1}{180}(15f^k_1 + 16f^k_2)\Lambda^3. \tag{S.93}$$

In this case, the sign of $\mathbb{P}^k_\circlearrowright$ can be different from the sign of $\mathbb{Q}^k_\circlearrowright$, provided that $-1 < f^k_2/f^k_1 < -15/16$.

Returning to Eq. (S.85), we can see that $\boldsymbol{F}_A$ can be written as the divergence of a bulk stress. Solving the hydrodynamics of the bulk fluid in such setup, which we shall refer hereinafter as the semi-microscopic description, is in general a difficult problem. However, we can use the fact that $\Lambda$ is much smaller than the size of the compartment. This allows us to reformulate Eq. (S.85) as a homogeneous problem, namely

$$\boldsymbol{\nabla}\cdot\boldsymbol{\Omega}_+ = \boldsymbol{0}, \tag{S.94}$$

with the active forces now entering as a boundary condition at membrane surface. In other words, we project the corresponding three-dimensional force distribution onto the membrane surface [28]. The general relationship between such a semi-microscopic model, as one described in Eq. (S.85), to an effective two-dimensional description has been carried out formally in Refs. [28–30], in the context of active membranes driven by ion pumps [31].

The connection between the two formulations follows from the fact that the net forces must be the same in both descriptions, once integrating their associated stress tensors. As a result, the integrated excess stress $\boldsymbol{\Omega}_A$ in the bulk due to the activity alone must equal the integrated traction force $\boldsymbol{T}_A$, that is,

$$\int_{\partial\mathcal{B}_\alpha} \mathrm{d}\tilde{S}\;\tilde{\boldsymbol{\nu}}\cdot\boldsymbol{\Omega}_A = \int_{\partial\mathcal{S}_\alpha} \mathrm{d}l\;\nu_\alpha\,\boldsymbol{T}^\alpha_A, \tag{S.95}$$

where $\partial\mathcal{B}_\alpha$ parameterizes the two independent side faces of a three-dimensional domain $\mathcal{B}$ whose base is given by a membrane surface patch $\mathcal{S}$, as illustrated in Fig. S.3. Concretely, the excess stress $\boldsymbol{\Omega}_A$ is defined as the bulk fluid stress tensor in the semi-microscopic description subtracting out the bulk fluid stress tensor in the corresponding two-dimensional formulation, with the boundary conditions for Eqs. (S.85) and (S.94) chosen such that the solutions for the bulk fluid are identical in the two formulations far away from the membrane at normal heights greater than boundary layer thickness $\Lambda$.

The set of points of $\partial\mathcal{B}_\alpha$ is given by

$$\partial\mathcal{B}_\alpha = \left\{\tilde{\boldsymbol{R}}(\xi^1, \xi^2, h)\;\middle|\; h \geq 0 \text{ and } \xi^{3-\alpha} \in \mathcal{S}\right\}, \tag{S.96}$$

with the other coordinate kept as a free parameter. Note that this becomes identical to the boundary curve $\partial\mathcal{S}_\alpha$ when we set the normal height $h = 0$, by construction. By parameterizing the curve $\partial\mathcal{S}_\alpha$ by $\lambda$, that is, we consider the function $\boldsymbol{R}\left[\xi^1(\lambda), \xi^2(\lambda)\right]$, then the unit vector $\tilde{\boldsymbol{\nu}}$ normal to the side surface $\partial\mathcal{B}_\alpha$ is given by

$$\tilde{\boldsymbol{\nu}} = \left(\tilde{\boldsymbol{e}}_\beta\tfrac{\mathrm{d}\xi^\beta}{\mathrm{d}\lambda}\times\boldsymbol{n}\right)\left|\tilde{\boldsymbol{e}}_\beta\tfrac{\mathrm{d}\xi^\beta}{\mathrm{d}\lambda}\right|^{-1}, \tag{S.97}$$

whereas the tangent basis are found to be

$$\tilde{\boldsymbol{e}}_\alpha = \tilde{\boldsymbol{R}}_{,\alpha} = \boldsymbol{e}_\beta\left(\delta^\beta_\alpha - h\,b^\beta_\alpha\right). \tag{S.98}$$

Furthermore, the area element $\mathrm{d}\tilde{S}$ can be expressed in this parametrization as $\mathrm{d}\tilde{S} = \mathrm{d}h\,\mathrm{d}\lambda\left|\tilde{\boldsymbol{e}}_\alpha\tfrac{\mathrm{d}\xi^\alpha}{\mathrm{d}\lambda}\right|$, leading to

$$\mathrm{d}\tilde{S}\;\tilde{\boldsymbol{\nu}} = \mathrm{d}h\,\mathrm{d}\lambda\;\varepsilon_{\alpha\gamma}\,\boldsymbol{e}^\alpha\left(\delta^\gamma_\beta - h\,b^\gamma_\beta\right)\frac{\mathrm{d}\xi^\beta}{\mathrm{d}\lambda}, \tag{S.99}$$

where the identity $\varepsilon_{\alpha\gamma}\boldsymbol{e}^\alpha = \boldsymbol{e}_\gamma\times\boldsymbol{n}$ is used, with $\varepsilon_{\alpha\gamma}$ as the Levi-Civita tensor. Similarly, the line element is

$$\mathrm{d}l\,\nu_\alpha = \varepsilon_{\alpha\beta}\frac{\mathrm{d}\xi^\beta}{\mathrm{d}\lambda}. \tag{S.100}$$

By substituting these two expressions into Eq. (S.95), then we find that the active stress vectors

$$\boldsymbol{T}_A^{\alpha} = \int_0^{\infty} \mathrm{d}h\,(\boldsymbol{e}_\beta \cdot \boldsymbol{\Omega}_A)\left[g^{\alpha\beta} - h\left(2Hg^{\alpha\beta} - b^{\alpha\beta}\right)\right]. \tag{S.101}$$

By using Eq. (S.435), we recognize the term in the round brackets, pre-multiplying $h$, as the cofactor of the curvature tensor, *i.e.* $\bar{b}^{\alpha\beta} = 2Hg^{\alpha\beta} - b^{\alpha\beta}$.

To find the form of the stress vectors in Eq. (S.101), we need to determine the excess stress, $\boldsymbol{\Omega}_A$, by solving the hydrodynamics in the semi-microscopic description. The latter is a difficult problem to solve in practice, with $\boldsymbol{\Omega}_A$ being generally unattainable in exact form [29, 30], and thus Eq. (S.101) may not seem to be that useful at first. However, one could use this result to analytically obtain a moment expansion of the stress vectors $\boldsymbol{T}_A^{\alpha}$ in terms of the first and second moments of the force distribution $\mathfrak{F}_A(h)$, as described subsequently.

From the linearity of the problem (assuming a Stokes flow for the ambient fluid, see forthcoming section), the contribution to the active force $\boldsymbol{F}_A$ that solely arises from an infinitesimal volume between $h$ and $h+\delta h$, and associated with an infinitesimal membrane area $\delta S$ (as depicted in Fig. S.4), is given by

$$\delta \boldsymbol{F}_A = \boldsymbol{n}\,\mathfrak{F}_A(h)\,\delta h\,\delta S. \tag{S.102}$$

By neglecting gradient contributions in the force distribution $\mathfrak{F}_A(h)$ and the mean curvature $H$, the effect of the active force $\delta\boldsymbol{F}_A$ to the excess stress tensor $\delta\boldsymbol{\Omega}_A$, associated with a small volume $\delta\tilde{V}$ between the heights $h$ and $h+\delta h$, and of area $\delta\tilde{S} = \delta S\left(1 - 2hH + h^2K\right)$, is only a small change $\delta p$ in the fluid pressure which balances the active forces. Namely, we can write that $\delta\boldsymbol{\Omega}_A = -\delta p\,\boldsymbol{I}$, where $\boldsymbol{I}$ is the identity tensor and the pressure jump is found to be $\delta p = -\delta h\left|\delta\boldsymbol{F}_A/\delta\tilde{V}\right|$ for heights between the membrane surface and $h$, and zero otherwise (that is, in the far field, the bulk fluid stresses are identical in both descriptions). Thus, to lowest order in the mean curvature, we have that

$$\delta\boldsymbol{\Omega}_A = \delta h\;\mathfrak{F}_A(h)\,(1 + 2hH)\,\boldsymbol{I} + \mathcal{O}[H^2], \tag{S.103}$$

which follows from expanding the volume $\delta\tilde{V} = \delta h\,\delta\tilde{S}$.

From Eq.(S.101), the infinitesimal stress vector $\delta\boldsymbol{T}_A^{\alpha}$ due to the jump in pressure $\delta p$ is given by

$$\delta\boldsymbol{T}_A^{\alpha} = (\boldsymbol{e}_\beta \cdot \delta\boldsymbol{\Omega}_A)\int_0^{h} \mathrm{d}h'\left(g^{\alpha\beta} - h'\,\bar{b}^{\alpha\beta}\right), \tag{S.104}$$

where we note that the excess stress in the bulk is given by $\delta\boldsymbol{\Omega}_A$ if $h' \in [0, h]$ and zero otherwise [29]. Thus, by substituting the expression of $\delta\boldsymbol{\Omega}_A$ from Eq. (S.103), the in-plane part of the active stress, *i.e.* $\delta T_A^{\alpha\beta} = \delta\boldsymbol{T}_A^{\alpha}\cdot\boldsymbol{e}^{\beta}$, is expressed as a cubic polynomial in $h$ that pre-multiplies

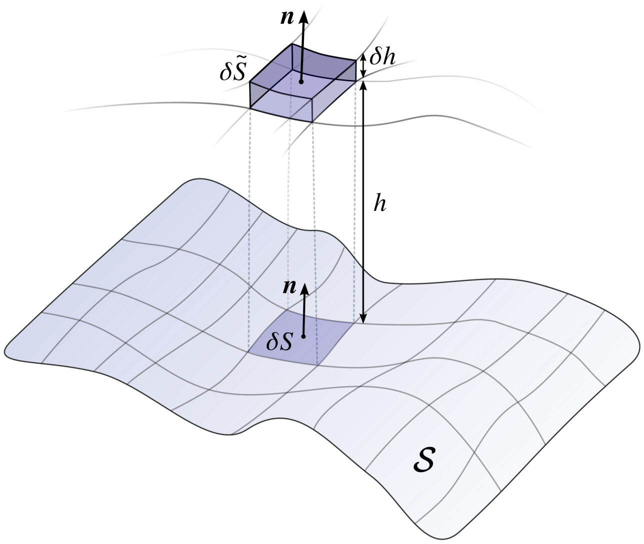


FIG. S.4. Schematic of an infinitesimal volume $\delta\tilde{V} = \delta h\,\delta\tilde{S}$ of the ambient fluid, which is positioned away from the membrane between the heights $h$ and $h+\delta h$, along the unit vector $\boldsymbol{n}$ normal to the membrane $\mathcal{S}$. Since the surface $\mathcal{S}$ is locally curved, with a mean curvature $H$, then the area element $\delta\tilde{S}$ at height $h$ is thus different than its associated membrane area $\delta S$; namely, we have that $\delta\tilde{S} \simeq \delta S\,(1 - 2hH)$.

the force density $\mathfrak{F}_A(h)$. By integrating over the small segment $\delta h$, we have the following moment expansion:

$$T_A^{\alpha\beta} = \int_0^{\Lambda} \mathrm{d}h\,\mathfrak{F}_A(h)\left[h\,g^{\alpha\beta} + h^2\left(Hg^{\alpha\beta} + \frac{1}{2}\,b^{\alpha\beta}\right)\right], \tag{S.105}$$

where we truncate to second-order in the moments of the force distribution $\mathfrak{F}_A$ for self-consistency. At third order in this expansion, we expect the gradients in the mean curvature and the force density to contribute to the in-plane stress, and thus Eq. (S.103) is no longer valid [29].

As a result, the active in-plane stress can be written in terms of the first moment,

$$\boxed{\mathbb{P} = \int_0^{\Lambda} \mathrm{d}h\,\left[h\,\mathfrak{F}_A(h)\right] = \sum_k \Phi_k\,\mathbb{P}^k_{\circlearrowleft}\,,} \tag{S.106}$$

as well as in terms of the second moment,

$$\boxed{\mathbb{Q} = \int_0^{\Lambda} \mathrm{d}h\,\left[h^2\,\mathfrak{F}_A(h)\right] = \sum_k \Phi_k\,\mathbb{Q}^k_{\circlearrowleft}\,,} \tag{S.107}$$

where $\mathbb{P}^k_{\circlearrowleft} = \int h\,\mathfrak{F}^k_{\circlearrowleft}(h)\,\mathrm{d}h$, and $\mathbb{Q}^k_{\circlearrowleft} = \int h^2\,\mathfrak{F}^k_{\circlearrowleft}(h)\,\mathrm{d}h$, as previously defined. Therefore, the in-plane stress from Eq. (S.105) can be expressed as follows [29, 30]:

$$T_A^{\alpha\beta} = g^{\alpha\beta}\,\mathbb{P} + \frac{1}{2}\left(2Hg^{\alpha\beta} + b^{\alpha\beta}\right)\mathbb{Q}. \tag{S.108}$$

$\mathbb{P}$ and $\mathbb{Q}$ are independent moments, and their sign may not be the same in general. However, within the simple point-force model, $\mathrm{sgn}\big[\mathbb{P}^k_{\circlearrowleft}\big] = \mathrm{sgn}\big[\mathbb{Q}^k_{\circlearrowleft}\big]$, which is determined by the sign of $f^k_0$ in Eq. (S.87). The latter is defined in terms of the momentum flux which is locally transferred to the *mother* membrane from a secretory vesicle, and thus we expect its sign to be negative for the fission events ($k = 1$), as it takes momentum away from the membrane, and positive for the fusion events ($k = 2$), which delivers momentum from the transport vesicle. This gives us that

$$\mathrm{sgn}\big(\mathbb{P}^k_{\circlearrowleft}\big) = \mathrm{sgn}\big(\mathbb{Q}^k_{\circlearrowleft}\big) = (-1)^k. \tag{S.109}$$

Since $\mathbb{P}$ renormalizes the surface tension, cf. Eq. (S.123), the above sign convention of $\mathbb{P}^k_{\circlearrowleft}$ leads to a reduction of surface tension by the fusogens (as more area is being added), as well as an increase of surface tension in the case of fissogens (as membrane area is removed), which is what we heuristically expect based on *in vitro* membrane experiments [32, 33]. Nevertheless, the sign of the second moment $\mathbb{Q}^k_{\circlearrowleft}$ might not be in general equal to the sign of $\mathbb{P}^k_{\circlearrowleft}$, as we saw in Eqs. (S.92) and (S.93).

As analogous to Eq. (S.101), the active contribution to the director traction $\boldsymbol{M}^\alpha_A$ can be computed via the excess stress tensor $\boldsymbol{\Omega}_A$, which is found to be [28]:

$$\boldsymbol{M}^\alpha_A = \int_0^\infty \mathrm{d}h\, h\,(\boldsymbol{e}_\beta \cdot \boldsymbol{\Omega}_A)\,\big(g^{\alpha\beta} - h\bar{b}^{\alpha\beta}\big), \tag{S.110}$$

which follows from requiring that the overall torques that result from integrating the corresponding angular stress tensors in the two descriptions must be the identical, *i.e.*

$$\int_{\partial\mathcal{B}_\alpha} \tilde{\boldsymbol{R}} \times (\tilde{\boldsymbol{\nu}} \cdot \boldsymbol{\Omega}_A)\,\mathrm{d}\tilde{S} = \int_{\partial\mathcal{S}_\alpha} \nu_\alpha\,(\boldsymbol{n} \times \boldsymbol{M}^\alpha_A + \boldsymbol{R} \times \boldsymbol{T}^\alpha_A)\,\mathrm{d}l.$$

By employing the same strategy as used to derive the in-plane stress in Eq. (S.105), the active bending moment tensor $M^{\alpha\beta}_A = -\boldsymbol{M}^\alpha_A \cdot \boldsymbol{e}^\beta$ can be determined as follows:

$$\boxed{M^{\alpha\beta}_A = -\int_0^\Lambda \mathrm{d}h\, \frac{h^2}{2}\,\mathfrak{F}_A(h)\,g^{\alpha\beta} = -\frac{1}{2}\,\mathbb{Q}\,g^{\alpha\beta}.} \tag{S.111}$$

In addition, due to the conservation of angular momentum, cf. Eq. (S.29), the active out-of-plane shear stress is thus found to be $N^\alpha_A = -M^{\alpha\beta}_{A\,;\beta} = \big(\frac{1}{2}\mathbb{Q}\big)^{;\alpha}$, whereas the symmetric in-plane stress tensor due to activity, which is defined by $\sigma^{\alpha\beta}_A = T^{\alpha\beta}_A - b^\beta_\gamma M^{\gamma\alpha}_A$, reduces to

$$\boxed{\sigma^{\alpha\beta}_A = \mathbb{P}\,g^{\alpha\beta} + \mathbb{Q}\left(Hg^{\alpha\beta} + b^{\alpha\beta}\right).} \tag{S.112}$$

This readily shows how the active force distribution can contribute to the total in-plane stress of the membrane, through its nonzero first and second moments.

### H. Membrane equations of motion

The free-energy density $F$ in Eq. (S.44) in the adiabatic approximation can be rewritten by substituting the form of the mass-fractions $\phi^k_n$ from Eq. (S.63), namely

$$F = \Sigma_{\text{eff}}(\Phi_k) + 2\kappa C_{\text{eff}}(\Phi_k)H + 2\kappa H^2. \tag{S.113}$$

Here, the effective surface tension $\Sigma_{\text{eff}}$ can be written as

$$\Sigma_{\text{eff}} = \bar{\Sigma} + \frac{k_BT}{b_0}\left[\ln\Phi_0 + \sum_k \Phi_k\left(E_k + \ln\frac{\Phi_k}{\Phi_0}\right)\right], \tag{S.114}$$

where we define the constants $\bar{\Sigma} = \Sigma + \frac{k_BT}{b_0}\mathcal{E}_0$, and

$$E_k = \sum_{n=0}^{\aleph_k} \frac{\mathcal{R}^k_{\circlearrowleft}}{\mathcal{R}^k_n}\left[\mathcal{E}^k_n - \mathcal{E}_0 + \ln\frac{\mathcal{R}^k_{\circlearrowleft}}{\mathcal{R}^k_n}\right]. \tag{S.115}$$

Since the dynamics of $\Phi_k$ depends on the active flux rates $\mathbb{M}^k_{\circlearrowleft}$, $\Sigma_{\text{eff}}$ represents a renormalized surface tension by the active addition and removal of membrane material.

In addition, the effective spontaneous curvature is found to be $C_{\text{eff}} = \mathcal{C}_0 + \sum_k \Phi_k C_k$, where $C_k$ is the local spontaneous curvature associated with each $k$, *i.e.*

$$C_k = \sum_{n=0}^{\aleph_k} \frac{\mathcal{R}^k_{\circlearrowleft}\,(\mathcal{C}^k_n - \mathcal{C}_0)}{\mathcal{R}^k_n}. \tag{S.116}$$

The effective term $C_{\text{eff}}$ represents a renormalized spontaneous curvature by the membrane fluxes $\mathbb{M}^k_{\circlearrowleft}$.

By using the free-energy in Eq. (S.113), the in-plane stress tensor $\sigma^{\alpha\beta}_0$ and the bending moment tensor $M^{\alpha\beta}_0$ are determined from Eq. (S.49) and (S.50), respectively. Furthermore, the overall in-plane (symmetrized) stress tensor $\sigma^{\alpha\beta} = \sigma^{\alpha\beta}_0 + \sigma^{\alpha\beta}_A$ is found to be

$$\sigma^{\alpha\beta} = (F + \mathbb{P} + H\mathbb{Q})\,g^{\alpha\beta} + \left(\mathbb{Q} - \frac{\partial F}{\partial H}\right)b^{\alpha\beta}, \tag{S.117}$$

whereas the moment bending tensor $M^{\alpha\beta}$ is given by

$$M^{\alpha\beta} = -\frac{1}{2}\left(\mathbb{Q} - \frac{\partial F}{\partial H}\right)g^{\alpha\beta}. \tag{S.118}$$

This allows us to obtain the equations of motion, as derived in Eq. (S.30) and (S.31), along the normal and tangential direction of the membrane, respectively.

As a result, the force balance equation along tangential direction is found to be

$$\begin{aligned}\rho\dot{\boldsymbol{v}} \cdot \boldsymbol{e}_\alpha = f_\alpha &+ \Sigma_{\text{eff}\,;\alpha} + 2\kappa H C_{\text{eff}\,;\alpha} \\ &+ \mathbb{P}_{;\alpha} + H\,\mathbb{Q}_{;\alpha} + 2\,\mathbb{Q}H_{;\alpha}\,,\end{aligned} \tag{S.119}$$

by substituting Eq. (S.113) into (S.117) and (S.118), and then expanding out the terms in Eq. (S.31). Moreover, by using the definitions in (S.106) and (S.107), as well as Eqs. (S.114–S.116), this can be further rewritten as

$$\rho\dot{\boldsymbol{v}}\cdot\boldsymbol{e}_\alpha = \sum_k \Phi_{k;\alpha}\left[\frac{k_BT}{b_0}\left(E_k+\ln\frac{\Phi_k}{\Phi_0}\right)+2H\left(\kappa C_k+\frac{\mathbb{Q}^k_{\circlearrowleft}}{2}\right)\right] + f_\alpha + \Sigma_{;\alpha} + \sum_k \left(\Phi_{k;\alpha}\mathbb{P}^k_{\circlearrowleft} + 2\Phi_k\mathbb{Q}^k_{\circlearrowleft}H_{;\alpha}\right). \quad (S.120)$$

Similarly, the force-balance along the normal direction to the membrane can be found via Eq. (S.30) as follows:

$$\rho\dot{\boldsymbol{v}}\cdot\boldsymbol{n} = f + 2H(\Sigma_{\text{eff}}+\mathbb{P}) - 2\kappa\left[\Delta H + 2H\left(H^2-K\right)\right] - \kappa(\Delta C_{\text{eff}} - 2KC_{\text{eff}}) + \frac{\Delta\mathbb{Q}}{2} + \mathbb{Q}\left(4H^2-K\right), \quad (S.121)$$

which can be subsequently expanded as

$$\rho\dot{\boldsymbol{v}}\cdot\boldsymbol{n} = f - 2\kappa\left[\Delta H + 2H\left(H^2-K\right)\right] + 2\kappa K\mathcal{C}_0 + 2H\left\{\bar{\Sigma} + \sum_k\Phi_k\left[\mathbb{P}^k_{\circlearrowleft} + \frac{k_BT}{b_0}\left(E_k+\ln\frac{\Phi_k}{\Phi_0}\right)\right]\right\} + \sum_k\Phi_k\left[2\kappa KC_k + \left(4H^2-K\right)\mathbb{Q}^k_{\circlearrowleft}\right] - \sum_k\Delta\Phi_k\left(\kappa C_k - \mathbb{Q}^k_{\circlearrowleft}/2\right), \quad (S.122)$$

by employing the results in Eqs. (S.114–S.116). From Eq. (S.121) and (S.122), the term which multiplies $2H$ represents a surface tension renormalized by both the active first moment $\mathbb{P}^k_{\circlearrowleft}$, as well as the active membrane flux $\mathbb{M}^k_{\circlearrowleft}$; namely we can write the activity-renomalized surface tension by

$$\boxed{\Sigma_A = \bar{\Sigma} + \sum_k\Phi_k\left[\mathbb{P}^k_{\circlearrowleft} + \frac{k_BT}{b_0}\left(E_k+\ln\frac{\Phi_k}{\Phi_0}\right)\right].} \quad (S.123)$$

By the sign convention in Eq. (S.109), $\text{sgn}(\mathbb{P}^k_{\circlearrowleft}) = (-1)^k$, we see that the first moment $\mathbb{P}^1_{\circlearrowleft}$ (associated with fissogens) leads to a reduction in the surface tension, while $\mathbb{P}^2_{\circlearrowleft}$ (associated with fusogens) increases it.

Similarly, we identify an activity-renormalized spontaneous curvature, namely

$$\boxed{C_A = \mathcal{C}_0 + \sum_k\Phi_k\left(C_k - \frac{\mathbb{Q}^k_{\circlearrowleft}}{2\kappa}\right),} \quad (S.124)$$

which is renormalized by both the active membrane flux $\mathbb{M}^k_{\circlearrowleft}$ through $\Phi_k$, and the active second moment $\mathbb{Q}^k_{\circlearrowleft}$. By the sign convention (S.109), this implies that spontaneous curvature increases (decreases) during fission (fusion), segregating of fissogens to highly curved regions.

#### I. Coupling to the ambient fluid

On either sides of the compartment, its closed membrane surface $\mathcal{M}$ is in contact with an ambient fluid. In general, this surrounding fluid provides the dominant dissipative mechanism, at large wavelengths, for the relaxation of shape deformations [34], which corresponds to length scales larger than the coarse-graining length in our model of the membrane system (*i.e.* 50–100 nm). Below this scale, the bulk fluid has an insignificant contribution, and the membrane dynamics is strongly dominated by the active mechanism of recycling, which leads to rapid conformational changes in the internal membrane constituents via the biochemical reaction cycles. Typically, under a submicron size, the friction between the monolayers of the membrane becomes important, and even under smaller scales, the shear viscosity of the membrane within each layer may also become significant [35]. Hence, based on this hierarchy of dissipative processes, at large length scales, the membrane is hydrodynamically coupled to the bulk fluid through interfacial conditions on both the interior and exterior faces of the membrane [36]. The fluid velocity along the normal surface pushes locally the membrane, resulting in a shape deformation, whereas the fluid motion along the tangential direction of the surface leads to flows within the membrane.

Here, we assume that the bulk fluid, which surrounds the membrane, is locally incompressible and obeys the Stokes equation. This is motivated by the small Reynolds number in our system (Re $\lessapprox 10^{-4}$), with the advective inertial terms being negligible, compared with the viscous forces. Thus, the fluid motion is governed by

$$\boxed{\eta\boldsymbol{\nabla}^2\boldsymbol{V}_\pm = \boldsymbol{\nabla}p_\pm, \quad \text{and} \quad \boldsymbol{\nabla}\cdot\boldsymbol{V}_\pm = 0,} \quad (S.125)$$

where the Laplacian $\boldsymbol{\nabla}^2 = \boldsymbol{\nabla}\cdot\boldsymbol{\nabla}$, with $\boldsymbol{\nabla}$ as the usual three-dimensional gradient operator, and $\eta$ is the shear viscosity of the bulk fluid. Here, $\boldsymbol{V}_\pm(\boldsymbol{r})$ and $p_\pm(\boldsymbol{r})$ are the fluid velocities and pressures at position $\boldsymbol{r}$, with the subscripts "−" and "+" indicating respectively whether the fluid is located inside and outside of the membrane compartment. The last relation in Eq. (S.125) is the incompressibility condition of the bulk fluid, whereas the former equation is the momentum conservation relationship, which follows from the stress balance condition:

$$\boldsymbol{\nabla}\cdot\boldsymbol{\Omega}_\pm = \boldsymbol{0}, \quad (S.126)$$

where the Stokesian stresses, which are denoted by $\boldsymbol{\Omega}_\pm$ for the respective inner and outer flows, are given by

$$\boxed{\boldsymbol{\Omega}_\pm = -p_\pm\boldsymbol{I} + \eta\left[(\boldsymbol{\nabla}\boldsymbol{V}_\pm) + (\boldsymbol{\nabla}\boldsymbol{V}_\pm)^{\mathsf{T}}\right].} \quad (S.127)$$

As discussed in the previous section (see § I.G.), the active forces in the semi-microscopic model would require the addition of an active source term in Eq. (S.126), namely we would write $\boldsymbol{\nabla}\cdot\boldsymbol{\Omega}_+ + \boldsymbol{F}_A = \boldsymbol{0}$. Nevertheless, this active force term can be projected onto the surface of the membrane. This leads to an effective two-dimensional formulation of the active forces, characterized by the membrane stresses $\sigma_A^{\alpha\beta}$ and $M_A^{\alpha\beta}$, which depend on the first and second moments of the transverse force distribution. In other words, in this description, the active forces $\boldsymbol{F}_A$ are absent from the momentum equation of the bulk fluid, as shown in Eq. (S.126), with the activity being included instead into the equations of motion of the membrane.

As a consequence, the stress vectors of the bulk fluid at the membrane surface must equal the membrane body forces $\boldsymbol{f}$, cf. Eq. (S.19). Hence, the force balance across the membrane is given by

$$\boxed{\boldsymbol{f} = (\boldsymbol{\Omega}_+ - \boldsymbol{\Omega}_-)\cdot\boldsymbol{n} \equiv \boldsymbol{\pi}_+ - \boldsymbol{\pi}_-, \text{ at } \boldsymbol{r} = \boldsymbol{R},} \tag{S.128}$$

where $\boldsymbol{R}$ is the position vector of the membrane surface, $\boldsymbol{n}$ is the normal unit vector to the membrane, and we define the stress vector acting across the surface as

$$\boldsymbol{\pi}_\pm = \boldsymbol{\Omega}_\pm \cdot \boldsymbol{n}. \tag{S.129}$$

In the local coordinate system of the membrane, we can decompose Eq. (S.128) into the normal component

$$f = \boldsymbol{n}\cdot\Big[\boldsymbol{\pi}_+(\boldsymbol{r}=\boldsymbol{R}) - \boldsymbol{\pi}_-(\boldsymbol{r}=\boldsymbol{R})\Big], \tag{S.130}$$

as well as along the in-plane membrane components

$$f^\alpha = \boldsymbol{e}^\alpha\cdot\Big[\boldsymbol{\pi}_+(\boldsymbol{r}=\boldsymbol{R}) - \boldsymbol{\pi}_-(\boldsymbol{r}=\boldsymbol{R})\Big]. \tag{S.131}$$

Lastly, we note that the latter procedure of projecting the active forces onto the membrane surface would also tangentially modify the boundary condition at the fluid–membrane interface [29]. This leads to a slip boundary condition, and we denote the additional contribution by the surface velocity $\boldsymbol{v}_S$. Hence, if the desired boundary condition in the semi-microscopic description is a non-slip condition at the membrane surface, then the slip velocity in the effective two-dimensional formulation (that respects the equivalence of the two descriptions) is [29]:

$$\boxed{\boldsymbol{v}_S = \frac{1}{2\eta}\boldsymbol{e}^\alpha\,\nabla_\alpha\mathbb{Q},} \tag{S.132}$$

where $\mathbb{Q}$ is the second moment of the active force distribution, as defined in Eq. (S.107). Thus, this leads to the following boundary condition:

$$\boxed{\boldsymbol{v} + \boldsymbol{v}_S - \boldsymbol{j}_V = \boldsymbol{V}_\pm(\boldsymbol{r} = \boldsymbol{R}),} \tag{S.133}$$

where $\boldsymbol{v}$ is the membrane velocity. Here, an additional term has been included on the left-hand-side, which is the local volume flux across the surface, denoted by $\boldsymbol{j}_V$. This accounts for the transport of bulk fluid across the surface of the membrane compartment through passive osmosis and active volume exchange due to the fusion and fission events (a detailed discussion of this volume flux will be deferred to the next section).

Additionally, we consider that the fluid flow is at rest at infinity and bounded at the origin of the compartment; namely, we have the boundary conditions:

$$\begin{cases} \boldsymbol{V}_-(|\boldsymbol{r}|\to 0) = \boldsymbol{0}, & p_-(|\boldsymbol{r}|\to 0) = P_-, \\ \boldsymbol{V}_+(|\boldsymbol{r}|\to \infty) = \boldsymbol{0}, & p_+(|\boldsymbol{r}|\to \infty) = P_+, \end{cases} \tag{S.134}$$

where $P_\pm$ are scalar constants to be determined via the osmolarity condition (see next section).

Therefore, these boundary conditions fully determine the fluid flow. Here, the flow is induced by the membrane movement, through Eq. (S.133), which is self-consistently determined via the stress balance condition at the membrane interface in Eq. (S.128). Hereinafter, the inertial terms in Eqs. (S.122) and (S.120) will be neglected, *i.e.*

$$\rho\dot{\boldsymbol{v}} = \boldsymbol{0}, \tag{S.135}$$

since the dynamics is overdamped by the ambient fluid, whose viscous forces, via $\boldsymbol{f}$, dominate over the inertial terms of the membrane. This assumption can be verified *a posteriori* by computing the corresponding Reynolds number, which must be much smaller than unity.

#### J. Boundary integral formulation

A more general approach to solve the Stokes equations in Eq. (S.125) in the presence of an active interface with a non-vanishing surface force, $\boldsymbol{f} \neq \boldsymbol{0}$, cf. Eq. (S.128), involves a boundary integral representation for the Stokes fluid flow [37]. A convenient starting point is the use of Lorentz reciprocal identity [38], which states that any two flows $\boldsymbol{V}$ and $\boldsymbol{V}'$ with their corresponding stress tensors $\boldsymbol{\Omega}$ and $\boldsymbol{\Omega}'$ satisfy the following identity:

$$\frac{\partial}{\partial x_k}\left[V_i'\,\Omega_{ik} - V_i\,\Omega_{ik}'\right] = 0, \tag{S.136}$$

which is written in Cartesian component form: $\boldsymbol{x} = x_k\,\hat{\boldsymbol{e}}_k$ with $\hat{\boldsymbol{e}}_k$ as basis vectors in a Cartesian coordinate system; similarly, we write the velocity $\boldsymbol{V} = V_i\,\hat{\boldsymbol{e}}_i$ and the stress $\boldsymbol{\Omega} = \Omega_{ij}\,\hat{\boldsymbol{e}}_i\otimes\hat{\boldsymbol{e}}_j$. If we identify $\boldsymbol{V}'$ with the fluid flow due to a point force of strength $\boldsymbol{g} = g_i\,\hat{\boldsymbol{e}}_i$ located at $\boldsymbol{x}_0$, we can write that

$$V_i'(\boldsymbol{x}) = \frac{1}{8\pi\eta}G_{ij}(\boldsymbol{x},\boldsymbol{x}_0)g_j, \tag{S.137}$$

$$\Omega_{ik}' = \frac{1}{8\pi}T_{ijk}(\boldsymbol{x},\boldsymbol{x}_0)g_j \tag{S.138}$$

where $G_{ij}$ and $T_{ijk}$ are the Green's functions for the flow and the stress at $\boldsymbol{x}$ due to a concentrated point force with the source at $\boldsymbol{x}_0$ [37]. Given that $\boldsymbol{g}$ is an arbitrary constant, we obtain that

$$\frac{\partial}{\partial x_k}\left[G_{ij}(\boldsymbol{x},\boldsymbol{x}_0)\Omega_{ik}(\boldsymbol{x}) - \eta V_i(\boldsymbol{x})T_{ijk}(\boldsymbol{x},\boldsymbol{x}_0)\right] = 0. \tag{S.139}$$

Integrating over a control volume $\mathcal{V}_C$ which is bounded by a closed surface $\mathcal{S}_C$ that can be either simply- or multiply-connected. If the point $\boldsymbol{x}_0$ lies outside the domain $\mathcal{V}_C$, the function within the square brackets in Eq. (S.139) is regular through $\mathcal{V}_C$ [37], and thus the volume integral can be turned into a surface integral over $\mathcal{S}_C$ by employing the divergence theorem. Hence, we have

$$\int_{\mathcal{S}_C}\mathcal{L}_{jk}(\boldsymbol{x},\boldsymbol{x}_0)\,n_k(\boldsymbol{x})\,\mathrm{d}S(\boldsymbol{x}) = 0, \tag{S.140}$$

where we define that

$$\mathcal{L}_{jk} = G_{ij}(\boldsymbol{x}, \boldsymbol{x}_0)\Omega_{ik}(\boldsymbol{x}) - \eta V_i(\boldsymbol{x}) T_{ijk}(\boldsymbol{x}, \boldsymbol{x}_0), \quad \text{(S.141)}$$

with the normal unit vector $\boldsymbol{n}$ being directed into the domain $\mathcal{V}_C$. Now, we consider that the point $\boldsymbol{x}_0$ is located within the interior of $\mathcal{V}_C$, and we construct a spherical volume $V_\varepsilon$ centered at $\boldsymbol{x}_0$ of small radius $\varepsilon$. Again, the function within the square brackets of Eq. (S.139) is regular throughout the domain of $\mathcal{V}_C$ that excludes $V_\varepsilon$. By divergence theorem, we obtain that

$$\int_{\mathcal{S}_C} \mathcal{L}_{jk}\, n_k \,\mathrm{d}S + \int_{\mathcal{S}_\varepsilon} \mathcal{L}_{jk}\, n_k \,\mathrm{d}S = 0, \quad \text{(S.142)}$$

where $\mathcal{S}_\varepsilon$ is the spherical surface that encloses the small spherical volume $\mathcal{V}_\varepsilon$. As the radius $\varepsilon$ tends to zero, we find that the tensors $G_{ij}$ and $T_{ijk}$ over $\mathcal{S}_\varepsilon$ reduce to the Stokeslet and stresslet tensors [38]; namely,

$$G_{ij} \simeq \frac{\delta_{ij}}{\varepsilon} + \frac{\hat{x}_i\hat{x}_j}{\varepsilon^3}, \quad \text{and} \quad T_{ijk} \simeq -6\frac{\hat{x}_i\hat{x}_j\hat{x}_k}{\varepsilon^5}, \quad \text{(S.143)}$$

where $\hat{\boldsymbol{x}} = \boldsymbol{x} - \boldsymbol{x}_0$ and $\delta_{ij}$ is the Kronecker delta. On the surface $\mathcal{S}_\varepsilon$, we have $\boldsymbol{n} = \hat{\boldsymbol{x}}/\varepsilon$ and $dS = \varepsilon^2 \sin\theta \,\mathrm{d}\theta \,\mathrm{d}\phi$. Therefore,

$$\int_{\mathcal{S}_C} \mathcal{L}_{jk}\, n_k \,\mathrm{d}S = -\int_{\mathcal{S}_\varepsilon} \left[\left(\delta_{ij} + \frac{\hat{x}_i\hat{x}_j}{\varepsilon^2}\right)\Omega_{ik}(\boldsymbol{x}) + 6\eta V_i(\boldsymbol{x})\frac{\hat{x}_i\hat{x}_j\hat{x}_k}{\varepsilon^4}\right]\hat{x}_k \sin\theta \,\mathrm{d}\theta \,\mathrm{d}\phi. \quad \text{(S.144)}$$

Note that the values of $\boldsymbol{V}$ and $\boldsymbol{\Omega}$ over $\mathcal{S}_\varepsilon$ tend to $\boldsymbol{V}(\boldsymbol{x}_0)$ and $\boldsymbol{\Omega}(\boldsymbol{x}_0)$ as $\varepsilon \to 0$. Also, $\hat{\boldsymbol{x}}$ decreases linearly with $\varepsilon$, as we approach $\hat{\boldsymbol{x}}_0$. This means that the stress term within the square brackets of Eq. (S.144) vanishes as $\varepsilon \to 0$, whilst the velocity term tends to a constant; namely,

$$\int_{\mathcal{S}_C} \mathcal{L}_{jk}\, n_k \,\mathrm{d}S = -6\eta V_i(\boldsymbol{x}_0)\int_{\mathcal{S}_\varepsilon} \frac{\hat{x}_i\hat{x}_j}{\varepsilon^4} \,\mathrm{d}S(\boldsymbol{x}). \quad \text{(S.145)}$$

To obtain that constant, we use the divergence theorem, $\int_{\mathcal{S}_\varepsilon} \hat{x}_i\hat{x}_j \,\mathrm{d}S = \varepsilon\int_{\mathcal{S}_\varepsilon} \hat{x}_i n_j \,\mathrm{d}S = \varepsilon\int_{\mathcal{V}_\varepsilon} \frac{\partial \hat{x}_i}{\partial \hat{x}_j} dV = \delta_{ij}\frac{4}{3}\pi\varepsilon^4$. Hence, we find that

$$\int_{\mathcal{S}_C} \mathcal{L}_{jk}(\boldsymbol{x}, \boldsymbol{x}_0)\, n_k(\boldsymbol{x}) \,\mathrm{d}S(\boldsymbol{x}) = -8\pi\eta V_j(\boldsymbol{x}_0), \quad \text{(S.146)}$$

which provides us with a representation of the flow in terms of two boundary integrals involving the Green's function $G_{ij}$ and the corresponding stress tensor $T_{ijk}$ [37].

As a result, Eq. (S.146) is valid when $\boldsymbol{x}_0$ lies inside the domain enclosed by the surface $\mathcal{S}_C$, whereas Eq. (S.140) is valid for any point $\boldsymbol{x}_0$ located outside that surface. As $\boldsymbol{x}_0$ approaches the bounding surface, it can be shown [37]:

$$\lim_{\boldsymbol{x}_0 \to \mathcal{S}_C} \int_{\mathcal{S}_C} V_i(\boldsymbol{x}) T_{ijk}(\boldsymbol{x}, \boldsymbol{x}_0) n_k(\boldsymbol{x}) \mathrm{d}S(\boldsymbol{x}) = \pm 4\pi V_j(\boldsymbol{x}_0) + \int_{\mathcal{S}_C}^{\mathscr{P}\mathscr{V}} V_i(\boldsymbol{x}) T_{ijk}(\boldsymbol{x}, \boldsymbol{x}_0) n_k(\boldsymbol{x}) \mathrm{d}S(\boldsymbol{x}), \quad \text{(S.147)}$$

where the plus and minus sign applies when the point $\boldsymbol{x}_0$ approaches the surface $\mathcal{S}_C$ from the interior and from the exterior side, respectively. Here, $\mathscr{P}\mathscr{V}$ denotes the principal value of the integral (defined as the improper integral when the point $\boldsymbol{x}_0$ is right on the surface $\mathcal{S}_C$). Therefore, substituting Eq. (S.147) with the minus sign into Eq. (S.140), and with the plus sign into Eq. (S.146), we obtain that for a point $\boldsymbol{x}_0$ that lies on the boundary,

$$V_j(\boldsymbol{x}_0) = \frac{1}{4\pi}\int_{\mathcal{S}_C}^{\mathscr{P}\mathscr{V}} V_i(\boldsymbol{x}) T_{ijk}(\boldsymbol{x}, \boldsymbol{x}_0) n_k(\boldsymbol{x}) \mathrm{d}S(\boldsymbol{x}) - \frac{1}{4\pi\eta}\int_{\mathcal{S}_C} G_{ij}(\boldsymbol{x}, \boldsymbol{x}_0)\Omega_{ik}(\boldsymbol{x}) n_k(\boldsymbol{x}) \mathrm{d}S(\boldsymbol{x}). \quad \text{(S.148)}$$

We consider an external and interior flow as discussed in section §I, which are denoted by $\boldsymbol{V}_\pm$, and delimited by an close membrane interface $\mathcal{M}$. As the flow in the exterior vanishes at infinity, from Eq. (S.146), we have

$$\boldsymbol{V}_+(\boldsymbol{r}) = \frac{1}{8\pi}\int_{\mathcal{M}} \boldsymbol{V}^\mathsf{T}(\boldsymbol{x})\, \boldsymbol{T}(\boldsymbol{x}, \boldsymbol{r})\, \boldsymbol{n}(\boldsymbol{x}) \,\mathrm{d}S(\boldsymbol{x}) - \frac{1}{8\pi\eta}\int_{\mathcal{M}} \boldsymbol{G}(\boldsymbol{r}, \boldsymbol{x})\, \boldsymbol{\Omega}_+(\boldsymbol{x}) \boldsymbol{n}(\boldsymbol{x}) \,\mathrm{d}S(\boldsymbol{x}), \quad \text{(S.149)}$$

where the symmetry property $G_{ij}(\boldsymbol{x}, \boldsymbol{x}_0) = G_{ji}(\boldsymbol{x}_0, \boldsymbol{x})$ is used to switch the order of indices [37]. By employing Eq. (S.140) for the internal flow $\boldsymbol{V}_-$ at a point $\boldsymbol{r}$ that is positioned in the exterior of the surface $\mathcal{M}$, we find

$$\int_{\mathcal{M}} \Big[\boldsymbol{G}(\boldsymbol{r}, \boldsymbol{x})\, \boldsymbol{\Omega}_-(\boldsymbol{x}) - \eta \boldsymbol{V}^\mathsf{T}(\boldsymbol{x})\, \boldsymbol{T}(\boldsymbol{x}, \boldsymbol{r})\Big] \boldsymbol{n}(\boldsymbol{x}) \mathrm{d}S(\boldsymbol{x}) = 0. \quad \text{(S.150)}$$

From the above equation and Eq. (S.149), we obtain

$$\boldsymbol{V}_+(\boldsymbol{r}) = -\frac{1}{8\pi\eta}\int_{\mathcal{M}} \boldsymbol{G}(\boldsymbol{r}, \boldsymbol{x})\, \boldsymbol{f}(\boldsymbol{x}) \,\mathrm{d}S(\boldsymbol{x}), \quad \text{(S.151)}$$

where $\boldsymbol{f} = (\boldsymbol{\Omega}_+ - \boldsymbol{\Omega}_-)\boldsymbol{n}$ is the interfacial surface force, as defined in Eq. (S.128). In an analogous way, we can derive the inner fluid flow to be

$$\boldsymbol{V}_-(\boldsymbol{r}) = -\frac{1}{8\pi\eta}\int_{\mathcal{M}} \boldsymbol{G}(\boldsymbol{r}, \boldsymbol{x})\, \boldsymbol{f}(\boldsymbol{x}) \,\mathrm{d}S(\boldsymbol{x}), \quad \text{(S.152)}$$

for any point $\boldsymbol{r}$ which is located inside the compartment. By approaching the interface, from either the interior or exterior side, and using the result of Eq. (S.147), we find

$$\boxed{\boldsymbol{V}_\pm(\boldsymbol{r}\!=\!\boldsymbol{R}) = -\frac{1}{8\pi\eta}\int_{\mathcal{M}} \boldsymbol{G}(\boldsymbol{r}, \boldsymbol{x})\, \boldsymbol{f}(\boldsymbol{x}) \,\mathrm{d}S(\boldsymbol{x}),} \quad \text{(S.153)}$$

which means that the fluid flow can be computed everywhere by prescribing the surface force $\boldsymbol{f}$. Moreover, the function $G_{ij}$ is the same throughout the whole domain, and thus is simply given by the free-space Green's function, so-called Stokeslet or Oseen tensor $\mathscr{S}_{ij}$, namely we write $G_{ij}(\boldsymbol{x},\boldsymbol{x}_0)=\mathscr{S}_{ij}(\boldsymbol{x}-\boldsymbol{x}_0)$, with the latter being

$$\boxed{\mathscr{S}_{ij}(\hat{\boldsymbol{x}})=\frac{\delta_{ij}}{|\hat{\boldsymbol{x}}|}+\frac{\hat{x}_i\hat{x}_j}{|\hat{\boldsymbol{x}}|^3}.} \tag{S.154}$$

A boundary integral representation can also be found for the inner and outer pressures as follows:

$$P_\pm(\boldsymbol{r})=-\frac{1}{8\pi}\int_{\mathcal{M}}\mathscr{P}(\boldsymbol{r}-\boldsymbol{x})\cdot\boldsymbol{f}(\boldsymbol{x})\,\mathrm{d}S(\boldsymbol{x}), \tag{S.155}$$

with $\mathscr{P}(\hat{\boldsymbol{x}})=2\hat{\boldsymbol{x}}\,/\,|\hat{\boldsymbol{x}}|^3$. For a more detailed discussion and a generalization to different viscosities inside and outside the compartment, the reader is referred to [37].

#### K. Osmosis and volume dynamics

The ambient fluid within cells is almost never a pure solvent, existing as a solution that contains various other molecules which are called generically as solutes. Many of these solutes are, in fact, involved in the biochemical reactions (discussed in § I.A.) that lead to the fussion and fission events. Membranes act as semi-permeable barriers that allow solvent molecules to pass through, but prohibit the solutes to permeate them [39]. If the osmolarity (*i.e.* the number of osmoles of solute per unit volume of solution) is different across the membrane, then the system tends to minimize the overall free-energy of the two solutions by either transporting the solvent molecules into the membrane compartment (hypotonicity), or out of the enclosed volume of the membrane (hypertonicity). This migration of solvent molecules is quantitatively described by an osmotic pressure, which is the force per unit area of membrane that needs to be applied to keep the solution at a constant volume [39]. In other words, an osmolarity jump across the closed membrane $\mathcal{M}$ results in a nonzero osmotic pressure at the interface, which in turn can lead to volume changes of the compartment.

Henceforth, we assume that the solute concentration is sufficiently low, so that the interaction among solutes can be neglected. The condition of equilibrium between the solutions is achieved when the chemical potentials associated with the solvents in them is the same. Note that the chemical potentials of the solute can be different across the semi-permeable membrane, as the solutes are prohibited to pass through [39]. By using a dilute limit approximation, the addition of solutes among the solvent molecules incurs only an entropic change that leads to the following chemical potential of solvent [40]:

$$\mu^{\mathfrak{B}}(T,P_\pm)=\mu_0^{\mathfrak{B}}(T,P_\pm)-k_BT\,\nu_0\,c_\pm, \tag{S.156}$$

where $c_\pm$ are the solute number densities, while $\mu_0^{\mathfrak{B}}$ is the bulk chemical potential associated with the pure solvent at a uniform pressure $P_\pm$ and temperature $T$, with the subscripts indicating the two regions, outside ("+") and inside ("−") of the membrane compartment, respectively. Note that the scalars $P_\pm$ are the homogeneous pressures defined in the boundary condition of Eq. (S.134). Also, $\nu_0$ is the molecular volume of the pure solvent, which can be computed at fixed $T$ in terms of the chemical potential of the pure solvent as follows: $\nu_0=\partial\mu_0^{\mathfrak{B}}/\partial P$.

At thermodynamic equilibrium, we must have that

$$\mu^{\mathfrak{B}}(T,P_+)=\mu^{\mathfrak{B}}(T,P_-), \tag{S.157}$$

with the difference in the hydrostatic pressure across the membrane being nonzero in general, $P_\vartriangle=P_+-P_-\neq 0$. However, this residual value is balanced at equilibrium by (and also defines) the osmotic pressure that we denote herein by $\Pi_\vartriangle$. For dilute solutions, the pressure drop $P_\vartriangle$ is relatively small; in the sense that we have $|P_\vartriangle|\ll P_\pm$. This allows us to expand Eq (S.157) to first order in $P_\vartriangle$, and to derive the so-called *van't Hoff's formula* [40]:

$$\Pi_\vartriangle=(c_+-c_-)\,k_BT,\quad\text{with}\quad P_\vartriangle=\Pi_\vartriangle. \tag{S.158}$$

In other words, this tells us that the Laplace pressure $P_\vartriangle$, the hydrostatic pressure drop, must equal at equilibrium the osmotic pressure $\Pi_\vartriangle$, which is linearly related to the difference in the solute concentrations across the membrane (if the solutions are in the dilute limit). Note that the corresponding osmolarities can be written as $c_\pm/\mathfrak{N}_A$, where $\mathfrak{N}_A$ is the Avogadro's constant ($\approx 6.022\times 10^{23}$).

Since the permeation dynamics through the membrane is typically much slower (on the order of minutes [41]) than the relaxation times of all other active and dissipative mechanisms, we assume that the solutions are sufficiently close to the equilibrium condition in Eq. (S.157), so that a nonzero excess of bulk chemical potential,

$$\delta\mu^{\mathfrak{B}}(T,P_\vartriangle)=\mu^{\mathfrak{B}}(T,P_+)-\mu^{\mathfrak{B}}(T,P_-), \tag{S.159}$$

can slowly drive changes to the enclosed volume. Thus, we expect to lowest order that the rate of change of volume to be linearly related to $\delta\mu^{\mathfrak{B}}=\nu_0\,(P_\vartriangle-\Pi_\vartriangle)$.

Moreover, the active processes of fusion and fission can actively add and remove bulk fluid, respectively, where the total rate of volume flux (in terms of the biochemical reaction rates of each $k$-th species) is given by

$$\mathbb{V}=\sum_{k,n,m}\mathbb{V}^k_{nm}r^k_{nm}\phi^k_m=\sum_k\mathbb{J}^k_{\circlearrowleft}\sum_{n=0}^{\aleph_k}\mathbb{V}^k_{n,n-1}, \tag{S.160}$$

where in the latter the adiabatic approximations of the cyclic transitions have been used (see § I.F.). By further using Eq. (S.65), we can write that

$$\boxed{\mathbb{V}=\sum_k\Phi^k\,\mathbb{V}^k_{\circlearrowleft}\,,} \tag{S.161}$$

where $\mathbb{V}^k_{\circlearrowleft}$ is the rate of the volume flux over each cycle,

$$\mathbb{V}^k_{\circlearrowleft}=\mathcal{R}^k_{\circlearrowleft}\sum_{n=0}^{\aleph_k}\mathbb{V}^k_{n,n-1}, \tag{S.162}$$

whose sign is negative for the fissogens species ($k = 1$) and positive for the fusogens ($k = 2$), that is,

$$\mathrm{sgn}\left(\mathbb{V}^k_{\circlearrowleft}\right) = (-1)^k. \tag{S.163}$$

As a result, by including the passive leakage of solvent as well as the active addition and removal of bulk fluid, the local volume flux can be written as follows:

$$\boxed{\boldsymbol{j}_V = \Big[\mathbb{V} - w_0\left(\Pi_\vartriangle - P_\vartriangle\right)\Big]\boldsymbol{n} = j_V\,\boldsymbol{n},} \tag{S.164}$$

where $w_0 > 0$ is a coefficient that characterizes the rate of solvent molecules transported per unit area of membrane in response to an excess of chemical potential $\delta\mu^{\mathfrak{B}}$. Note that the volume flux acts only along the membrane normal, so that the rate of change of the enclosed volume,

$$\frac{\mathrm{d}V}{\mathrm{d}t} = \int_{\mathcal{M}} j_V\,\mathrm{d}S, \tag{S.165}$$

equals the overall flux of volume carried across the membrane surface. Here, the osmotic pressure $\Pi_\vartriangle$ is given by the *van't Hoff's formula*, as in Eq. (S.158), but its value may be different to the Laplace pressure $P_\vartriangle$, with the exact equality holding only at thermodynamic equilibrium.

Herein we assume that the external osmolarity is kept at a fixed value, and thus $c_+$ is a constant (which is set by an external reservoir). On the other hand, the number density of solute molecules inside the compartment is given by a fixed number of solutes $\mathfrak{n}_{\mathcal{S}}$ and its volume $V$; namely, we have $c_- = \mathfrak{n}_{\mathcal{S}}/V$. Thus, the osmotic pressure is a function of volume, and it can be written as

$$\Pi_\vartriangle = k_B T\left[c_+ - \frac{\mathfrak{n}_{\mathcal{S}}}{V}\right]. \tag{S.166}$$

In general, the solute number $\mathfrak{n}_{\mathcal{S}}$ could also depend on the activity; nonetheless, we ignore this here, considering that the net volume transported by the fusion and fission events consists of mostly solvent molecules.

By employing the Reynolds transport theorem on the volume $V = \iiint \mathrm{d}V = \frac{1}{3}\int_{\mathcal{M}}(\boldsymbol{n}\cdot\boldsymbol{R})\,\mathrm{d}S$, and then by using integration by parts on the terms that contain gradients in the membrane velocity, the rate of change of volume can also be expressed as follows:

$$\frac{\mathrm{d}V}{\mathrm{d}t} = \int_{\mathcal{M}} v\,\mathrm{d}S, \tag{S.167}$$

where $v$ is the normal velocity (this identity is derived in Appendix C). Hence, the volume dynamics in Eq. (S.165) can be written in terms of the following integral equation:

$$\int_{\mathcal{M}} \mathrm{d}S\left[v + w_0\left(\Pi_\vartriangle - P_\vartriangle\right) - \sum_k \Phi^k\,\mathbb{V}^k_{\circlearrowleft}\right] = 0. \tag{S.168}$$

By solving this, the Laplace pressure can be determined as a function of the osmotic pressure, the normal membrane velocity, and the active sources of volume due to the biochemical reaction cycles. Note that the integrand in Eq. (S.168) may not vanish in general, and it holds only as an integral condition. In fact, this expression can be obtained by integrating over the normal component of the boundary condition in Eq. (S.133), and by noting that $\int_{\mathcal{M}}(\boldsymbol{n}\cdot\boldsymbol{V}_\pm)\,\mathrm{d}S = \iiint(\boldsymbol{\nabla}\cdot\boldsymbol{V}_\pm)\,\mathrm{d}V = 0$ due to the incompressibility condition, where the latter integration is over the entire volume that is enclosed by $\mathcal{M}$.

### II. QUASI-SPHERICAL PERTURBATION ANALYSIS

#### A. Surface parameterization

The results derived so far are coordinate-invariant and valid for any closed surface $\mathcal{M}$, that is given by an embedding function $\boldsymbol{R}$ (its position vector). We now assume that the surface of the membrane may be characterized by a slightly deformed sphere, which is described by an equation of the form:

$$\boxed{\boldsymbol{R} = R\left[1 + \varepsilon\, u(\theta, \varphi)\right] \hat{\boldsymbol{r}},} \tag{S.169}$$

in which we employ the spherical coordinates $(r, \theta, \varphi)$ having their origin at the center of the fixed, undeformed sphere of radius $r = R$. Here, $\hat{\boldsymbol{r}}$ is the unit vector along the radial direction, which is explicitly given by

$$\hat{\boldsymbol{r}} = (\sin\theta\cos\varphi,\, \sin\theta\sin\varphi,\, \cos\theta)^{\mathsf{T}}, \tag{S.170}$$

where the inclination angle $\theta \in [0, \pi)$, and the azimuth angle $\varphi \in [0, 2\pi)$. Furthermore, the unit tangent vectors along the spherical angular directions $\theta$ and $\varphi$ are denoted by $\hat{\boldsymbol{\theta}}$ and $\hat{\boldsymbol{\varphi}}$, respectively. By using Eq. (S.405), the explicit form of these two unit vectors are found to be

$$\hat{\boldsymbol{\varphi}} = (-\sin\varphi,\, \cos\varphi,\, 0)^{\mathsf{T}}, \qquad \text{and} \tag{S.171}$$

$$\hat{\boldsymbol{\theta}} = (\cos\theta\cos\varphi,\, \cos\theta\sin\varphi,\, -\sin\theta)^{\mathsf{T}}. \tag{S.172}$$

Herein, $\varepsilon \ll 1$ is a small dimensionless parameter which can ultimately be absorbed into the arbitrary angular function $u\,(\theta, \varphi)$. The nature of $\varepsilon$ is immaterial, and only included here in order to keep track of the perturbation order of the results that will be subsequently computed for the shape parameterization in Eq. (S.169).

We assume that $u(\theta, \varphi)$ is a well-behaved function that can be further expanded in a series of surface spherical harmonics, denoted by the subscript $\ell$; namely, we have

$$\boldsymbol{R} = R\left[1 + \varepsilon \sum_{\ell=0}^{\infty} u_\ell(\theta, \varphi)\right] \hat{\boldsymbol{r}}, \tag{S.173}$$

where the surface harmonics $u_\ell(\theta, \varphi)$ can be written as

$$u_\ell(\theta, \varphi) = \sum_{m=-\ell}^{\ell} u_{\ell,m}\, Y_{\ell,m}(\theta, \varphi), \tag{S.174}$$

with $u_{\ell,m}$ as some arbitrary amplitude, associated with each spherical harmonic function $Y_{\ell,m}$ [42]. Note that the zeroth harmonic $u_0$ results in a rescaling of the radius over which we perturb. The first harmonics correspond to a translation of the whole sphere, while the second harmonics describe an ellipsoidal elongation of the sphere.

#### B. Lamb's general solution of Stokes equations

In spherical coordinates, H. Lamb [43] has outlined a general solution for the Stokes equations in the absence of body forces within the bulk fluid. In order to derive this solution, we first employ the vector identity of the Laplacian $\boldsymbol{\nabla}^2 \boldsymbol{V} = \boldsymbol{\nabla}(\boldsymbol{\nabla} \cdot \boldsymbol{V}) - \boldsymbol{\nabla} \times (\boldsymbol{\nabla} \times \boldsymbol{V})$, and then by taking the divergence of the first vector equation in Eq. (S.125), we find that

$$\boldsymbol{\nabla} \cdot \left(\boldsymbol{\nabla}^2 \boldsymbol{V}\right) = \boldsymbol{\nabla}^2(\boldsymbol{\nabla} \cdot \boldsymbol{V}) = \frac{1}{\eta} \boldsymbol{\nabla}^2 p, \tag{S.175}$$

omitting for brevity the subscript "±". This implies that the pressure field $p$ must be a harmonic function, *i.e.*

$$\boldsymbol{\nabla}^2 p = 0, \tag{S.176}$$

due to incompressibility condition $\boldsymbol{\nabla} \cdot \boldsymbol{V} = 0$. We expand the pressure in a series of solid spherical harmonics:

$$p(r, \theta, \varphi) = \sum_{\ell=-\infty}^{\infty} p_\ell(r, \theta, \varphi), \tag{S.177}$$

where $p_\ell$ is a solid spherical harmonic of order $\ell$, which can be explicitly written as

$$p_\ell(r, \theta, \varphi) = \sum_{m=-\ell}^{\ell} p_{\ell,m}\, r^\ell\, Y_{\ell,m}(\theta, \varphi), \tag{S.178}$$

where $p_{\ell,m}$ is the spherical harmonic amplitude corresponding to the angular mode numbers $m$ and $\ell$.

This expansion forms the basis of Lamb's method [43], which allows us to find a general solution for the bulk fluid velocity $\boldsymbol{V}$ in terms of the pressure harmonics (as the inhomogeneous part), and two other solid spherical harmonics (representing the homogeneous solutions) which we denote by $\Upsilon_\ell$ and $\Xi_\ell$. Thus, this reads

$$\boxed{\begin{aligned} \boldsymbol{V}(r, \theta, \varphi) = &\sum_{\substack{\ell=-\infty \\ \ell \neq -1}}^{\infty} \frac{(\ell + 3)\, r^2\, \boldsymbol{\nabla} p_\ell - 2\ell\, r\, \hat{\boldsymbol{r}}\, p_\ell}{2\eta\, (\ell + 1)(2\ell + 3)} \\ &+ \sum_{\ell=-\infty}^{\infty} \left[\boldsymbol{\nabla} \Upsilon_\ell + \boldsymbol{\nabla} \times (r\, \hat{\boldsymbol{r}}\, \Xi_\ell)\right]. \end{aligned}} \tag{S.179}$$

These three harmonic functions can be easily computed if the velocity field is prescribed on a spherical surface, say, at a radius $r = R$. Nonetheless, in many studies of membrane vesicles, or droplets, one typically must satisfy the boundary conditions on a deformed (time-dependent) surface that deviates slightly from a fixed sphere. A common approximation, which has been carried out in such cases, is to assume that the prescribed velocity field on (or the stress vector across) the deformable surface acts on a sphere of an appropriately chosen mean radius. To motivate this approximation, we develop a perturbation scheme that allows one to compute the solid harmonics

$p_\ell$, $\Upsilon_\ell$ and $\Xi_\ell$ in Eq. (S.179) when the velocity and the stress across the membrane is prescribed on a deformed spherical surface, as one parameterized by Eq. (S.169).

Instead of directly working with the individual components of the velocity field $\boldsymbol{V}$, the boundary-value problem is more suitably addressed by the following quantities: the radial velocity, $\hat{\boldsymbol{r}}\cdot\boldsymbol{V}$, the radial component of the fluid vorticity, $(r\,\hat{\boldsymbol{r}})\cdot\boldsymbol{\nabla}\times\boldsymbol{V}$, and the Lamb term $\mathfrak{L}=(r\,\hat{\boldsymbol{r}})\cdot\boldsymbol{\nabla}(\hat{\boldsymbol{r}}\cdot\boldsymbol{V})-r\,\boldsymbol{\nabla}\cdot\boldsymbol{V}$. Note that the divergence of the velocity in the latter quantity is identically zero in the bulk fluid due to incompressibility; however, this will, in general, not vanish at the boundaries where the velocity field, or the stress vector, is prescribed.

The radial component of the velocity can be readily calculated from Eq. (S.179); namely,

$$\hat{\boldsymbol{r}}\cdot\boldsymbol{V}=\sum_{\ell=-\infty}^{\infty}\left[\frac{\ell\,r\,p_\ell}{2\eta\,(2\ell+3)}+\frac{\ell\,\Upsilon_\ell}{r}\right],\tag{S.180}$$

where the following identity has been used [43]:

$$r\frac{\partial h_\ell}{\partial r}=\ell\,h_\ell,\tag{S.181}$$

with $h_\ell$ being any solid spherical harmonic of order $\ell$. Similarly, the radial component of the curl of the velocity field $\boldsymbol{V}$ can be determined as follows:

$$\hat{\boldsymbol{r}}\cdot\boldsymbol{\nabla}\times\boldsymbol{V}=\sum_{\ell=-\infty}^{\infty}\frac{\ell\,(\ell+1)\,\Xi_\ell}{r}.\tag{S.182}$$

The usefulness of the third term, $\mathfrak{L}$, is more difficult to immediately assess, but its motivation will become clearer, shortly. To obtain its expression, we begin by differentiating Eq. (S.180) with respect to $r$, and then by utilizing again the identity in Eq. (S.181), we find that

$$\frac{\partial(\hat{\boldsymbol{r}}\cdot\boldsymbol{V})}{\partial r}=\sum_{\ell=-\infty}^{\infty}\left[\frac{\ell\,(\ell+1)\,p_\ell}{2\eta\,(2\ell+3)}+\frac{\ell\,(\ell-1)\Upsilon_\ell}{r^2}\right].\tag{S.183}$$

Due to the incompressibility condition, the above equation also allows us to compute the Lamb term $\mathfrak{L}$ in terms of the solid harmonic functions, that is,

$$\mathfrak{L}=\sum_{\ell=-\infty}^{\infty}\left[\frac{\ell\,(\ell+1)\,r\,p_\ell}{2\eta\,(2\ell+3)}+\frac{\ell\,(\ell-1)\Upsilon_\ell}{r}\right].\tag{S.184}$$

Hence, by evaluating the equations (S.180), (S.182), and (S.184) at the radius $r=R$, and by employing the identity $[h_\ell]_{r=R}=h_\ell\,(r/R)^{-\ell}$, for every solid spherical harmonics of order $\ell$, then we find that

$$[\hat{\boldsymbol{r}}\cdot\boldsymbol{V}]_{r=R}=\sum_{\ell=-\infty}^{\infty}\left[\frac{\ell\,p_\ell\,R^{\ell+1}}{2\eta(2\ell+3)\,r^\ell}+\frac{\ell\,\Upsilon_\ell}{R^{1-\ell}\,r^\ell}\right],\tag{S.185}$$

the radial part of the vorticity

$$[\hat{\boldsymbol{r}}\cdot\boldsymbol{\nabla}\times\boldsymbol{V}]_{r=R}=\sum_{\ell=-\infty}^{\infty}\frac{\ell\,(\ell+1)\,R^{\ell-1}\,\Xi_\ell}{r^\ell},\tag{S.186}$$

and, lastly, the Lamb divergence term

$$[\mathfrak{L}]_{r=R}=\sum_{\ell=-\infty}^{\infty}\left[\frac{\ell(\ell+1)p_\ell\,R^{\ell+1}}{2\eta\,(2\ell+3)\,r^\ell}+\frac{\ell\,(\ell-1)\,\Upsilon_\ell}{R^{1-\ell}\,r^\ell}\right].\tag{S.187}$$

We denote the prescribed velocity field at the radial position $r=R$ as $\boldsymbol{v}_{\rm m}(R,\theta,\varphi)$. By construction, we can trivially rewrite that $[\hat{\boldsymbol{r}}\cdot\boldsymbol{V}]_{r=R}=\hat{\boldsymbol{r}}\cdot\boldsymbol{v}_{\rm m}(R,\theta,\varphi)$ and $[r\,\hat{\boldsymbol{r}}\cdot\boldsymbol{\nabla}\times\boldsymbol{V}]_{r=R}=(r\,\hat{\boldsymbol{r}})\cdot\boldsymbol{\nabla}\times\boldsymbol{v}_{\rm m}(R,\theta,\varphi)$. Interestingly, the Lamb term at $r=R$ reduces to

$$[\mathfrak{L}]_{r=R}=-r\,\boldsymbol{\nabla}\cdot\boldsymbol{v}_{\rm m}(R,\theta,\varphi).\tag{S.188}$$

The velocity $\boldsymbol{v}_{\rm m}(R,\theta,\varphi)$ is a vector field that lives solely on a spherical surface of radius $R$. Hence, by following Lamb's approach, this vector field may be decomposed in a series of surface spherical harmonics; specifically,

$$\hat{\boldsymbol{r}}\cdot\boldsymbol{v}_{\rm m}(R,\theta,\varphi)=\sum_{\ell=1}^{\infty}\mathcal{X}_\ell\,(\theta,\varphi),\tag{S.189}$$

$$-r\,\boldsymbol{\nabla}\cdot\boldsymbol{v}_{\rm m}(R,\theta,\varphi)=\sum_{\ell=1}^{\infty}\mathcal{Y}_\ell\,(\theta,\varphi),\tag{S.190}$$

$$(r\,\hat{\boldsymbol{r}})\cdot\boldsymbol{\nabla}\times\boldsymbol{v}_{\rm m}(R,\theta,\varphi)=\sum_{\ell=1}^{\infty}\mathcal{Z}_\ell\,(\theta,\varphi),\tag{S.191}$$

where $\mathcal{X}_\ell$, $\mathcal{Y}_\ell$ and $\mathcal{Z}_\ell$ are surface harmonics, which are defined in an analogous way as shown in Eq. (S.174). Note that the summation runs only over the positive integers, as $\mathcal{X}_0=\mathcal{Y}_0=\mathcal{Z}_0=0$, identically. Without any loss of generality, we can rewrite the sums in Eqs. (S.185–S.187) from $\ell=1$ to infinity [43]. By means of the orthogonality condition of the solid spherical harmonics, we find

$$\begin{aligned}\mathcal{X}_\ell=\;&p_{-(\ell+1)}\frac{(\ell+1)\,r^{\ell+1}}{2\eta(2\ell-1)\,R^\ell}-\Upsilon_{-(\ell+1)}\frac{(\ell+1)\,r^{\ell+1}}{R^{\ell+2}}\\&+p_\ell\,\frac{\ell\,R^{\ell+1}}{2\eta(2\ell+3)\,r^\ell}+\Upsilon_\ell\,\frac{\ell\,R^{\ell-1}}{r^\ell},\end{aligned}\tag{S.192}$$

$$\begin{aligned}\mathcal{Y}_\ell=\;&p_{-(\ell+1)}\frac{\ell\,(\ell+1)\,r^{\ell+1}}{2\eta(2\ell-1)\,R^\ell}-\Upsilon_{-(\ell+1)}\frac{(\ell+1)(\ell+2)\,r^{\ell+1}}{R^{\ell+2}}\\&+p_\ell\,\frac{\ell\,(\ell+1)\,R^{\ell+1}}{2\eta(2\ell+3)\,r^\ell}+\Upsilon_\ell\,\frac{\ell\,(\ell-1)\,R^{\ell-1}}{r^\ell},\end{aligned}\tag{S.193}$$

$$\mathcal{Z}_\ell=\Xi_{-(\ell+1)}\frac{\ell\,(\ell+1)\,r^{\ell+1}}{R^{\ell+1}}+\Xi_\ell\,\frac{\ell\,(\ell+1)\,R^\ell}{r^\ell},\tag{S.194}$$

where the integer mode $\ell\geq 1$. It is noteworthy to mention that the negative harmonics (for which $\ell$ is negative) are formally determined by interchanging $\ell$ with $-\ell-1$.

Since the fluid is required to be at rest at infinity, in the case of the outer velocity flow (*i.e.* $\boldsymbol{V}=\boldsymbol{V}_+$), we must exclude the strictly positive harmonics, namely

$$p_\ell=\Upsilon_\ell=\Xi_\ell=0,\quad\text{for}\quad\ell\geq 1,\tag{S.195}$$

as the corresponding terms diverge as $r\to\infty$. Therefore, the resulting equations can be simultaneously solved for the three unknown harmonic functions: $p_{-(\ell+1)}$, $\Upsilon_{-(\ell+1)}$, and $\Xi_{-(\ell+1)}$. Consequently, this yields that

$$p_{-(\ell+1)} = \frac{\eta\,(2\ell-1)\,R^{\ell}}{(\ell+1)\,r^{\ell+1}}\,[\mathcal{Y}_\ell + (\ell+2)\,\mathcal{X}_\ell], \tag{S.196}$$

$$\Upsilon_{-(\ell+1)} = \frac{R^{\ell+2}}{2\,(\ell+1)\,r^{\ell+1}}\,[\mathcal{Y}_\ell + \ell\mathcal{X}_\ell], \tag{S.197}$$

$$\Xi_{-(\ell+1)} = \frac{R^{\ell+1}}{\ell\,(\ell+1)\,r^{\ell+1}}\,\mathcal{Z}_\ell, \tag{S.198}$$

for $\ell \geq 1$, and we also set that $p_{-1} = \Upsilon_{-1} = \Xi_{-1} = 0$, as there are no explicit sources or sinks for the ambient fluid. This completely determines both the velocity and pressure fields outside the closed membrane.

Similarly, for the inner flow (*i.e.* $\boldsymbol{V} = \boldsymbol{V}_-$), bounded by the membrane, the negative harmonics must be set to zero, as these terms diverge at the origin ($r \to 0$), *i.e.*

$$p_{-(\ell+1)} = \Upsilon_{-(\ell+1)} = \Xi_{-(\ell+1)} = 0, \quad \text{for} \quad \ell \geq 0. \tag{S.199}$$

Thus, we find that

$$p_\ell = \frac{\eta\,(2\ell+3)\,r^{\ell}}{\ell R^{\ell+1}}\,[\mathcal{Y}_\ell - (\ell-1)\,\mathcal{X}_\ell], \tag{S.200}$$

$$\Upsilon_\ell = \frac{r^\ell}{2\ell R^{\ell-1}}\,[(\ell+1)\,\mathcal{X}_\ell - \mathcal{Y}_\ell], \tag{S.201}$$

$$\Xi_\ell = \frac{r^\ell}{\ell\,(\ell+1)\,R^\ell}\,\mathcal{Z}_\ell, \tag{S.202}$$

for integer modes $\ell \geq 1$. This allows us to solve for both the velocity and pressure fields of an inner fluid flow.

#### C. Flows around a slightly deformed sphere

Herein, we assume that velocity and pressure fields of the ambient fluid can be written as a power series of the perturbation parameter $\varepsilon$, that is,

$$\boldsymbol{V}_\pm = \sum_{\mathtt{i}=0}^{\infty} \varepsilon^{\mathtt{i}}\,\boldsymbol{V}_\pm^{(\mathtt{i})}, \quad \text{and} \quad p_\pm = \sum_{\mathtt{i}=0}^{\infty} \varepsilon^{\mathtt{i}}\,p_\pm^{(\mathtt{i})}, \tag{S.203}$$

respectively, where the subscripts "+" and "−" denote accordingly the region inside and outside of the closed membrane. Due to the linearity of the Stokes equations, by substituting the above expansions into Eq. (S.125), we find that each of the individual perturbation fields must also satisfy the Stokes equations, *i.e.*

$$\eta\,\boldsymbol{\nabla}^2\boldsymbol{V}_\pm^{(\mathtt{i})} = \boldsymbol{\nabla}p_\pm^{(\mathtt{i})}, \quad \text{and} \quad \boldsymbol{\nabla}\cdot\boldsymbol{V}_\pm^{(\mathtt{i})} = 0. \tag{S.204}$$

Similarly, by equating the terms corresponding to each order in $\varepsilon$, the boundary conditions in Eq. (S.134) yield

$$\begin{cases} \boldsymbol{V}_-^{(\mathtt{i})}(|\boldsymbol{r}| \to 0) = \boldsymbol{0}, & p_-^{(\mathtt{i})}(|\boldsymbol{r}| \to 0) = P_-^{(\mathtt{i})}, \\ \boldsymbol{V}_+^{(\mathtt{i})}(|\boldsymbol{r}| \to \infty) = \boldsymbol{0}, & p_+^{(\mathtt{i})}(|\boldsymbol{r}| \to \infty) = P_+^{(\mathtt{i})}. \end{cases} \tag{S.205}$$

Likewise, the boundary condition in Eq. (S.133) at $\boldsymbol{r} = \boldsymbol{R}$ can be expressed as follows:

$$\boldsymbol{v}_{\mathrm{m}} = \boldsymbol{v} + \boldsymbol{v}_S - \boldsymbol{j}_V = \sum_{\mathtt{i}=0}^{\infty} \varepsilon^{\mathtt{i}}\,\boldsymbol{V}_\pm^{(\mathtt{i})}(\boldsymbol{r} = \boldsymbol{R}), \tag{S.206}$$

where we define the composite velocity $\boldsymbol{v}_{\mathrm{m}}$ as being the prescribed velocity field on the surface that matches the flow of the surrounding fluid. In addition, the membrane velocity $\boldsymbol{v} = \frac{\mathrm{d}}{\mathrm{d}t}\boldsymbol{R}$, the tangential slip velocity $\boldsymbol{v}_S$, and the volume flux $\boldsymbol{j}_V$ could all be also rewritten as a power series in the perturbation parameter; namely,

$$\boldsymbol{v} = \sum_{\mathtt{i}=0}^{\infty} \varepsilon^{\mathtt{i}}\boldsymbol{v}^{(\mathtt{i})}, \;\; \boldsymbol{v}_S = \sum_{\mathtt{i}=0}^{\infty} \varepsilon^{\mathtt{i}}\boldsymbol{v}_S^{(\mathtt{i})}, \text{ and } \boldsymbol{j}_V = \sum_{\mathtt{i}=0}^{\infty} \varepsilon^{\mathtt{i}}\,\boldsymbol{j}_V^{(\mathtt{i})}.$$

Hence, we have that

$$\boldsymbol{v}_{\mathrm{m}}^{(\mathtt{i})} = \boldsymbol{v}^{(\mathtt{i})} + \boldsymbol{v}_S^{(\mathtt{i})} - \boldsymbol{j}_V^{(\mathtt{i})} \tag{S.207}$$

at each order in the perturbation parameter. By Taylor expanding the boundary condition in Eq. (S.206) about the fixed radius $r = R$, we find that

$$\boldsymbol{V}_\pm^{(\mathtt{i})}(\boldsymbol{r} = \boldsymbol{R}) = \sum_{\mathtt{j}=0}^{\infty} \frac{(r-R)^{\mathtt{j}}}{\mathtt{j}!}\left[\frac{\partial^{(\mathtt{j})}}{\partial r^{(\mathtt{j})}}\,\boldsymbol{V}_\pm^{(\mathtt{i})}\right]_{r=R} \tag{S.208}$$

in which the term $r - R = \varepsilon R\,u(\theta,\varphi)$ at the membrane. By substituting (S.208) into Eq. (S.206), we have that

$$\boldsymbol{v}^{(\mathtt{i})} + \boldsymbol{v}_S^{(\mathtt{i})} - \boldsymbol{j}_V^{(\mathtt{i})} = \sum_{\mathtt{j}=0}^{\mathtt{i}} \frac{(R\,u)^{\mathtt{j}}}{\mathtt{j}!}\left[\frac{\partial^{(\mathtt{j})}}{\partial r^{(\mathtt{j})}}\,\boldsymbol{V}_\pm^{(\mathtt{i}-\mathtt{j})}\right]_{r=R} \tag{S.209}$$

which is derived by collecting and then equating all terms of the same order of $\varepsilon$. Satisfying the boundary condition of the velocities on the deformed surface of the membrane requires us to match an infinite number of perturbation fields in the velocity at a fixed undeformed sphere of radius $r = R$. Since each of the perturbation fields of the bulk fluid also satisfies the Stokes equations, Lamb's approach could be readily applied. The first few terms in Eq. (S.209) can be explicitly written as

$$\boldsymbol{v}^{(0)} + \boldsymbol{v}_S^{(0)} - \boldsymbol{j}_V^{(0)} = \left[\boldsymbol{V}_\pm^{(0)}\right]_{r=R} \quad \text{and} \tag{S.210}$$

$$\boldsymbol{v}^{(1)} + \boldsymbol{v}_S^{(1)} - \boldsymbol{j}_V^{(1)} = \left[\boldsymbol{V}_\pm^{(1)}\right]_{r=R} + (R\,u)\left[\frac{\partial\,\boldsymbol{V}_\pm^{(0)}}{\partial r}\right]_{r=R} \tag{S.211}$$

with the next term being quadratic in $u(\theta,\varphi)$, and so on. From Eq. (S.71), the surface divergence of the membrane velocity field is related to the overall mass flux $\mathbb{M}$, which may be expanded as $\mathbb{M} = \sum_{\mathtt{i}=0}^{\infty} \varepsilon^{\mathtt{i}}\mathbb{M}^{(\mathtt{i})}$, allowing us to rewrite the mass conservation law in Eq. (S.71) as

$$\mathbb{M}^{(\mathtt{i})} = (\operatorname{div}\boldsymbol{v})^{(\mathtt{i})}, \tag{S.212}$$

where $\operatorname{div}(\cdot)$ denotes the surface divergence, cf. (S.437).

Moreover, the perturbation expansion in Eq. (S.203) leads to the following stress tensors:

$$\boldsymbol{\Omega}_{\pm}^{(\mathtt{i})} = -p_{\pm}^{(\mathtt{i})}\boldsymbol{I} + \eta\left\{\left[\boldsymbol{\nabla}\boldsymbol{V}_{\pm}^{(\mathtt{i})}\right] + \left[\boldsymbol{\nabla}\boldsymbol{V}_{\pm}^{(\mathtt{i})}\right]^{\mathsf{T}}\right\}, \quad \text{(S.213)}$$

which also results in the perturbed stress vectors $\boldsymbol{\pi}_{\pm}^{(\mathtt{i})}$ acting across the surface of the membrane, namely

$$\boldsymbol{\pi}_{\pm}^{(\mathtt{i})} = \sum_{\mathtt{j}=0}^{\mathtt{i}} \boldsymbol{\Omega}_{\pm}^{(\mathtt{j})}\cdot \boldsymbol{n}^{(\mathtt{i}-\mathtt{j})}, \quad \text{(S.214)}$$

that follows from the series expansions in $\varepsilon$ of the stress vector $\boldsymbol{\pi}_{\pm} = \sum_{\mathtt{i}=0}^{\infty}\left[\varepsilon^{\mathtt{i}}\,\boldsymbol{\pi}_{\pm}^{(\mathtt{i})}\right]$, as well as the unit vector normal to the surface, $\boldsymbol{n} = \sum_{\mathtt{i}=0}^{\infty}\varepsilon^{\mathtt{i}}\,\boldsymbol{n}^{(\mathtt{i})}$. In particular, the zeroth and first order terms of Eq. (S.214) are

$$\boldsymbol{\pi}_{\pm}^{(0)} = \boldsymbol{\Omega}_{\pm}^{(0)}\cdot \boldsymbol{n}^{(0)}, \quad \text{and} \quad \text{(S.215)}$$

$$\boldsymbol{\pi}_{\pm}^{(1)} = \boldsymbol{\Omega}_{\pm}^{(0)}\cdot \boldsymbol{n}^{(1)} + \boldsymbol{\Omega}_{\pm}^{(1)}\cdot \boldsymbol{n}^{(0)}. \quad \text{(S.216)}$$

Similarly, the body forces $\boldsymbol{f}$ on the membrane can also be expanded in a power series of the parameter $\varepsilon$, that is, $\boldsymbol{f} = \sum_{\mathtt{i}=0}^{\infty}\varepsilon^{\mathtt{i}}\boldsymbol{f}^{(\mathtt{i})}$. Hence, this allows us to recast the stress boundary condition in Eq. (S.128), as follows:

$$\boldsymbol{f}^{(\mathtt{i})} = \sum_{\mathtt{j}=0}^{\mathtt{i}} \frac{(R\,u)^{\mathtt{j}}}{\mathtt{j}!}\left[\frac{\partial^{(\mathtt{j})}}{\partial r^{(\mathtt{j})}}\,\boldsymbol{\pi}_{\vartriangle}^{(\mathtt{i}-\mathtt{j})}\right]_{r=R}, \quad \text{(S.217)}$$

where we define that

$$\boldsymbol{\pi}_{\vartriangle}^{(\mathtt{i})} = \boldsymbol{\pi}_{+}^{(\mathtt{i})} - \boldsymbol{\pi}_{-}^{(\mathtt{i})}. \quad \text{(S.218)}$$

Eq. (S.217) is derived as analogously to Eq. (S.208), with the stress vectors $\boldsymbol{\pi}_{\pm}^{(\mathtt{i})}(\boldsymbol{r} = \boldsymbol{R})$ being expanded about a fixed sphere of radius $R$. This gives us the explicit forms of the zeroth and first order terms of Eq. (S.217):

$$\boldsymbol{f}^{(0)} = \left[\boldsymbol{\pi}_{\vartriangle}^{(0)}\right]_{r=R}, \quad \text{and} \quad \text{(S.219)}$$

$$\boldsymbol{f}^{(1)} = \left[\boldsymbol{\pi}_{\vartriangle}^{(1)}\right]_{r=R} + (R\,u)\left[\frac{\partial}{\partial r}\,\boldsymbol{\pi}_{\vartriangle}^{(0)}\right]_{r=R}. \quad \text{(S.220)}$$

Lastly, it is noteworthy to mention that $\boldsymbol{\nabla}\times\boldsymbol{f} = \boldsymbol{0}$; this follows from the fact that the membrane body force is given by $\boldsymbol{f} = -\boldsymbol{T}^{\alpha}_{;\alpha}$, with $\boldsymbol{T}$ as the traction, and by employing the identity that the curl of the gradient of any smooth tensor field is identically zero. As the traction is written as $\boldsymbol{T} = \sum_{\mathtt{i}=0}^{\infty}\varepsilon^{\mathtt{i}}\,\boldsymbol{T}^{(\mathtt{i})}$, we find that $\boldsymbol{\nabla}\times\boldsymbol{f}^{(\mathtt{i})} = \boldsymbol{0}$. Hence, by using Eq. (S.217), this leads to

$$\left[\boldsymbol{\nabla}\times\boldsymbol{\pi}_{\vartriangle}^{(0)}\right]_{r=R} = \boldsymbol{0}, \quad \text{and} \quad \text{(S.221)}$$

$$\left[\boldsymbol{\nabla}\times\boldsymbol{\pi}_{\vartriangle}^{(1)}\right]_{r=R} = -\,(R\,\boldsymbol{\nabla}u)\times\left[\frac{\partial}{\partial r}\,\boldsymbol{\pi}_{\vartriangle}^{(0)}\right]_{r=R}. \quad \text{(S.222)}$$

By starting with the leading-order perturbation field, each of the other fields could successively be solved by appropriately matching the boundary conditions.

#### 1. Zeroth order perturbation

At the leading-order in $\varepsilon$, the position vector $\boldsymbol{R}$ of the membrane is given by $\boldsymbol{R}^{(0)} = R\,\hat{\boldsymbol{r}}$. Thus, the membrane velocity $\boldsymbol{v}^{(0)} = \frac{\mathrm{d}}{\mathrm{d}t}\boldsymbol{R}^{(0)}$ may further be rewritten as

$$\boldsymbol{v}^{(0)} = \frac{\partial \boldsymbol{R}^{(0)}}{\partial t} + [v^{\alpha}]^{(0)}\boldsymbol{R}^{(0)}_{,\alpha} = v^{(0)}\boldsymbol{n}^{(0)} + [v^{\alpha}]^{(0)}\boldsymbol{e}^{(0)}_{\alpha}, \quad \text{(S.223)}$$

where the first equality follows from the definition of the material derivative, noting that $\boldsymbol{R}^{(0)}_{,\alpha} = \partial_{\alpha}\boldsymbol{R}^{(0)}$, and $[v^{\alpha}]^{(0)}$ is the membrane tangential velocity at zeroth-order, with the index $\alpha\in\{\theta,\varphi\}$. On the other hand, the latter equality is the decomposition of $\boldsymbol{v}^{(0)}$ into the basis $\{\boldsymbol{e}^{(0)}_{\alpha}, \boldsymbol{n}^{(0)}\}$, with $v^{(0)}$ being the normal velocity of the membrane.

Since $\boldsymbol{n}^{(0)} = \hat{\boldsymbol{r}}$, $\boldsymbol{e}^{(0)}_{\theta} = R\,\hat{\boldsymbol{\theta}}$, and $\boldsymbol{e}^{(0)}_{\varphi} = R\sin\theta\,\hat{\boldsymbol{\varphi}}$, this yields that $v^{(0)} = 0$, and consequently we find

$$\boldsymbol{v}^{(0)} = \hat{v}^{(0)}_{\theta}\,\hat{\boldsymbol{\theta}} + \hat{v}^{(0)}_{\varphi}\,\hat{\boldsymbol{\varphi}}, \quad \text{(S.224)}$$

where we additionally define that

$$\hat{v}^{(0)}_{\theta} = R\left[v^{\theta}\right]^{(0)}, \quad \text{and} \quad \hat{v}^{(0)}_{\varphi} = R\sin\theta\,[v^{\varphi}]^{(0)}. \quad \text{(S.225)}$$

As expected, the membrane flow at this order is solely within the tangential plane of a fixed sphere of radius $R$, and thus movements along the normal are restricted.

Using Lamb's method as described in previous section, we need to compute the radial velocity of the prescribed field on the membrane, which is given by

$$\hat{\boldsymbol{r}}\cdot\boldsymbol{v}^{(0)}_{\mathtt{m}} = \sum_{\ell=0}^{\infty}\mathcal{X}^{(0)}_{\ell} = 0, \quad \text{(S.226)}$$

where the surface harmonic functions $\mathcal{X}^{(0)}_{\ell}$ are identically zero for each mode, as the left-hand-side of the equation is a scalar constant, which is independent of the angular coordinates. Similarly, the slip velocity $\boldsymbol{v}^{(0)}_{S} = \boldsymbol{0}$, as it only depends on the surface gradients of $\mathbb{Q}^{(0)}$ that is a scalar at this order. Also, since $v^{(0)} = 0$ and $\mathcal{X}^{(0)}_{0} = 0$, we find that the volume flux vanishes identically; that is,

$$j^{(0)}_{V} = \mathbb{V}^{(0)} + w_0\left[P^{(0)}_{\vartriangle} - \Pi^{(0)}_{\vartriangle}\right] = 0, \quad \text{(S.227)}$$

where $\mathbb{V}^{(0)}$ is the net active volume flux at zeroth order in the perturbation, while $\Pi^{(0)}_{\vartriangle}$ and $P^{(0)}_{\vartriangle}$ are the corresponding osmotic and Laplace pressures at zeroth order. Note that the Lamb term, $\mathfrak{L}^{(0)} = -r\boldsymbol{\nabla}\cdot\boldsymbol{v}^{(0)}$, is given by

$$\mathfrak{L}^{(0)} = -R\,\mathrm{div}\,\boldsymbol{v}^{(0)}_{\mathtt{m}} = -R\,\mathrm{div}\,\boldsymbol{v}^{(0)}_{S} + 2j^{(0)}_{V} - R\mathbb{M}^{(0)}, \quad \text{(S.228)}$$

where the final equality follows directly from Eq. (S.212). Here, $\mathbb{M}^{(0)}$ is the homogeneous steady-state of Eq. (S.73), and its value is found to be zero, cf. Eq. (S.273). Thus, we have that $\mathfrak{L}^{(0)} = \sum_{\ell=0}^{\infty}\mathcal{Y}^{(0)}_{\ell} = 0$, which implies that each of the surface harmonics $\mathcal{Y}^{(0)}_{\ell} = 0$ for all $\ell \geq 0$.

Consequently, $p_\ell^{(0)}$ and $\Upsilon_\ell^{(0)}$ corresponding to inner flow $\boldsymbol{V}_-^{(0)}$ can be obtained from Eq. (S.200) and (S.201) to be

$$p_\ell^{(0)} = \Upsilon_\ell^{(0)} = 0, \quad \text{for all} \quad \ell \geq 1. \tag{S.229}$$

Additionally, the negative solid harmonics $p_{-(\ell+1)}^{(0)}$ and $\Upsilon_{-(\ell+1)}^{(0)}$ which are associated with the outer fluid $\boldsymbol{V}_+^{(0)}$ are found as well to be identically zero, that is,

$$p_{-(\ell+1)}^{(0)} = \Upsilon_{-(\ell+1)}^{(0)} = 0, \quad \text{for all} \quad \ell \geq 0. \tag{S.230}$$

Again, as the membrane normal $\boldsymbol{n}^{(0)} = \hat{\boldsymbol{r}}$, the stress vectors $\boldsymbol{\pi}_\pm^{(0)}$ acting across the membrane are given by

$$\boldsymbol{\pi}_\pm^{(0)} = -\hat{\boldsymbol{r}}\, p_\pm^{(0)} + \eta\, r \frac{\partial}{\partial r}\left[\frac{\boldsymbol{V}_\pm^{(0)}}{r}\right] + \frac{\eta}{r}\boldsymbol{\nabla}\Big[r\hat{\boldsymbol{r}}\cdot\boldsymbol{V}_\pm^{(0)}\Big], \tag{S.231}$$

where the inner pressure field is written as

$$p_-^{(0)} = P_-^{(0)} + \sum_{\ell=1} p_\ell^{(0)} = P_-^{(0)}, \tag{S.232}$$

while the outer pressure is given by

$$p_+^{(0)} = P_+^{(0)} + \sum_{\ell=1} p_{-(\ell+1)}^{(0)} = P_+^{(0)}, \tag{S.233}$$

with $P_\pm^{(0)}$ as the zeroth solid spherical harmonic associated with the fluid outside ("+") and inside ("−") of the closed membrane. Note that the scalars $P_\pm^{(0)}$ do not contribute to the velocity fields $\boldsymbol{V}_\pm^{(0)}$, and only the harmonics of a nonzero degree affects the fluid flow, cf. Eq. (S.179).

The other solid spherical harmonics, $\Xi_\ell^{(0)}$ and $\Xi_{-(\ell+1)}^{(0)}$, which accordingly correspond to the inner and outer fluid flows, can be found from the stress boundary condition. By taking the curl of the stress vectors in Eq. (S.231), and then by using the condition in Eq. (S.221), the radial component of the latter gives us that $\mathcal{Z}_\ell^{(0)} = 0$ for all $\ell$. Hence, from Eq. (S.198) and (S.202), we have that both harmonics $\Xi_\ell^{(0)}$ and $\Xi_{-(\ell+1)}^{(0)}$ vanish identically.

As a result, the velocity fields $\boldsymbol{V}_\pm^{(0)} = \boldsymbol{0}$ everywhere within the bulk fluid, including at $r = R$, which means that the membrane velocity $\boldsymbol{v}^{(0)} = \boldsymbol{0}$, and thus $\boldsymbol{v}_{\mathrm{m}}^{(0)} = \boldsymbol{0}$. By means of Eq. (S.231), we find that the stress vectors $\boldsymbol{\pi}_\pm^{(0)} = -\hat{\boldsymbol{r}}\, P_\pm^{(0)}$, whereas the stress tensors are given by

$$\boxed{\boldsymbol{\Omega}_\pm^{(0)} = -P_\pm^{(0)}\boldsymbol{I}.} \tag{S.234}$$

Hence, the stress boundary condition in Eq. (S.217) may be written as follows:

$$\boldsymbol{f}^{(0)} = -\hat{\boldsymbol{r}}\left[P_+^{(0)} - P_-^{(0)}\right] = -\hat{\boldsymbol{r}}\, P_\vartriangle^{(0)}, \tag{S.235}$$

where, in the latter equality, we define the pressure drop across the membrane to be $P_\vartriangle^{(0)} = P_+^{(0)} - P_-^{(0)}$. Thus, by decomposing the body force $\boldsymbol{f}^{(0)}$ onto the local membrane basis, we have that

$$\boxed{f^{(0)} = -P_\vartriangle^{(0)}, \quad \text{and} \quad \boldsymbol{f}_S^{(0)} = \boldsymbol{e}_\alpha^{(0)}\,[f^\alpha]^{(0)} = \boldsymbol{0},} \tag{S.236}$$

where $f^{(0)}$ and $[f^\alpha]^{(0)}$ are the normal and tangential components of the body force, respectively, and $\boldsymbol{f}_S^{(0)}$ representing the surface part of the body force at zeroth order.

### 2. *First order perturbation*

At first-order in the perturbation, the position vector of the membrane is given by $\boldsymbol{R}^{(1)} = R\, u(\theta,\varphi)\,\hat{\boldsymbol{r}}$, and the corresponding membrane velocity $\boldsymbol{v}^{(1)}$ is found to be

$$\boldsymbol{v}^{(1)} = \frac{\partial \boldsymbol{R}^{(1)}}{\partial t} + [v^\alpha]^{(1)}\boldsymbol{R}_{,\alpha}^{(0)} = v^{(1)}\boldsymbol{n}^{(0)} + [v^\alpha]^{(1)}\boldsymbol{e}_\alpha^{(0)}, \tag{S.237}$$

which is derived by using that $v^{(0)} = 0$ and $[v^\alpha]^{(0)} = 0$. As before, we can rewrite the above expression as follows:

$$\boldsymbol{v}^{(1)} = \hat{v}_r^{(1)}\,\hat{\boldsymbol{r}} + \hat{v}_\theta^{(1)}\,\hat{\boldsymbol{\theta}} + \hat{v}_\varphi^{(1)}\,\hat{\boldsymbol{\varphi}}, \tag{S.238}$$

in which we normalize the tangent vectors $\boldsymbol{e}_\theta^{(0)}$ and $\boldsymbol{e}_\varphi^{(0)}$, and then we define the tangential velocities

$$\hat{v}_\theta^{(1)} = R\,\left[v^\theta\right]^{(1)} \quad \text{and} \quad \hat{v}_\varphi^{(1)} = R\sin\theta\,[v^\varphi]^{(1)}, \tag{S.239}$$

while the radial velocity of $\boldsymbol{v}^{(1)}$ is found to be

$$\hat{v}_r^{(1)} = R\,\frac{\partial u}{\partial t} = R\sum_{\ell=0}^{\infty}\frac{\partial u_\ell}{\partial t}. \tag{S.240}$$

As the fluid velocities $\boldsymbol{V}_\pm^{(0)} = \boldsymbol{0}$, the boundary condition in Eq. (S.211) reduces to

$$\boldsymbol{v}^{(1)} + \boldsymbol{v}_S^{(1)} - \boldsymbol{j}_V^{(1)} = \left[\boldsymbol{V}_\pm^{(1)}\right]_{r=R}, \tag{S.241}$$

which allows us to readily apply Lamb's method in order to obtain both the inner and outer fluid flows. Thus, by employing Eq. (S.189), this gives us the surface spherical harmonics at the first-order in the expansion:

$$\boxed{\mathcal{X}_\ell^{(1)} = R\,\frac{\partial u_\ell}{\partial t} - \mathbb{V}_\ell^{(1)}, \quad \text{with} \quad \ell \geq 1,} \tag{S.242}$$

where $u_\ell$ and $\mathbb{V}_\ell^{(1)}$ are the surface harmonics of degree $\ell$ which are associated with $u(\theta,\varphi)$ and the active volume flux-rate $\mathbb{V}^{(1)}(\theta,\varphi)$ at first order in the perturbation.

From Eq. (S.212), we have $\operatorname{div}\boldsymbol{v}^{(1)} = \mathbb{M}^{(1)}$, and since $R\operatorname{div}\boldsymbol{v}_{\mathrm{m}}^{(1)} = r\boldsymbol{\nabla}\cdot\boldsymbol{v}_{\mathrm{m}}^{(1)}$, we find, via Eq. (S.190), that

$$\boxed{\mathcal{Y}_\ell^{(1)} = -R\,\mathbb{M}_\ell^{(1)} + 2\mathbb{V}_\ell^{(1)} + \frac{\ell(\ell+1)}{2\eta R}\,\mathbb{Q}_\ell^{(1)},} \tag{S.243}$$

where $\mathbb{M}_\ell^{(1)}$ are surface spherical harmonics of degree $\ell$ corresponding to the overall mass flux-rate $\mathbb{M}^{(1)}(\theta,\varphi)$ at first order. Last two terms in Eq. (S.243) are obtained by using that $\operatorname{div} \boldsymbol{v}_{\mathrm{m}}^{(1)} = \operatorname{div}\left[\boldsymbol{v}^{(1)} + \boldsymbol{v}_S^{(1)}\right] + 2H^{(0)} j_V^{(1)}$, where the mean curvature $H^{(0)} = -1/R$, and the slip velocity

$$\boxed{\boldsymbol{v}_S^{(1)} = \frac{1}{2\eta}[\boldsymbol{e}^\alpha]^{(0)}\,\partial_\alpha \mathbb{Q}^{(1)}.} \tag{S.244}$$

This gives that $\operatorname{div} \boldsymbol{v}_S^{(1)} = R^2\Delta^{(0)}\mathbb{Q}^{(1)}/(2\eta R)$, with $\Delta^{(0)}$ as the Laplacian at zeroth order. By employing

$$R^2\Delta^{(0)} Y_{\ell,m}(\theta,\varphi) = -\ell(\ell+1) Y_{\ell,m}(\theta,\varphi)\,, \tag{S.245}$$

we derive the form of the last term in Eq (S.243), with $\mathbb{Q}_\ell^{(1)}$ being the surface spherical harmonics of degree $\ell$ associated with the second moment $\mathbb{Q}^{(1)}$.

As both $\mathcal{X}_0^{(1)}$ and $\mathcal{Y}_0^{(1)}$ are zero, due to the incompressibility condition, their associated equations reduce to

$$\boxed{R\frac{\partial u_0}{\partial t} = \frac{1}{2}R\,\mathbb{M}_0^{(1)} = \mathbb{V}_0^{(1)} + w_0\left[P_\triangle^{(1)} - \Pi_\triangle^{(1)}\right],} \tag{S.246}$$

where $P_\triangle^{(1)}$ and $\Pi_\triangle^{(1)}$ are the first order perturbations to the Laplace pressure and osmotic pressure, respectively.

Therefore, the surface harmonics $\mathcal{X}_\ell^{(1)}$ and $\mathcal{Y}_\ell^{(1)}$ allows us to obtain both the functions $p_\ell^{(1)}$ and $\Upsilon_\ell^{(1)}$ which are associated with the interior fluid flow $\boldsymbol{V}_-^{(1)}$, that is,

$$p_\ell^{(1)} = \frac{\eta\,(2\ell+3)\,r^\ell}{\ell R^{\ell+1}}\left[\mathcal{Y}_\ell^{(1)} - (\ell-1)\,\mathcal{X}_\ell^{(1)}\right], \tag{S.247}$$

$$\Upsilon_\ell^{(1)} = \frac{r^\ell}{2\ell R^{\ell-1}}\left[(\ell+1)\,\mathcal{X}_\ell^{(1)} - \mathcal{Y}_\ell^{(1)}\right], \tag{S.248}$$

with $\ell \geq 1$, and the functions $p_{-(\ell+1)}^{(1)}$ and $\Upsilon_{-(\ell+1)}^{(1)}$ that correspond to the outer fluid flow $\boldsymbol{V}_+^{(1)}$, namely

$$p_{-(\ell+1)}^{(1)} = \frac{\eta\,(2\ell-1)\,R^\ell}{(\ell+1)\,r^{\ell+1}}\left[\mathcal{Y}_\ell^{(1)} + (\ell+2)\,\mathcal{X}_\ell^{(1)}\right], \tag{S.249}$$

$$\Upsilon_{-(\ell+1)}^{(1)} = \frac{R^{\ell+2}}{2(\ell+1)\,r^{\ell+1}}\left[\mathcal{Y}_\ell^{(1)} + \ell\mathcal{X}_\ell^{(1)}\right], \tag{S.250}$$

where $\ell \geq 1$. On the other hand, $\Xi_{-(\ell+1)}^{(1)}$ and $\Xi_\ell^{(1)}$ can be found as before from the stress boundary condition.

The stress vectors $\boldsymbol{\pi}_\pm^{(1)}$ at first order in the perturbation expansion may be expressed in the form

$$\boldsymbol{\pi}_\pm^{(1)} = \eta r\frac{\partial}{\partial r}\left[\frac{\boldsymbol{V}_\pm^{(1)}}{r}\right] + \frac{\eta}{r}\,\boldsymbol{\nabla}\left[r\hat{\boldsymbol{r}}\cdot\boldsymbol{V}_\pm^{(1)}\right] - \boldsymbol{n}^{(1)}P_\pm^{(0)} - \hat{\boldsymbol{r}}\,p_\pm^{(1)}, \tag{S.251}$$

where the normal vector $\boldsymbol{n}^{(1)}$ is found to be

$$\boldsymbol{n}^{(1)} = -\frac{\partial u}{\partial\theta}\,\hat{\boldsymbol{\theta}} - \frac{1}{\sin\theta}\frac{\partial u}{\partial\varphi}\,\hat{\boldsymbol{\varphi}} = -r\boldsymbol{\nabla}u, \tag{S.252}$$

where the last equality follows from the definition of the three-dimensional gradient operator $\boldsymbol{\nabla}$ written in spherical coordinates. As before, the pressure inside the closed membrane is given by

$$p_-^{(1)} = P_-^{(1)} + \sum_{\ell=1} p_\ell^{(1)}, \tag{S.253}$$

whilst the outside pressure is

$$p_+^{(1)} = P_+^{(1)} + \sum_{\ell=1} p_{-(\ell+1)}^{(1)}\,, \tag{S.254}$$

where the scalars $P_\pm^{(1)}$ are the associated solid spherical harmonics of degree zero of the pressure fields.

Since $\boldsymbol{\nabla}\times\hat{\boldsymbol{r}} = \boldsymbol{0}$ and $\boldsymbol{\nabla}\times\boldsymbol{n}^{(1)} = -\hat{\boldsymbol{r}}\times\boldsymbol{\nabla}u$, the radial component of the curl of the stress vectors $\boldsymbol{\pi}_\pm^{(1)}$ corresponding to the inner and outer flows are given by

$$\hat{\boldsymbol{r}}\cdot\left[\boldsymbol{\nabla}\times\boldsymbol{\pi}_-^{(1)}\right]_{r=R} = \sum_{\ell=1}^{\infty}\frac{\eta\ell(\ell-1)(\ell+1)}{r^\ell R^{-\ell+1}}\,\Xi_\ell, \tag{S.255}$$

and

$$\hat{\boldsymbol{r}}\cdot\left[\boldsymbol{\nabla}\times\boldsymbol{\pi}_+^{(1)}\right]_{r=R} = \sum_{\ell=1}^{\infty}\frac{-\eta\ell(\ell+1)(\ell+2)}{r^{-(\ell+1)}R^{\ell+2}}\,\Xi_{-(\ell+1)}, \tag{S.256}$$

respectively. By using the form of $\Xi_\ell^{(1)}$ and $\Xi_{-(\ell+1)}^{(1)}$ from Eq. (S.202) and Eq. (S.198), together with Eq. (S.222), we find that

$$\sum_{\ell=1}^{\infty}\frac{\eta\,(2\ell+1)}{R}\mathcal{Z}_\ell^{(1)} = 0. \tag{S.257}$$

This yields that the surface harmonics $\mathcal{Z}_\ell^{(1)} = 0$, which implies that both $\Xi_\ell^{(1)}$ and $\Xi_{-(\ell+1)}^{(1)}$ are identically zero.

By employing Eqs. (S.235) and (S.251), the normal component of the body force $\boldsymbol{f}^{(1)}$ is given by

$$f^{(1)} = \boldsymbol{n}^{(0)}\cdot\boldsymbol{f}^{(1)} + \boldsymbol{n}^{(1)}\cdot\boldsymbol{f}^{(0)} = \hat{\boldsymbol{r}}\cdot\left[\boldsymbol{\pi}_\triangle^{(1)}\right]_{r=R}, \tag{S.258}$$

where the identity $\hat{\boldsymbol{r}}\cdot\boldsymbol{n}^{(1)} = 0$ and Eq. (S.220) are used to derive the last equation. To find $f^{(1)}$, we need to compute the radial part of the stress vectors. On the inner side of the closed membrane, we have

$$\hat{\boldsymbol{r}}\cdot\left[\boldsymbol{\pi}_-^{(1)}\right]_{r=R} = -P_-^{(1)} + \sum_{\ell=1}^{\infty}\left[\frac{\left(\ell^2-\ell-3\right)R^\ell}{(2\ell+3)\,r^\ell}\,p_\ell^{(1)} + \frac{2\eta\ell\,(\ell-1)\,R^{\ell-2}}{r^\ell}\,\Upsilon_\ell^{(1)}\right], \tag{S.259}$$

whereas, on the outer side of the membrane, we get

$$\hat{\boldsymbol{r}}\cdot\left[\boldsymbol{\pi}_+^{(1)}\right]_{r=R} = -P_+^{(1)} - \sum_{\ell=1}^{\infty}\left[\frac{\left(\ell^2+3\ell-1\right)r^{\ell+1}}{(2\ell-1)\,R^{\ell+1}}\,p_{-(\ell+1)}^{(1)} - \frac{2\eta(\ell+2)(\ell+1)\,r^{\ell+1}}{R^{\ell+2}}\,\Upsilon_{-(\ell+1)}^{(1)}\right]. \tag{S.260}$$

Hence, the normal body force $f^{(1)}$ in Eq. (S.258) can be written in terms of the surface harmonics $\mathcal{X}_\ell^{(1)}$ and $\mathcal{Y}_\ell^{(1)}$, by substituting Eq. (S.247) and (S.248) into Eq. (S.259), and Eq. (S.249) and (S.250) into Eq. (S.260), which yields

$$\boxed{f^{(1)} = -P_\Delta^{(1)} + \sum_{\ell=1}^{\infty} f_\ell^{(1)},} \tag{S.261}$$

where the (Laplace) pressure drop across the membrane at first order in the perturbation reads $P_\Delta^{(1)} = P_+^{(1)} - P_-^{(1)}$, and the surface harmonics $f_\ell^{(1)}$ are found to be

$$\boxed{f_\ell^{(1)} = \frac{\eta(2\ell+1)\left\{3\mathcal{Y}_\ell^{(1)} - [2\ell(\ell+1)-3]\mathcal{X}_\ell^{(1)}\right\}}{\ell(\ell+1)R}.} \tag{S.262}$$

The tangential contribution of the body force $\boldsymbol{f}^{(1)}$, at first order in $\varepsilon$, has the components

$$[f^\alpha]^{(1)} = \boldsymbol{f}^{(1)} \cdot [\boldsymbol{e}^\alpha]^{(0)} + \boldsymbol{f}^{(0)} \cdot [\boldsymbol{e}^\alpha]^{(1)}, \tag{S.263}$$

where $\boldsymbol{f}^{(0)} = -\hat{\boldsymbol{r}}\, P_\Delta^{(0)}$ from Eq. (S.235), while the explicit form of the body force $\boldsymbol{f}^{(1)}$ is found to be

$$\boldsymbol{f}^{(1)} = -\boldsymbol{n}^{(1)} P_\Delta^{(0)} + \left[\boldsymbol{\pi}_\Delta^{(1)}\right]_{r=R} \tag{S.264}$$

which readily follows by projecting stress tensor $\boldsymbol{\Omega}_\pm^{(1)}$ onto the surface of the membrane. This leads to

$$[f^\alpha]^{(1)} = [\boldsymbol{e}^\alpha]^{(0)} \cdot \left[\boldsymbol{\pi}_\Delta^{(1)}\right]_{r=R}. \tag{S.265}$$

Thus, the surface component of the body force at first order in $\varepsilon$ can be calculated through

$$\boldsymbol{f}_S^{(1)} = -\boldsymbol{n}^{(1)} P_\Delta^{(0)} + \left[\boldsymbol{\pi}_\Delta^{(1)} - \hat{\boldsymbol{r}}\left(\hat{\boldsymbol{r}} \cdot \boldsymbol{\pi}_\Delta^{(1)}\right)\right]_{r=R}, \tag{S.266}$$

where the term within the square brackets represents the surface part of the stress vectors acting across the membrane. The surface vectors associated with the outer and inner flows can be expressed as follows:

$$\left[\boldsymbol{\pi}_-^{(1)} - \hat{\boldsymbol{r}}\left(\hat{\boldsymbol{r}} \cdot \boldsymbol{\pi}_-^{(1)}\right)\right]_{r=R} = \sum_{\ell=1}^{\infty}\left[\frac{\ell(\ell+2)R^\ell}{(\ell+1)(2\ell+3)\,r^\ell}\boldsymbol{\nabla}_S\, p_\ell^{(1)} + \frac{2\eta(\ell-1)R^{\ell-2}}{r^\ell}\boldsymbol{\nabla}_S \Upsilon_\ell^{(1)}\right] + P_-^{(0)}\boldsymbol{\nabla}_S u, \tag{S.267}$$

and

$$\left[\boldsymbol{\pi}_+^{(1)} - \hat{\boldsymbol{r}}\left(\hat{\boldsymbol{r}} \cdot \boldsymbol{\pi}_+^{(1)}\right)\right]_{r=R} = \sum_{\ell=1}^{\infty}\left[\frac{(\ell^2-1)r^{\ell+1}}{\ell(2\ell-1)R^{\ell+1}}\boldsymbol{\nabla}_S\, p_{-(\ell+1)}^{(1)} - \frac{2\eta(\ell+2)\,r^{\ell+1}}{R^{\ell+2}}\boldsymbol{\nabla}_S \Upsilon_{-(\ell+1)}^{(1)}\right] + P_+^{(0)}\boldsymbol{\nabla}_S u, \tag{S.268}$$

respectively, where $\boldsymbol{\nabla}_S$ is the surface gradient on a unit sphere, which is defined as follows:

$$\boldsymbol{\nabla}_S(\cdot) = \frac{\partial(\cdot)}{\partial\theta}\hat{\boldsymbol{\theta}} + \frac{1}{\sin\theta}\frac{\partial(\cdot)}{\partial\varphi}\hat{\boldsymbol{\varphi}}. \tag{S.269}$$

The above expressions in Eq. (S.267) and (S.268) are derived by further employing the identities:

$$\boldsymbol{\nabla} h_\ell = \frac{\ell\, h_\ell\, \hat{\boldsymbol{r}}}{r} + \frac{\boldsymbol{\nabla}_S h_\ell}{r}, \quad \text{and} \tag{S.270}$$

$$[\boldsymbol{\nabla}_S h_\ell]_{r=R} = \left[\frac{r}{R}\right]^{-\ell} \boldsymbol{\nabla}_S h_\ell, \tag{S.271}$$

where $h_\ell$ is a solid spherical harmonic of degree $\ell$. Hence, by substituting Eq. (S.247) and (S.248) into Eq. (S.267), and Eq. (S.249) and (S.250) into Eq. (S.268), we find

$$\boxed{\boldsymbol{f}_S^{(1)} = -\sum_{\ell=1}^{\infty}\boldsymbol{\nabla}_S\left[\frac{\eta(2\ell+1)}{\ell(\ell+1)R}\left(\mathcal{X}_\ell^{(1)} + 2\mathcal{Y}_\ell^{(1)}\right)\right],} \tag{S.272}$$

which shows that the tangential part of $\boldsymbol{f}^{(1)}$ can be purely written as the surface gradient of a function, whose spherical harmonics are the terms in the square brackets.

#### D. Dynamics of the excess areal fractions

Herein, the homogeneous steady-states corresponding to Eq. (S.73) are denoted by $\bar{\Phi}_k$, and they must satisfy

$$\boxed{\mathbb{M}^{(0)} = \sum_k \bar{\Phi}_k\, \mathbb{M}_{\circlearrowleft}^k = 0.} \tag{S.273}$$

On the other hand, the steady-state mass fraction $\bar{\Phi}_0$ that is associated with Eq. (S.72) is obtained via the mass conservation condition, *i.e.* $1 - \bar{\Phi}_0 = \bar{\Phi}_1 + \bar{\Phi}_2$. By using this constraint, $\bar{\Phi}_k$ can be expressed as follows:

$$\boxed{\bar{\Phi}_k = \frac{\mathbb{M}_{\circlearrowleft}^{k'}}{\mathbb{M}_{\circlearrowleft}^{k'} - \mathbb{M}_{\circlearrowleft}^{k}}\left(1 - \bar{\Phi}_0\right), \quad \text{with } k \neq k',} \tag{S.274}$$

where both the indices $k,\, k' \in \{1, 2\}$. In deriving this homogeneous steady-state, we assume that the shape is also fixed, corresponding to the undeformed sphere of radius $R$ at zeroth order in the perturbation expansion.

The system is subsequently perturbed about this non-equilibrium steady-state by a small excess of membrane areal density; namely, we consider that

$$\Phi_0(\xi^\alpha, t) = \bar{\Phi}_0 - \varepsilon\, \Psi_0(\xi^\alpha, t) + \mathcal{O}[\varepsilon^2], \tag{S.275}$$

where $\varepsilon$ is the perturbation parameter and $\Psi_0$ is an arbitrary function which is of order $\mathcal{O}[\varepsilon^0]$ with respect to the uniform state $\bar{\Phi}_0$. This linear perturbation also leads to a linear response in the other slow variables,

$$\Phi_k(\xi^\alpha, t) = \bar{\Phi}_k + \varepsilon\, \Psi_k(\xi^\alpha, t) + \mathcal{O}[\varepsilon^2], \tag{S.276}$$

where $\Psi_k$ is the excess in the net mass fraction of the $k$-th membrane species. Due to the conservation requirement in Eq. (S.15), the functions $\Psi_k$ must satisfy that

$$\Psi_1 + \Psi_2 = \Psi_0. \tag{S.277}$$

Moreover, Eq. (S.276) implies the following perturbation expansion of the mass flux-rate $\mathbb{M}$ from Eq. (S.68):

$$\mathbb{M} = \mathbb{M}^{(0)} + \varepsilon\, \mathbb{M}^{(1)} + \mathcal{O}[\varepsilon^2], \tag{S.278}$$

which is truncated to linear order in $\varepsilon$. Hence, the first order contribution is found to be

$$\mathbb{M}^{(1)} = \sum_k \Psi_k\, \mathbb{M}^k_{\circlearrowleft}. \tag{S.279}$$

Thus, the dynamical equation of $\Psi_k$ associated with the first order perturbation of Eq. (S.78) is given by

$$\dot{\Psi}_k = 2\Omega_k\, [\Delta H]^{(1)} - \bar{\Phi}_k \mathbb{M}^{(1)} + \gamma_k\, \Delta^{(0)}\left[\frac{\Psi_0}{\bar{\Phi}_0} + \frac{\Psi_k}{\bar{\Phi}_k}\right], \tag{S.280}$$

where $[\Delta H]^{(1)} = \Delta^{(0)} H^{(1)} + \Delta^{(1)} H^{(0)}$ is the first order contribution to the Laplacian of the mean curvature $H$. From the shape parameterization in Eq. (S.169), we have that $H^{(0)} = -1/R$, and thus $\Delta^{(1)} H^{(0)} = 0$. Also, we find

$$H^{(1)} = \frac{u(\theta,\varphi)}{R} + \frac{\partial_\theta[\sin\theta\, \partial_\theta\, u(\theta,\varphi)]}{2R\sin\theta} + \frac{\partial^2_\varphi\, u(\theta,\varphi)}{2R\sin^2\theta}, \tag{S.281}$$

where $\partial^2_\alpha(\cdot) = \partial_\alpha[\partial_\alpha(\cdot)]$ is the second derivative with respect to the local surface coordinates. As the Laplacian operator $\Delta^{(0)}$ in spherical coordinates is given by

$$\Delta^{(0)}(\cdot) = \frac{\partial_\theta[\sin\theta\, \partial_\theta(\cdot)]}{R^2\sin\theta} + \frac{\partial^2_\varphi(\cdot)}{R^2\sin^2\theta}, \tag{S.282}$$

we can express the term $[\Delta H]^{(1)} = \Delta^{(0)} H^{(1)}$ as follows:

$$[\Delta H]^{(1)} = \frac{\Delta^{(0)} u}{R} + \frac{R\, \Delta^{(0)}\Delta^{(0)} u}{2}. \tag{S.283}$$

Since $\sum_k \Psi_k = \Psi_0$, the last term in Eq. (S.280) can be written in terms of the variables $\Psi_1$ and $\Psi_2$, *i.e.*

$$\frac{\Delta^{(0)}\Psi_k}{\bar{\Phi}_k} + \frac{\Delta^{(0)}\Psi_0}{\bar{\Phi}_0} = \Delta^{(0)}\Psi_k\left[\frac{1}{\bar{\Phi}_0} + \frac{1}{\bar{\Phi}_k}\right] + \frac{\Delta^{(0)}\Psi_{k'}}{\bar{\Phi}_0},$$

with $k' \neq k$. Also, the total derivative of $\Psi_k$ reduces to

$$\dot{\Psi}_k = \frac{\partial \Psi_k}{\partial t} + [v^\alpha]^{(0)}\Psi_{k,\alpha} + [v^\alpha]^{(1)}\bar{\Phi}_{k,\alpha} = \frac{\partial \Psi_k}{\partial t}, \tag{S.284}$$

as the membrane velocity at zeroth order $\boldsymbol{v}^{(0)} = \mathbf{0}$. Thus, Eq. (S.280) can be expressed in a matrix form as follows:

$$\boxed{\frac{\partial \boldsymbol{\Psi}}{\partial t} = -\mathsf{M}\,\boldsymbol{\Psi} + \mathsf{D}\,\Delta^{(0)}\boldsymbol{\Psi} + 2\boldsymbol{\Omega}\,[\Delta H]^{(1)},} \tag{S.285}$$

by defining the vectors $\boldsymbol{\Psi} = (\Psi_1, \Psi_2)^{\mathsf{T}}$, $\boldsymbol{\Omega} = (\Omega_1, \Omega_2)^{\mathsf{T}}$, where the superscript $\mathsf{T}$ indicates a vector transpose. On the other hand, $\mathsf{M}$ is a matrix which is defined by

$$\mathsf{M} = \begin{bmatrix} \bar{\Phi}_1 \mathbb{M}^1_{\circlearrowleft} & \bar{\Phi}_1 \mathbb{M}^2_{\circlearrowleft} \\ \bar{\Phi}_2 \mathbb{M}^1_{\circlearrowleft} & \bar{\Phi}_2 \mathbb{M}^2_{\circlearrowleft} \end{bmatrix}, \tag{S.286}$$

whilst the diffusion matrix $\mathsf{D}$ is given by

$$\mathsf{D} = \begin{bmatrix} \gamma_1\big(1/\bar{\Phi}_0 + 1/\bar{\Phi}_1\big) & \gamma_1/\bar{\Phi}_0 \\ \gamma_2/\bar{\Phi}_0 & \gamma_2\big(1/\bar{\Phi}_0 + 1/\bar{\Phi}_2\big) \end{bmatrix}. \tag{S.287}$$

By employing the quasi–spherical parametrization in Eq. (S.169), and assuming that $\Psi_k(\theta,\varphi)$ are well-behaved functions of the angular variables, we could then expand, as done previously for $u(\theta,\varphi)$, the excess areal fractions $\Psi_k$ in a series of surface spherical harmonics, that is,

$$\Psi_k(\theta,\varphi) = \sum_{\ell=0}^{\infty} \Psi^k_\ell(\theta,\varphi). \tag{S.288}$$

Here, the surface harmonics can also be rewritten as

$$\Psi^k_\ell(\theta,\varphi) = \sum_{m=-\ell}^{\ell} \Psi^k_{\ell,m}\, Y_{\ell,m}(\theta,\varphi), \tag{S.289}$$

where $\Psi^k_{\ell,m}$ is the amplitude associated with each of the spherical harmonics $Y_{\ell,m}$. Hence, the governing equation in (S.285) can be re-expressed as follows:

$$\boxed{\frac{\partial \boldsymbol{\Psi}_\ell}{\partial t} = -\mathsf{M}\,\boldsymbol{\Psi}_\ell - \frac{\ell(\ell+1)}{R^2}\mathsf{D}\,\boldsymbol{\Psi}_\ell + \frac{\ell(\ell+2)(\ell^2-1)}{R^3}\boldsymbol{\Omega}\, u_\ell,} \tag{S.290}$$

where we define the vectors $\boldsymbol{\Psi}_\ell = \big(\Psi^1_\ell, \Psi^2_\ell\big)^{\mathsf{T}}$, and we make use of the identity,

$$R^2\Delta^{(0)} Y_{\ell,m}(\theta,\varphi) = -\ell(\ell+1) Y_{\ell,m}(\theta,\varphi)\,, \tag{S.291}$$

in order to expand the Laplacian terms in the spherical harmonics. In particular, we note that

$$[\Delta H]^{(1)} = \sum_{\ell=0}^{\infty} \frac{\ell\,(\ell+2)(\ell^2-1)\, u_\ell}{2R^3}, \tag{S.292}$$

which shows that the zeroth and first harmonics of the Laplacian of the mean curvature vanish identically.

#### E. Area and volume dynamics

The Laplace pressure difference, $P_\vartriangle$, between the two sides of the membrane depends on the osmotic pressure, as well as the active transport of volume due to the biochemical cycles as prescribed by Eq. (S.168). At zeroth order in the perturbation expansion, this becomes

$$\int_{\mathcal{M}} \mathrm{d}S^{(0)} \left\{ v^{(0)} + w_0 \left[\Pi^{(0)}_\vartriangle - P^{(0)}_\vartriangle\right] - \mathbb{V}^{(0)} \right\} = 0. \tag{S.293}$$

Note that the area element $\mathrm{d}S$ also depends on the perturbation parameter $\varepsilon$; namely, we have that

$$\mathrm{d}S^{(0)} = [\sqrt{g}]^{(0)}\, \mathrm{d}\varphi\, \mathrm{d}\theta\ = R^2\sin\theta\ \mathrm{d}\varphi\, \mathrm{d}\theta, \tag{S.294}$$

for the shape parameterization in Eq. (S.169). The total rate of volume flux at zeroth order in $\varepsilon$ is given by

$$\mathbb{V}^{(0)} = \sum_k \bar{\Phi}_k \mathbb{V}^k_{\circlearrowleft}. \tag{S.295}$$

Moreover, the osmotic pressure is found to be

$$\Pi^{(0)}_{\Delta} = k_B T \left[c_+ - \frac{\mathfrak{n}_{\mathcal{S}}}{V^{(0)}}\right] = k_B T \left[c_+ - c_-^{(0)}\right], \tag{S.296}$$

where $\mathfrak{n}_{\mathcal{S}}$ is the number of solutes within the membrane compartment, the volume $V^{(0)} = \frac{4\pi}{3} R^3$, and the number density at zeroth order is $c_-^{(0)} = \mathfrak{n}_{\mathcal{S}} / V^{(0)}$.

As the normal velocity $v^{(0)} = 0$, the integral equation in (S.293) gives us an expression for the Laplace pressure,

$$\boxed{P^{(0)}_{\Delta} = \Pi^{(0)}_{\Delta} - w_0^{-1} \mathbb{V}^{(0)},} \tag{S.297}$$

which in terms of Eq. (S.295) and (S.296) now reads

$$P^{(0)}_{\Delta} = k_B T \left[c_+ - c_-^{(0)}\right] - w_0^{-1} \sum_k \bar{\Phi}_k \mathbb{V}^k_{\circlearrowleft}. \tag{S.298}$$

This is equivalent to volume flux being zero, $j_V^{(0)} = 0$, cf. Eq. (S.227). Note that the active terms in Eq. (S.298) can be seen as an effective renormalization of the solute number densities in the outer and inner solutions.

If the characteristic time corresponding to the passive leakage is much slower than the those associated with the active rates $\mathbb{V}^k_{\circlearrowleft}$ (*i.e.* $w_0 \to \infty$), then Eq. (S.298) retrieves in this limit the *van't Hoff's formula*, cf. Eq. (S.158), that holds only at equilibrium. This gives $\mathbb{V}^{(0)} = \sum_k \bar{\Phi}_k \mathbb{V}^k_{\circlearrowleft} = 0$, which together with Eq. (S.273) implies that the active rates $\mathbb{M}^k_{\circlearrowleft}$ and $\mathbb{V}^k_{\circlearrowleft}$ cannot be independent of each other if a homogeneous steady state is ought to exist, with the mass fractions $\bar{\Phi}_k \neq 0$. This leads to the following linear relationship: $\mathbb{V}^1_{\circlearrowleft}/\mathbb{M}^1_{\circlearrowleft} = \mathbb{V}^2_{\circlearrowleft}/\mathbb{M}^2_{\circlearrowleft}$, required if $w_0^{-1} \to 0$.

The osmotic pressure at first order in $\varepsilon$ can be determined by expanding the volume in Eq. (S.166), namely

$$\Pi^{(1)}_{\Delta} = c_-^{(0)} k_B T \frac{V^{(1)}}{V^{(0)}}, \tag{S.299}$$

where the first order perturbation in the enclosed volume of the membrane can be computed as follows:

$$V^{(1)} = \frac{1}{3} \iint \mathrm{d}\varphi \, \mathrm{d}\theta \, \Big\{ [\sqrt{g}]^{(0)} \boldsymbol{n}^{(1)} \cdot \boldsymbol{R}^{(0)} + [\sqrt{g}]^{(0)} \boldsymbol{n}^{(0)} \cdot \boldsymbol{R}^{(1)} + [\sqrt{g}]^{(1)} \boldsymbol{n}^{(0)} \cdot \boldsymbol{R}^{(0)} \Big\}, \tag{S.300}$$

where the Jacobian term at first order in $\varepsilon$ is given by

$$[\sqrt{g}]^{(1)} = 2R^2 u(\theta, \varphi) \sin\theta. \tag{S.301}$$

Since the lowest perturbation fields of the position vector are given by $\boldsymbol{R}^{(0)} = R\hat{\boldsymbol{r}}$ and $\boldsymbol{R}^{(1)} = R\, u(\theta,\varphi)\, \hat{\boldsymbol{r}}$, and the normal vector $\boldsymbol{n}^{(0)}$ is the radial unit vector $\hat{\boldsymbol{r}}$, the excess volume $V^{(1)}$ can be further simplified to

$$V^{(1)} = R^3 \iint \mathrm{d}\varphi \, \mathrm{d}\theta \; u(\theta, \varphi) \sin\theta = 4\pi R^3 u_0, \tag{S.302}$$

by also using that $\boldsymbol{n}^{(1)}$ is orthogonal to $\hat{\boldsymbol{r}}$, cf. Eq. (S.252). The latter equality in Eq. (S.302) is found by expanding $u(\theta,\varphi)$ in terms of the spherical harmonics $u_\ell(\theta,\varphi)$, and by employing thereupon the orthonormality condition of the spherical harmonics, namely

$$\iint \mathrm{d}\varphi \, \mathrm{d}\theta \sin\theta \, Y_{\ell,m} \, Y_{\ell',m'} = \delta_{\ell,\ell'} \, \delta_{m,m'}. \tag{S.303}$$

Since $\mathcal{Y}_{0,0} = 1/\sqrt{4\pi}$ and the zeroth spherical harmonic is given by $u_0 = u_{0,0}\, \mathcal{Y}_{0,0}$, the second result in Eq. (S.302) readily follows from the orthonormality condition. Thus, the osmotic pressure at first order can be rewritten as:

$$\Pi^{(1)}_{\Delta} = 3\, u_0\, c_-^{(0)} k_B T. \tag{S.304}$$

The total rate of volume flux-rate at first order in the perturbation parameter can be computed by substituting Eq. (S.276) into Eq. (S.161), which yields

$$\mathbb{V}^{(1)} = \sum_k \Psi_k \mathbb{V}^k_{\circlearrowleft}. \tag{S.305}$$

The last two equations can be used to attain the form of the integral condition in Eq. (S.168) at first order in $\varepsilon$, which can be written as

$$\int_{\mathcal{M}} \mathrm{d}S^{(0)} \left\{ v^{(1)} + w_0 \left[\Pi^{(1)}_{\Delta} - P^{(1)}_{\Delta}\right] - \mathbb{V}^{(1)} \right\} = 0, \tag{S.306}$$

where the term pre-multiplying the area element $\mathrm{d}S^{(1)}$ vanishes identically due to the result in Eq. (S.297). As the normal membrane velocity $v^{(1)} = \hat{v}_r^{(1)} = R\, \frac{\partial}{\partial t} u(\theta,\varphi)$, due to Eq. (S.238) and Eq. (S.239), we get

$$\begin{aligned} P^{(1)}_{\Delta} &= \Pi^{(1)}_{\Delta} + \frac{1}{4\pi w_0} \iint \mathrm{d}\varphi \, \mathrm{d}\theta \sin\theta \left( R \frac{\partial u}{\partial t} - \sum_k \Psi_k \mathbb{V}^k_{\circlearrowleft} \right), \\ &= \Pi^{(1)}_{\Delta} + w_0^{-1} \left( R \frac{\partial u_0}{\partial t} - \sum_k \Psi_0^k \mathbb{V}^k_{\circlearrowleft} \right), \end{aligned} \tag{S.307}$$

where the latter equation is found via the orthonormality condition in Eq. (S.303), and thus only the zeroth spherical harmonics of the functions $u(\theta,\varphi)$ and $\Psi_k(\theta,\varphi)$ contribute to the overall integral. Eq. (S.307) is the same as the result derived in Eq. (S.246) by using the Lamb's solution; however, the latter also gives us that

$$\boxed{\frac{\partial u_0}{\partial t} = \frac{1}{2} \mathbb{M}_0^{(1)} = \frac{1}{2} \sum_k \Psi_0^k \, \mathbb{M}^k_{\circlearrowleft},} \tag{S.308}$$

which accounts for the area surface variation in response to a volume change. Thus, we can rewrite the first-order perturbation of the Laplace pressure as

$$P_{\triangle}^{(1)} = \Pi_{\triangle}^{(1)} - w_0^{-1} \sum_k \Psi_0^k \left( \mathbb{V}_{\circlearrowleft}^k - \frac{1}{2} R \mathbb{M}_{\circlearrowleft}^k \right). \tag{S.309}$$

The dynamics of the excess areal fractions $\Psi_k$ in terms of the surface spherical harmonics is given by Eq. (S.290). For $\ell = 0$, this reduces to $\frac{\partial}{\partial t}\Psi_0^k = -\bar{\Phi}_k \mathbb{M}_0^{(1)}$, which gives

$$\frac{\partial \mathbb{M}_0^{(1)}}{\partial t} = -\mathbb{M}^{(0)} \mathbb{M}_0^{(1)} = 0.$$

This means that the net flux-rate at first order in $\varepsilon$ does not change with time, being a fixed constant set by the initial perturbation (say, at $t = 0$). Hence, we find that

$$\mathbb{M}_0^{(1)}(t) = \tilde{\mathbb{M}}_0^{(1)} \quad \text{and} \quad u_0(t) = \tilde{u}_0 + \frac{t}{2} \tilde{\mathbb{M}}_0^{(1)}, \tag{S.310}$$

where $\tilde{u}_0$ and $\tilde{\mathbb{M}}_0^{(1)}$ are the initial values at $t = 0$ for their corresponding functions. Note that the second equation is simply obtained by integrating Eq. (S.308). Thus, $u_0$ grows linearly with time if the initial perturbation in the overall flux of membrane mass is non-zero; meaning that the system is always unstable if $\mathbb{M}_0^{(1)} \neq 0$. To attain a stable steady-state under isotropic perturbations ($\ell = 0$), we require the incoming flux due to fission to balance the outgoing flux of fusion vesicles; that is, $\mathbb{M}_0^{(1)} = 0$.

Since the volume of the homogeneous steady-state is given by $V^{(0)} = \frac{4\pi}{3} R^3$, we find from Eq. (S.302) that

$$\frac{\partial V^{(1)}}{\partial t} = \frac{3}{2} \mathbb{M}_0^{(1)} V^{(0)}. \tag{S.311}$$

Similarly, the total area at zeroth order is $A^{(0)} = 4\pi R^2$, and its first order contribution is given by

$$A^{(1)} = 2R^2 \iint \mathrm{d}\varphi \, \mathrm{d}\theta \; u(\theta, \varphi) \sin\theta = 8\pi R^2 u_0, \tag{S.312}$$

where the former equality follows from Eq. (S.301), and the latter is obtained by employing the orthogonality condition of the surface spherical harmonics. Hence, the rate of change in the total area at first order is found to be

$$\frac{\partial A^{(1)}}{\partial t} = \mathbb{M}_0^{(1)} A^{(0)}. \tag{S.313}$$

Interestingly, we can see that the rate of change in $u_0$ is related to variations in both the area and the volume:

$$\frac{\partial u_0}{\partial t} = \frac{1}{2A^{(0)}} \frac{\partial A^{(1)}}{\partial t} = \frac{1}{3V^{(0)}} \frac{\partial V^{(1)}}{\partial t}. \tag{S.314}$$

If the system is isotropically stable, *i.e.* $\mathbb{M}_0^{(1)} = 0$, then both the area and the volume do not change as a function of time, being entirely set by the initial perturbation $\tilde{u}_0$. On the other hand, in an unstable steady-state with $\mathbb{M}_0^{(1)} \neq 0$, we find that both the area and the volume of the membrane compartment grows linearly with time.

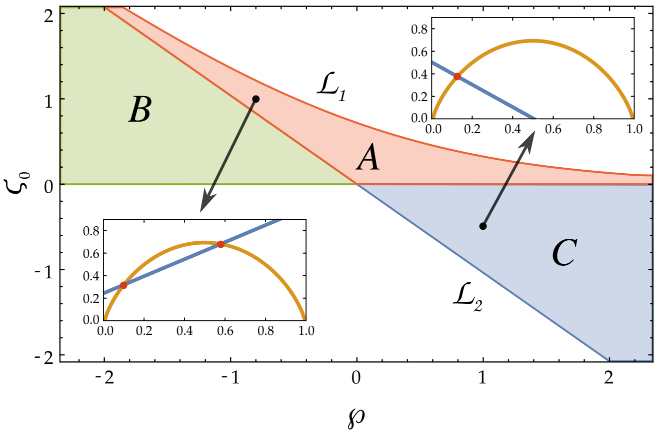


FIG. S.5. Phase diagram of the mass fraction $\bar{\Phi}_0$ associated with the background membrane component, as a function of the parameters $(\varsigma_0, \wp)$. Moreover, the phase boundary $\mathcal{L}_1$ is given by the curve $\varsigma_0 = \ln(1 + e^{-\wp})$, while the boundary $\mathcal{L}_2$ is defined by $\varsigma_0 = -\wp$. In region $A$, we have two distinct real solutions for the fraction $\bar{\Phi}_0$, as shown in the inset plot, which corresponds to the point ($\varsigma_0 = 1$, $\wp = -0.8$), and is indicated by the respective arrow. On the other hand, in the regions $B$ and $C$, we find only one solution, as shown in the other inset plot corresponding to the point ($\varsigma_0 = -0.5$, $\wp = 1$). Here, the insets show the two curves given by the left-hand-side (in blue) and the right-hand-side (in orange) of Eq. (S.278) as a function of $\bar{\Phi}_0$, with the point(s) of intersection (in red) being the corresponding solution(s) for the mass fraction $\bar{\Phi}_0$.

### F. Membrane shape equations

The surface tension $\Sigma(\xi^\alpha)$ acts as a Lagrange multiplier for the membrane area, due to the local incompressibility condition in Eq. (S.71). Since this requirement is satisfied perturbedly, cf. Eq. (S.212), the surface tension must also be expanded in the perturbation parameter $\varepsilon$, that is,

$$\Sigma(\theta, \varphi) = \sum_{\mathtt{i}=0}^{\infty} \varepsilon^{\mathtt{i}} \, \Sigma^{(\mathtt{i})}(\theta, \varphi), \tag{S.315}$$

thus ensuring the local incompressibility condition on the membrane at all orders in the perturbation expansion. Their explicit expressions must be self-consistently determined from the local stress balance tangential to the membrane. In other words, the incompressibility condition in Eq. (S.212) fixes the local surface tension $\Sigma^{(\mathtt{i})}$.

#### 1. At zeroth order

By utilizing the shape parameterization in Eq. (S.169), as well as the expansion in Eq. (S.276) of the mass fractions, the equations of motion of the membrane may be computed at each order in the perturbation parameter. Along the tangential direction, we find that $[f_\alpha]^{(0)} = \Sigma_{;\alpha}^{(0)}$, which together with the result in Eq. (S.236), yields

$$\Sigma_{;\alpha}^{(0)} = 0. \tag{S.316}$$

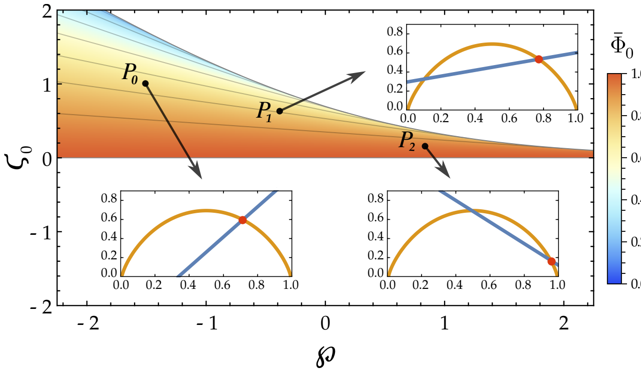


FIG. S.6. Density plot of the background mass fraction $\bar{\Phi}_0$ as a function of the parameters $\varsigma_0$ and $\wp$, showing only one of the roots of Eq. (S.278). Specifically, we display the root that lives solely in the union set of $A$ and $B$ regions (by using the notation introduced in Fig. S.3), namely $0 \leq \varsigma_0 \leq \ln\left(1 + e^{-\wp}\right)$ for any real value of $\wp$. Furthermore, we illustrate a few explicit points on this phase-space $(\varsigma_0, \wp)$. The point $P_0 = (1, -1.5)$ corresponds to a single isolated solution, as displayed by the intersection point (in red) of the respective inset plot, where the blue and orange curves are as those defined in Fig. S.3. On the other hand, the points $P_1 = (0.6, 0.3)$ and $P_2 = (0.1, 1)$, as shown in their insets, correspond to two solutions, but the chosen root is the right-most one. In Fig. S.5, the same points are depicted again, with the other root of $\bar{\Phi}_0$ being plotted.

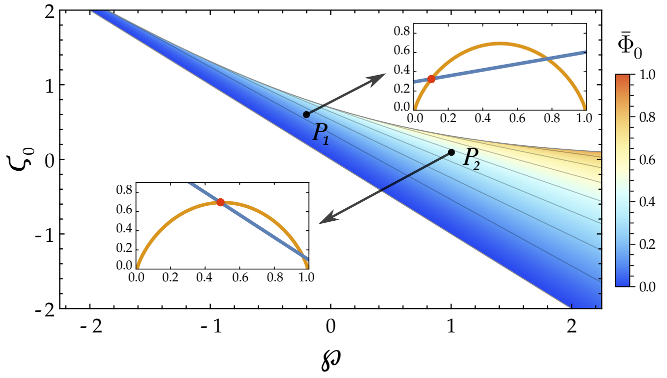


FIG. S.7. Density plot of the mass fraction $\bar{\Phi}_0$ on the phase-space of $\varsigma_0$ and $\wp$, where we display only one of the roots of Eq. (S.278); namely, the root that is defined on the union of $A$ and $C$ regions (using the notation of Fig. S.3). Two explicit points are shown, $P_1 = (0.6, 0.3)$ and $P_2 = (0.1, 1)$, which are the same as those depicted in Fig. S.4. Their respective inset plots show that this root of $\bar{\Phi}_0$ corresponds to the left-most solution, as illustrated by the red points, with the blue and orange curves as defined before in Fig. S.3. This particular root is unphysical since it does not respect the requirement that the mass fraction $\bar{\Phi}_0$ must tend to unity as we approach equilibrium, *i.e.* in the limit of both $\varsigma_0 \to 0$ and $\wp \to 0$.

This means that the surface tension at zeroth order is a scalar quantity, which has a homogeneous value $\Sigma^{(0)}$. On the other hand, by using Eq. (S.122), the zeroth order contribution to the normal body force is found to be

$$f^{(0)} = \sum_k \frac{2\bar{\Phi}_k}{R}\left[\mathbb{P}^k_{\circlearrowleft} + \frac{k_BT}{b_0}\left(E_k + \ln\frac{\bar{\Phi}_k}{\bar{\Phi}_0}\right) - \frac{2\kappa C_k + 3\mathbb{Q}^k_{\circlearrowleft}}{2R}\right] + \frac{2}{R}\left[\Sigma^{(0)} + \frac{k_BT}{b_0}(\mathcal{E}_0 + \ln\bar{\Phi}_0) - \frac{\kappa\mathcal{C}_0}{R}\right]. \quad \text{(S.317)}$$

Here, the body force $f^{(0)}$ along the normal is given by the pressure difference across the interface, *i.e.* $f^{(0)} = -P^{(0)}_{\Delta}$. By also using Eq. (S.297), then Eq. (S.317) becomes

$$\Pi^{(0)}_{\Delta} + \sum_k \frac{2\bar{\Phi}_k}{R}\left[\mathbb{P}^k_{\circlearrowleft} + \frac{k_BT}{b_0}\left(E_k + \ln\frac{\bar{\Phi}_k}{\bar{\Phi}_0}\right) - \frac{2\kappa C_k + 3\mathbb{Q}^k_{\circlearrowleft}}{2R}\right] + \frac{2}{R}\left(\bar{\Sigma}^{(0)} + \frac{k_BT}{b_0}\ln\bar{\Phi}_0 - \frac{\kappa\mathcal{C}_0}{R}\right) = \sum_k \frac{\bar{\Phi}_k\mathbb{V}^k_{\circlearrowleft}}{w_0}, \quad \text{(S.318)}$$

where we define $\bar{\Sigma}^{(0)} = \Sigma^{(0)} + \frac{k_BT}{b_0}\mathcal{E}_0$, and $\Pi^{(0)}_{\Delta}$ is the osmotic pressure, whose expression is found in Eq. (S.296).

Note that the steady state values of the mass fractions $\bar{\Phi}_k$ are given by Eq. (S.274), and therefore $\bar{\Phi}_k \propto \left(1 - \bar{\Phi}_0\right)$. Hence, if the surface tension $\Sigma^{(0)}$ is set at a fixed value, then Eq. (S.318) gives us with a transcendental equation for the mass fraction $\bar{\Phi}_0$ that can be rewritten as follows:

$$\boxed{\varsigma_0 + \wp(1 - \bar{\Phi}_0) = -\bar{\Phi}_0\ln\bar{\Phi}_0 - (1 - \bar{\Phi}_0)\ln(1 - \bar{\Phi}_0),} \quad \text{(S.319)}$$

where the parameters $\varsigma_0$ and $\wp$ are given by

$$\varsigma_0 = \frac{b_0}{k_BT}\left(\bar{\Sigma}^{(0)} + \frac{R}{2}\Pi^{(0)}_{\Delta} - \frac{\kappa\mathcal{C}_0}{R}\right), \quad \text{(S.320)}$$

and $\wp = \wp_1 + \wp_2$, respectively, and we define that

$$\wp_k = \frac{|\mathbb{M}^{k'}_{\circlearrowleft}|}{|\mathbb{M}^k_{\circlearrowleft}| + |\mathbb{M}^{k'}_{\circlearrowleft}|}\left[E_k + \ln\left(\frac{|\mathbb{M}^{k'}_{\circlearrowleft}|}{|\mathbb{M}^k_{\circlearrowleft}| + |\mathbb{M}^{k'}_{\circlearrowleft}|}\right) + \frac{b_0}{k_BT}\left(\mathbb{P}^k_{\circlearrowleft} - \frac{R\mathbb{V}^k_{\circlearrowleft}}{2w_0} - \frac{2\kappa C_k + 3\mathbb{Q}^k_{\circlearrowleft}}{2R}\right)\right], \quad \text{(S.321)}$$

with the index $k \neq k'$. The right-hand-side of Eq. (S.319) is greater than zero for all $\bar{\Phi}_0 \in (0, 1)$. At thermodynamic equilibrium, we have that the mass fractions $\bar{\Phi}_k = 0$, and therefore $\bar{\Phi}_0 = 1$. By applying this limit in Eq. (S.319), we get that $\varsigma_0 = 0$ at equilibrium; namely, we retrieve a version of the *Young–Laplace* equation:

$$\Pi^{(0)}_{\Delta} = 2H^{(0)}\bar{\Sigma}^{(0)} + 2\kappa\mathcal{C}_0K^{(0)}, \quad \text{(S.322)}$$

with the Laplace pressure being $P^{(0)}_{\Delta} = \Pi^{(0)}_{\Delta}$, where $H^{(0)}$ and $K^{(0)}$ is the mean and Gaussian curvature at zeroth order in the perturbation expansion. Moreover, we have that $\wp = \wp_1 = \wp_2 = 0$ at equilibrium, since the loop currents $\mathbb{J}^k_{\circlearrowleft}$ vanish identically (and $\mathcal{R}^k_{\circlearrowleft} = 0$, as well).

In Figure S.5, we display the region of the parameter space $(\varsigma_0, \wp)$ in which Eq. (S.319) gives a real solution for the mass fraction $\bar{\Phi}_0$. The upper phase-boundary, that is denoted by $\mathcal{L}_1$ in Fig. S.5, is found by seeking the points where the tangent of the right-hand-side of Eq. (S.319) is the same as the slope of the left-hand-side of Eq. (S.319). This requirement yields that $\bar{\Phi}_0 = 1/(1+e^{-\wp})$, which by substitution into Eq. (S.319) gives us the phase-boundary curve $\mathcal{L}_1 = \{(\varsigma_0, \wp) \,|\, \varsigma_0 = \ln(1+e^{-\wp})\}$. Hence, in order to find a solution to Eq. (S.319), we need that

$$\varsigma_0 \le \ln\big(1+e^{-\wp}\big), \quad \text{for any } \wp \in \mathbb{R}. \tag{S.323}$$

To get the lower bounds, we seek the straight lines that are given by the left-hand-side of Eq. (S.319), which cross the extreme points $\bar{\Phi}_0 = 0$ and $\bar{\Phi}_0 = 1$. For $\wp \le 0$, we find the lower bound of $\varsigma_0$ to be the line $\varsigma_0 = 0$, whilst, if $\wp > 0$, we obtain the curve $\mathcal{L}_2 = \{(\varsigma_0, \wp) \,|\, \varsigma_0 = -\wp\}$ as the lower phase-boundary. Therefore, we require

$$\begin{cases} \varsigma_0 \ge -\wp, & \text{if } \wp \ge 0, \\ \varsigma_0 \ge 0, & \text{if } \wp < 0. \end{cases} \tag{S.324}$$

The inequalities in Eq. (S.323) and (S.324) defines the phase-region where Eq. (S.319) has at least one real solution. However, within this domain, we identify a region in which two real district solutions can exist, and which we denote by $A$ in Fig. S.5. This is strictly defined by

$$A = \left\{ (\varsigma_0, \wp) \left| \begin{matrix} 0 \le \varsigma_0 < \ln(1+e^{-\wp}), \text{ if } \wp < 0 \\ -\wp \le \varsigma_0 < \ln(1+e^{-\wp}), \text{ if } \wp \ge 0 \end{matrix} \right. \right\} \tag{S.325}$$

excluding curve $\mathcal{L}_1$ where the two solutions are identical.

In the other regions, indicated by the letters $B$ and $C$, we find that Eq. (S.323) has only a single real solution, which are defined as follows:

$$B = \{(\varsigma_0, \wp) \,|\, 0 < \varsigma_0 < -\wp, \text{ with } \wp \in \mathbb{R}\}, \tag{S.326}$$

$$C = \{(\varsigma_0, \wp) \,|\, -\wp < \varsigma_0 < 0, \text{ with } \wp \in \mathbb{R}\}. \tag{S.327}$$

By numerically tracing out the two roots, we find that one of the roots is purely defined on the union of $A$ and $C$ (see Fig. S.6), while the other belongs solely to the union of $A$ and $B$ (see Fig. S.7). The latter solution is found to be unphysical, as it does not respect the requirement that we must have $\wp = 0$ and $\bar{\Phi}_0 = 1$ at equilibrium, which is respected only by the former solution, see Fig. S.6.

As a result, the necessary condition for the existence of a homogeneous steady-state solution is given by

$$\boxed{0 \le \varsigma_0 \le \ln\big(1+e^{-\wp}\big),} \tag{S.328}$$

for any real value of the activity coefficient $\wp$. In other words, provided that the parameters $\varsigma_0$ and $\wp$ respect this condition, then Eq. (S.319) allows us to find a unique value for the mass fraction $\bar{\Phi}_0$, which in turn determines the values of the other mass fractions $\bar{\Phi}_k$ via Eq. (S.274).

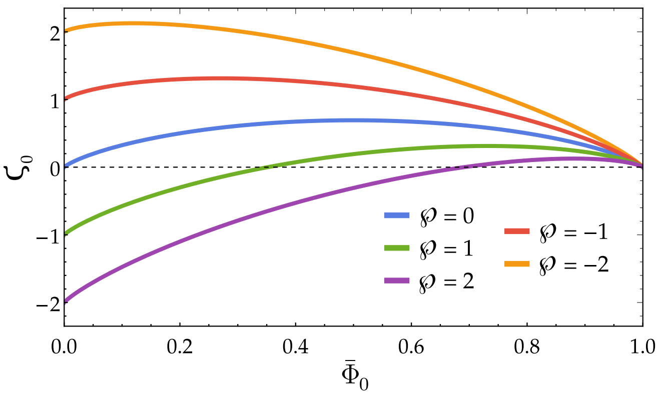


FIG. S.8. The reduced surface tension $\varsigma_0$ as a function of the mass fraction $\bar{\Phi}_0$ of the background membrane component in terms of various values of the activity coefficient $\wp$.

The stability condition in Eq. (S.328) applies only if the surface tension at zeroth order in the perturbation is fixed, and $\bar{\Phi}_0$ is allowed to adjust. Instead, if we consider a thermodynamic ensemble in which the mass fraction $\bar{\Phi}_0$ of the background component is prescribed, then the surface tension is given by the force-balance in Eq. (S.319). The dependence of the reduced surface tension $\varsigma_0$ on the homogeneous mass fraction $\bar{\Phi}_0$ is shown in Fig. (S.8), which shows that $\varsigma_0$ vanishes in the limit of $\Phi_0 \to 1$, as the system approaches thermodynamic equilibrium.

#### 2. *At first order*

Neglecting the membrane inertial terms in Eq. (S.120) as before, the equation of motion at first order in $\varepsilon$, along the tangential directions, can be written as follows:

$$[f^\alpha]^{(1)} = \sum_k -\Psi_k^{;\alpha}\left[\mathbb{P}_\circlearrowleft^k + \frac{k_B T}{b_0}\left(E_k + \ln\frac{\bar{\Phi}_k}{\bar{\Phi}_0}\right) - \frac{2\kappa C_k + \mathbb{Q}_\circlearrowleft^k}{R}\right] - \big[\Sigma^{(1)}\big]^{;\alpha} - \sum_k \bar{\Phi}_k \mathbb{Q}_\circlearrowleft^k \big[2H^{(1)}\big]^{;\alpha}, \tag{S.329}$$

At first order, the surface contribution to the body force is given by $\boldsymbol{f}_s^{(1)} = \boldsymbol{e}_\alpha^{(0)} [f^\alpha]^{(1)}$, since $[f^\alpha]^{(0)} = 0$. Hence,

$$\boldsymbol{f}_s^{(1)} = \sum_{\ell=1}^{\infty} \boldsymbol{\nabla}_s \left\{ -\frac{\Sigma_\ell^{(1)}}{R} + \sum_k \bar{\Phi}_k \mathbb{Q}_\circlearrowleft^k \frac{(\ell-1)(\ell+2)\, u_\ell}{R^2} - \frac{\Psi_\ell^k}{R}\left[\mathbb{P}_\circlearrowleft^k + \frac{k_B T}{b_0}\left(E_k + \ln\frac{\bar{\Phi}_k}{\bar{\Phi}_0}\right) - \frac{2\kappa C_k + \mathbb{Q}_\circlearrowleft^k}{R}\right]\right\}, \tag{S.330}$$

with $\Sigma^{(1)}$ and $H^{(1)}$ being expanded in the basis of surface spherical harmonics. The result in Eq. (S.330) tells us that the body force $\boldsymbol{f}_s^{(1)}$ can be expressed as a surface gradient of a scalar function that lives solely on the membrane. At the same time, $\boldsymbol{f}_s^{(1)}$ can be written in terms of the ambient fluid stresses acting across the membrane surface, as derived in Eq. (S.272) from Lamb's solution. This expression is also a surface gradient of a function, and due to the orthogonality of spherical harmonics, each surface mode associated with the two expressions of $\boldsymbol{f}_S^{(1)}$ can be equated accordingly. Namely, we obtain that

$$\frac{\Sigma_\ell^{(1)}}{R} = u_\ell \sum_k \bar{\Phi}_k \left[\frac{(\ell-1)(\ell+2)\,\mathbb{Q}_{\circlearrowright}^k}{R^2}\right] + \frac{\eta(2\ell+1)}{\ell(\ell+1)}\left[\frac{\partial u_\ell}{\partial t} - \sum_k \Psi_\ell^k\left(2\mathbb{M}_{\circlearrowright}^k - \frac{3\mathbb{V}_{\circlearrowright}^k}{R} - \frac{\ell(\ell+1)\mathbb{Q}_{\circlearrowright}^k}{\eta R^2}\right)\right] - \sum_k \frac{\Psi_\ell^k}{R}\left[\mathbb{P}_{\circlearrowright}^k + \frac{k_B T}{b_0}\left(E_k + \ln\frac{\bar{\Phi}_k}{\bar{\Phi}_0}\right) - \frac{2\kappa C_k + \mathbb{Q}_{\circlearrowright}^k}{R}\right], \tag{S.331}$$

by using the explicit forms of $\mathcal{X}_\ell^{(1)} = R\frac{\partial u_\ell}{\partial t} - \sum_k \Psi_\ell^k \mathbb{V}_{\circlearrowright}^k$, and $\mathcal{Y}_\ell^{(1)} = \sum_k \Psi_\ell^k\left[2\mathbb{V}_{\circlearrowright}^k - R\mathbb{M}_{\circlearrowright}^k + \frac{\ell(\ell+1)\mathbb{Q}_{\circlearrowright}^k}{2\eta R}\right]$.

The equation of motion along the normal direction at first order in $\varepsilon$ can be determined from Eq. (S.122). The normal body force on the membrane at first order in the perturbation expansion, $f^{(1)}$, can be computed as:

$$f^{(1)} = \frac{2\Sigma^{(1)}}{R} - 2H^{(1)}\left\{\left(\bar{\Sigma}^{(0)} - \frac{2\kappa\mathcal{C}_0}{R} + \frac{k_B T}{b_0}\ln\bar{\Phi}_0\right) + \sum_k \bar{\Phi}_k\left[\mathbb{P}_{\circlearrowright}^k + \frac{k_B T}{b_0}\left(E_k + \ln\frac{\bar{\Phi}_k}{\bar{\Phi}_0}\right) - \frac{2\kappa C_k + 3\mathbb{Q}_{\circlearrowright}^k}{R}\right]\right\} + \sum_k \frac{2\Psi_k}{R}\left[\mathbb{P}_{\circlearrowright}^k + \frac{k_B T}{b_0}\left(E_k + \ln\frac{\bar{\Phi}_k}{\bar{\Phi}_0}\right) - \frac{2\kappa C_k + 3\mathbb{Q}_{\circlearrowright}^k}{2R}\right] + \sum_k \Delta^{(0)}\Psi_k\left(\kappa C_k - \frac{1}{2}\mathbb{Q}_{\circlearrowright}^k\right) + 2\kappa[\Delta H]^{(1)}, \tag{S.332}$$

where we use that $H^2 - K = 0 + \mathcal{O}[\varepsilon^2]$, which gives us the Gaussian curvature at first order: $K^{(1)} = 2H^{(0)}H^{(1)}$. The term in the curly brackets can be further simplified by using the expression in Eq. (S.317), which yields

$$f^{(1)} = 2H^{(1)}\left[\frac{R}{2}P_\Delta^{(0)} + \frac{\kappa\mathcal{C}_0}{R} + \sum_k \frac{\bar{\Phi}_k\left(2\kappa C_k + 3\mathbb{Q}_{\circlearrowright}^k\right)}{2R}\right] + \sum_k \frac{2\Psi_k}{R}\left[\mathbb{P}_{\circlearrowright}^k + \frac{k_B T}{b_0}\left(E_k + \ln\frac{\bar{\Phi}_k}{\bar{\Phi}_0}\right) - \frac{2\kappa C_k + 3\mathbb{Q}_{\circlearrowright}^k}{2R}\right] + \sum_k \Delta^{(0)}\Psi_k\left(\kappa C_k - \frac{\mathbb{Q}_{\circlearrowright}^k}{2}\right) + 2\kappa[\Delta H]^{(1)} + \frac{2\Sigma^{(1)}}{R}, \tag{S.333}$$

where $P_\Delta^{(0)}$ is the Laplace pressure at zeroth order in $\varepsilon$. Moreover, this can be expanded as before in the basis of spherical harmonics, which now reads:

$$f^{(1)} = \sum_{\ell=0}^{\infty}\left\{\frac{2\Sigma_\ell^{(1)}}{R} + \frac{\ell(\ell+2)(\ell^2-1)\kappa u_\ell}{R^3} - (\ell-1)(\ell+2)u_\ell\left[\frac{P_\Delta^{(0)}}{2} + \frac{\kappa\mathcal{C}_0}{R^2} + \sum_k \frac{\bar{\Phi}_k\left(2\kappa C_k + 3\mathbb{Q}_{\circlearrowright}^k\right)}{2R^2}\right] + \sum_k \frac{2\Psi_\ell^k}{R}\left[\mathbb{P}_{\circlearrowright}^k + \frac{k_B T}{b_0}\left(E_k + \ln\frac{\bar{\Phi}_k}{\bar{\Phi}_0}\right) - \frac{2\kappa C_k + 3\mathbb{Q}_{\circlearrowright}^k}{2R}\right] - \sum_k \frac{\ell(\ell+1)\Psi_\ell^k\left(2\kappa C_k - \mathbb{Q}_{\circlearrowright}^k\right)}{2R^2}\right\}. \tag{S.334}$$

For an isotropically stable solution, cf. Eq. (S.311) and Eq. (S.313), we must demand that $\ell = 0$ contribution to the vesicular flux to be zero: $\mathbb{M}_0^{(1)} = \sum_k \Psi_0^k \mathbb{M}_{\circlearrowright}^k = 0$. Hence, Eq. (S.290) for $\ell = 0$ becomes: $\dot{\Psi}_0^1 = \dot{\Psi}_0^2 = 0$. This implies that the excess areal fractions $\Psi_0^k$ are constants independent of time, which are set by the initial perturbation. Since we perturb about the homogeneous steady-states $\bar{\Phi}_k$, we can choose that $\Psi_0^1 = \Psi_0^2 = 0$, such that the uniform area fractions $\bar{\Phi}_k$ are the starting points. In a similar way, we can also choose that $u_0 = 0$, cf. Eq. (S.246). Using Eq. (S.261) and (S.309), we find that the zeroth mode of Eq. (S.334) simply reduces to $\Sigma_0^{(1)} = 0$, as $P_\Delta^{(1)} = \Pi_\Delta^{(1)} = 0$. Hence, the sum in Eq. (S.334) may start from the first surface mode ($\ell = 1$) without any loss of generality. Nonetheless, this is a simplification, as in general $\Sigma_0^{(1)}$ is linearly related to the initial value of $u_0$ and one of the initial values of either $\Psi_0^1$ or $\Psi_0^2$, as the latter are linearly dependent via $\mathbb{M}_0^{(1)} = 0$.

By employing Eq. (S.261) and the orthogonality of the spherical harmonics, each of the surface harmonics can be separately analyzed; namely, for every $\ell \geq 1$, we have

$$\frac{\eta(2\ell+1)}{\ell(\ell+1)}\left\{[3 - 2\ell(\ell+1)]\frac{\partial u_\ell}{\partial t} + \frac{3 + 2\ell(\ell+1)}{R}\sum_k \Psi_\ell^k \mathbb{V}_{\circlearrowright}^k - \sum_k 3\Psi_\ell^k\left(\mathbb{M}_{\circlearrowright}^k - \frac{\ell(\ell+1)\mathbb{Q}_{\circlearrowright}^k}{2\eta R^2}\right)\right\} = \frac{2\Sigma_\ell^{(1)}}{R} + - (\ell-1)(\ell+2)u_\ell\left[\frac{P_\Delta^{(0)}}{2} + \frac{\kappa\mathcal{C}_0}{R^2} + \sum_k \frac{\bar{\Phi}_k\left(2\kappa C_k + 3\mathbb{Q}_{\circlearrowright}^k\right)}{2R^2}\right] + \frac{\ell(\ell+2)(\ell^2-1)\kappa u_\ell}{R^3} + \sum_k \frac{2\Psi_\ell^k}{R}\left[\mathbb{P}_{\circlearrowright}^k + \frac{k_B T}{b_0}\left(E_k + \ln\frac{\bar{\Phi}_k}{\bar{\Phi}_0}\right) - \frac{2(\ell^2+\ell+2)\kappa C_k - (\ell-2)(\ell+3)\mathbb{Q}_{\circlearrowright}^k}{4R}\right]. \tag{S.335}$$

The tangential stress balance in Eq. (S.331) provides us an explicit expression for $\Sigma_\ell^{(1)}$, which allows us to obtain the following shape equation:

$$\boxed{\begin{aligned}\frac{\partial u_\ell}{\partial t} &= \frac{\kappa u_\ell(2\ell+\ell^2)(\ell^2-1)(R\mathcal{C}_0-\ell-\ell^2)}{\eta R^3(2\ell+1)(2\ell^2+2\ell-1)} + \frac{(2\ell+\ell^2)(\ell^2-1)\,u_\ell}{2\eta(2\ell+1)(2\ell^2+2\ell-1)}\left[\Pi_\Delta^{(0)} - \frac{1}{w_0}\sum_k \bar{\Phi}_k\,\mathbb{V}^k_{\circlearrowleft}\right] \\ &+ \sum_k \Psi^k_\ell\left[\frac{R\,\mathbb{M}^k_{\circlearrowleft} + (2\ell^2+2\ell-3)\,\mathbb{V}^k_{\circlearrowleft} + \frac{\ell(\ell+1)}{2\eta R}\,\mathbb{Q}^k_{\circlearrowleft}}{R\,(2\ell^2+2\ell-1)}\right] - \sum_k \frac{(2\ell+\ell^2)(\ell^2-1)\big(\Psi^k_\ell + u_\ell\,\bar{\Phi}_k\big)\left(\mathbb{Q}^k_{\circlearrowleft} - 2\kappa C_k\right)}{2\eta R^2(2\ell+1)\,(2\ell^2+2\ell-1)},\end{aligned}} \tag{S.336}$$

while the surface tension, associated with each of the surface spherical harmonics, is given by:

$$\begin{aligned}\Sigma^{(1)}_\ell &= \frac{\kappa u_\ell(\ell+2)(\ell-1)(R\mathcal{C}_0-\ell-\ell^2)}{R^2(2\ell^2+2\ell-1)} - \sum_k \Psi^k_\ell\left[\mathbb{P}^k_{\circlearrowleft} + \frac{k_BT}{b_0}\left(E_k + \ln\frac{\bar{\Phi}_k}{\bar{\Phi}_0}\right) + \frac{(3+2\ell)\big(4\ell^2-1\big)\big(\eta R\,\mathbb{M}^k_{\circlearrowleft} - 2\eta\mathbb{V}^k_{\circlearrowleft}\big)}{\ell(\ell+1)(2\ell^2+2\ell-1)}\right] \\ &+ \sum_k \frac{2\kappa C_k\left[\big(5\ell^2+5\ell-4\big)\Psi^k_\ell + (\ell-1)(\ell+2)\,u_\ell\,\bar{\Phi}_k\right] + \mathbb{Q}^k_{\circlearrowleft}\left[\big(8\ell^3+15\ell^2+\ell-3\big)\Psi^k_\ell + \big(4\ell^4+8\ell^3-7\ell^2-11\ell+6\big)u_\ell\,\bar{\Phi}_k\right]}{2R\,(2\ell^2+2\ell-1)} \\ &+ \frac{(\ell+2)(\ell-1)Ru_\ell}{(4\ell^2+4\ell-2)}\left[\Pi^{(0)}_\Delta - \frac{1}{w_0}\sum_k \bar{\Phi}_k\,\mathbb{V}^k_{\circlearrowleft}\right],\end{aligned} \tag{S.337}$$

which shows that both $\mathbb{M}^k_{\circlearrowleft}$ and $\mathbb{P}^k_{\circlearrowleft}$ tend to increase the surface tension $\Sigma^{(1)}_\ell$ for $k=1$, as a result of the fission events, and reduce the surface tension for $k=2$, corresponding to the fusion events. This readily follows from the sign convention that $\mathrm{sgn}(\mathbb{M}^k_{\circlearrowleft}) = \mathrm{sgn}(\mathbb{P}^k_{\circlearrowleft}) = (-1)^k$.

#### G. Linear stability analysis

The dynamics of the excess areal fractions $\Psi^k_\ell$ is given by the vector equation (S.290), whilst the shape equation is shown in Eq. (S.336). Hence, these equations form a closed system of first-order linear differential equations, and the stability of the corresponding solutions can be analyzed for all spherical harmonics with $\ell \geq 1$; namely, by expressing the system of equations in a matrix form:

$$\frac{\partial}{\partial t}\Big[u_\ell \;\; \Psi^1_\ell \;\; \Psi^2_\ell\Big]^{\mathsf{T}} = \mathsf{S}_\ell\,\Big[u_\ell \;\; \Psi^1_\ell \;\; \Psi^2_\ell\Big]^{\mathsf{T}}, \tag{S.338}$$

where the superscript $\mathsf{T}$ denotes a transpose, and $\mathsf{S}_\ell$ is the stability matrix associated with each $\ell$-th mode.

For the first surface harmonics ($\ell=1$), the membrane shape equation is simply given by

$$\frac{\partial u_1}{\partial t} = \sum_k \Psi_k\left[\frac{R\,\mathbb{M}^k_{\circlearrowleft} + \mathbb{V}^k_{\circlearrowleft}}{3R} - \frac{\mathbb{Q}^k_{\circlearrowleft}}{3\eta R^2}\right], \tag{S.339}$$

whereas the membrane compositional equation, written in the vector form $\boldsymbol{\Psi}_\ell = \big(\Psi^1_\ell, \Psi^2_\ell\big)^{\mathsf{T}}$, reduces to

$$\frac{\partial \boldsymbol{\Psi}_1}{\partial t} = -\mathsf{M}\boldsymbol{\Psi}_1 - \frac{2}{R^2}\,\mathsf{D}\boldsymbol{\Psi}_1. \tag{S.340}$$

As a result, the stability matrix $\mathsf{S}_1$ is found to be

$$\begin{bmatrix} 0 & -\frac{\mathbb{Q}^1_{\circlearrowleft}}{3\eta R^2} + \frac{R\,\mathbb{M}^1_{\circlearrowleft} + \mathbb{V}^1_{\circlearrowleft}}{3R} & -\frac{\mathbb{Q}^2_{\circlearrowleft}}{3\eta R^2} + \frac{R\,\mathbb{M}^2_{\circlearrowleft} + \mathbb{V}^2_{\circlearrowleft}}{3R} \\ 0 & -\frac{\bar{\Phi}_0(\bar{\Phi}_1 R)^2\,\mathbb{M}^1_{\circlearrowleft} + 2\gamma_1\big(1-\bar{\Phi}_2\big)}{\bar{\Phi}_0\bar{\Phi}_1 R^2} & -\frac{2\gamma_1 + \bar{\Phi}_0\bar{\Phi}_1 R^2\,\mathbb{M}^2_{\circlearrowleft}}{\bar{\Phi}_0 R^2} \\ 0 & -\frac{2\gamma_2 + \bar{\Phi}_0\bar{\Phi}_2 R^2\,\mathbb{M}^1_{\circlearrowleft}}{\bar{\Phi}_0 R^2} & -\frac{\bar{\Phi}_0(\bar{\Phi}_2 R)^2\,\mathbb{M}^2_{\circlearrowleft} + 2\gamma_2\big(1-\bar{\Phi}_1\big)}{\bar{\Phi}_0\bar{\Phi}_2 R^2} \end{bmatrix}$$

from which the corresponding eigenvalues of the matrix can be computed. We denote the matrix elements of $\mathsf{S}_\ell$ to be $s^{(i,j)}_\ell$, where $i$ and $j$ are the positions of the rows and columns, respectively, (for instance, $s^{(1,1)}_1 = s^{(2,1)}_1 = 0$). We define $\lambda_\ell$ to be the eigenvalues of $\mathsf{S}_\ell$, which satisfy

$$\det(-\lambda_\ell \mathsf{I} + \mathsf{S}_\ell) = 0, \tag{S.341}$$

with $\mathsf{I}$ as the identity matrix. Hence, the eigenvalues of the stability matrix $\mathsf{S}_1$ can be computed as the roots of the following polynomial:

$$\lambda_1^3 - \lambda_1^2\left(s^{(2,2)}_1 + s^{(3,3)}_1\right) + \lambda_1\left(s^{(2,2)}_1 s^{(3,3)}_1 - s^{(2,3)}_1 s^{(3,2)}_1\right) = 0.$$

Note that one of the eigenvalues is zero, while the other two do not depend on the first row elements of $\mathsf{S}_1$.

Although we could obtain their expressions in exact form, it is more instructive to work with a number of reduced dimensionless parameters. We choose the surface diffusivities $\gamma_k$ to be the same for both the fissogens and fusogens, namely $\gamma_k = \gamma$. We also re-scale the mass flux-rates $\mathbb{M}^k_{\circlearrowleft}$ by the membrane diffusion time, that is,

$$\mathbb{M}^1_{\circlearrowleft} = -\frac{\gamma\,A}{R^2}, \quad \text{and} \quad \mathbb{M}^2_{\circlearrowleft} = \zeta\,\frac{\gamma\,A}{R^2}, \tag{S.342}$$

where $A$ is a dimensionless coefficient that captures the number of fission events within the diffusion time $R^2/\gamma$, while $\zeta$ is the ratio of the magnitude of the mass fluxes; namely, $\zeta = -\mathbb{M}^2_{\circlearrowleft}/\mathbb{M}^1_{\circlearrowleft}$, describing the strength of the fusion membrane flux compared to the fission flux. Estimates of $A$ and $\zeta$ are shown in Table II.

The steady-state mass fractions, $\bar{\Phi}_k$, from Eq. (S.274), can be rewritten as follows:

$$\bar{\Phi}_1 = \zeta \, \frac{1-\bar{\Phi}_0}{1+\zeta}, \quad \text{and} \quad \bar{\Phi}_2 = \frac{1-\bar{\Phi}_0}{1+\zeta}, \tag{S.343}$$

where $\bar{\Phi}_0$ is mass fraction of the background component. This shows that the mass areal fractions $\bar{\Phi}_k$ are independent of $A$. For simplicity, we also neglect the spontaneous curvature terms and the volume fluxes, by setting both of them to zero; namely, $C_k = 0$ and $\mathbb{V}^k_{\circlearrowleft} = 0$. In other words, we choose to focus only on the membrane mass exchange and the active momentum transfer. Since only the the second moments $\mathbb{Q}^k_{\circlearrowleft}$ directly contribute to the shape equation in Eq. (S.336), we choose

$$\mathbb{Q}^1_{\circlearrowleft} = -\gamma\eta \, Q_1, \quad \text{and} \quad \mathbb{Q}^2_{\circlearrowleft} = \gamma\eta \, Q_2, \tag{S.344}$$

rescaling them by a characteristic force $\gamma\eta$, with $Q_k$ dimensionless. Here, the choice of the minus sign in the first equation is to reflect the sign convention in Eq. (S.109), which is motivated by the dipolar point-force model from Eq. (S.87). However, we argue that in general the the sign of the second moments is completely independent from the first moments, and thus $Q_k$ can in general be any real number. Here we consider two limiting cases. First, we take $Q_1 = Q_2 = Q$, with $Q$ as some real number, and thus the magnitude of the moments $\mathbb{Q}^k_{\circlearrowleft}$ is the same, but have opposite signs. Second, we assume that both of them have the same sign and magnitude, that is, we take $Q_1 = -Q_2 = Q$, with $Q$ as any real. Estimates of the range of $Q$ is shown in Table II.

Using this rescaling of parameters, the nonzero eigenvalues of $\mathsf{S}_1$ are found to be:

$$\lambda_1^{\pm} = \frac{-2\zeta-(1+\zeta^2)\,\bar{\Phi}_0 \pm \sqrt{4\zeta^2+2\left[A(\zeta-1)-4\right]\zeta^2\,\bar{\Phi}_0 + \left[1+2(1+2m)\zeta^2-4A\zeta^3+\zeta^4\right]\bar{\Phi}_0^2 + 2A(\zeta-1)\zeta^2\,\bar{\Phi}_0^3}}{\zeta\,\bar{\Phi}_0(1-\bar{\Phi}_0)}. \tag{S.345}$$

This shows that the real part of $\lambda_1^- < 0$ for any value of parameters, and only $\lambda_1^+$ can switch sign, and it can be strictly greater than zero provided that the term within the square-root term is a positive real number. Hence, this leads to the following condition for instability:

$$A \geq A^{\star}_{\ell=1} = \frac{2(1+\zeta)^2}{\zeta(\zeta-1)(1-\bar{\Phi}_0)^2}, \quad \text{with} \quad \zeta > 1, \tag{S.346}$$

where $A^{\star}_{\ell=1}(\zeta)$ defines the instability boundary at which the composition fields become spontaneously unstable. By using Eq. (S.339), we can see that this also leads to an instability in the shape. For the first spherical harmonics, the instability of these modes ($\ell = 1$) corresponds to a spontaneous drift of the membrane compartment.

Herein, the drift velocity $\boldsymbol{V}_{\mathrm{cm}}$ of the centre-of-mass of the membrane compartment is defined by

$$\boldsymbol{V}_{\mathrm{cm}} = \frac{\mathrm{d}}{\mathrm{d}t}\boldsymbol{R}_{\mathrm{cm}}, \tag{S.347}$$

where $\boldsymbol{R}_{\mathrm{cm}}$ is its centre-of-mass position, and given by

$$\boldsymbol{R}_{\mathrm{cm}} = \frac{1}{V}\iiint \mathrm{d}V\,\boldsymbol{R}, \tag{S.348}$$

with the integration being taken over the enclosed volume $V$ of the compartment, and $\boldsymbol{R}$ is the position vector that describes its membrane surface $\mathcal{M}$. We can express the centre-of-mass position $\boldsymbol{R}_{\mathrm{cm}}$, as well as $\boldsymbol{V}_{\mathrm{cm}}$, in terms of a power series in terms of the perturbation parameter $\varepsilon$; that is, $\boldsymbol{R} = \sum_{\mathtt{i}=0}^{\infty} \varepsilon^{\mathtt{i}}\boldsymbol{R}^{(\mathtt{i})}$ and $\boldsymbol{V}_{\mathrm{cm}} = \sum_{\mathtt{i}=0}^{\infty} \varepsilon^{\mathtt{i}}\boldsymbol{V}^{(\mathtt{i})}_{\mathrm{cm}}$. At zeroth order in $\varepsilon$, we find that

$$\boldsymbol{R}^{(0)}_{\mathrm{cm}} = \boldsymbol{0}, \quad \text{and} \quad \boldsymbol{V}^{(0)}_{\mathrm{cm}} = \boldsymbol{0}. \tag{S.349}$$

At first order in $\varepsilon$, the centre-of-mass position is given by

$$\boldsymbol{R}^{(1)}_{\mathrm{cm}} = \frac{3R}{4\pi}\iint \mathrm{d}\varphi\,\mathrm{d}\theta\sin\theta\,\hat{\boldsymbol{r}}\,u(\theta,\varphi), \tag{S.350}$$

and therefore the centre-of-mass velocity is found to be

$$\boldsymbol{V}^{(1)}_{\mathrm{cm}} = \frac{3R}{4\pi}\iint \mathrm{d}\varphi\,\mathrm{d}\theta\sin\theta\,\hat{\boldsymbol{r}}\,\frac{\partial}{\partial t}u(\theta,\varphi). \tag{S.351}$$

By expanding, $u(\theta,\varphi)$ in spherical harmonics, only $u_{\ell=1}$ contributes to the integrals in Eqs. (S.350) and (S.351); for instance, $\iint \mathrm{d}\varphi\,\mathrm{d}\theta\sin\theta\,\hat{\boldsymbol{r}}\,Y_{1,0}(\theta,\varphi) = \sqrt{4\pi/3}\,\hat{\boldsymbol{z}}$, with $\hat{\boldsymbol{z}}$ as the unit vector pointing in the $z$-direction.

Thus, for the $m = 0$ perturbation of the $\ell = 1$ modes, centre-of-mass velocity can be written as

$$\boldsymbol{V}^{(1)}_{\mathrm{cm}} = \hat{\boldsymbol{z}}\,R\,\frac{\partial u_{1,0}}{\partial t}\sqrt{\frac{3}{4\pi}}. \tag{S.352}$$

This can be generalized to the other perturbations in the modes $m = \pm 1$ that result in velocities along $x$- and $y$-direction. Similarly, the centre-of-mass position $\boldsymbol{R}^{(1)}_{\mathrm{cm}}$ can be computed; that is, $\boldsymbol{R}^{(1)}_{\mathrm{cm}} = \hat{\boldsymbol{z}}\,R\,u_{1,0}\sqrt{\frac{3}{4\pi}}$, for $m = 0$.

As a result, Eq. (S.352), and generalizations thereof, shows that $\boldsymbol{V}^{(1)}_{\mathrm{cm}}$ depends linearly on the rate of change

| | | |
|---|---|---|
| $A$ | Dimensionless fission rate | $10-400$ |
| $\zeta$ | Ratio of fusion and fission rates | $1-4$ |
| $Q$ | Dimensionless second moment | $10-10^5$ |
| $\mathbb{M}^k_{\circlearrowleft}$ | Active rates of fusion/fission | $0.5-10\ \mathrm{s}^{-1}$ |
| $\mathbb{P}^k_{\circlearrowleft}$ | Active force first moment | $0.02-0.2$ mN/m |
| $\mathbb{Q}^k_{\circlearrowleft}$ | Active force second moment | $0.5-5$ pN |

TABLE II. Range estimation of model parameters. The rates $\mathbb{M}^k_{\circlearrowleft}$ are estimated as $1/T_a$, where the trafficking time of fission and fusion $T_a \approx 0.1$–$2$ s, as computed in Table I. This estimate, together with the surface diffusion $\gamma$ and the organelle size $R$ from Table I, allows us to compute the typical range of $A$ and $\zeta$. The estimates of $\mathbb{P}^k_{\circlearrowleft}$ and $\mathbb{Q}^k_{\circlearrowleft}$ are obtained by considering an net energy barrier for fission/fusion to be between $10-50\ k_BT$ over a coarse-graining length which is given by the vesicle size $r \approx 30$–$40$ nm. Using the estimate of viscosity $\eta$ in Table I, we determine $Q$. Note that fission/fusion energy barrier must consider both elastic deformation (membrane pore opening) and hydrodynamic lubrication (displacement of ambient fluid by transport vesicles).

of $u_1$, which in turn depends through Eq. (S.339) to the compositional fields $\mathbf{\Psi}_1$. The latter becomes linearly unstable if the condition in Eq. (S.346) is satisfied, resulting in the spontaneous motion of the compartment.

The condition for linear stability is given by $A < A^\star_{\ell=1}$ if $\zeta > 1$, and for every $A > 0$ if $\zeta \in (0,1)$. In the latter domain, we also find a region in which the solutions show damped oscillations. The boundary of this regime is found by setting the term within the square-root of Eq. (S.345) to zero (since less than zero would give rise to complex eigeinvalues, and thus oscillatory behavior). This reduces to the following condition:

$$A \geq A^o_{\ell=1} = \frac{4\zeta^2 - 8\zeta^2\,\bar{\Phi}_0 + \left(1+\zeta^2\right)^2\bar{\Phi}_0^2}{2\bar{\Phi}_0\left(1-\bar{\Phi}_0\right)^2(1-\zeta)\,\zeta^2}, \qquad \text{(S.353)}$$

with $\zeta \in (0,1)$. The phase regions are shown in Fig. S.9.

For higher surface harmonics ($\ell > 1$), we also choose to neglect the curvature couplings to the compositional fields, by setting $\Omega_k = 0$. Thus, the dynamics of $\Psi^k_\ell$ is decoupled from the shape, which means that the matrix elements $s^{(2,1)}_\ell = s^{(3,1)}_\ell = 0$. Hence, the eigenvalues associated with the compositional fields again do not depend on the first row of the stability matrix $S_\ell$, and we denote them by $\lambda^\pm_\ell$. We find that the real part of $\lambda^-_\ell < 0$ for all $\ell > 1$, and $\lambda^+_\ell > 0$ corresponds to the instability condition of all the other spherical harmonic modes:

$$\boxed{A \geq A^\star_\ell = \frac{\ell(\ell+1)(1+\zeta)^2}{\zeta(\zeta-1)(1-\bar{\Phi}_0)^2}, \quad \text{with} \quad \zeta > 1,} \qquad \text{(S.354)}$$

where $A^\star_\ell(\zeta)$ gives the $\ell$-th instability boundary at which the compositional fields become unstable. The region of stable damped oscillations can be found as before, which

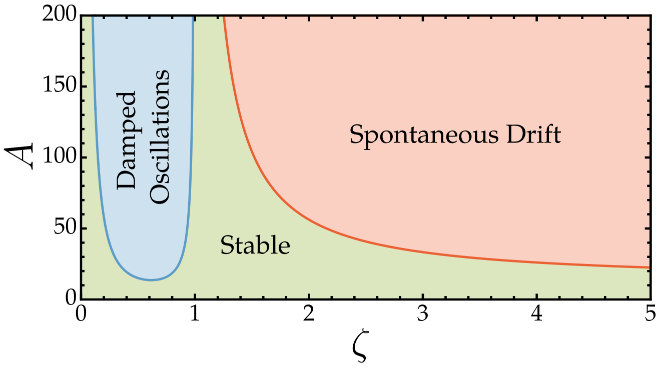


FIG. S.9. Phase diagram shows in red the linearly unstable region associated with the first spherical harmonics of the compositional fields, which leads to a spontaneous drift of the membrane compartment. The stable region is shown in green and blue, with the latter displaying damped oscillations.

leads to the following condition:

$$A \geq A^o_\ell = \frac{\ell(\ell+1)\left[4\zeta^2 - 8\zeta^2\,\bar{\Phi}_0 + \left(1+\zeta^2\right)^2\bar{\Phi}_0^2\right]}{2\bar{\Phi}_0\left(1-\bar{\Phi}_0\right)^2(1-\zeta)\,\zeta^2}, \qquad \text{(S.355)}$$

where $\zeta \in (0,1)$, and $A^o_\ell(\zeta)$ defines the phase boundary of this region. The instability in the composition readily leads to an unstable growth of the shape modes, through the respective shape equation.

Another instability can arise directly from the shape equation, where the corresponding eigenvalues $\lambda^u_\ell$ can be written as follows:

$$\lambda^u_\ell = -\frac{\ell^2(\ell+1)^2(\ell-1)(\ell+2)\left[\mathcal{K} + \frac{(Q_2-\zeta Q_1)(1-\bar{\Phi}_0)}{2\ell(\ell+1)(1+\zeta)}\right]}{(4\ell^3 + 6\ell^2 - 1)} \qquad \text{(S.356)}$$

where we assume an isotonic solution, with the osmotic pressure $\Pi_\Delta = 0$, and $\mathcal{K} = \kappa/(\gamma\eta R)$ is a rescaled bending rigidity, non-dimensionalized by the energy scale $\gamma\eta R$. Hence, a band of unstable modes can be obtained (for which $\lambda^u_\ell > 0$) when the following condition holds:

$$Q_2 < \zeta Q_1 - \frac{2\ell(\ell+1)(1+\zeta)\mathcal{K}}{1-\bar{\Phi}_0}. \qquad \text{(S.357)}$$

If we consider that $Q_1 = Q_2 = Q$ (the second moments have the same magnitude but opposite signs), the condition for instability can be further simplified as

$$\begin{cases} Q < -\frac{2\ell(\ell+1)(1+\zeta)\mathcal{K}}{(1-\zeta)(1-\bar{\Phi}_0)}, & \text{if} \quad \zeta < 1, \\ Q > \frac{2\ell(\ell+1)(1+\zeta)\mathcal{K}}{(\zeta-1)(1-\bar{\Phi}_0)}, & \text{if} \quad \zeta > 1, \end{cases} \qquad \text{(S.358)}$$

and their respective phase regions are plotted in Fig. 2(c) of the main text. If we consider the other case, with equal magnitudes and signs ($Q_1 = -Q_2 = Q$), we find that

$$Q > \frac{2\ell(\ell+1)\mathcal{K}}{1-\bar{\Phi}_0} \qquad \text{(S.359)}$$

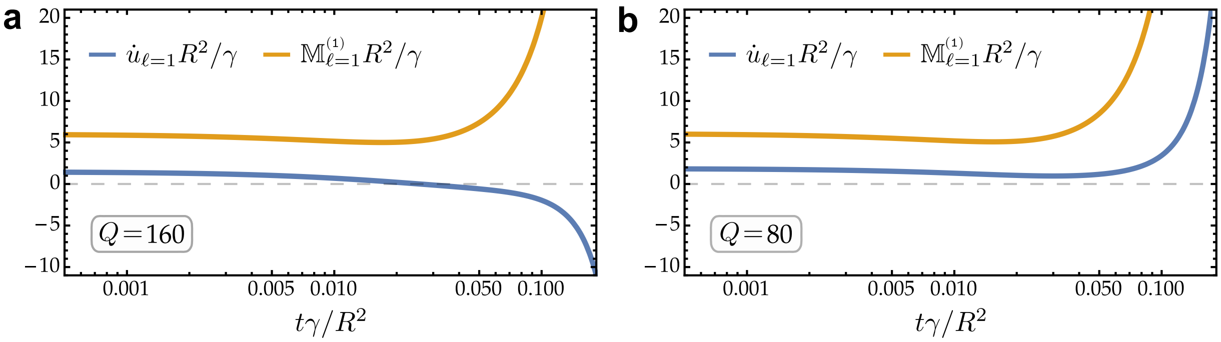


FIG. S.10. Temporal evolution of the mass-flux rate $\mathbb{M}^{(1)}_{\ell=1}$ and the rate of change of the shape, $\dot{u}_{\ell=1}$, both being measured in units of $R^2/\gamma$. Here, $\zeta = 3$, $A = 300$, $\bar{\Phi}_0 = 3/5$, $\mathcal{K} = 10$, whilst the initial conditions are chosen to be $u_{\ell=1}(t=0) = 0$, $\Psi^{k=1}_{\ell=1}(t=0) = 0$, and $\Psi^{k=2}_{\ell=1}(t=0) = 0.01$. (a) We set $Q = 160$. In this case, the motion of the compartment is anterograde; it moves in a direction opposite to the axis of the membrane flux (away from the source). (b) We choose $Q = 80$, which results in a retrograde motion; the membrane compartment moves towards the source of the vesicular mass flux.

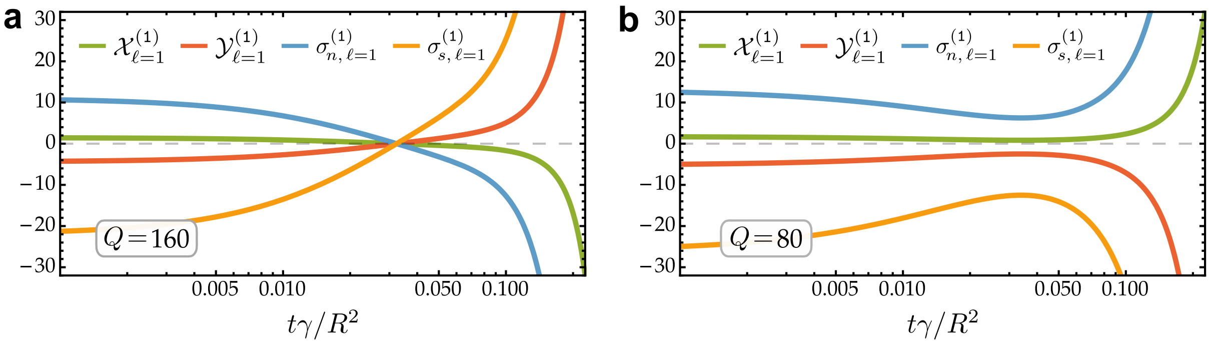


FIG. S.11. Dynamics of the surface harmonics $\mathcal{X}^{(1)}_{\ell=1}$ and $\mathcal{Y}^{(1)}_{\ell=1}$ (which determine the surrounding Stokesian flows from Lamb's solution), and the time evolution of the normal stress $\sigma^{(1)}_{n,\,\ell=1}$ and tangential stress $\sigma^{(1)}_{s,\,\ell=1}$ of the membrane interface, with all of these functions being measured in units of $R/\gamma$. The initial conditions are set to be $u_{\ell=1}(t=0) = 0$, $\Psi^{k=1}_{\ell=1}(t=0) = 0$, and $\Psi^{k=2}_{\ell=1}(t=0) = 0.01$. Here, $\zeta = 3$, $A = 300$, $\bar{\Phi}_0 = 3/5$, $\mathcal{K} = 10$, while $Q = 160$ and $Q = 80$ in subfigure (a) and (b), respectively.

corresponds to the unstable region, which is independent of the parameter $\zeta$. However, in what follows we will only consider the case of $Q_1 = Q_2 = Q$.

#### H. Solutions to membrane shape and composition

The governing equations at first order in the perturbation expansion, from Eq. (S.338), can be solved exactly by starting with some initial conditions in the shape $u_\ell(t)$ and the compositional fields $\Psi^k_\ell(t)$ at time $t = 0$. This allows to compute the transient evolution of each of these fields, which we know that in the unstable regime they asymptotically diverge, growing exponentially with a rate that is given by the respective eigenvalue $\lambda_\ell$. Once the solutions to $u_\ell(t)$ and $\Psi^k_\ell(t)$ are determined we can compute all of the other quantities of interest; for instance, the corresponding surface tension from Eq. (S.337).

We begin by solving the simultaneous set of linear differential equations of the $\ell = 1$ mode, which are shown in Eqs. (S.339) and (S.340). By employing the initial condition that $u_{\ell=1}(t = 0) = 0$ (that is, there is no distortion of the membrane at $t = 0$), we perturb one (or both) of the compositional fields at $t = 0$. The quantities of interest are the local rate of change of the shape, $\dot{u}_{\ell=1}$, and the net membrane flux-rate $\mathbb{M}^{(1)}_{\ell=1}$, which are shown in Fig. S.10 for some fixed values of the parameters $A$, $\zeta$, $\mathcal{K}$, $Q$, and the area fraction $\bar{\Phi}_0$ of the membrane background component. Here, $A$ and $\zeta$ are chosen such that the system is compositionally unstable, as given by Eq. (S.346). Note that this instability condition is independent of the parameter $Q$. This could also be seen in the overall membrane flux-rate $\mathbb{M}^{(1)}_{\ell=1}$, which grows exponentially, as plotted in Fig. S.10, being entirely insensitive to value of the active second moment. On the other hand, the shape depends on $Q$, and the sign of $\dot{u}_{\ell=1}$ at large times can switch depending on its value.

Since $\dot{u}_{\ell=1}$ gives us the velocity $\boldsymbol{V}^{(1)}_{\rm cm}$ of the centre of mass of the membrane compartment, cf. Eq. (S.352), the temporal evolution of $\dot{u}_{\ell=1}$ and in particular its asymptotic behavior tells us the direction in which the compartment moves with respect to the axis of the net membrane flux. If the compartment drifts towards the source of the mass flux, the motion is called *retrograde*; it moves in the

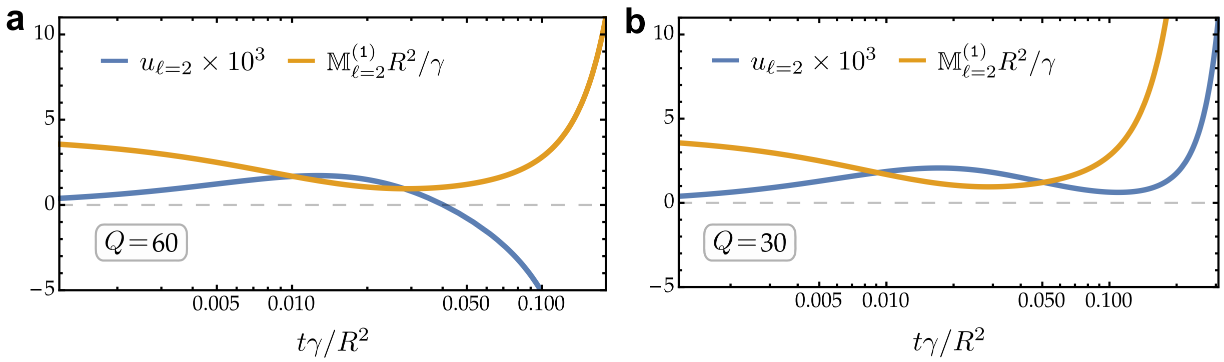


FIG. S.12. The time evolution of the mass-flux rate $\mathbb{M}^{(1)}_{\ell=2}$, measured in units of $R^2/\gamma$, and the temporal dynamics of the distortion shape field $u_{\ell=2}$ (scaled by $10^3$ for clarity). Here, $\zeta=3$, $A=300$, $\bar{\Phi}_0=3/5$, $\mathcal{K}=10$, whereas the initial conditions are chosen to be $u_{\ell=2}(t=0)=0$, $\Psi^{k=1}_{\ell=2}(t=0)=0.01$, and $\Psi^{k=2}_{\ell=2}(t=0)=0.01$. (a) We choose $Q=60$, which leads to an oblate deformation of the compartment. (b) We set $Q=30$, which results in prolate deformation at large time.

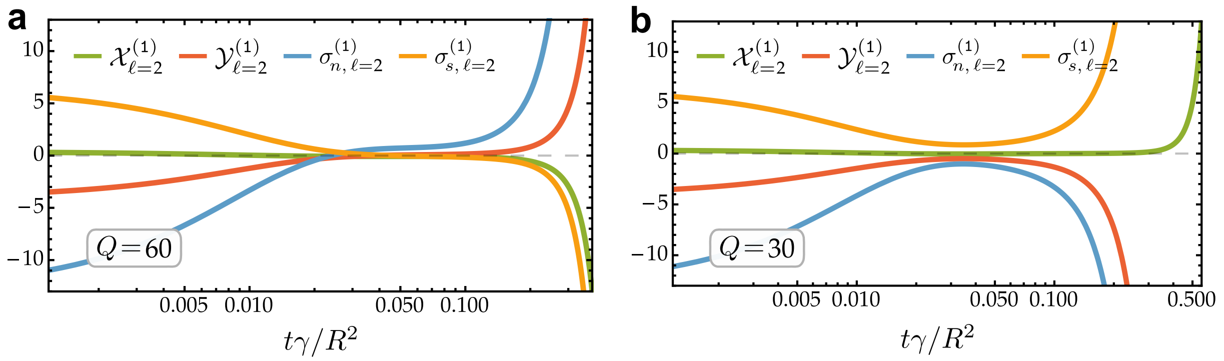


FIG. S.13. The dynamics of the spherical harmonics $\mathcal{X}^{(1)}_{\ell=2}$ and $\mathcal{Y}^{(1)}_{\ell=2}$, and the time evolution of the membrane stresses, $\sigma^{(1)}_{n,\,\ell=2}$ and $\sigma^{(1)}_{s,\,\ell=2}$, which are associated with the normal and tangential components, respectively. All of these functions are measured in units of $R/\gamma$. The initial conditions are set to be $u_{\ell=1}(t=0)=0$, $\Psi^{k=1}_{\ell=2}(t=0)=0.01$, and $\Psi^{k=2}_{\ell=2}(t=0)=0.01$. Here, we choose that $\zeta=3$, $A=300$, $\bar{\Phi}_0=3/5$, $\mathcal{K}=10$, while $Q=60$ and $Q=30$ in subfigure (a) and (b), respectively.

direction in which the mass is being added, similar to the phenomenon of treadmilling. On the other hand, if the compartment moves away from the source of the membrane flux, we call this motion to be *anterograde*, moving in a direction opposite to treadmilling.

The solution to $u_{\ell=1}(t)$ and $\Psi^k_{\ell=1}(t)$ also allows us to compute the temporal evolution of the surface spherical harmonics $\mathcal{X}^{(1)}_{\ell=1}$ and $\mathcal{Y}^{(1)}_{\ell=1}$, which fully determine the surrounding Stokesian flows via Lamb's solution. Namely, for the mode $\ell=1$, we have that

$$\mathcal{X}^{(1)}_{\ell=1}=R\,\frac{\partial u_1}{\partial t}-\sum_k \Psi^k_1\mathbb{V}^k_{\circlearrowleft},\quad \text{and} \tag{S.360}$$

$$\mathcal{Y}^{(1)}_{\ell=1}=\sum_k \Psi^k_1\left[2\mathbb{V}^k_{\circlearrowleft}-R\mathbb{M}^k_{\circlearrowleft}+\frac{\mathbb{Q}^k_{\circlearrowleft}}{\eta R}\right], \tag{S.361}$$

which are plotted in Fig. S.11, by neglecting the active volume fluxes as before. Note that the sign of both $\mathcal{X}^{(1)}_{\ell=1}$ and $\mathcal{Y}^{(1)}_{\ell=1}$ at large time could switch as we increase the value of the parameter $Q$ (as a reminder, the choice $Q=Q_1=Q_2$ is used throughout; interestingly, the other case for $Q$ shows qualitatively the same results).

Moreover, the membrane stresses could be calculated once the solutions to the shape and compositional fields are determined; namely, the stress jumps at the membrane interface which are balanced by the fluid stresses across the membrane. In the tangential direction, this is given by the surface component of the body force at first order; that is, we define that $\sigma^{(1)}_{s,\,\ell,m}=\boldsymbol{f}^{(1)}\cdot\boldsymbol{\Theta}_{\ell,m}$, where the vector $\boldsymbol{\Theta}_{l,m}=r\boldsymbol{\nabla}Y_{l,m}(\theta,\varphi)$. By using Eq. (S.272), and summing over each of the $m$-modes, this leads to

$$\boxed{\sigma^{(1)}_{s,\,\ell}=-\frac{\eta(2\ell+1)}{\ell(\ell+1)R}\left[\mathcal{X}^{(1)}_{\ell}+2\mathcal{Y}^{(1)}_{\ell}\right].} \tag{S.362}$$

Along the normal to the surface, the stress jump across the interface is given by $f^{(1)}=\boldsymbol{n}\cdot\boldsymbol{f}^{(1)}$, cf. Eq. (S.261). For clarity, we rewrite its projection onto each harmonics by $\sigma^{(1)}_{n,\ell,m} = \left[f^{(1)}\right]_{\ell,m}$. Hence, using Eq. (S.262), we have

$$\boxed{\sigma^{(1)}_{n,\ell} = \frac{\eta(2\ell+1)\left\{3\mathcal{Y}^{(1)}_{\ell} - [2\ell(\ell+1)-3]\mathcal{X}^{(1)}_{\ell}\right\}}{\ell(\ell+1)R}.} \quad \text{(S.363)}$$

Therefore, the membrane stresses, $\sigma^{(1)}_{s,\ell=1}$ and $\sigma^{(1)}_{n,\ell=1}$, can be readily obtained by computing the surface spherical harmonics $\mathcal{X}^{(1)}_{\ell=1}$ and $\mathcal{Y}^{(1)}_{\ell=1}$, as shown in Fig. S.11.

The phase regions corresponding to anterograde and retrograde motion (in the asymptotic limit of large time) are illustrated in Fig. S.14(a), where we fix the value of $Q$, and we vary $\zeta$ and $A$ provided that they respect the instability condition in Eq. (S.346). The dashed magenta curve in Fig. S.14(a) corresponds to the phase boundary between the anterograde and retrograde regimes, which is given by $Q^{\star}_{\ell=1}(A,\zeta)$. For $Q > Q^{\star}_{\ell=1}$, the motion of the compartment is retrograde, while for $Q < Q^{\star}_{\ell=1}$ it is anterograde. The form of $Q^{\star}_{\ell=1}(A,\zeta)$ can be written as

$$Q^{\star}_{\ell=1} = \frac{A(\zeta+1)\left\{\bar{\Phi}_0\zeta^2 + 2\zeta + \bar{\Phi}_0 - \sqrt{\bar{\Phi}_0\left[2A(1-\bar{\Phi}_0)^2\zeta^3 - 2\zeta^2\left(A(1-\bar{\Phi}_0)^2 - \bar{\Phi}_0 + 4\right) + \bar{\Phi}_0\zeta^4 + \bar{\Phi}_0\right] + 4\zeta^2}\,\right\}}{\bar{\Phi}_0\left[2\zeta(\zeta+2) + 2 - A\zeta(\zeta-1)(1-\bar{\Phi}_0)^2\right]}. \quad \text{(S.364)}$$

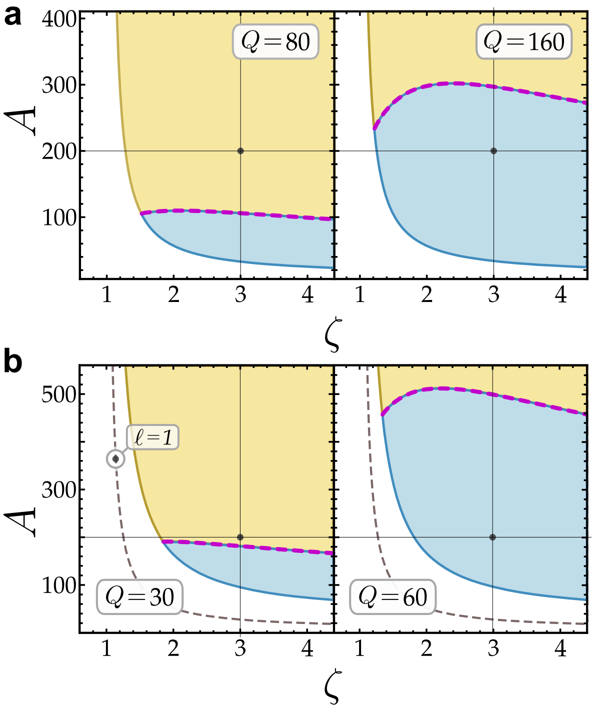


FIG. S.14. (a) Diagrams of the phase regions corresponding to retrograde (in yellow, above dashed) and anterograde (in blue, below dashed) motion of the membrane compartment, for two different values of $Q$. Black points indicate the parameters used to render Fig. S.10 and S.11. (b) Plots of the phase regions in which the the shape distortion of compartment is a prolate (in yellow) and an oblate (in blue). Black points indicate the parameters used in Fig. S.12 and S.13.

The dependence on $Q^{\star}_{\ell=1}$ on $\zeta$ and $A$ in the unstable region is plotted in Fig. S.15(a). We find that the limit

$$\lim_{\zeta\to\infty} Q^{\star}_{\ell=1}(A,\zeta) = A/\bar{\Phi}_0, \quad \text{(S.365)}$$

is strictly greater than zero for $A > 0$, which means that the antegrograde motion can only exist for $Q > 0$. As

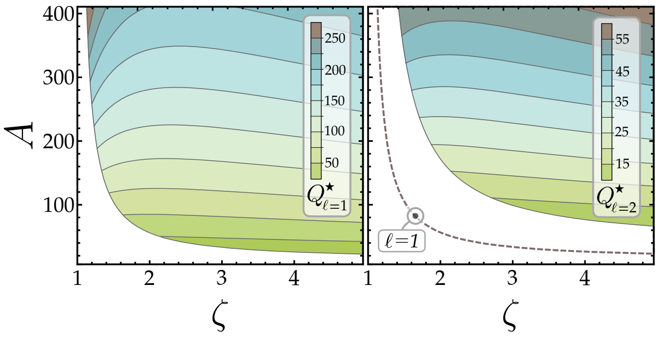


FIG. S.15. Contour plot of the phase boundaries between the anterograde to retrograde regions (left), with $Q = Q^{\star}_{\ell=1}(\zeta, A)$. (b) Contour plot of the phase boundaries between regions of prolate and oblate deformations (right), with $Q = Q^{\star}_{\ell=2}(\zeta, A)$.

we will see in the later, this asymptotic behaviour is only true for the linear theory.

Similarly, we can solve the set of differential equations in Eq. (S.338) associated with the mode $\ell = 2$, by using initial conditions at time $t = 0$ for the compositional fields $\Psi^{k}_{\ell=2}(t)$, and assuming an initial undeformed shape with $u_{\ell=2}(t=0) = 0$. The time evolution of shape distortion $u_2$ is plotted in Fig. S.12 for two different values of $Q$, which show distinct asymptotic divergences. If the sign of $u_2(t)$ is different to the sign of the mass flux $\mathbb{M}^{(1)}_{\ell=2}$ as time $t \to \infty$, then the membrane compartment suffers an oblate deformation; otherwise, if $\text{sgn}(u_2) = \text{sgn}\left[\mathbb{M}^{(1)}_{\ell=2}\right]$, then it results in a prolate deformation. Note that this nomenclature is based on shape distortions with mode number $m = 0$, which revolve about the $z$-axis, as illustrated in Fig. 3 of the main text, and Fig S.16.

The solution to $u_{\ell=2}(t)$ and $\Psi^{k}_{\ell=2}(t)$ also allows us to compute the spherical harmonics $\mathcal{X}^{(1)}_{\ell=2}$ and $\mathcal{Y}^{(1)}_{\ell=2}$, which in turns allows to find the local membrane stresses $\sigma^{(1)}_{s,\ell=2}$ and $\sigma^{(1)}_{n,\ell=2}$, as shown in Fig. S.13. The phase regions associated with prolate and oblate deformations are shown

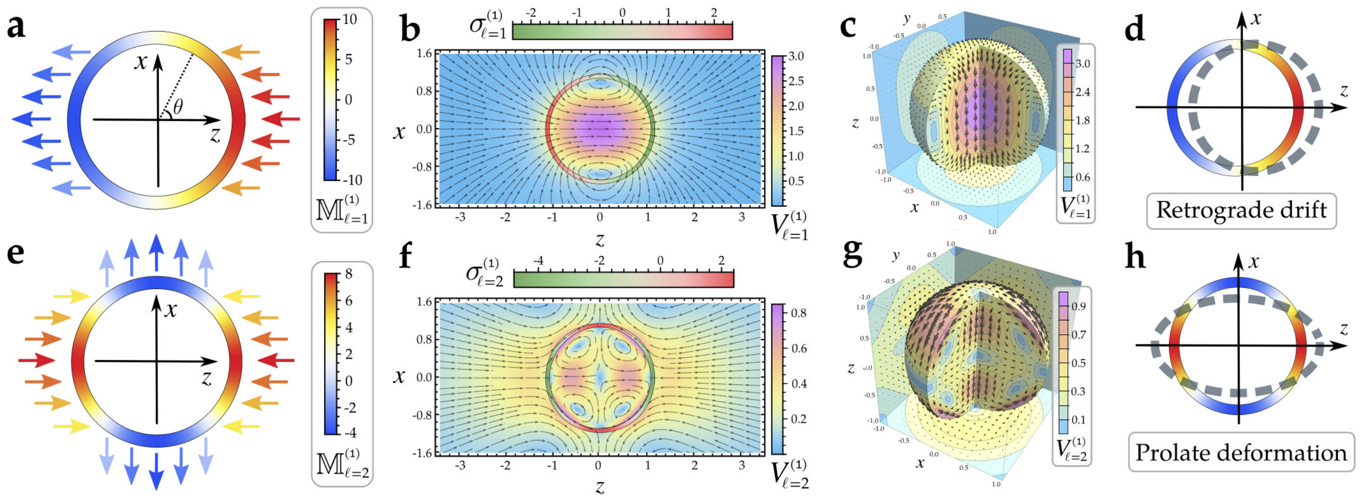


FIG. S.16. Here, $\zeta = 3$, $A = 300$, $\bar{\Phi}_0 = 3/5$, $\mathcal{K} = 10$, and $Q = 30$, with every plot shown at a snapshot time $t = 0.1\,\gamma/R^2$ (a) Total membrane mass flux-rate $\mathbb{M}^{(1)}_{\ell=1}(\theta, \varphi = 0)$, being measured in units of $\gamma/R^2$. (b) Cross-sectional view of the surrounding fluid flows for the mode $\ell = 1$ and $m = 0$, where the color indicates the magnitude of the velocity. The normal stress $\sigma^{(1)}_{\ell=1}$ at the membrane interface is also plotted as a color density. (c) Three-dimensional view of the fluid flows for mode $\ell = 1$ and $m = 0$; same as flows shown in the previous subfigure. (d) The vesicular membrane flux induces stresses and flows within the membrane which in turn lead to shape changes; in this case, the membrane compartment has a retrograde motion, drifting in the direction of the source (along the positive $z$-axis). (e) Overall membrane mass flux-rate $\mathbb{M}^{(1)}_{\ell=2}(\theta, \varphi = 0)$, measured again in units of $\gamma/R^2$. (f) The leading-order outer and inner fluid flows for the mode $\ell = 2$ and $m = 0$; same cross-sectional cut as in subfigure (b). The normal membrane stress $\sigma^{(1)}_{\ell=2}$ induced by the vesicular flux is depicted as a density plot at the interface. (g) Three-dimensional view of the fluid flows. (h) In this case, the membrane compositional instability leads to a shape change that deforms the comparment into a prolate, with the long axis along the $z$-axis.

in Fig. S.14(b) at fixed $Q$ and varying values of $\zeta$ and $A$. The dashed magenta curve is the boundary between these two regimes, and described by $Q^{\star}_{\ell=2}(A, \zeta)$, which is shown in Fig. S.15(b). Similarly, we can show that oblate deformation can only exist in the linear theory if $Q > 0$, as $Q^{\star}_{\ell=2}(A, \zeta)$ is strictly greater than zero.

#### I. Solutions to the surrounding fluid flows

By computing the surface harmonics $\mathcal{X}^{(1)}_{\ell}$ and $\mathcal{Y}^{(1)}_{\ell}$, in terms of the shape distortion $u_\ell(t)$ and the compositional fields $\Psi^k_\ell(t)$, we could readily find the surrounding fluid flows by using Lamb's solution in Eq. (S.179). By substituting the solid hamonics $p^{(1)}_{\ell}$ and $\Upsilon^{(1)}_{\ell}$ from Eqs. (S.247) and (S.248), respectively, we find the following expression for the velocity of the inner fluid flow associated with each spherical harmonic mode $\ell > 0$ and $m$ (with $|m| \leq \ell$):

$$\left[\boldsymbol{V}^{(1)}_{-}\right]_{\ell,m} = \frac{\hat{\boldsymbol{r}} Y_{\ell,m} r^{\ell-1}}{2R^{\ell-1}}\left[\mathcal{X}_{\ell,m}\left(1+\ell-(\ell-1)\frac{r^2}{R^2}\right) - \mathcal{Y}_{\ell,m}\left(1-\frac{r^2}{R^2}\right)\right] - \frac{\boldsymbol{\Theta}_{\ell,m}\, r^{\ell-1}}{2\ell R^{\ell-1}}\left[\mathcal{Y}_{\ell,m}\left(1-\frac{(\ell+3)\,r^2}{(\ell+1)\,R^2}\right) - \mathcal{X}_{\ell,m}\left(1+\ell-\frac{(\ell-1)(\ell+3)\,r^2}{(\ell+1)\,R^2}\right)\right], \tag{S.366}$$

where the vector spherical harmonics $\boldsymbol{\Theta}_{l,m}$ are defined by $\boldsymbol{\Theta}_{l,m} = r\boldsymbol{\nabla} Y_{l,m}(\theta, \varphi)$. Similarly, by substituting $p^{(1)}_{-(\ell+1)}$ and $\Upsilon^{(1)}_{-(\ell+1)}$ from the equations (S.249) and (S.250), respectively, we obtain the following expression:

$$\left[\boldsymbol{V}^{(1)}_{+}\right]_{\ell,m} = \frac{\hat{\boldsymbol{r}} Y_{\ell,m} R^{\ell+2}}{2\,r^{\ell+2}}\left[\mathcal{X}_{\ell,m}\left(-\ell+(\ell+2)\frac{r^2}{R^2}\right) - \mathcal{Y}_{\ell,m}\left(1-\frac{r^2}{R^2}\right)\right] + \frac{\boldsymbol{\Theta}_{\ell,m}\, R^{\ell+2}}{2(\ell+1)\,r^{\ell+2}}\left[\mathcal{Y}_{\ell,m}\left(1-\frac{(\ell-2)\,r^2}{\ell\,R^2}\right) + \mathcal{X}_{\ell,m}\left(\ell-\frac{(\ell+2)(\ell-2)\,r^2}{\ell\,R^2}\right)\right]. \tag{S.367}$$

The fluid velocity $\boldsymbol{v}^{(1)}_{\mathrm{m}}$ at the membrane surface ($r = R$) for each surface harmonic mode $\ell$ and $m$ is found to be

$$\boxed{\left[\boldsymbol{v}^{(1)}_{\mathrm{m}}\right]_{\ell,m} = \mathcal{X}_{\ell,m}\,\hat{\boldsymbol{r}}\, Y_{\ell,m} + \frac{2\mathcal{X}_{\ell,m}+\mathcal{Y}_{\ell,m}}{\ell(\ell+1)}\,\boldsymbol{\Theta}_{\ell,m}.} \tag{S.368}$$

The outer and inner fluid flows are plotted in Fig. S.16(b) and (c) for $\ell = 1$ and $m = 0$, and in Fig. S.16(f) and (g) for $\ell = 2$ and $m = 0$. Here, $\zeta$, $A$, and $Q$ are chosen such that the system is linearly unstable, resulting in a retrograde motion and a prolate distortion of the membrane compartment for $\ell = 1$ and $\ell = 2$, respectively. The case when the motion is anterograde and the shape deforms into an oblate is shown in Fig. 3 of the main text.

From Eq. (S.367) we can find the far field flow of the outside fluid, which is given by

$$\boldsymbol{V}_{+}^{(1)} = \sum_{|m|\leq 1} \frac{3\mathcal{X}_{1,m} + \mathcal{Y}_{1,m}}{4(r/R)} \left(2\hat{\boldsymbol{r}}\, Y_{1,m} + \boldsymbol{\Theta}_{1,m}\right) + \sum_{|m|\leq 2} \frac{4\mathcal{X}_{2,m} + \mathcal{Y}_{2,m}}{2(r/R)^2}\, \hat{\boldsymbol{r}}\, Y_{2,m} \; + \mathcal{O}[r^{-3}]. \quad \text{(S.369)}$$

Using the definitions of $\mathcal{X}_1$ and $\mathcal{Y}_1$ in Eq. (S.360) and (S.361), respectively, together with the membrane shape equation of $u_1$ in (S.339), we can show that the term associated with $1/r$ is identically zero, where the factor $3\mathcal{X}_{1,m} + \mathcal{Y}_{1,m} = 0$ for all $m \leq 1$. Thus, the far-field flow of the outside fluid is found to be

$$\boldsymbol{V}_{+}^{(1)} = \sum_{|m|\leq 2} \frac{4\mathcal{X}_{2,m} + \mathcal{Y}_{2,m}}{2(r/R)^2}\, \hat{\boldsymbol{r}}\, Y_{2,m} \; + \mathcal{O}[r^{-3}], \quad \text{(S.370)}$$

where the factor

$$4\mathcal{X}_2 + \mathcal{Y}_2 = 4R\,\partial_t u_2 - 2\mathbb{V}_2 - R\,\mathbb{M}_2 + \frac{2\mathbb{Q}_2}{\eta R}. \quad \text{(S.371)}$$

The drag force on the membrane compartment can be computed by integrating the fluid stress vectors at the interface over the whole surface. By integrating first along the normal component of the stress vectors, namely

$$\boldsymbol{F}_n = \iint \mathrm{d}\varphi\, \mathrm{d}\theta\, R^2 \sin\theta \sum_{\ell,m} \sigma_{n,\,\ell,m}^{(1)}\, \hat{\boldsymbol{r}}\, Y_{\ell,m}(\theta,\varphi), \quad \text{(S.372)}$$

with $\sigma_{n,\,\ell,m}^{(1)}$ as defined in Eq. (S.363), we find that only the mode $\ell = 1$ contributes to the integral:

$$\boldsymbol{F}_n = 10\sqrt{\frac{\pi}{3}}\,\hat{\boldsymbol{z}}\, \left[\mathbb{Q} + 2\eta R\mathbb{V} - \eta R^2\mathbb{M}\right]_{\ell=1,m=0} + \dots \quad \text{(S.373)}$$

where the dots imply a summation over the other two modes $m = \pm 1$ associated with $\hat{\boldsymbol{x}}$ and $\hat{\boldsymbol{y}}$ vectors, having the same prefactor as in Eq. (S.373). By integrating the surface component of the stress vectors,

$$\boldsymbol{F}_s = \iint \mathrm{d}\varphi\, \mathrm{d}\theta\, R^2 \sin\theta \sum_{\ell,m} \sigma_{s,\,\ell,m}^{(1)}\, \boldsymbol{\Theta}_{\ell,m}(\theta,\varphi), \quad \text{(S.374)}$$

we obtain that

$$\boldsymbol{F}_s = 10\sqrt{\frac{\pi}{3}}\,\hat{\boldsymbol{z}}\, \left[\eta R^2\mathbb{M} - \mathbb{Q} - 2\eta R\mathbb{V}\right]_{\ell=1,m=0} + \dots \quad \text{(S.375)}$$

where the dots denote all of the other terms corresponding to $m = \pm 1$, as before. This gives us that $\boldsymbol{F}_n + \boldsymbol{F}_s = \boldsymbol{0}$, which implies that the membrane compartment is force-free, as expected. In other words, under the action of a vesicular membrane flux, the compartment is a force-free swimmer, moving either retrograde or anterograde with respect to the direction of the mass flux.

### III. NONLINEAR PERTURBATIVE ANALYSIS

The linear perturbation analysis indicates that the homogeneous steady-state of the membrane compartment becomes unstable, where the compositional instability drives a shape distortion (for all modes $\ell > 1$), as well as a spontaneous drift instability (for $\ell = 1$). Note that $\ell = 0$ corresponds to a expansion or dilation of the membrane compartment, as a result of the addition or removal of membrane material via the vesicular flux.

#### A. Weakly nonlinear compositional dynamics

The compositional instability brings the system in a nonlinear regime, where the higher-order terms in the perturbation parameter $\varepsilon$ are no longer negligible. Since the governing equations of the compositional fields $\Phi_k$ drive the linear instability, we expand Eq. (S.78) up to second order in $\varepsilon$, with the mass fractions being given by $\Phi_k = \bar{\Phi}_k + \varepsilon\Psi_k$, and the shape distortion of the membrane compartment $\boldsymbol{R} = R(1+\varepsilon u)\hat{\boldsymbol{r}}$. By retaining all of the terms up to quadratic order in the perturbation parameter $\varepsilon$, this leads to the following expression:

$$\begin{aligned}\dot{\Psi}_k = &-\left(\bar{\Phi}_k + \Psi_k\right)\mathbb{M}^{(1)} + 2\Omega_k\left[\Delta^{(0)}H^{(1)} + \Delta^{(1)}H^{(1)}\right]\\ &+\gamma_k\,\Delta^{(0)}\left[\frac{\Psi_0}{\bar{\Phi}_0}+\frac{\Psi_k}{\bar{\Phi}_k}\right] + \gamma_k\,\Delta^{(1)}\left[\frac{\Psi_0}{\bar{\Phi}_0}+\frac{\Psi_k}{\bar{\Phi}_k}\right]\\ &+\gamma_k\left[\frac{\Psi_{0\,;\alpha}\Psi_{0\,;\beta}}{\left(\bar{\Phi}_0\right)^2} - \frac{\Psi_{k\,;\alpha}\Psi_{k\,;\beta}}{\left(\bar{\Phi}_k\right)^2}\right]\left[g^{\alpha\beta}\right]^{(0)}, \qquad \text{(S.376)}\end{aligned}$$

where $\Delta^{(0)}$ is the Laplacian at zeroth order, as defined in Eq. (S.282), and $\Delta^{(1)} = -2u(\theta,\varphi)\,\Delta^{(0)}$. Here, $\left[g^{\alpha\beta}\right]^{(0)}$ is the inverse metric tensor at zeroth order in $\varepsilon$, that is,

$$\left[g^{\alpha\beta}\right]^{(0)} = \begin{bmatrix} 1/R^2 & 0 \\ 0 & 1/(R^2\sin\theta)\end{bmatrix}. \qquad \text{(S.377)}$$

Moreover, the last term in Eq. (S.376) could be simplified by noting that

$$\frac{\Psi_{0\,;\alpha}\Psi_{0\,;\beta}}{\left(\bar{\Phi}_0\right)^2} = \frac{(r\boldsymbol{\nabla}\Psi_0)^2}{R^2\left(\bar{\Phi}_0\right)^2}, \quad \text{and} \quad \frac{\Psi_{k\,;\alpha}\Psi_{k\,;\beta}}{\left(\bar{\Phi}_k\right)^2} = \frac{(r\boldsymbol{\nabla}\Psi_k)^2}{R^2\left(\bar{\Phi}_k\right)^2},$$

which can be written in terms of a total derivative:

$$\frac{\Psi_{0\,;\alpha}\Psi_{0\,;\beta}}{\left(\bar{\Phi}_0\right)^2} = \boldsymbol{\nabla}\cdot\left[\frac{r^2\Psi_0\boldsymbol{\nabla}\Psi_0}{R^2\left(\bar{\Phi}_0\right)^2}\right] - \frac{\Psi_0\,\Delta^{(0)}\Psi_0}{\left(\bar{\Phi}_0\right)^2}, \qquad \text{(S.378)}$$

and similarly we have that

$$\frac{\Psi_{k\,;\alpha}\Psi_{k\,;\beta}}{\left(\bar{\Phi}_k\right)^2} = \boldsymbol{\nabla}\cdot\left[\frac{r^2\Psi_k\boldsymbol{\nabla}\Psi_k}{R^2\left(\bar{\Phi}_k\right)^2}\right] - \frac{\Psi_k\,\Delta^{(0)}\Psi_k}{\left(\bar{\Phi}_k\right)^2}. \qquad \text{(S.379)}$$

Note that the mass fraction $\Psi_0 = \Psi_1 + \Psi_2$, due to the mass conservation, as shown in Eq. (S.277).

The material derivative in Eq. (S.376) is given by

$$\dot{\Psi}_k = \frac{\partial\Psi_k}{\partial t} + \left[v^\alpha\right]^{(1)}\partial_\alpha\Psi_k, \qquad \text{(S.380)}$$

where the convective term can be expressed as follows:

$$\left[v^\alpha\right]^{(1)}\partial_\alpha\Psi_k = \frac{r}{R}\,\boldsymbol{v}^{(1)}\cdot\boldsymbol{\nabla}\Psi_k. \qquad \text{(S.381)}$$

This allows us to determine the dynamics of the excess mass fractions $\Psi_k$ in the weakly nonlinear regime:

$$\begin{aligned}\frac{\partial\Psi_k}{\partial t} = &-\left(\bar{\Phi}_k + \Psi_k\right)\mathbb{M}^{(1)} + 2\Omega_k\,(1-2u)\,\Delta^{(0)}H^{(1)}\\ &+\gamma_k\left\{(1-2u)\,\Delta^{(0)}\left[\frac{\Psi_0}{\bar{\Phi}_0}+\frac{\Psi_k}{\bar{\Phi}_k}\right] - \frac{\Psi_0\,\Delta^{(0)}\Psi_0}{\left(\bar{\Phi}_0\right)^2} + \frac{\Psi_k\,\Delta^{(0)}\Psi_k}{\left(\bar{\Phi}_k\right)^2}\right\}\\ &+\gamma_k\,\Delta^{(0)}\left[\frac{(\Psi_0)^2}{2\left(\bar{\Phi}_0\right)^2} - \frac{(\Psi_k)^2}{2\left(\bar{\Phi}_k\right)^2}\right] - \frac{\boldsymbol{v}^{(1)}\cdot r\boldsymbol{\nabla}\Psi_k}{R}. \qquad \text{(S.382)}\end{aligned}$$

#### B. Galerkin projection and truncation method

The compositional fields $\Psi_k(\theta,\varphi)$ can be decomposed into the spherical harmonics; namely, we write

$$\Psi_k(\theta,\varphi) = \sum_{\ell=0}^{\infty}\sum_{m=-\ell}^{\ell}\Psi^k_{\ell,m}\,Y_{\ell,m}(\theta,\varphi). \qquad \text{(S.383)}$$

Therefore, the dynamical equation associated with each spherical mode is found via the orthogonality relation:

$$\frac{\partial\Psi^k_{\ell,m}}{\partial t} = \iint \mathrm{d}\varphi\,\mathrm{d}\theta\sin\theta\;Y^*_{\ell,m}(\theta,\varphi)\,\frac{\partial\Psi_k}{\partial t}, \qquad \text{(S.384)}$$

with $Y^*_{\ell,m}(\theta,\varphi) = (-1)^m\,Y_{\ell,-m}(\theta,\varphi)$. By directly substituting Eq. (S.382) into the above expression, we see that the dynamics of a spherical harmonic mode is coupled to all of the other modes, in contrast to the linear perturbation for which every mode is independent.

We define the projection operation of some angular function $h(\theta,\varphi)$ onto the spherical harmonics by

$$\left\lfloor h(\theta,\varphi)\right\rfloor_{\ell,m} = \iint \mathrm{d}\varphi\,\mathrm{d}\theta\sin\theta\;Y^*_{\ell,m}(\theta,\varphi)\,h(\theta,\varphi). \qquad \text{(S.385)}$$

First nonlinear term in Eq. (S.382) is $\Psi_k\,\mathbb{M}^{(1)}$, which can be decomposed into the spherical basis as follows:

$$\begin{aligned}\left\lfloor\Psi_k\,\mathbb{M}^{(1)}\right\rfloor_{\ell,m} = &\sum_{k'}\sum_{\ell',m'}\sum_{\ell'',m''}\mathbb{M}^{k'}_{\circlearrowleft}\,\Psi^k_{\ell',m'}\Psi^{k'}_{\ell'',m''}\times\\ &\times(-1)^m\,\mathcal{W}\left(\ell',m'|\ell'',m''|\ell,-m\right), \qquad \text{(S.386)}\end{aligned}$$

where the coefficient $\mathcal{W}\left(\ell_1,m_1|\ell_2,m_2|\ell_3 m_3\right)$ is defined by the following integral

$$\mathcal{W} = \iint \mathrm{d}\varphi\,\mathrm{d}\theta\sin\theta\;Y_{\ell_1,m_1}(\theta,\varphi)\,Y_{\ell_2,m_2}(\theta,\varphi)\,Y_{\ell_3,m_3}(\theta,\varphi),$$

which is resolved in terms of the Wigner $3j$-symbol [42]:

$$\mathcal{W} = \sqrt{\tfrac{(2\ell_1+1)(2\ell_2+1)(2\ell_3+1)}{4\pi}} \begin{pmatrix} \ell_1 & \ell_2 & \ell_3 \\ 0 & 0 & 0 \end{pmatrix} \begin{pmatrix} \ell_1 & \ell_2 & \ell_3 \\ m_1 & m_2 & m_3 \end{pmatrix},$$

where the $3j$-symbols $\begin{pmatrix} \ell_1 & \ell_2 & \ell_3 \\ m_1 & m_2 & m_3 \end{pmatrix}$ can be computed using the *Racah formula* [44]. In order for the $3j$-symbols to be non-zero, we require both $m_1 + m_2 + m_3 = 0$ and the triangle inequality $|\ell_1 - \ell_2| \leq \ell_3 \leq \ell_1 + \ell_2$ to hold; otherwise, if any of these conditions are not satisfied, $3j$-symbols are identically zero. Hence, the double infinite sum in Eq. (S.385) is effectively reduced to a single sum, which in practice needs to be truncated to a finite mode. Herein, we choose to truncate this summation to the second mode; in order words, only $\ell = 0$, 1, and 2 are used, neglecting the contribution for the higher-order modes. This projection procedure onto some local orthogonal basis (in this case, we use the surface spherical harmonics) is formally known as the Galerkin projection method, which leads to a set of coupled nonlinear differential equations for the mode amplitudes [45].

The projection of the other nonlinear terms can be similarly determined. Second nonlinear term in Eq. (S.382), that is, $2u\,\Delta^{(0)}H^{(1)}$, can be decomposed as

$$\left\lfloor u\,\Delta^{(0)}H^{(1)}\right\rceil_{\ell,m} = \sum_{\ell',m'}\sum_{\ell'',m''} \frac{\ell'(\ell'+2)(\ell'-1)(\ell'+1)}{2R^3} \times \\ \times (-1)^m\, u_{\ell',m'} u_{\ell'',m''}\, \mathcal{W}\,(\ell', m'|\ell'', m''|\ell, -m), \quad \text{(S.387)}$$

where Eq. (S.292) is used to expand the Laplacian term in terms of spherical harmonics. Moreover, we have

$$\left\lfloor u\,\Delta^{(0)}\Psi_k\right\rceil_{\ell,m} = \sum_{\ell',m'}\sum_{\ell'',m''} \frac{\ell'(\ell'+1)}{R^2}\,\Psi^k_{\ell',m'}\, u_{\ell'',m''} \times \\ \times (-1)^{m+1}\, \mathcal{W}\,(\ell', m'|\ell'', m''|\ell, -m), \quad \text{(S.388)}$$

where we use the identity in Eq. (S.291). Similarly,

$$\left\lfloor \Psi_k\,\Delta^{(0)}\Psi_k\right\rceil_{\ell,m} = \sum_{\ell',m'}\sum_{\ell'',m''} \frac{\ell'(\ell'+1)}{R^2}\,\Psi^k_{\ell',m'}\, \Psi^k_{\ell'',m''} \times \\ \times (-1)^{m+1}\, \mathcal{W}\,(\ell', m'|\ell'', m''|\ell, -m). \quad \text{(S.389)}$$

The expansion of $\Delta^{(0)}\big[(\Psi_k)^2\big]$ term in Eq. (S.382) can be computed as follows:

$$\left\lfloor \Delta^{(0)}\big[(\Psi_k)^2\big]\right\rceil_{\ell,m} = -\frac{\ell(\ell+1)}{R^2}\,\left\lfloor(\Psi_k)^2\right\rceil_{\ell,m} \quad \text{(S.390)}$$

which can be further expanded as

$$\left\lfloor \Delta^{(0)}\big[(\Psi_k)^2\big]\right\rceil_{\ell,m} = \frac{\ell(\ell+1)}{R^2}\sum_{\ell',m'}\sum_{\ell'',m''} \Psi^k_{\ell',m'}\,\Psi^k_{\ell'',m''} \times \\ \times (-1)^{m+1}\, \mathcal{W}\,(\ell', m'|\ell'', m''|\ell, -m). \quad \text{(S.391)}$$

Analogous expressions can be derived for the mass fraction $\Psi_0$, as those in Eqs. (S.388),(S.389), and (S.391).

Lastly, we rewrite the convective term in Eq. (S.382) by using the boundary condition in Eq. (S.241), as well as the expressions of the fluid velocity in Eq. (S.368) and the slip velocity in Eq. (S.244), which yields:

$$\left\lfloor \frac{\boldsymbol{v}^{(1)}\cdot r\boldsymbol{\nabla}\Psi_k}{R}\right\rceil_{\ell,m} = \sum_{\ell',m'}\sum_{\ell'',m''} \frac{\big[\frac{\partial}{\partial t}u_{\ell',m'} - \frac{1}{2}\mathbb{M}^{(1)}_{\ell',m'}\big]2\Psi^k_{\ell'',m''}}{\ell'(\ell'+1)} \times \\ \times \left\lfloor \boldsymbol{\Theta}_{\ell',m'}(\theta,\varphi)\cdot\boldsymbol{\Theta}_{\ell'',m''}(\theta,\varphi)\right\rceil_{\ell,m} \quad \text{(S.392)}$$

where the vector harmonics $\boldsymbol{\Theta}_{\ell,m}(\theta,\varphi) = r\boldsymbol{\nabla}Y_{\ell,m}(\theta,\varphi)$, as previously defined, whilst the projection term

$$\left\lfloor \boldsymbol{\Theta}_{\ell_1,m_1}\cdot\boldsymbol{\Theta}_{\ell_2,m_2}\right\rceil_{\ell,m} = (-1)^m\,\mathcal{W}(\ell_1,m_1|\ell_2,m_2|\ell,-m)\times \\ \times \tfrac{1}{2}\left[\ell_1(\ell_1+1)+\ell_2(\ell_2+1)-\ell(\ell+1)\right], \quad \text{(S.393)}$$

which is determined by an integration-by-parts method. Here, $\mathbb{M}^{(1)}_{\ell,m} = \sum_k \mathbb{M}^k_{\circlearrowleft}\Psi^k_{\ell,m}$ is the total membrane mass flux-rate, and $\frac{\partial}{\partial t}u_{\ell,m}$ is the rate of change of the shape distortion, which is given by Eq. (S.336) at first order in the perturbation, associated with each of $\ell$ and $m$ modes. We neglect the nonlinearities corresponding to the shape equation; thus, preserving only the weak nonlinear terms associated with the membrane compositional dynamics of the fissogens and fusogens.

As a result, the dynamics of each spherical harmonics $\frac{\partial}{\partial t}\Psi^k_{\ell,m}$ can be determined as an infinite summation over all of the other modes, with the coefficients expressed in terms of the Wigner $3j$-symbols. As mentioned earlier, the series is truncated to the second mode, which allows us to find a complete set of coupled differential equations. In addition, we consider, for simplicity, only the $m = 0$ mode that is associated with $\ell = 0$, 1, and 2, which in effect corresponds to choosing the same direction or axis for each of the modes. Hence, we have six nonlinear differential equations to solve for the composition, and three linear equations for the shape. However, we seek for solutions that conserve the total mass of the system; namely, the integrated excess membrane mass must be

$$\int_{\mathcal{M}} \mathbb{M}\,\mathrm{d}S = 0. \quad \text{(S.394)}$$

This condition can be expanded to second-order in the perturbation parameter $\varepsilon$, as follows:

$$\iint \mathrm{d}\varphi\,\mathrm{d}\theta\,\Big[\mathbb{M}^{(1)}\,[\sqrt{g}]^{(0)} + \mathbb{M}^{(1)}[\sqrt{g}]^{(1)}\Big] = 0, \quad \text{(S.395)}$$

where $\big[\sqrt{g}\big]^{(0)} = R^2\sin\theta$, and $\big[\sqrt{g}\big]^{(1)} = 2R^2u(\theta,\varphi)\sin\theta$. By using the orthogonality condition, we find that

$$\mathbb{M}^{(1)}_{0,0} + \frac{1}{\sqrt{\pi}}\sum_{\ell,m} u^*_{\ell,m}\,\mathbb{M}^{(1)}_{\ell,m} = 0, \quad \text{(S.396)}$$

with $u^*_{\ell,m} = (-1)^m\,u_{\ell,-m}$. Again, this leads to an infinite series over the surface modes, which we truncate here to

$\ell = 2$, and set $m = 0$ for all $\ell$-th modes. For consistency, we also must require that

$$\frac{\mathrm{d}}{\mathrm{d}t}\int_{\mathcal{M}} \mathbb{M}\,\mathrm{d}S = 0, \tag{S.397}$$

which follows in general from the conservation of all membrane components, cf. Eqs. (S.6) and (S.10). Hence,

$$\frac{\partial}{\partial t}\mathbb{M}^{(1)}_{0,0} + \frac{1}{\sqrt{\pi}}\frac{\partial}{\partial t}\sum_{\ell,m} u^{*}_{\ell,m}\,\mathbb{M}^{(1)}_{\ell,m} = 0. \tag{S.398}$$

These conditions guarantee that the overall membrane mass of the system remains constant and does not change as a function of time, despite the local addition and removal of membrane via the vesicular trafficking fluxes.

#### C. Dynamics of the centre-of-mass velocity

The centre-of-mass velocity $\boldsymbol{V}^{(1)}_{\mathrm{cm}}$ at first-order in the perturbation is linearly related to $\frac{\partial}{\partial t}u_{\ell=1,m}$, as shown in Eq. (S.352). To obtain the second-order contribution in $\varepsilon$ for the centre-of-mass velocity, we make use the following identity [46]: $\frac{\mathrm{d}}{\mathrm{d}t}\iiint \mathrm{d}V\boldsymbol{R} = \iint \mathrm{d}S\; v\boldsymbol{R}$, which follows from Reynolds transport theorem, with $v$ being the normal velocity of the membrane that encloses a volume $V$. Therefore, we have that

$$\frac{\mathrm{d}}{\mathrm{d}t}(\boldsymbol{R}_{\mathrm{cm}}V) = \iint \mathrm{d}S\; v\boldsymbol{R}, \tag{S.399}$$

which can be readily expanded to quadratic order in $\varepsilon$. The second-order term is found to be

$$\begin{aligned}\boldsymbol{V}^{(2)}_{\mathrm{cm}}\,V^{(0)} + \boldsymbol{V}^{(1)}_{\mathrm{cm}}\,V^{(1)} + \boldsymbol{R}^{(1)}_{\mathrm{cm}}\frac{\mathrm{d}}{\mathrm{d}t}V^{(1)} &= \\ = \iint \mathrm{d}\varphi\mathrm{d}\theta\left([\sqrt{g}]^{(0)}v^{(1)}\boldsymbol{R}^{(1)} + [\sqrt{g}]^{(1)}v^{(1)}\boldsymbol{R}^{(0)}\right)&,\end{aligned} \tag{S.400}$$

with $V^{(0)} = 4\pi R^3/3$ and $V^{(1)} = 4\pi R^3\,u_{0,0}\,Y_{0,0}$. Note that $v^{(0)} = 0$ and also $v^{(2)} = 0$. Since $v^{(1)} = R\frac{\partial}{\partial t}u(\theta,\varphi)$, and position vectors $\boldsymbol{R}^{(0)} = R\hat{\boldsymbol{r}}$ and $\boldsymbol{R}^{(1)} = Ru(\theta,\varphi)\hat{\boldsymbol{r}}$, then

$$\begin{aligned}\boldsymbol{V}^{(2)}_{\mathrm{cm}} = &-\frac{3u_{0,0}}{\sqrt{4\pi}}\,\boldsymbol{V}^{(1)}_{\mathrm{cm}} - \frac{3\frac{\partial}{\partial t}u_{0,0}}{\sqrt{4\pi}}\boldsymbol{R}^{(1)}_{\mathrm{cm}} \\ &+ \frac{9R}{4\pi}\iint \mathrm{d}\varphi\,\mathrm{d}\theta\sin\theta\;\hat{\boldsymbol{r}}\,u(\theta,\varphi)\frac{\partial u(\theta,\varphi)}{\partial t}.\end{aligned} \tag{S.401}$$

By writing $u(\theta,\varphi) = \sum_{\ell,m} u_{\ell,m}Y_{\ell,m}(\theta,\varphi)$, and truncating to the second $\ell$-th mode, the centre of mass velocity is found to be

$$\boldsymbol{V}^{(2)}_{\mathrm{cm}} = \frac{3R\sqrt{15}}{10\pi}\left(u_{1,0}\frac{\partial u_{2,0}}{\partial t} + u_{2,0}\frac{\partial u_{1,0}}{\partial t}\right)\hat{\boldsymbol{z}} + \cdots \tag{S.402}$$

where $m = 0$, and the dots refer to a summation over the other associated $m$-th modes.

Therefore, the weak nonlinear form of the velocity of the centre of mass, for $m = 0$, is given by

$$\boxed{V_{\mathrm{cm}} = \frac{R\sqrt{3}}{2\sqrt{\pi}}\frac{\partial}{\partial t}(u_{1,0}) + \frac{3R\sqrt{15}}{10\pi}\frac{\partial}{\partial t}(u_{1,0}\,u_{2,0}),} \tag{S.403}$$

which corresponds to the $z$-component of the velocity of the membrane compartment. By using the expression of $\frac{\partial}{\partial t}u_{\ell,m}$ from Eq. (S.336), the nonlinear form of $V_{\mathrm{cm}}$ can be obtained in terms of the mass fractions $\Psi^k_{\ell,m}$. Hence, a dynamical equation for the centre of mass velocity can be found by evaluating $\frac{\partial}{\partial t}V_{\mathrm{cm}}$. Thus, given the nonlinear differential equations for the membrane composition $\Psi^k_{\ell,m}$ from Eq. (S.384), and the constraints from Eqs. (S.396) and (S.398) due to net membrane mass conservation, we determine the fixed points of the dynamical system by solving for

$$\frac{\partial \Psi^k_{\ell,m}}{\partial t} = 0 \quad \text{and} \quad \frac{\partial V_{\mathrm{cm}}}{\partial t} = 0, \tag{S.404}$$

where $\ell = 0$, 1, and 2, and every $m = 0$. We denote the fixed points for the mass fractions by $\bar{\Psi}^k_{\ell,m}$ and the fixed point of the membrane centre of mass velocity by $\bar{V}_{\mathrm{cm}}$. Here, the stationary points are computed by a homotopy continuation method [47], which allows us to find numerically all real roots of the algebraic equations in (S.404). Note that we do not demand $\frac{\partial}{\partial t}u_{\ell,m} = 0$, the shape distortions being allowed to change in order to accommodate the mass constraints in Eqs. (S.396) and (S.398), as well as the form of $\frac{\partial}{\partial t}V_{\mathrm{cm}} = 0$ at the fixed point. We denote the stationary values of the shape deviations by $\bar{u}_{\ell,m}$.

Once these roots are numerically determined, we study the linear stability of the fixed points by determining the eigenvalues of the Jacobin matrix, corresponding to the dynamical system, at each of the resolved fixed points. Namely, we perform a linear perturbation in the membrane compositional fields, $\Psi^k_{\ell,m} = \bar{\Psi}^k_{\ell,m} + \delta\Psi^k_{\ell,m}$, and the centre of mass velocity, $V_{\mathrm{cm}} = \bar{V}_{\mathrm{cm}} + \delta V_{\mathrm{cm}}$, where $\delta\Psi^k_{\ell,m}$ is a small perturbation in the mass fractions, whereas $\delta V_{\mathrm{cm}}$ is the corresponding perturbation in the velocity that is self-consistently obtained through the shape distortions $u_{\ell,m} = \bar{u}_{\ell,m} + \delta u_{\ell,m}$, with $\delta u_{\ell,m}$ and $\frac{\partial}{\partial t}\delta u_{\ell,m}$ respecting the constrains due to the overall mass conservation.

This shows us that the homogeneous steady-state is not only linearly unstable, but can also be quadratically unstable, which means that a small perturbation in any of the compositional modes (other than $\ell = 1$) could drive the drift instability of the membrane compartment.

#### D. Bifurcation diagrams and phase portraits

By fixing the mass fraction $\bar{\Phi}_0$, and the parameters $\zeta$, $A$, and $Q$, as defined in Eqs. (S.342) and (S.344), we find stable solutions for the centre of mass velocity $\bar{V}_{\mathrm{cm}} \neq 0$. Note that these non-trivial real solutions always come as a pair of positive and negative values for $\bar{V}_{\mathrm{cm}}$ with the same magnitude. The stationary stable values of the mass fractions $\bar{\Psi}^k_{\ell,m}$ can be used to find the fixed points of the corresponding mass fluxes $\bar{\mathbb{M}}^{(1)}_{\ell,m} = \sum_k \mathbb{M}^k_{\circlearrowleft} \bar{\Psi}^k_{\ell,m}$ of each spherical harmonic mode. The dynamics of these mass fluxes, $\frac{\partial}{\partial t}\mathbb{M}^{(1)}_{\ell,m}$, can be also determined. Hence, the phase portraits of the velocity $V_{\rm cm}$ versus $\frac{\partial}{\partial t}\mathbb{M}^{(1)}_{\ell,m}$ can be visualized, if we allow two of the mass fractions, say $\Psi^{k=1}_{\ell=1}$ and $\Psi^{k=2}_{\ell=1}$, to vary, and the the remaining compositional fields to be pinned at their fixed point value $\bar{\Psi}^k_{\ell,m}$ where the mode $\ell = 0$ and 2 (with $m = 0$). Hence, this gives us a two-dimensional projection of the steamlines around the stable fixed point, as plotted in Fig. S.18. Only the flows around the positive fixed value of $V_{\rm cm}$ are depicted; similar flows can be found for the negative solution.

Another way to visualize the dynamics of the system near its stable fix points is by solving the initial value problem of these coupled differential equations, starting with $V_{\rm cm}(t = 0) = 0$ and choosing the other mass fractions such that the orbit (or trajectory) lies within the basin of attraction of the fixed point [45]. This is illustrated in Fig. S.20, which shows the velocity $V_{\rm cm}$ and the active membrane fluxes $\mathbb{M}^{(1)}_{\ell=1,2}$ as function of time, with the latter being scaled by the diffusion time $R^2/\gamma$.

Furthermore, the product of the centre of mass velocity $V_{\rm cm}$ and the mass flux $\mathbb{M}^{(1)}_{\ell=1}$, which we denote hereinafter by $\Xi$, provides us with an order parameter which differentiates between stable retrograde drift ($\Xi > 0$) and the stable anterograde motion ($\Xi < 0$). In the example shown in Fig S.20, the dynamical variable $\Xi(t)$ saturates to a positive value, which means that the membrane compartment drifts in a retrograde fashion.

The fixed points of $\Xi$ and $V_{\rm cm}$ change as we vary the active parameters $A$ and $\zeta$. This is shown in the bifurcation diagrams in Fig. (S.19), where we trace the value of the fixed points as we vary $\zeta$, at fixed $A$ and $Q$. The values of the latter are chosen so that the order parameter $\Xi$ shows a sign switch at some critical value of $\zeta$, depicted in Fig. (S.19) by the third vertical line (around $\zeta \approx 3.25$). This is the transition point between retrograde motion and anterograde drift of the membrane compartment. This phase transition happens smoothly, meaning that $\Xi = 0$ is a second-order phase boundary. The second vertical dashed line in Fig. (S.19) corresponds to the linearly unstable boundary as given by Eq. (S.346), where the homogeneous steady-state (with velocity $V_{\rm cm} = 0$ and vanishing mass fluxes) is no longer a stable solution, as shown by the green line. This also indicates that at this point we have a subcritical bifurcation, with the equilibrium solution becoming unstable, and thus the system makes a finite jump to the nearest attractor. The first vertical line in Fig. (S.19) shows the initial point at which the nontrivial solutions become stable, representing at a line of metastability. In other words, the homogeneous steady-state can coexist with the other (nonlinear) stable solutions within a region near the linearly unstable line.

These different phase boundaries can be visualized by using a phase diagram as a function of $\zeta$ and $A$, at fixed value of $Q$, as illustrated in Fig. S.21. The stability is found only for large negative values of $Q$; here, we make use of the convention that $Q_1 = Q_2 = Q$, where $Q_{1,2}$ are defined in Eq. (S.344). A region of stable retrograde and anterograde solutions is found, as depicted in Fig. S.21 by the yellow and blue regions, respectively. A nonlinearly unstable domain is also found at large $A$, which is shown by the red regions in Fig. S.21. As increasing $Q$ (from a large negative value), this region of instability tends to cover a larger area of the phase space, which ultimately leads to the complete absence of any stable fixed points for $Q > 0$. All of these regions are qualitatively the same for the other convention in which $Q_1 = -Q_2 = Q$, where the stable solutions are only found for negative values of $Q$, and an absence of stability for $Q > 0$.

The linearly unstable boundary is a first-order line, at which the system shows a discontinuity in the dynamical variables. This line is given by $A = A^\star_{\ell=1}(\zeta)$ with $A^\star_{\ell=1}$ being defined in Eq. (S.346). By setting the parameter $A$ to be on the unstable line, the variation of $\Xi$ with $\zeta$ and $Q$ can be obtained as shown in Fig. (S.17). This shows that $\Xi$ displays a maximum value, above which the system is unstable. From its maximum, $\Xi$ decreases as a function of $\zeta$, vanishing at some critical value of $\zeta_c$, which corresponds to the phase transition from a retrograde drift to an anterograde motion. As the points are chosen to lie to the unstable line $A = A^\star_{\ell=1}(\zeta)$, the point $\zeta = \zeta_c$ is a tricritical point, with a first order line intersecting a second-order line. Thus, the curve $\zeta_c(Q)$ at which $\Xi = 0$ describes a tricritical line.

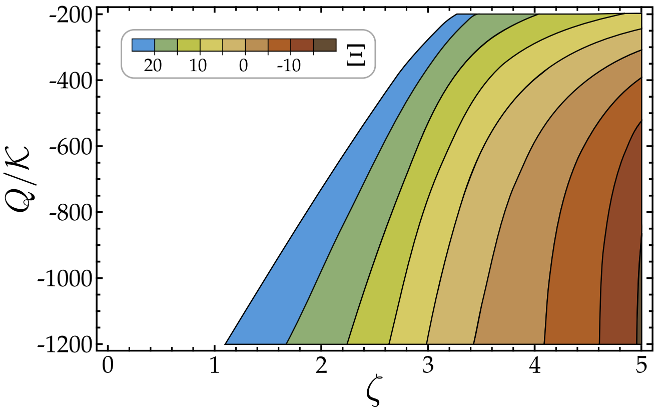


FIG. S.17. Phase plot of the order parameter $\Xi$, which is the product between $V_{\rm cm}$ and the local mass flux $\mathbb{M}^{(1)}_{\ell=1}$ associated with the first spherical harmonic mode. The value of $A$ and $\zeta$ are set to be on the linearly unstable line, namely $A = A^\star_{\ell=1}(\zeta)$, see main text. $\Xi$ shows a maximum (blue contour region), which varies as a function of $Q$. In the white region, we find only unstable fixed points. Note that $\Xi$ switches sign at value of $\zeta = \zeta_c(Q)$. This represents a tricitical line, where the linearly unstable line meets with the second order line $\Xi = 0$.

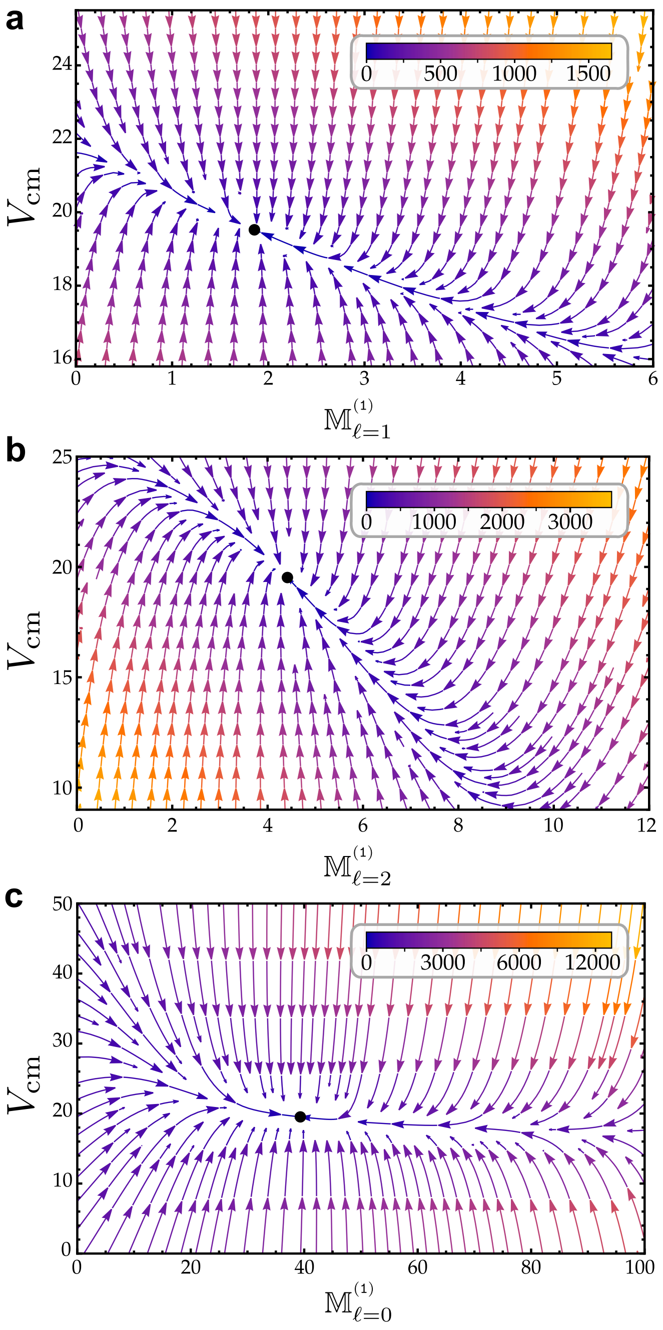


FIG. S.18. Streamline plots of the dynamical variables, showing the centre of mass velocity $V_{\rm cm}$ versus the local, net membrane flux $\mathbb{M}^{(1)}_{\ell} = \sum_k \mathbb{M}^k_{\circlearrowleft} \Psi^k_{\ell}$ which are associated with the spherical harmonic modes $\ell = 0$, 1, and 2. Here, the streamlines represent the flows of $\frac{\partial}{\partial t} V_{\rm cm}$ and $\frac{\partial}{\partial t} \mathbb{M}^{(1)}_{\ell}$, with only $\Psi^k_{\ell=1}$ being allowed to vary, and keeping the other mass fractions at their fixed point values. The color represents the magnitude of the flow fields, whereas the black points are the stable stationary points of the system, with parameters $\zeta = 2$, $A = 250$, $Q = -8000$, $\mathcal{K} = 10$, and $\bar{\Phi}_0 = 3/5$. Note that the centre of mass velocity is measured in units of $\gamma/R$, whilst the rates $\mathbb{M}^{(1)}_{\ell}$ are all non-dimensionalized by the diffusion time $R^2/\gamma$. This shows only the stream flows in the neighborhood around the positive fixed point of the velocity. A similar flow profile can be found around the negative velocity fixed point.

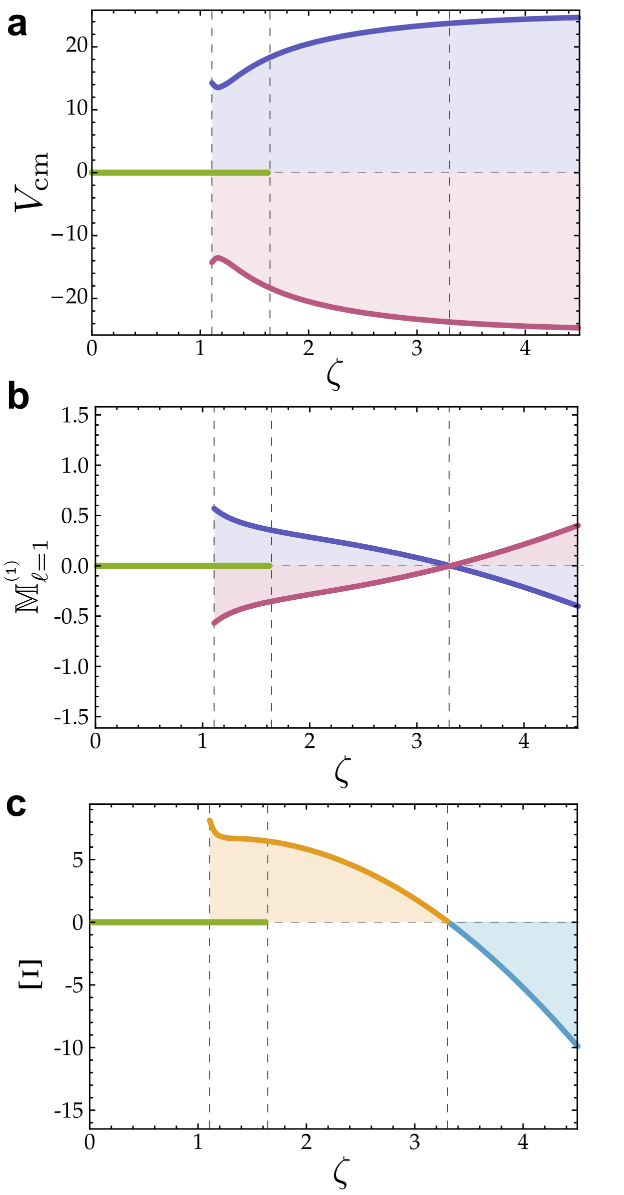


FIG. S.19. Bifurcation diagrams as a function of the parameter $\zeta$ of the fixed point values of (a) the velocity $V_{\rm cm}$, (b) the mass flux $\mathbb{M}^{(1)}_{\ell=1}$, and (c) the order parameter $\Xi$, which is the product of the previous two dynamical variables. The blue and purple curves in (a) represent the two stable branches of velocity, which can be positive or negative. Similarly, for subfigure (b), which in this case switches sign at a particular value of $\zeta$. The product of $V_{\rm cm}$ and $\mathbb{M}^{(1)}_{\ell=1}$ merges these two branches into one curve that we denote by $\Xi$, which is colored in orange for $\Xi > 0$ (retrograde motion) and is shown in blue for $\Xi < 0$ (anterograde motion). The green line shows the stable homogeneous steady-state solutions, with a stationary centre-of-mass velocity and a vanishing mass flux. The first vertical dashed line indicates the initial point at which the nonlinear solution becomes stable, while the vertical second dashed line gives the point of linear instability at which the green line ends. Lastly, the third vertical line indicates the transition point between the retrograde drift to the anterograde motion of the membrane compartment. Here, we choose that $A = 85$, $Q = -8000$, $\mathcal{K} = 10$, and $\bar{\Phi}_0 = 3/5$.

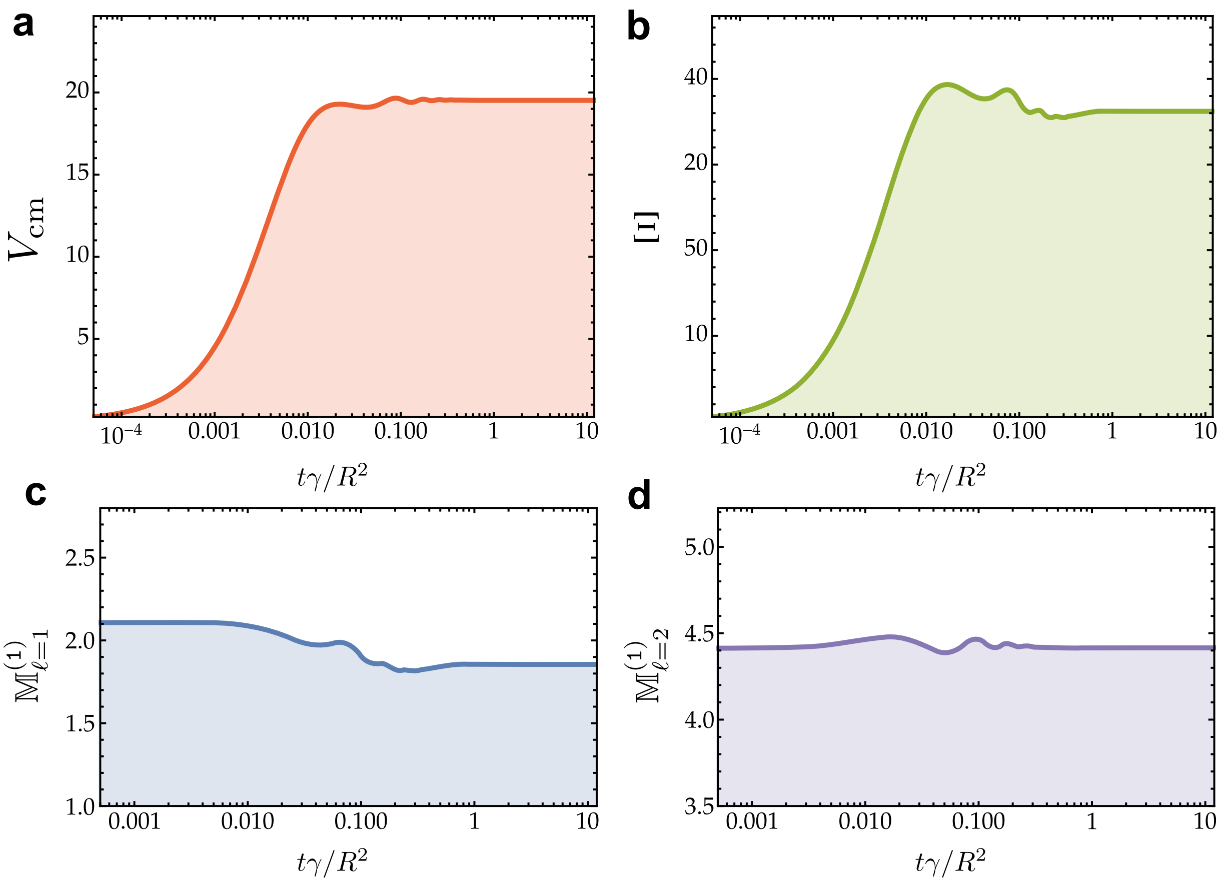


FIG. S.20. Temporal evolution of the dynamical variables: (a) the centre of mass velocity $V_{\rm cm}(t)$; (b) the order parameter $\Xi(t)$ which is given by the product $V_{\rm cm}\,\mathbb{M}^{(1)}_{\ell=1}$; (c) the mass flux $\mathbb{M}^{(1)}_{\ell=1}(t)$, associated with the first spherical harmonic mode; and (d) the mass flux $\mathbb{M}^{(1)}_{\ell=2}(t)$ corresponding to the second spherical harmonic mode. The fluxes $\mathbb{M}^{(1)}_{\ell=1,2}$ are rescaled by $R^2/\gamma$, whilst the velocity $V_{\rm cm}$ is measured in units of $\gamma/R$. Here, $\zeta = 2$, $A = 250$, $Q = -8000$, $\mathcal{K} = 10$, and $\bar{\Phi}_0 = 3/5$.

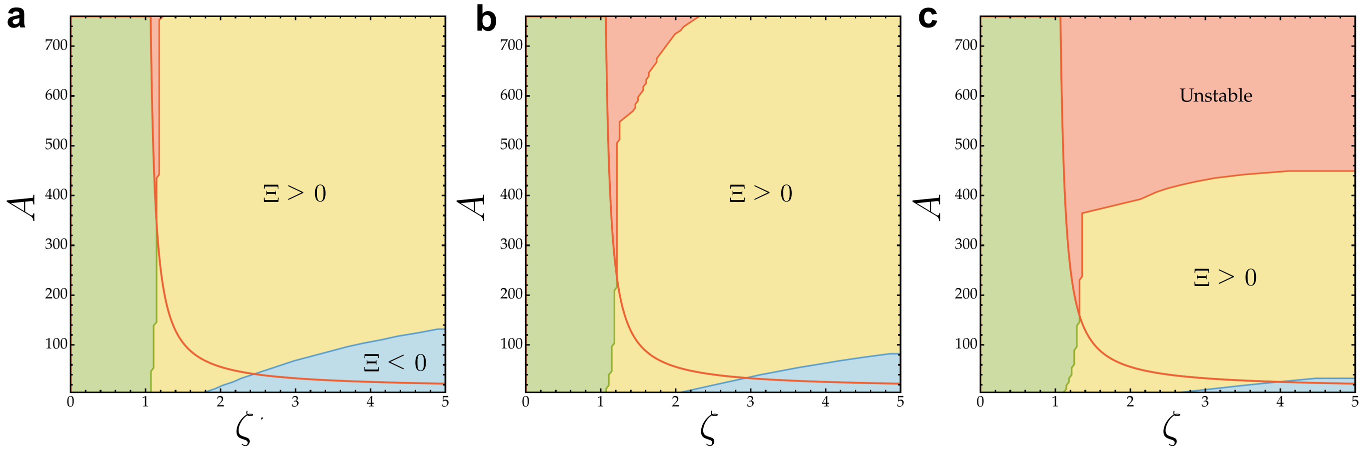


FIG. S.21. Phase diagrams as function of $\zeta$ and $m$ for three distinct $Q$-values: (a) $Q = -8000$; (b) $Q = -6000$; (c) $Q = -4000$. The mass fraction $\bar{\Phi}_0 = 3/5$ and the rescaled bending rigidity $\mathcal{K} = 10$. Yellow regions show the stable retrograde solutions with $\Xi > 0$, while the blue regions show the stable anterograde solutions with $\Xi < 0$. The green region lies below the red curve (that is the linearly unstable boundary), and depicts the region in which the homogeneous steady-state solution ($V_{\rm cm} = 0$) is stable. Note that the yellow and blue regions also superimpose on top of the green region for $\zeta > 1$, which means that stable retrograde and anterograde motion can coexist with the stationary state $V_{\rm cm} = 0$. The red regions are the unstable domains, where none of the fixed points are stable; meaning that the weak nonlinearities truncated up to the second spherical harmonic mode are not sufficient to stabilize the motion of the membrane compartment for a particular triplet $(A, \zeta, Q)$.

### IV. MATHEMATICAL PREREQUISITES

#### A. Differential Geometry of Surfaces

We briefly outline the mathematical language of differential geometry of two-dimensional surfaces, concentrating on the practical aspects of the subject rather than mathematical rigour. This brief review assumes that the reader is acquainted with some basic knowledge of vector and tensor calculus. It is noteworthy to mention that an extensive literature exists on differential geometry, with emphasis on both pure mathematics (e.g. Ref. [48]) and applications in physics (e.g. Refs. [17] and [49]).

***Surface parametrization*** — A two-dimensional surface, say $\mathcal{S}$, embedded in the three-dimensional space $\mathbb{R}^3$ can be uniquely determined by a three-dimensional vector $\boldsymbol{R} = (X, Y, Z)^{\mathsf{T}}$, where $X$, $Y$, and $Z$ are the Cartesian coordinates, and $\mathsf{T}$ denotes a transpose [21]. However, these coordinates are not independent, but they satisfy a condition which dictates the precise form of the two-dimensional surface (in other words, the choice of two coordinates gives exactly the value of the remaining one).

Therefore, the embedding surface function $\boldsymbol{R}$ can be described by a pair of two variables, denoted by $(\xi^1, \xi^2)$, which represent the internal *curvilinear* coordinates associated with a parametrization of the surface. At each point on the surface, a basis can be formed by constructing the tangent vectors and their corresponding normal vector, as shown in Fig. S.22. The tangent vector associated with the internal coordinate $\xi^\alpha$ is defined by

$$\boldsymbol{e}_\alpha = \frac{\partial \boldsymbol{R}}{\partial \xi^\alpha} \equiv \partial_\alpha \boldsymbol{R}, \tag{S.405}$$

with index $\alpha = 1, 2$, whilst the unit vector $\boldsymbol{n}$ normal to the surface (*i.e.* $\boldsymbol{n} \cdot \boldsymbol{e}_\alpha = 0$ and $\boldsymbol{n} \cdot \boldsymbol{n} = 1$) is given by

$$\boldsymbol{n} = (\boldsymbol{e}_1 \times \boldsymbol{e}_2) / \|\boldsymbol{e}_1 \times \boldsymbol{e}_2\|, \tag{S.406}$$

where $\times$ is the three-dimensional cross product, and $\|\cdot\|$ gives the norm of the vector. The vectors $\boldsymbol{e}_1$ and $\boldsymbol{e}_2$ are not orthonormal in general, namely the four elements of

$$g_{\alpha\beta} = \boldsymbol{e}_\alpha \cdot \boldsymbol{e}_\beta, \tag{S.407}$$

known as the *metric tensor*, when written in matrix form, typically give $[g_{\alpha\beta}] \neq \begin{bmatrix} 1 & 0 \\ 0 & 1 \end{bmatrix}$. Nevertheless, the condition of orthonormality can be restored by constructing a dual set of tangent vectors, say $\boldsymbol{e}^1$ and $\boldsymbol{e}^2$, which satisfy

$$\boldsymbol{e}_\alpha \cdot \boldsymbol{e}^\beta = \delta_\alpha^\beta, \tag{S.408}$$

where $\delta_\alpha^\beta$ is the Kronecker delta, *i.e.* $[\delta_\alpha^\beta] = \begin{bmatrix} 1 & 0 \\ 0 & 1 \end{bmatrix}$. Note that metric tensor is a symmetric tensor, *i.e.* $g_{\alpha\beta} = g_{\beta\alpha}$. Also, the inverse of the metric tensor, $g^{\alpha\beta}$, satisfies

$$g^{\alpha\beta} g_{\beta\gamma} = \delta_\gamma^\alpha, \tag{S.409}$$

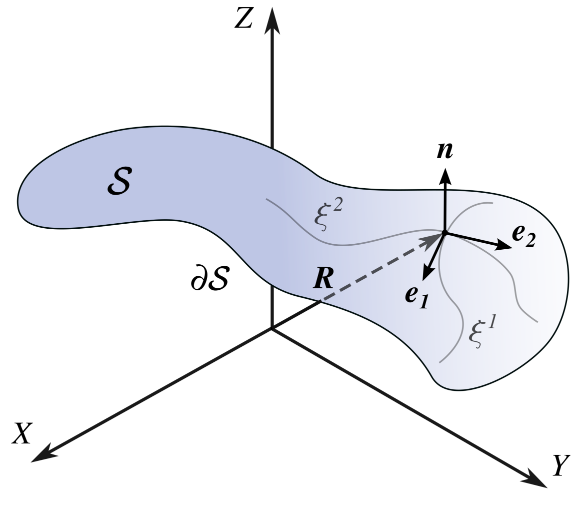


FIG. S.22. Schematic diagram of a two-dimensional surface $\mathcal{S}$, whose boundary is denoted by $\partial\mathcal{S}$. Every point on the surface can be identified by a three-dimensional vector $\boldsymbol{R}$ (described by the Cartesian coordinates $X$, $Y$ and $Z$). Also, for each vector $\boldsymbol{R}$, we define the tangent vectors $\boldsymbol{e}_1$ and $\boldsymbol{e}_2$, corresponding to the internal coordinates $\xi^1$ and $\xi^2$, respectively, as well as the unit vector $\boldsymbol{n}$ normal to the tangent plane.

where Einstein convention is employed, *i.e.* summation over repeated super- and sub-script indices is implied. It follows that $g_{\alpha\beta}$ and $g^{\alpha\beta}$ can be used to lower and raise indices, respectively; for example,

$$\boldsymbol{e}^\alpha = g^{\alpha\beta} \boldsymbol{e}_\beta, \quad \text{and} \quad \boldsymbol{e}_\alpha = g_{\alpha\beta} \boldsymbol{e}^\beta, \tag{S.410}$$

The metric tensor captures the distance information on the surface, being an intrinsic geometrical property of that surface, and defines the scalar product on $\mathcal{S}$; namely, for two vectors $\boldsymbol{v} = v^\alpha \boldsymbol{e}_\alpha$ and $\boldsymbol{w} = w^\alpha \boldsymbol{e}_\alpha$, we have that $\boldsymbol{v} \cdot \boldsymbol{w} = (v^\alpha \boldsymbol{e}_\alpha) \cdot (w^\beta \boldsymbol{e}_\beta) = g_{\alpha\beta} v^\alpha w^\beta = v_\alpha w^\alpha = v^\alpha w_\alpha$.

***Surface decomposition*** — Both triads $\{\boldsymbol{e}_1, \boldsymbol{e}_2, \boldsymbol{n}\}$ and $\{\boldsymbol{e}^1, \boldsymbol{e}^2, \boldsymbol{n}\}$ can be used as bases to decompose any three-dimensional vector $\boldsymbol{v}$ into its normal and in-plane vectorial parts, namely $\boldsymbol{v} = \boldsymbol{v}_{\boldsymbol{n}} + \boldsymbol{v}_0$, where $\boldsymbol{v}_{\boldsymbol{n}} = v\,\boldsymbol{n}$ and $\boldsymbol{v}_0 = v^\alpha \boldsymbol{e}_\alpha = v_\alpha \boldsymbol{e}^\alpha$, in which we define the normal vector component by $v = \boldsymbol{v} \cdot \boldsymbol{n}$, the co- and contra-variant vector components by $v_\alpha = \boldsymbol{v} \cdot \boldsymbol{e}_\alpha$ and $v^\alpha = \boldsymbol{v} \cdot \boldsymbol{e}^\alpha$, respectively. From (S.409) and (S.410), we find that the metric tensor and its inverse can be used to raise and lower the index of vector components, *i.e.* $v^\alpha = g^{\alpha\beta} v_\beta$ and $v_\alpha = g_{\alpha\beta} v^\beta$.

Furthermore, we define the surface identity tensor,

$$\boldsymbol{I}_{\mathcal{S}} = \boldsymbol{e}_\alpha \otimes \boldsymbol{e}^\alpha = \boldsymbol{e}^\alpha \otimes \boldsymbol{e}_\alpha, \tag{S.411}$$

where $\otimes$ is the tensor product. This second-order tensor acts as projection operator onto the tangent plane of the surface, extracting the in-plane vector $\boldsymbol{v}_0$ from $\boldsymbol{v}$ (that is, $\boldsymbol{v}_0 = \boldsymbol{I}_{\mathcal{S}} \cdot \boldsymbol{v} = \boldsymbol{v} \cdot \boldsymbol{I}_{\mathcal{S}}$). Similarly, we define the three-dimensional identity tensor $\boldsymbol{I}$ as follows:

$$\boldsymbol{I} = \boldsymbol{I}_{\mathcal{S}} + \boldsymbol{n} \otimes \boldsymbol{n}, \tag{S.412}$$

which satisfies that $\boldsymbol{I} \cdot \boldsymbol{v} = \boldsymbol{v} \cdot \boldsymbol{I} = \boldsymbol{v}$ for any vector $\boldsymbol{v}$.

The surface identity can also be used to extract the surface part of any tensor; for instance, the in-plane content of a second-order tensor $\boldsymbol{T} \in \mathbb{R}^3 \times \mathbb{R}^3$ is given by $\boldsymbol{T}_{\mathcal{S}} = (\boldsymbol{I}_{\mathcal{S}} \cdot \boldsymbol{T})\, \boldsymbol{I}_{\mathcal{S}}$, which in index notation is written as

$$\begin{aligned} \boldsymbol{T}_{\mathcal{S}} &= T_{\alpha\beta}\, \boldsymbol{e}^\alpha \otimes \boldsymbol{e}^\beta, \quad T_{\alpha\beta} = (\boldsymbol{e}_\alpha \cdot \boldsymbol{T}) \cdot \boldsymbol{e}_\beta \\ &= T^\alpha_{\ \beta}\, \boldsymbol{e}_\alpha \otimes \boldsymbol{e}^\beta, \quad T^\alpha_{\ \beta} = (\boldsymbol{e}^\alpha \cdot \boldsymbol{T}) \cdot \boldsymbol{e}_\beta \\ &= T_\alpha^{\ \beta}\, \boldsymbol{e}^\alpha \otimes \boldsymbol{e}_\beta, \quad T_\alpha^{\ \beta} = (\boldsymbol{e}_\alpha \cdot \boldsymbol{T}) \cdot \boldsymbol{e}^\beta \\ &= T^{\alpha\beta}\, \boldsymbol{e}_\alpha \otimes \boldsymbol{e}_\beta, \quad T^{\alpha\beta} = (\boldsymbol{e}^\alpha \cdot \boldsymbol{T}) \cdot \boldsymbol{e}^\beta. \end{aligned} \tag{S.413}$$

Notice that the order of upper and lower indices of the tensor components generally does matter; however, if $\boldsymbol{T}$ is symmetric ($T_{\alpha\beta} = T_{\beta\alpha}$), then we have the property that $T_\alpha^{\ \beta} = T^\alpha_{\ \beta} \equiv T^\alpha_\beta$. In a similar fashion, the normal component of the tensor $\boldsymbol{T}$ can be written as

$$\boldsymbol{T}_{\boldsymbol{n}} = T\, \boldsymbol{n} \otimes \boldsymbol{n}, \quad T = (\boldsymbol{n} \cdot \boldsymbol{T}) \cdot \boldsymbol{n}. \tag{S.414}$$

In addition to these, a second-order tensor may also have in general components along $\boldsymbol{n} \otimes \boldsymbol{e}_\alpha$ and $\boldsymbol{e}_\alpha \otimes \boldsymbol{n}$.

***Surface differentiation*** — In tensor calculus, the derivative introduced in (S.405) is known as the parametric derivative, and is typically denoted by a comma (that is, we write $\partial_\alpha \boldsymbol{R} = \boldsymbol{R}_{,\alpha}$). By differentiating again Eq. (S.405) with respect to $\xi^\beta$, this yields the parametric derivative $\boldsymbol{e}_{\alpha,\beta} = \boldsymbol{R}_{,\alpha\beta}$, which in general could have both normal and in-plane components. As only the former is needed to understand the notion of surface curvature (heuristically, curvature can be understood as instantaneous rate of change of the normal vector as one moves along a path on the surface), we write the normal component of the derivative $\boldsymbol{e}_{\alpha,\beta}$ by using the semi-colon notation, *i.e.*

$$\boldsymbol{e}_{\alpha;\beta} = (\boldsymbol{n} \otimes \boldsymbol{n})\, \boldsymbol{e}_{\alpha,\beta}. \tag{S.415}$$

This is known as a covariant derivative, and by using the equations (S.411) and (S.412), we retrieve the common form that is usually introduced in the literature:

$$\boldsymbol{e}_{\alpha;\beta} = \boldsymbol{e}_{\alpha,\beta} - \Gamma^\gamma_{\alpha\beta} \boldsymbol{e}_\gamma, \tag{S.416}$$

where $\Gamma^\gamma_{\alpha\beta} = \boldsymbol{e}^\gamma \cdot \boldsymbol{e}_{\alpha,\beta}$ are the *Christoffel symbols* of the second kind, which can be also rewritten in terms of the derivatives of the metric tensor, namely

$$\Gamma^\gamma_{\alpha\beta} = \frac{g^{\gamma\delta}}{2} \left(g_{\alpha\delta,\beta} + g_{\delta\beta,\alpha} - g_{\alpha\beta,\delta}\right), \tag{S.417}$$

which shows the symmetry property that $\Gamma^\gamma_{\alpha\beta} = \Gamma^\gamma_{\beta\alpha}$.

Similarly, for the basis vector $\boldsymbol{e}^\alpha$, its covariant derivative is $\boldsymbol{e}^\alpha_{;\beta} = (\boldsymbol{n} \otimes \boldsymbol{n})\, \boldsymbol{e}^\alpha_{,\beta}$. By (S.411) and (S.412), we find

$$\boldsymbol{e}^\alpha_{;\beta} = \boldsymbol{e}^\alpha_{,\beta} + \Gamma^\alpha_{\beta\gamma} \boldsymbol{e}^\gamma. \tag{S.418}$$

The covariant derivative for scalars $w \in \mathbb{R}$ is equivalent to the parametric derivative, *i.e.* $w_{;\alpha} = w_{,\alpha}$. This is also true for three-dimensional vectors; namely, for $\boldsymbol{v} \in \mathbb{R}^3$, we have $(v^\alpha \boldsymbol{e}_\alpha)_{;\beta} = \boldsymbol{v}_{;\beta} = \boldsymbol{v}_{,\beta} = (v^\alpha \boldsymbol{e}_\alpha)_{,\beta}$. On the other hand, the covariant derivatives of the co- and contravariant components of a vector $\boldsymbol{v}$ are not the same as their comma derivative, and they are given by

$$v_{\alpha;\beta} = v_{\alpha,\beta} - \Gamma^\gamma_{\alpha\beta} v_\gamma, \quad \text{and} \quad v^\alpha_{;\beta} = v^\alpha_{,\beta} + \Gamma^\alpha_{\beta\gamma} v^\gamma. \tag{S.419}$$

A useful property of the covariant derivative is the fact that the metric tensor and its inverse is constant with respect to the covariant derivative, *i.e.*

$$g_{\alpha\beta;\gamma} = g^{\alpha\beta}_{\ \ \ ;\gamma} = 0, \tag{S.420}$$

which is known as *metrinilic property*. In practice, this means that the process of raising and lowering of indices commutes with the covariant differentiation.

Although the covariant differentiation has restored the tensorial structure of the (parametric) derivative $\boldsymbol{e}_{\alpha,\beta}$, a resulting drawback is the loss of commutativity, namely

$$v_{\gamma;\alpha\beta} - v_{\gamma;\beta\alpha} = v^\delta R_{\alpha\beta\gamma\delta}, \tag{S.421}$$

where $R_{\alpha\beta\gamma\delta}$ is the *Riemann curvature tensor*. This is, in general, a nonzero tensor, vanishing only for locally flat surfaces, and its components are given by

$$R_{\alpha\beta\gamma\delta} = \Gamma_{\beta\delta\alpha,\gamma} - \Gamma_{\beta\gamma\alpha,\delta} + \Gamma^\omega_{\beta\gamma}\Gamma_{\alpha\delta\omega} - \Gamma^\omega_{\beta\delta}\Gamma_{\alpha\gamma\omega}, \tag{S.422}$$

where $\Gamma_{\gamma\alpha\beta} = g_{\gamma\delta}\Gamma^\delta_{\ \alpha\beta}$ are the Christoffel symbols of the first kind. Since $R_{\alpha\beta\gamma\delta}$ depends only on the metric tensor, it is then an intrinsic quantity of the surface.

With the help of the covariant derivative, we can now define the surface analogous of the regular differential operators, encountered in vector calculus: gradients, divergences, and Laplacian operators. The surface gradient of a scalar function $\phi$ that lives on $\mathcal{S}$ is defined by

$$\nabla\phi = \boldsymbol{I}_{\mathcal{S}} \cdot \boldsymbol{\nabla}\phi = \phi_{,\alpha}\, \boldsymbol{e}^\alpha = \phi_{;\alpha}\, \boldsymbol{e}^\alpha, \tag{S.423}$$

where $\boldsymbol{\nabla}$ is the usual gradient operator in $\mathbb{R}^3$. Similarly, the gradient of a vector function $\boldsymbol{\Phi}$ can be written as follows $\nabla\boldsymbol{\Phi} = \boldsymbol{I}_{\mathcal{S}} \cdot \boldsymbol{\nabla}\boldsymbol{\Phi} = \boldsymbol{\Phi}_{,\alpha} \otimes \boldsymbol{e}^\alpha$. As a corollary, this allows us to define the surface divergence of $\boldsymbol{\Phi}$, namely

$$\operatorname{div}\boldsymbol{\Phi} = \nabla\boldsymbol{\Phi} : \boldsymbol{I} = \boldsymbol{\Phi}_{,\alpha} \cdot \boldsymbol{e}^\alpha = \boldsymbol{\Phi}_{;\alpha} \cdot \boldsymbol{e}^\alpha, \tag{S.424}$$

where $:$ denotes the double dot product. Let $\boldsymbol{\Phi}$ be solely defined on $\mathcal{S}$, with $\boldsymbol{\Phi} = \phi^\alpha \boldsymbol{e}_\alpha$, then Eq. (S.424) can be written in component form as follows:

$$\operatorname{div}\boldsymbol{\Phi} = \phi^\alpha_{;\alpha} = \partial_\alpha(\sqrt{g}\,\phi^\alpha)/\sqrt{g}\,, \tag{S.425}$$

where the former follows from (S.416) and (S.417), while the latter is a result known as the *Voss–Weyl formula*, in which we define that

$$g = \det[g_{\alpha\beta}]. \tag{S.426}$$

Note that $g$ is also covariantly invariant, *i.e.* $g_{;\alpha} = 0$, due to the metrinilic property. The surface Laplacian for a scalar function $\phi$ is given by

$$\Delta\phi = \operatorname{div}\nabla\phi = g^{\alpha\beta}\phi_{;\alpha\beta}. \tag{S.427}$$

It is noteworthy to stress that $\phi_{;\alpha\beta} \neq \phi_{,\alpha\beta}$ in general, as

$$\phi_{;\alpha\beta} = \phi_{,\alpha\beta} - \Gamma^{\gamma}_{\alpha\beta}\,\phi_{,\gamma}. \tag{S.428}$$

Lastly, we note that the most effective way to compute the surface Laplacian of a scalar function is via the Voss–Weyl formula, which allows us to rewrite (S.427) as

$$\Delta\phi = \partial_\alpha\big(\sqrt{g}\,g^{\alpha\beta}\phi_{,\beta}\big)\,/\sqrt{g}. \tag{S.429}$$

***Surface curvature*** — As previously mentioned, the curvature of a surface can be described via the normal component of the second (parametric) derivatives of $\boldsymbol{R}$. Consequently, a second-order tensor can be constructed as $\boldsymbol{B} = b_{\alpha\beta}\,\boldsymbol{e}^\alpha \otimes \boldsymbol{e}^\beta$, which is known as the curvature tensor, and its covariant elements $b_{\alpha\beta}$ are defined by

$$b_{\alpha\beta} = \boldsymbol{n}\cdot\boldsymbol{R}_{,\alpha\beta} = \boldsymbol{n}\cdot\boldsymbol{e}_{\alpha,\beta} = \boldsymbol{n}\cdot\boldsymbol{e}_{\alpha;\beta} \tag{S.430}$$

which is a symmetric tensor, and the sign of $b_{\alpha\beta}$ depends on the choice of the normal. The mixed components of $\boldsymbol{B}$ are thus given by $b^{\beta}_{\alpha} = b_{\alpha\gamma}\,g^{\gamma\beta}$, whereas its contravariant elements are found to be $b^{\alpha\beta} = b^{\alpha}_{\gamma}\,g^{\gamma\beta}$. This tensor measures the extrinsic curvature, *i.e.* the bending of the two-dimensional surface in the three-dimensional space. The connection between intrinsic and extrinsic geometry is captured by the *Gauss–Weingarten equations*, *i.e.*

$$\boldsymbol{e}_{\alpha;\beta} = b_{\alpha\beta}\,\boldsymbol{n}, \quad \text{and} \quad \boldsymbol{n}_{;\alpha} = -b^{\beta}_{\alpha}\,\boldsymbol{e}_\beta, \tag{S.431}$$

where the former equation follows from the definition of curvature tensor and covariant differentiation, while the latter is derived by differentiating the identities $\boldsymbol{n}\cdot\boldsymbol{e}_\alpha = 0$ and $\boldsymbol{n}\cdot\boldsymbol{n} = 1$, and then solving for the derivative $\boldsymbol{n}_{;\alpha}$.

The eigenvalues of the matrix $[b^{\alpha}_{\beta}]$ are commonly called the *principal curvatures*, and their associated eigenvectors yield the *principal directions*. Instead of working with the principal curvatures, we construct two equivalent scalar invariants: the mean curvature $H$, and the Gaussian curvature $K$, which are defined by

$$H = \frac{1}{2}\mathrm{tr}\,[b^{\alpha}_{\beta}] = \frac{1}{2}b^{\alpha}_{\alpha}, \quad \text{and} \tag{S.432}$$

$$K = \det[b^{\alpha}_{\beta}], \tag{S.433}$$

respectively. The Gaussian curvature can also be written as $K = \det[b_{\alpha\beta}]\,/\det[g_{\alpha\beta}]$, or in index notation,

$$K = \frac{1}{2}\varepsilon^{\alpha\beta}\varepsilon^{\gamma\delta}b_{\alpha\gamma}\,b_{\beta\delta}, \tag{S.434}$$

where the permutation tensor $\varepsilon^{\alpha\beta}$ is defined by vanishing diagonal terms, *i.e.* $\varepsilon^{11} = \varepsilon^{22} = 0$, while its off-diagonal components are given by $\varepsilon^{12} = -\varepsilon^{21} = 1/\sqrt{g}$. From Eq. (S.434), we can see that the tensor $\varepsilon^{\alpha\beta}\varepsilon^{\gamma\delta}b_{\beta\delta} = \bar{b}^{\alpha\gamma}$ acts as the cofactor of the curvature tensor, which can be rewritten as follows:

$$\bar{b}^{\alpha\beta} = \varepsilon^{\alpha\gamma}\varepsilon^{\beta\delta}b_{\gamma\delta} = 2Hg^{\alpha\beta} - b^{\alpha\beta}, \tag{S.435}$$

resulting in $\bar{b}^{\alpha\gamma}b_{\gamma\beta} = K\delta^{\alpha}_{\beta}$. This gives us yet another way to write the Gaussian curvature, *i.e.* $K = 2H^2 - \frac{1}{2}b^{\alpha}_{\beta}b^{\beta}_{\alpha}$, which leads to the following useful formula:

$$b_{\alpha\beta}\,b^{\alpha\beta} = b^{\alpha}_{\beta}\,b^{\beta}_{\alpha} = 4H^2 - 2K. \tag{S.436}$$

In equation (S.425), the domain of definition of the vector field $\boldsymbol{\Phi}$ has been restricted to the tangent plane of the surface. However, Eqs. (S.431) and (S.432) provide now the tools to extend this to vector fields in $\mathbb{R}^3$. By using the definition of the surface divergence, as in (S.424), for a vector $\boldsymbol{v} = v^\alpha\,\boldsymbol{e}_\alpha + v\,\boldsymbol{n}$, then $\mathrm{div}(\boldsymbol{v})$ is found to be

$$\mathrm{div}(\boldsymbol{v}) = v^{\alpha}_{;\alpha} - 2Hv. \tag{S.437}$$

We note that the tensors $g_{\alpha\beta}$ and $b_{\alpha\beta}$ are not independent (in other words, not all choices of tensor fields $g_{\alpha\beta}$ and $b_{\alpha\beta}$ can describe a surface). This can be seen as an integrability condition on the Gauss–Weingarten equations, namely one needs to make sure that the identity

$$\boldsymbol{e}_{\gamma,\alpha\beta} = \boldsymbol{e}_{\gamma,\beta\alpha} \tag{S.438}$$

is satisfied. Thus, the normal and tangential components of (S.438) leads to *Gauss–Codazzi–Mainardi equations*:

$$b_{\beta\gamma;\alpha} - b_{\alpha\gamma;\beta} = 0, \quad \text{and} \tag{S.439}$$

$$b_{\alpha\gamma}b_{\beta\delta} - b_{\alpha\delta}b_{\beta\gamma} = R_{\alpha\beta\gamma\delta}. \tag{S.440}$$

The latter equation tells us that the combination given by $b_{\alpha\gamma}b_{\beta\delta} - b_{\alpha\delta}b_{\beta\gamma}$ is an intrinsic quantity of the surface, which can be purely computed from the metric (as $R_{\alpha\beta\gamma\delta}$ depends only on the metric tensor), despite the fact that the curvature tensor $b_{\alpha\beta}$ itself is an extrinsic quantity that requires an embedding. Moreover, by contracting twice the Riemann tensor, we obtain a scalar quantity known as the Ricci scalar, *i.e.* $R = g^{\alpha\gamma}g^{\beta\delta}R_{\alpha\beta\gamma\delta}$. By using Eq. (S.440), we have $R = 4H^2 - b^{\alpha}_{\beta}b^{\beta}_{\alpha} = 2K$, where the last equality follows from Eq. (S.436), and thus the Gaussian curvature is an intrinsic geometrical quantity. In fact, for two-dimensional surfaces, $K$ is the only independent component of the Riemann tensor, as

$$R_{\alpha\beta\gamma\delta} = K\,(g_{\alpha\gamma}\,g_{\beta\delta} - g_{\alpha\delta}\,g_{\beta\gamma}). \tag{S.441}$$

On the other hand, Eq. (S.439), which is often called the *Codazzi equation*, states that $b_{\alpha\gamma;\beta}$ must be symmetric with respect to the indices $\alpha$ and $\beta$. Since the curvature tensor $b_{\alpha\beta}$ is also symmetric, then we find that $b_{\alpha\gamma;\beta}$ is fully symmetric with respect to all its indices.

#### B. Calculus of Moving Surfaces

The formulation of kinetic and kinematic relationships on a deformable membrane is much easier achieved by parameterizing the surface through a convected coordinate system $\varrho^\alpha$ with respect to a fixed reference system $\xi^\alpha$, where the latter is depicted in Fig. S.22. Specifically, we identify by $\hat{\boldsymbol{R}}(\varrho^\alpha, t)$ the current position at time $t$ of a material point that was located at

$$\hat{\boldsymbol{R}}(\varrho^\alpha, t_0) = \boldsymbol{R}(\xi^\alpha), \tag{S.442}$$

with the initial (reference) time $t_0 \leq t$. In other words, we specify the fixed coordinates $\xi^\alpha$ as functions which depend on both the time and the convected coordinates $\varrho^\alpha$, subject to the condition that

$$\varrho^\alpha = \xi^\alpha(\varrho^\beta, t_0). \tag{S.443}$$

In addition, we assume that these mappings between $\varrho^\alpha$ and $\xi^\alpha$ are invertible, reflecting the notion that, at any fixed time $t$, a material point that is identified by constant $\varrho^\alpha$ can be uniquely associated with the local surface coordinates $\xi^\alpha$ (at time $t_0$). Thus, we write

$$\hat{\boldsymbol{R}}(\varrho^\alpha, t) = \boldsymbol{R}\big(\xi^\beta(\varrho^\alpha, t), t\big). \tag{S.444}$$

The velocity $\boldsymbol{v}$ of a material point that lives solely on the surface is defined by the rate of change of the position vector $\hat{\boldsymbol{R}}(\varrho^\alpha, t)$ at fixed $\varrho^\alpha$, namely

$$\boldsymbol{v} = \left.\frac{\partial \hat{\boldsymbol{R}}}{\partial t}\right|_{\{\varrho^\beta\}} = \left.\frac{\partial \boldsymbol{R}}{\partial t}\right|_{\{\xi^\beta\}} + \boldsymbol{e}_\alpha \left.\frac{\partial \xi^\alpha}{\partial t}\right|_{\{\varrho^\beta\}}. \tag{S.445}$$

where the latter equality follows via the chain rule. Hereinafter, we define the tangential velocity components $v^\alpha$ as the rate of change of the fixed coordinates $\xi^\alpha$ for a given material point of constant $\varrho^\alpha$, that is,

$$v^\alpha = \left.\frac{\partial \xi^\alpha}{\partial t}\right|_{\{\varrho^\beta\}}. \tag{S.446}$$

On the other hand, the normal velocity is given by

$$v = \boldsymbol{n} \cdot \left.\frac{\partial \boldsymbol{R}}{\partial t}\right|_{\{\xi^\alpha\}} \tag{S.447}$$

so that the surface points of constant $\xi^\alpha$ change only along the normal direction to the surface, without any loss of generality. Thus, the velocity $\boldsymbol{v}$ can be written as

$$\boldsymbol{v} = v\boldsymbol{n} + v^\alpha \boldsymbol{e}_\alpha. \tag{S.448}$$

Any scalar function $\phi(\xi^\alpha, t)$ that lives on the surface can also be written as a function in terms of the convected coordinates, say $\hat{\phi}(\varrho^\alpha, t)$, where we have that

$$\hat{\phi}(\varrho^\alpha, t) = \phi\big(\xi^\beta(\varrho^\alpha, t), t\big). \tag{S.449}$$

The material derivative of $\phi$ is simply the rate of change of $\hat{\phi}$ for a given material point of constant $\varrho^\alpha$, that is,

$$\dot{\phi} = \frac{\mathrm{d}\phi}{\mathrm{d}t} = \left.\frac{\partial \hat{\phi}}{\partial t}\right|_{\{\varrho^\alpha\}} \tag{S.450}$$

where the *overdot* symbol denotes the material derivative of a function. By utilizing the chain rule, this leads to

$$\dot{\phi} = \left.\frac{\partial \phi}{\partial t}\right|_{\{\xi^\beta\}} + \phi_{,\alpha} \left.\frac{\partial \xi^\alpha}{\partial t}\right|_{\{\varrho^\beta\}} = \frac{\partial \phi}{\partial t} + v^\alpha \phi_{,\alpha}. \tag{S.451}$$

Similarly, the material derivative of the in-plane basis vectors $\boldsymbol{e}_\alpha$ that are defined in Eq. (S.405) can be expressed in terms of the convected coordinates, as follows:

$$\dot{\boldsymbol{e}}_\alpha = \frac{\partial}{\partial t}\left[\frac{\partial \boldsymbol{R}}{\partial \xi^\alpha}\right]_{\{\varrho^\beta\}}. \tag{S.452}$$

This could be further expressed in terms of $\hat{\boldsymbol{R}}(\varrho^\alpha, t)$ by using Eq. (S.444), and by chain rule we find

$$\begin{aligned} \dot{\boldsymbol{e}}_\alpha &= \frac{\partial}{\partial t}\left[\frac{\partial \varrho^\gamma}{\partial \xi^\alpha}\frac{\partial \hat{\boldsymbol{R}}}{\partial \varrho^\gamma}\right]_{\{\varrho^\beta\}} = \frac{\partial \varrho^\gamma}{\partial \xi^\alpha}\frac{\partial}{\partial \varrho^\gamma}\left[\frac{\partial \hat{\boldsymbol{R}}}{\partial t}\right]_{\{\varrho^\beta\}} \\ &= \frac{\partial \boldsymbol{v}}{\partial \varrho^\gamma}\frac{\partial \varrho^\gamma}{\partial \xi^\alpha} = \frac{\partial \boldsymbol{v}}{\partial \xi^\alpha} = \boldsymbol{v}_{,\alpha}. \end{aligned} \tag{S.453}$$

Moreover, this can be decomposed into the local surface basis, *i.e.* $\{\boldsymbol{n}, \boldsymbol{e}_\alpha\}$, which yields

$$\dot{\boldsymbol{e}}_\alpha = \boldsymbol{v}_{,\alpha} = \big(v^\beta_{;\alpha} - v\, b^\beta_\alpha\big)\boldsymbol{e}_\beta + \big(v^\beta b_{\alpha\beta} + v_{,\alpha}\big)\boldsymbol{n}, \tag{S.454}$$

by employing the form of the velocity field in Eq. (S.448), and by also using the Gauss–Weingarten equations.

By applying the material derivative onto the orthogonality condition $\boldsymbol{n} \cdot \boldsymbol{e}_\alpha = 0$, we have $\boldsymbol{e}_\alpha \cdot \dot{\boldsymbol{n}} = -\boldsymbol{n} \cdot \dot{\boldsymbol{e}}_\alpha$. As $\boldsymbol{n} \cdot \dot{\boldsymbol{n}} = 0$, the material derivative of the normal unit vector acts only within the tangent plane of the surface, and thus from Eq. (S.454) we obtain that

$$\dot{\boldsymbol{n}} = -\big(v^\beta b^\alpha_\beta + v^{,\alpha}\big)\boldsymbol{e}_\alpha. \tag{S.455}$$

The material derivatives in Eqs. (S.454) and (S.455) allows us to determine the expression of the acceleration,

$$\begin{aligned} \dot{\boldsymbol{v}} \;=\; & \left[\frac{\partial v^\alpha}{\partial t} - \big(v^\beta b^\alpha_\beta + v^{,\alpha}\big)v + \big(v^\alpha_{;\beta} - v\, b^\alpha_\beta\big)v^\beta\right]\boldsymbol{e}_\alpha \\ &+ \left[\frac{\partial v}{\partial t} + \big(v^\beta b_{\alpha\beta} + v_{,\alpha}\big)\, v^\alpha\right]\boldsymbol{n}, \end{aligned} \tag{S.456}$$

as well as the explicit forms of the material derivatives of the metric tensor $g_{\alpha\beta}$ and the curvature tensor $b_{\alpha\beta}$, *i.e.*

$$\dot{g}_{\alpha\beta} = v_{\beta;\alpha} + v_{\alpha;\beta} - 2v\, b_{\alpha\beta}, \quad \text{and} \tag{S.457}$$

$$\dot{b}_{\alpha\beta} = b_{\gamma\beta}\left(v^\gamma_{;\alpha} - v\, b^\gamma_\alpha\right) + \left(v^\gamma b_{\gamma\alpha} + v_{,\alpha}\right)_{;\beta}, \tag{S.458}$$

respectively. The above expressions have been derived by applying the product rule of the material derivative, and by expanding out the definitions of $g_{\alpha\beta}$ and $b_{\alpha\beta}$ in terms of the local tangent basis and/or the normal. For instance, we have that $\dot{g}_{\alpha\beta} = \boldsymbol{e}_\alpha \cdot \dot{\boldsymbol{e}}_\beta + \dot{\boldsymbol{e}}_\alpha \cdot \boldsymbol{e}_\beta$, and then by using Eq. (S.454), we obtain the expression in Eq. (S.457). In an analogous way, from Eq. (S.430), we can write that $\dot{b}_{\alpha\beta} = \boldsymbol{n} \cdot \dot{\boldsymbol{e}}_{\alpha,\beta} + \dot{\boldsymbol{n}} \cdot \boldsymbol{e}_{\alpha,\beta} = -\boldsymbol{n}_{,\beta} \cdot \dot{\boldsymbol{e}}_\alpha - \dot{\boldsymbol{n}}_{,\beta} \cdot \boldsymbol{e}_\alpha$, which readily leads to Eq. (S.458) after some tedious algebra, by substituting the results in Eq. (S.454) and (S.455).

#### C. Integral and Transport Theorems

The area element $\mathrm{d}S$ of a surface that is parameterized by the fixed curvilinear coordinates $(\xi^1, \xi^2)$ is given by

$$\mathrm{d}S = \mathrm{d}\xi^1 \mathrm{d}\xi^2 \sqrt{g}, \tag{S.459}$$

where $g = \det[g_{\alpha\beta}]$, as previously defined in Eq. (S.426). This allows us to compute the total area of a surface $\mathcal{M}$ by integrating over the fixed internal coordinates $\xi^\alpha$, *i.e.*

$$A = \int_{\mathcal{M}} \mathrm{d}S = \iint \mathrm{d}\xi^1 \mathrm{d}\xi^2 \sqrt{g}\,. \tag{S.460}$$

To determine the rate of change of area, we thus need to compute $\frac{\mathrm{d}}{\mathrm{d}t}\sqrt{g}$. We make use of Jacobi's formula that allows us to express the derivative of the determinant of the matrix $[g_{\alpha\beta}]$ in terms of its inverse, $[g_{\alpha\beta}]^{-1}$, as well as its derivative; namely, this reads

$$\frac{\mathrm{d}}{\mathrm{d}t}\det[g_{\alpha\beta}] = \det[g_{\alpha\beta}]\ \mathrm{tr}\left\{[g_{\alpha\gamma}]^{-1}\frac{\mathrm{d}}{\mathrm{d}t}[g_{\gamma\beta}]\right\}, \tag{S.461}$$

with $\mathrm{tr}[\bullet]$ denoting the trace of a matrix. In terms of our previous notation, this formula can be rewritten as

$$\frac{1}{\sqrt{g}}\frac{\mathrm{d}(\sqrt{g})}{\mathrm{d}t} = \frac{1}{2}\,g^{\alpha\beta}\dot{g}_{\alpha\beta} = v^\alpha_{;\alpha} - 2vH = \mathrm{div}(\boldsymbol{v}), \tag{S.462}$$

where the third and the forth equality follows by means of Eq. (S.457) and Eq. (S.437), respectively. Hence,

$$\frac{\mathrm{d}}{\mathrm{d}t}\sqrt{g} = \mathrm{div}(\boldsymbol{v})\sqrt{g}, \tag{S.463}$$

which implies that the rate of change of area is given by

$$\frac{\mathrm{d}A}{\mathrm{d}t} = \iint \mathrm{d}\xi^1 \mathrm{d}\xi^2\,\frac{\mathrm{d}}{\mathrm{d}t}\sqrt{g} = \int_{\mathcal{M}} \mathrm{div}(\boldsymbol{v})\ \mathrm{d}S. \tag{S.464}$$

The identity in Eq. (S.463) could also be used to derive the so-called *Reynolds transport theorem*, that is,

$$\frac{\mathrm{d}}{\mathrm{d}t}\int_{\mathcal{S}} \boldsymbol{T}\,\mathrm{d}S = \int_{\mathcal{S}} \mathrm{d}S\left[\boldsymbol{T}\,\mathrm{div}(\boldsymbol{v}) + \frac{\mathrm{d}\boldsymbol{T}}{\mathrm{d}t}\right], \tag{S.465}$$

where $\mathcal{S}$ is an arbitrary surface patch, and $\boldsymbol{T}$ can be any tensor field that is defined on the domain $\mathcal{S}$, *e.g.* surface densities, or velocity fields that live on a membrane.

Furthermore, the volume $V$ enclosed by an arbitrary closed (orientable) surface, say $\mathcal{M}$, can be obtained from the position vector $\boldsymbol{R}$ as follows:

$$V = \iiint \mathrm{d}V = \frac{1}{3}\int_{\mathcal{M}} (\boldsymbol{n}\cdot\boldsymbol{R})\,\mathrm{d}S, \tag{S.466}$$

where the latter is found by employing the vector calculus identity, $\boldsymbol{\nabla}\cdot\boldsymbol{r} = 3$, with $\boldsymbol{r}$ being any position vector, and lastly making use of the divergence theorem (by also noting that boundary of the volume $\partial V$ is trivially the same as the closed surface of the membrane $\mathcal{M}$).

To find the rate of change of volume, we can simply apply the Reynolds transport theorem, which yields

$$\frac{\mathrm{d}V}{\mathrm{d}t} = \frac{1}{3}\int_{\mathcal{M}} \Big\{v + \boldsymbol{R}\cdot[\dot{\boldsymbol{n}} + \boldsymbol{n}\,\mathrm{div}(\boldsymbol{v})]\Big\}\,\mathrm{d}S, \tag{S.467}$$

where the term in the square brackets can be expressed as: $\dot{\boldsymbol{n}} + \boldsymbol{n}\,\mathrm{div}(\boldsymbol{v}) = -\big(v^\beta b^\alpha_\beta + v^{,\alpha}\big)\boldsymbol{e}_\alpha + \big(v^\alpha_{;\alpha} - 2Hv\big)\boldsymbol{n}$. Notice that the term $\boldsymbol{R}\cdot(v^{,\alpha}\boldsymbol{e}_\alpha)$ can be written in terms of a total derivative, namely

$$\boldsymbol{R}\cdot(v^{,\alpha}\boldsymbol{e}_\alpha) = [v\,(\boldsymbol{e}_\alpha\cdot\boldsymbol{R})]^{,\alpha} - 2v(1 + H\boldsymbol{n}\cdot\boldsymbol{R}), \tag{S.468}$$

where Eq. (S.430) is used to derive the latter. Since $\mathcal{M}$ is a closed surface, we obtain that

$$\frac{\mathrm{d}V}{\mathrm{d}t} = \frac{1}{3}\int_{\mathcal{M}} \Big\{3v + \boldsymbol{R}\cdot\big[\boldsymbol{n}\,v^\alpha_{;\alpha} - v^\beta b^\alpha_\beta\boldsymbol{e}_\alpha\big]\Big\}\,\mathrm{d}S. \tag{S.469}$$

By using the Gauss–Weingarten equation, $\boldsymbol{n}_{;\beta} = -b^\alpha_\beta\boldsymbol{e}_\alpha$, the second term in the integrand of Eq. (S.469) becomes

$$\boldsymbol{R}\cdot\big[\boldsymbol{n}\,v^\alpha_{;\alpha} + v^\alpha\boldsymbol{n}_{;\alpha}\big] = [\boldsymbol{n}\cdot\boldsymbol{R}\,v^\alpha]_{;\alpha}, \tag{S.470}$$

which is a total derivative that vanishes identically by integrating over the closed surface. Therefore, this leads to the following simple (and geometrically intuitive) result:

$$\frac{\mathrm{d}V}{\mathrm{d}t} = \int_{\mathcal{M}} v\ \mathrm{d}S, \tag{S.471}$$

that is, the rate of change of the volume $V$ is identical to the integrated value of the normal velocity over the total area of the membrane surface.

[21] E. Kreyszig, Differential Geometry (Dover Pub., 1991).

[37] C. Pozrikidis, Boundary Integral and Singularity Methods for Linearized Viscous Flow (Cambridge University Press, 1992).

[38] S. Kim and S. J. Karrila, Microhydrodynamics: Principles and Selected Applications (Elsevier, 1991).

[39] M. Doi, Soft Matter Physics (Oxford University Press, 2013).

[40] L. D. Landau and E. M. Lifshitz, Statistical Physics, 3rd ed. (Butterworth-Heinemann, 1980).

[41] M. Staykova, M. Arroyo, M. Rahimi, and H. A. Stone, Confined bilayers passively regulate shape and stress, Phys. Rev. Lett. **110**, 028101 (2013).

[42] M. Abramowitz and I. Stegun, Handbook of Mathematical Functions (Dover Pub., 1965).

[43] J. Happel and H. Brenner, Low Reynolds number hydrodynamics (Springer, 1981).

[44] G. Racah, Theory of complex spectra. ii, Physical Review **62**, 438 (1942).

[45] J. Guckenheimer and P. Holmes, Nonlinear Oscillations, Dynamical Systems, and Bifurcations of Vector Fields, Vol. 42 (Springer New York, 1983).

[46] N. Yoshinaga, Spontaneous motion and deformation of a self-propelled droplet, Phys. Rev. E **89**, 012913 (2014).

[47] M. Kuno and J. D. Seader, Computing all real solutions

to systems of nonlinear equations with a global fixed-point homotopy, Industrial and Engineering Chemistry Research **27**, 1320 (1988).

[48] M. Spivak, A Comprehensive Introduction to Differential Geometry, 3rd ed. (Publish or Perish, 1999).

[49] R. Kamien, The geometry of soft materials: a primer, Rev. Mod. Phys. **74**, 953 (2002).
